## Supplementary Data 1 for "Systematic functional analysis of *Leishmania* protein kinases identifies regulators of differentiation or survival"

### AGC FAMILY

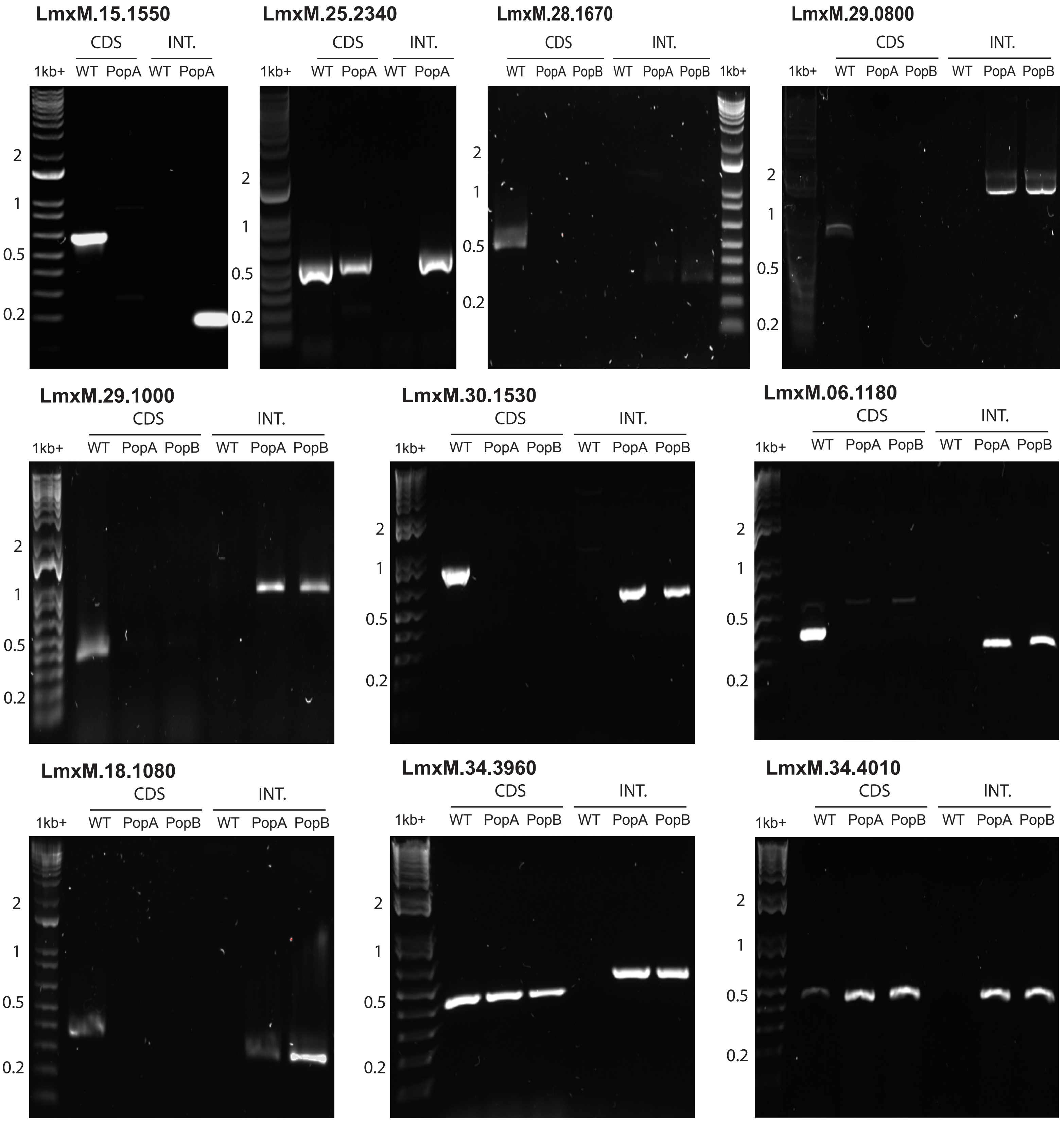
LmxM.15.1550
CDS INT.
1kb+ WT PopA WT PopA
2
1
0.5
0.2
LmxM.25.2340
CDS INT.
1kb+ WT PopA WT PopA
2
1
0.5
0.2
LmxM.28.1670
CDS INT.
WT PopA PopB WT PopA PopB 1kb+
2
1
0.5
0.2
LmxM.29.0800
CDS INT.
1kb+ WT PopA PopB WT PopA PopB
2
1
0.5
0.2
LmxM.29.1000
CDS INT.
1kb+ WT PopA PopB WT PopA PopB
2
1
0.5
0.2
LmxM.30.1530
CDS INT.
1kb+ WT PopA PopB WT PopA PopB
2
1
0.5
0.2
LmxM.06.1180
CDS INT.
1kb+ WT PopA PopB WT PopA PopB
2
1
0.5
0.2
LmxM.18.1080
CDS INT.
1kb+ WT PopA PopB WT PopA PopB
2
1
0.5
0.2
LmxM.34.3960
CDS INT.
1kb+ WT PopA PopB WT PopA PopB
2
1
0.5
0.2
LmxM.34.4010
CDS INT.
1kb+ WT PopA PopB WT PopA PopB
2
1
0.5
0.2

### CAMK FAMILY

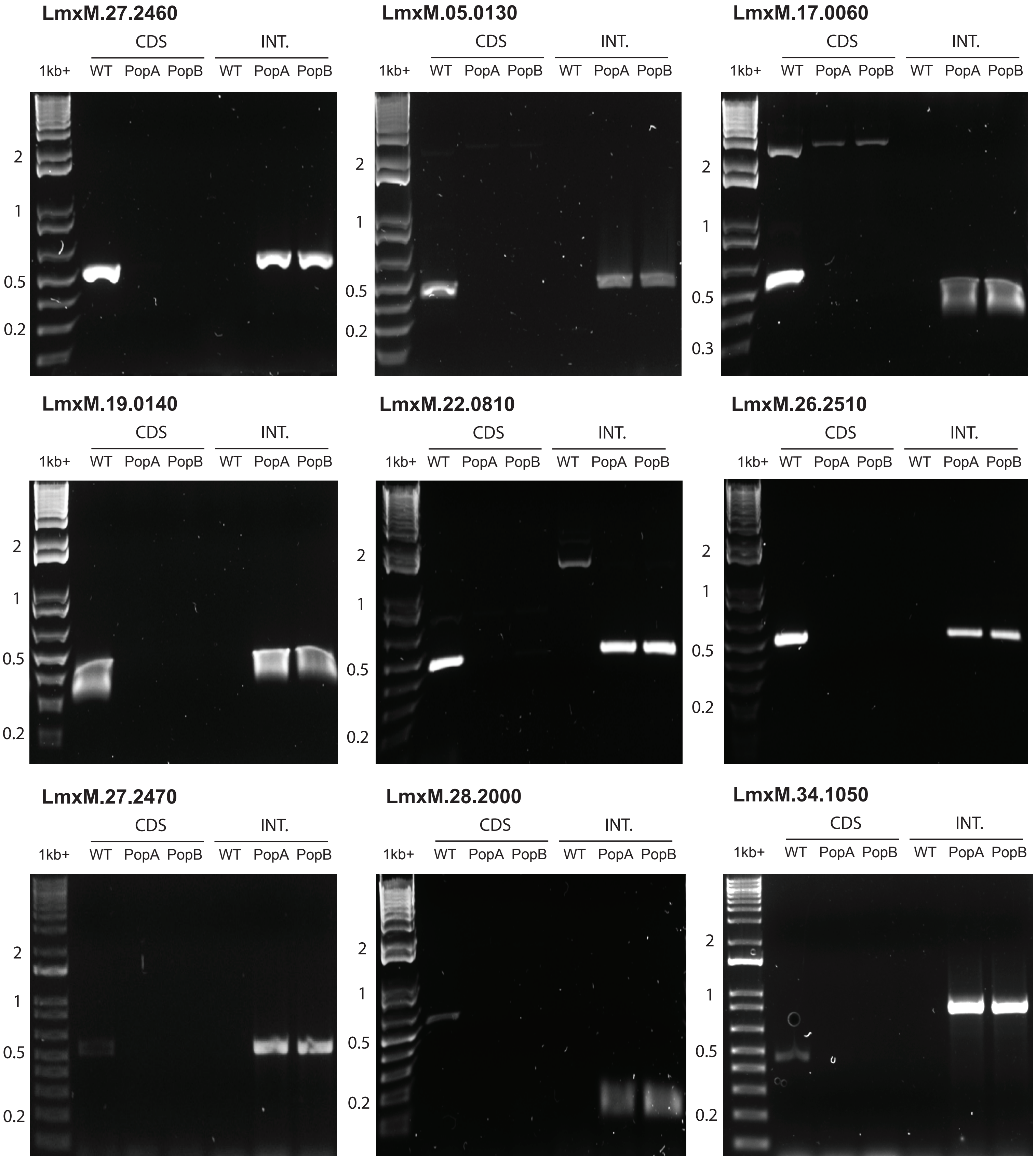
LmxM.27.2460
CDS
INT.
1kb+
WT PopA PopB WT PopA PopB
2
1
0.5
0.2
LmxM.05.0130
CDS
INT.
1kb+
WT PopA PopB WT PopA PopB
2
1
0.5
0.2
LmxM.17.0060
CDS
INT.
1kb+
WT PopA PopB WT PopA PopB
2
1
0.5
0.3
LmxM.19.0140
CDS
INT.
1kb+
WT PopA PopB WT PopA PopB
2
1
0.5
0.2
LmxM.22.0810
CDS
INT.
1kb+
WT PopA PopB WT PopA PopB
2
1
0.5
0.2
LmxM.26.2510
CDS
INT.
1kb+
WT PopA PopB WT PopA PopB
2
1
0.5
0.2
LmxM.27.2470
CDS
INT.
1kb+
WT PopA PopB WT PopA PopB
2
1
0.5
0.2
LmxM.28.2000
CDS
INT.
1kb+
WT PopA PopB WT PopA PopB
2
1
0.5
0.2
LmxM.34.1050
CDS
INT.
1kb+
WT PopA PopB WT PopA PopB
2
1
0.5
0.2

### CAMK FAMILY

##### LmxM.07.0900

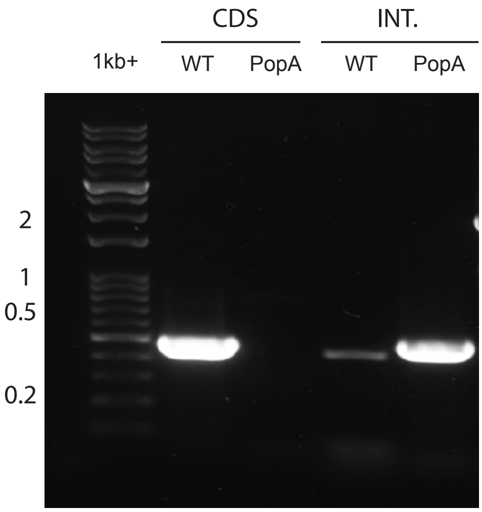


##### LmxM.18.0640

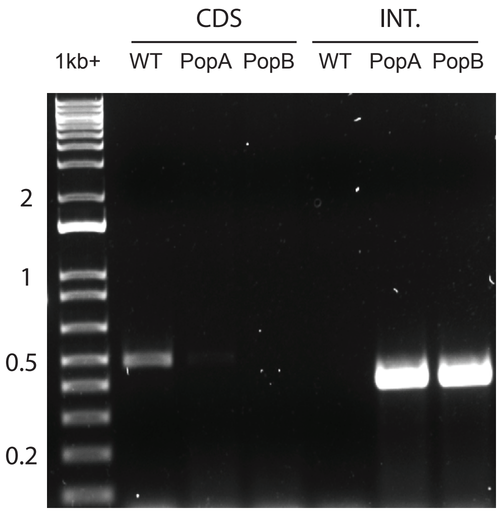


##### LmxM.24.0230

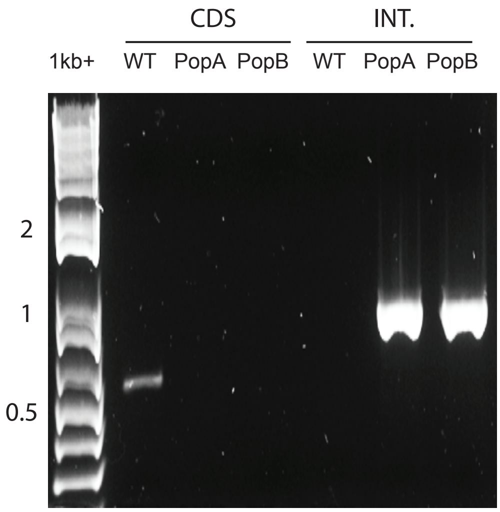


##### LmxM.08_29.2020

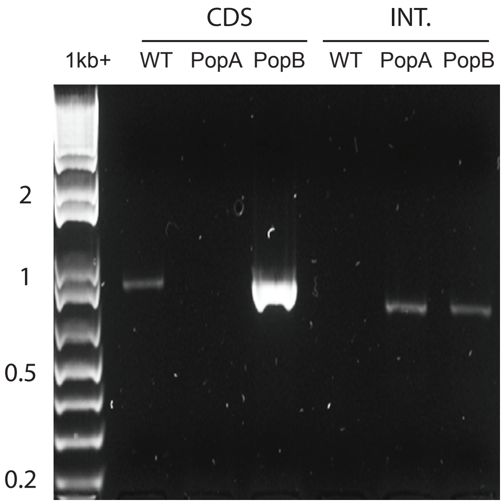


##### LmxM.32.1710

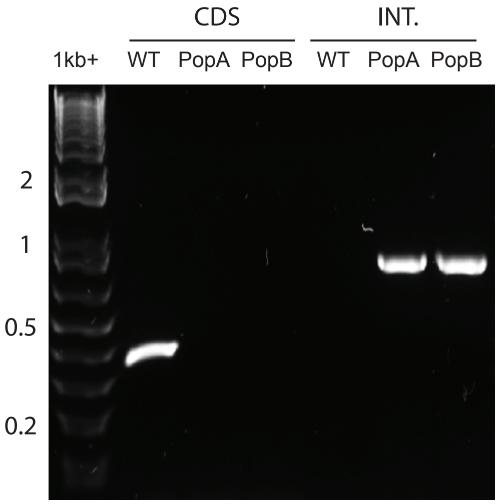


##### LmxM.34.0490

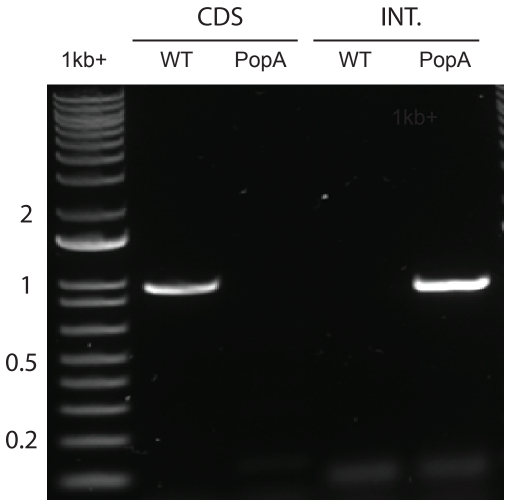


##### LmxM.36.0900

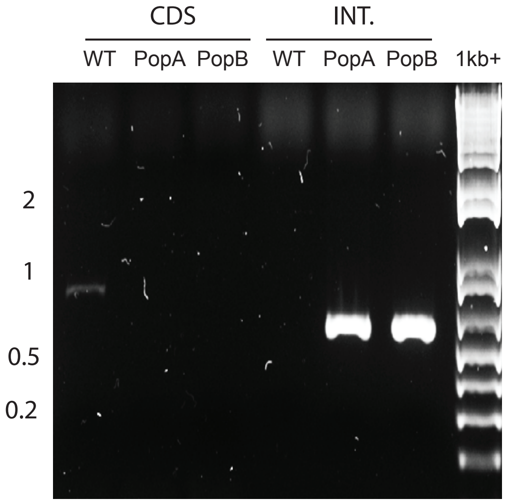


##### LmxM.21.0150

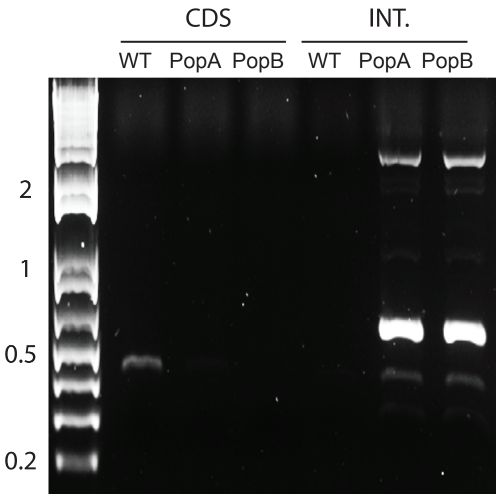


##### LmxM.22.1170

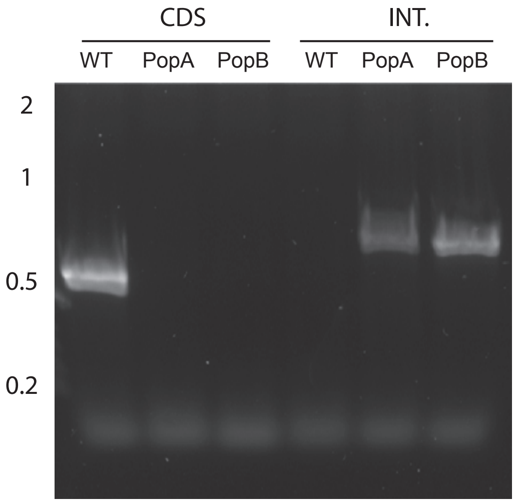

#### CAMK FAMILY

**LmxM.19.1470**

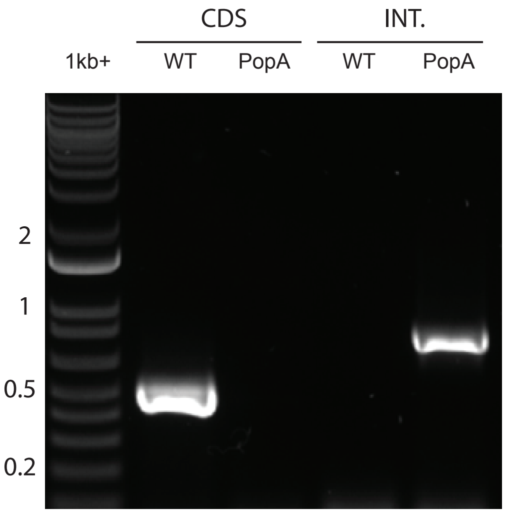


**LmxM.23.0490**

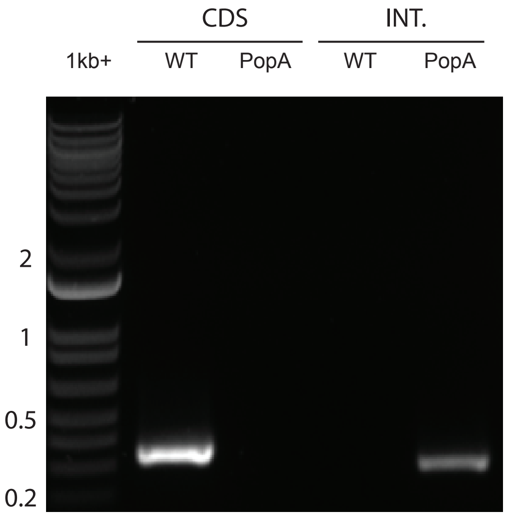


#### CK1 FAMILY

**LmxM.04.1210**

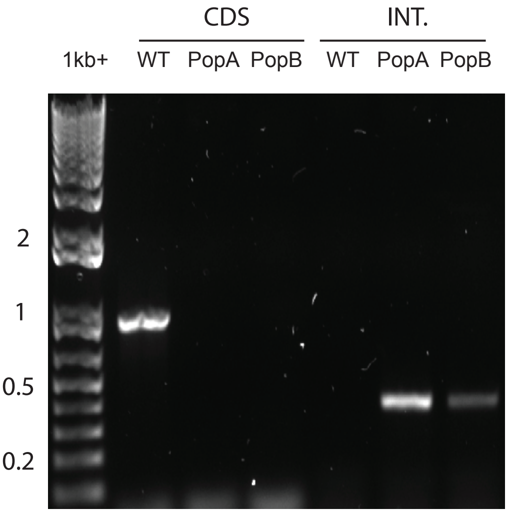


**LmxM.25.1580**

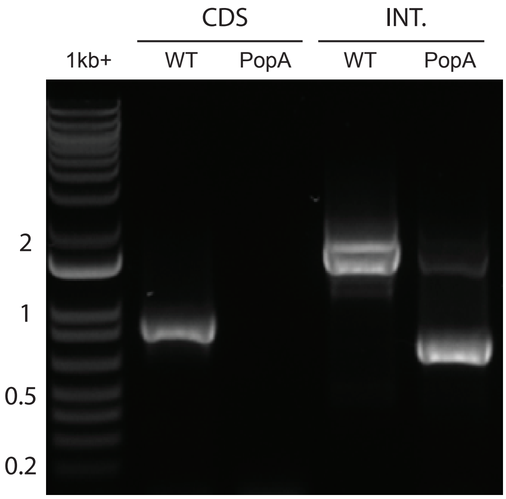


**LmxM.29.3470**

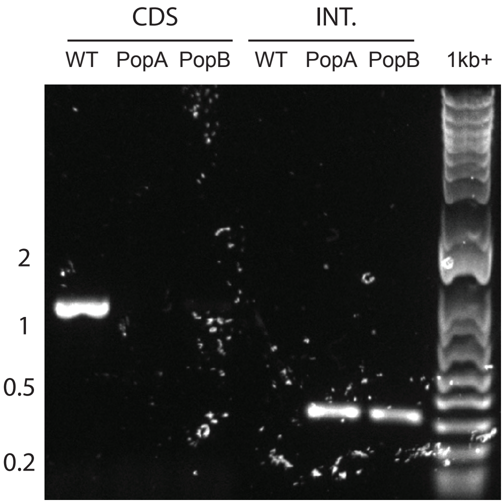


**LmxM.34.1000**

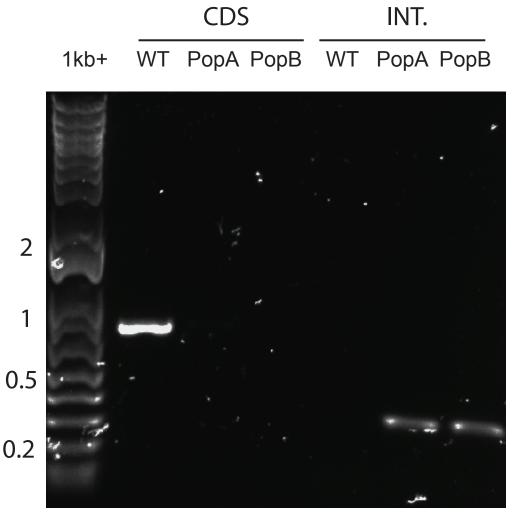


**LmxM.33.3020**

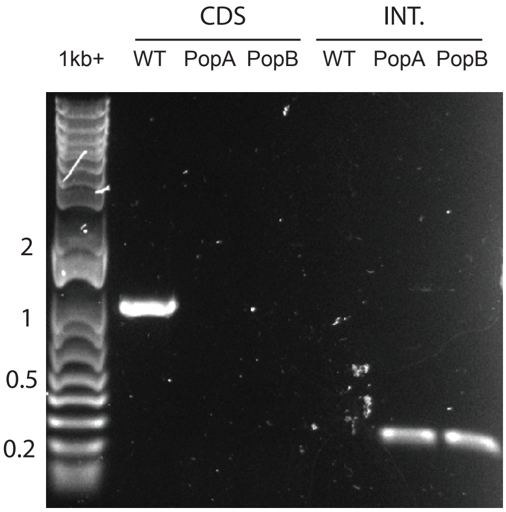

### CMGC FAMILY

**LmxM.32.2070**

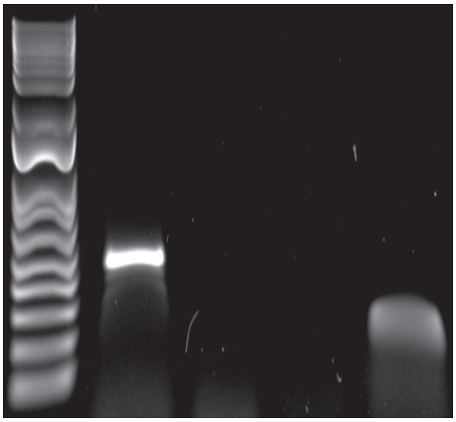


**LmxM.26.0040**

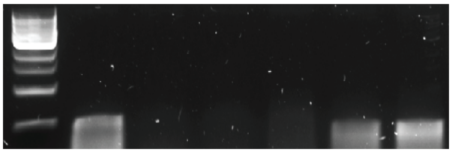


**LmxM.16.0990**

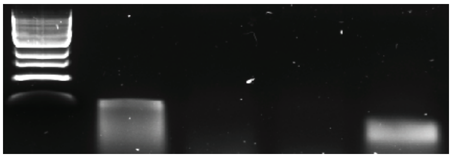


**LmxM.11.0110**

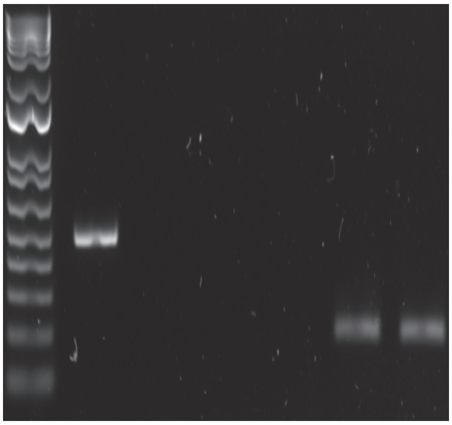


**LmxM.27.0560**

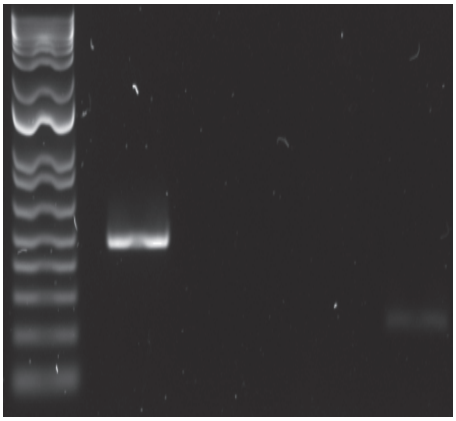


**LmxM.08_29.2150**

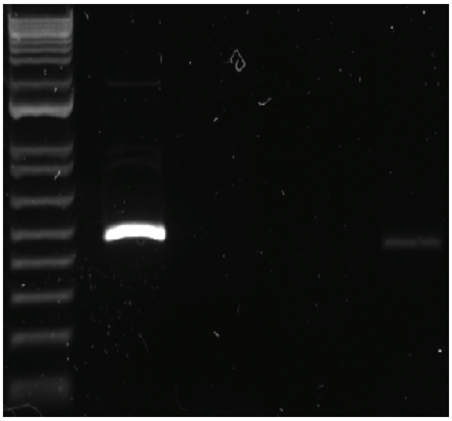


**LmxM.09.0410**

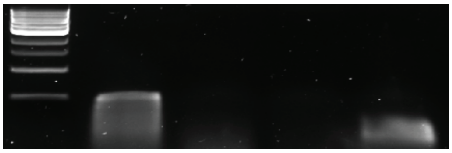


**LmxM.25.1560**

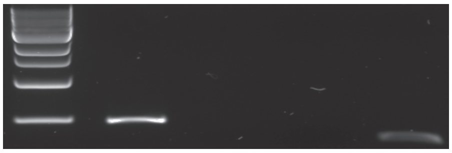


**LmxM.27.1800**

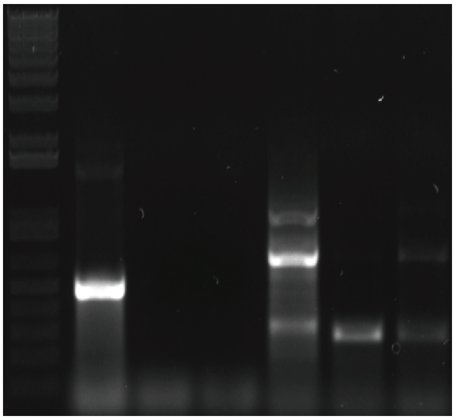


**LmxM.14.1070**

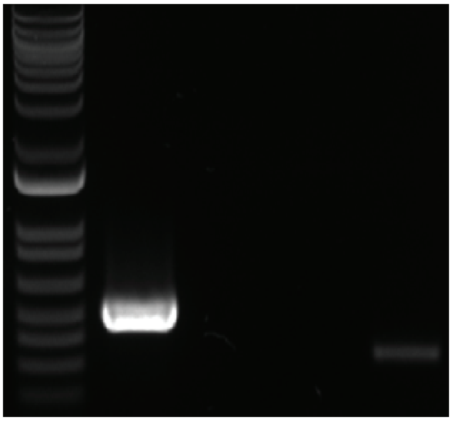


**LmxM.19.0360**

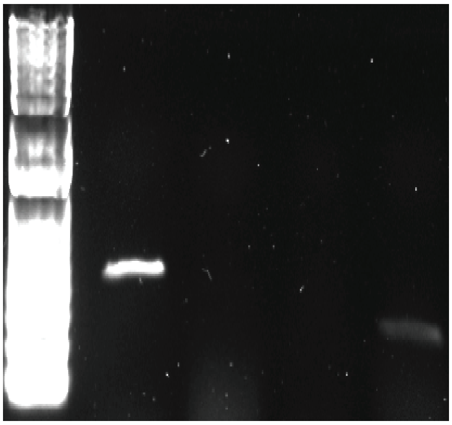

#### CMGC FAMILY

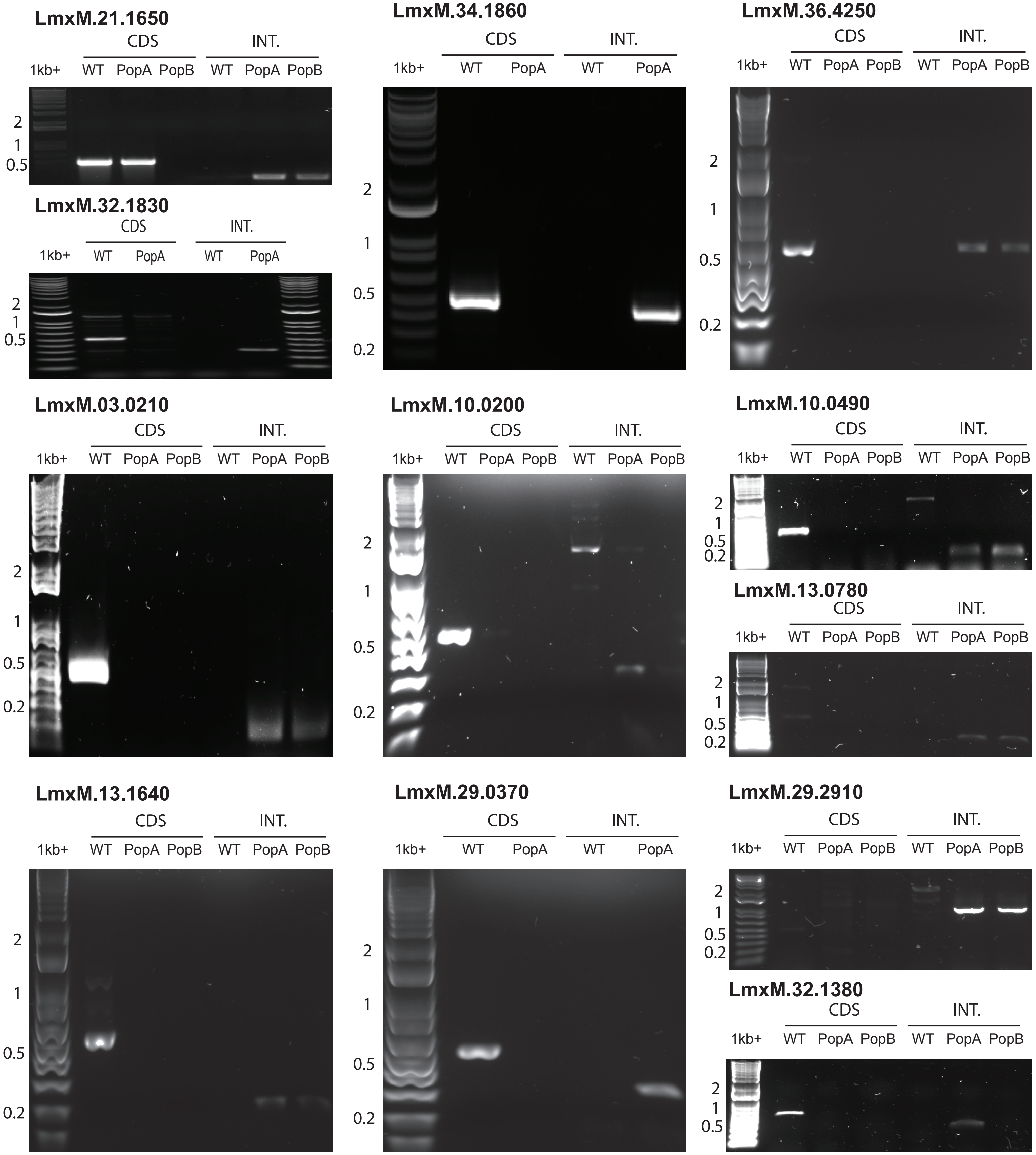
LmxM.21.1650
CDS
INT.
1kb+
WT
PopA
PopB
WT
PopA
PopB
2
1
0.5
LmxM.32.1830
CDS
INT.
1kb+
WT
PopA
WT
PopA
2
1
0.5
LmxM.34.1860
CDS
INT.
1kb+
WT
PopA
WT
PopA
2
1
0.5
0.2
LmxM.36.4250
CDS
INT.
1kb+
WT
PopA
PopB
WT
PopA
PopB
2
1
0.5
0.2
LmxM.03.0210
CDS
INT.
1kb+
WT
PopA
PopB
WT
PopA
PopB
2
1
0.5
0.2
LmxM.10.0200
CDS
INT.
1kb+
WT
PopA
PopB
WT
PopA
PopB
2
1
0.5
0.2
LmxM.10.0490
CDS
INT.
1kb+
WT
PopA
PopB
WT
PopA
PopB
2
1
0.5
0.2
LmxM.13.0780
CDS
INT.
1kb+
WT
PopA
PopB
WT
PopA
PopB
2
1
0.5
0.2
LmxM.13.1640
CDS
INT.
1kb+
WT
PopA
PopB
WT
PopA
PopB
2
1
0.5
0.2
LmxM.29.0370
CDS
INT.
1kb+
WT
PopA
WT
PopA
2
1
0.5
0.2
LmxM.29.2910
CDS
INT.
1kb+
WT
PopA
PopB
WT
PopA
PopB
2
1
0.5
0.2
LmxM.32.1380
CDS
INT.
1kb+
WT
PopA
PopB
WT
PopA
PopB
2
1
0.5

#### CMGC FAMILY

##### LmxM.20_36.6470

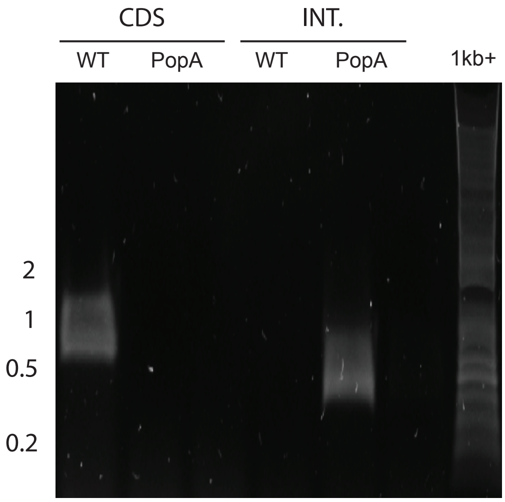


##### LmxM.28.0580

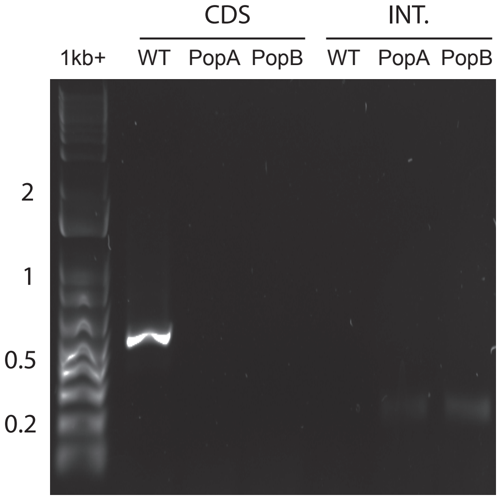


##### LmxM.31.3250

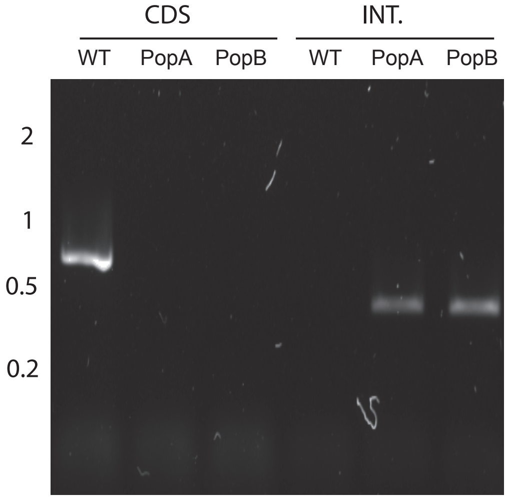


##### LmxM.19.0180

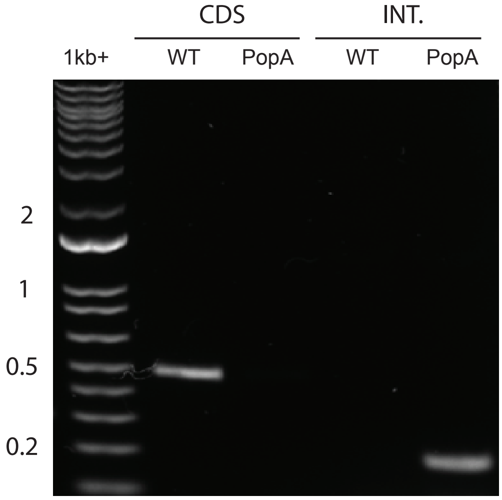


##### LmxM.27.0100

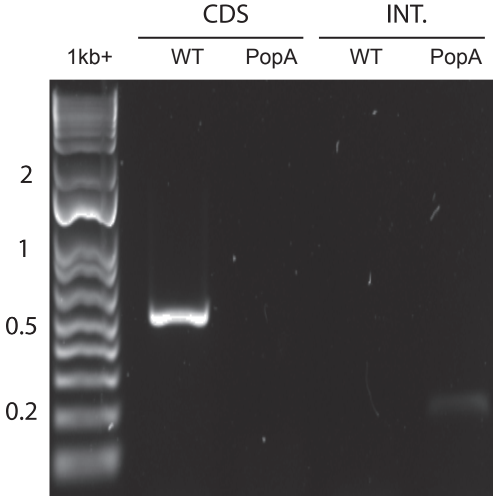


##### LmxM.34.5010

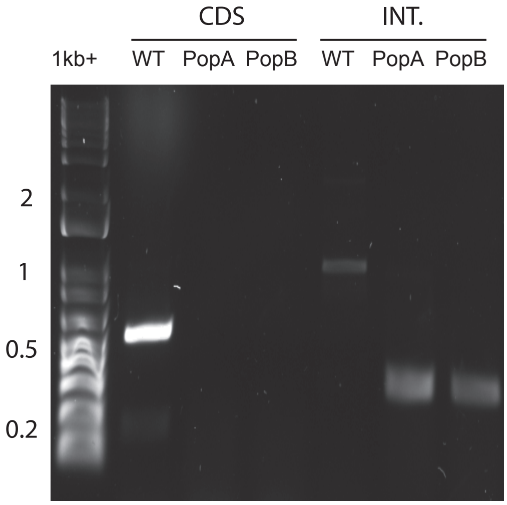


##### LmxM.26.0980

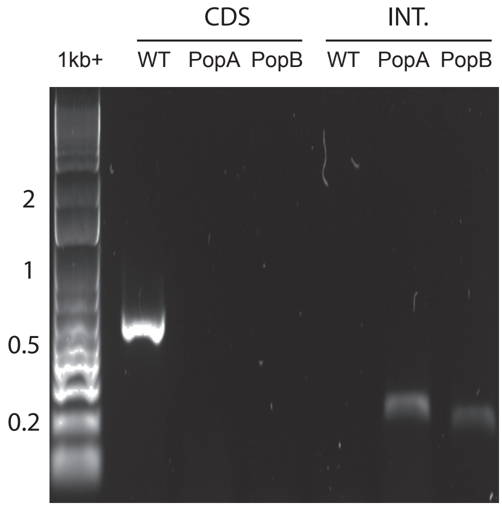


##### LmxM.36.0720

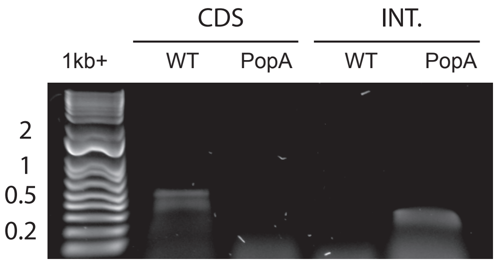


##### LmxM.17.0670

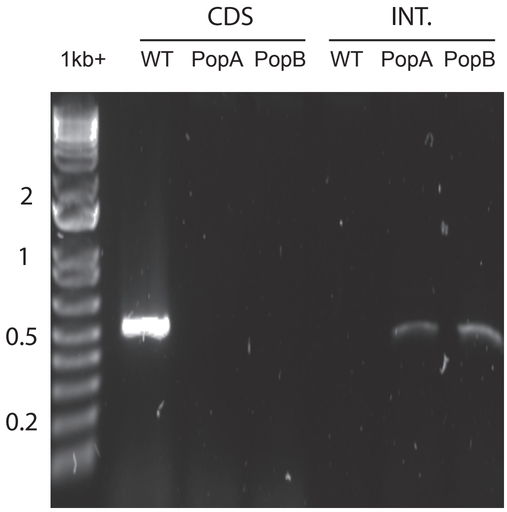

#### CMGC FAMILY

##### LmxM.01.0750

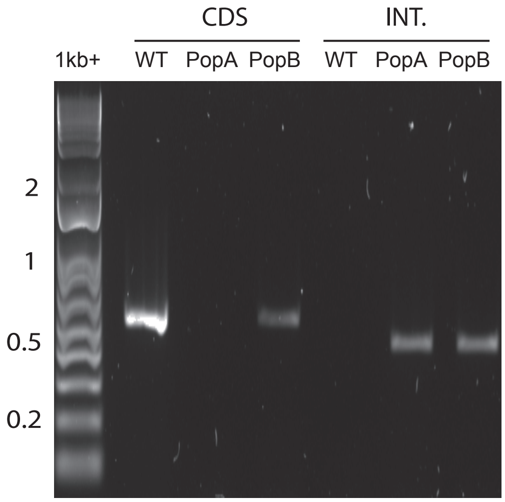


##### LmxM.12.0130

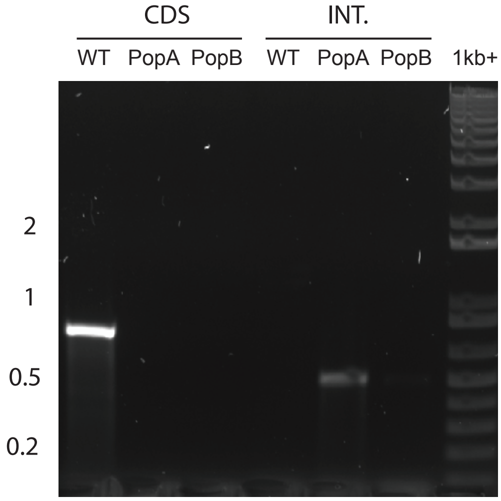


#### STE FAMILY

##### LmxM.08_29.2320

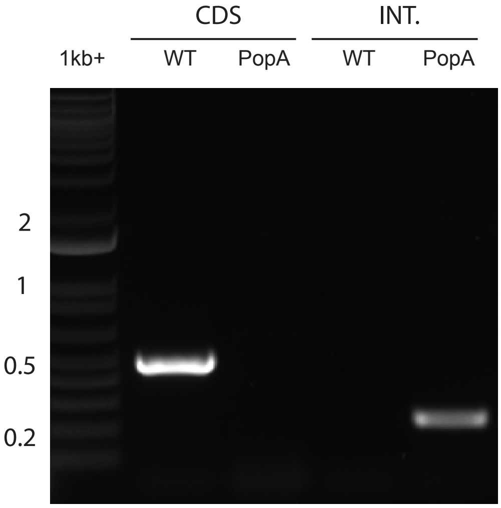


##### LmxM.08.1228

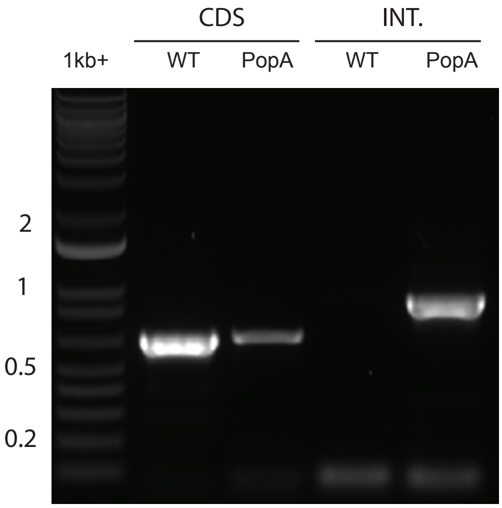


##### LmxM.15.1200

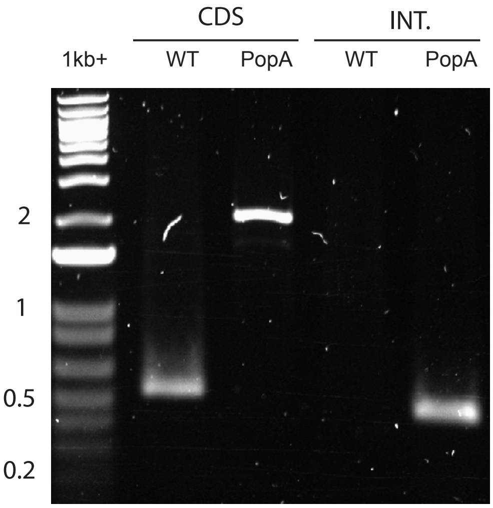


##### LmxM.17.0390

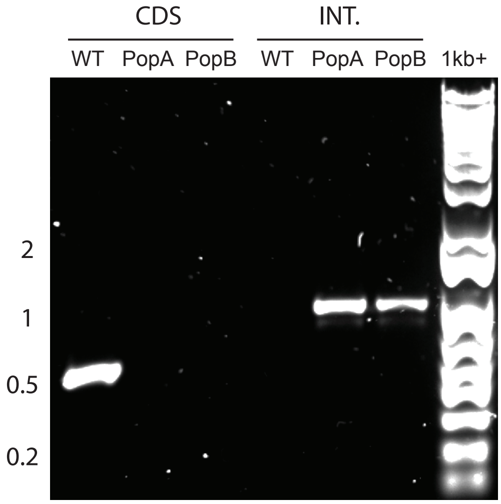


##### LmxM.20.0770

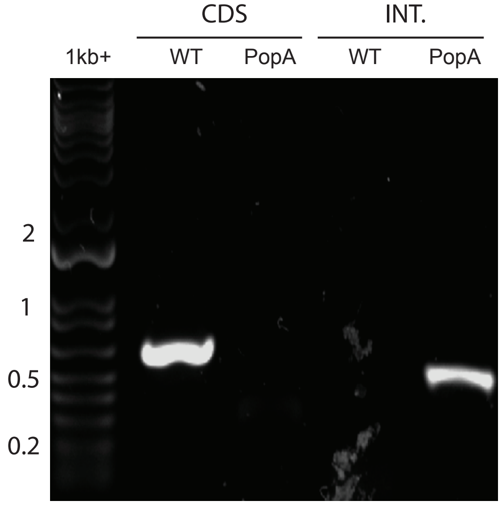


##### LmxM.21.0270

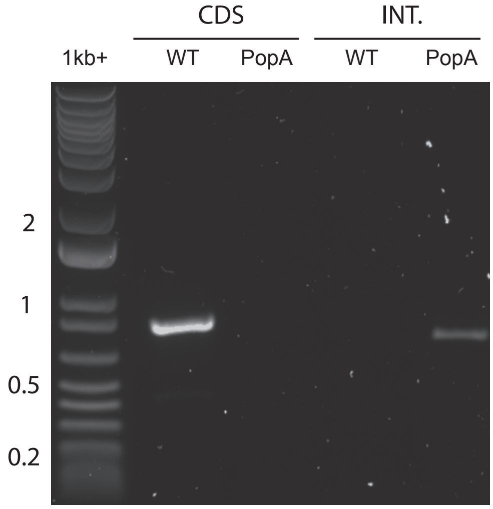

#### STE FAMILY

##### LmxM.25.1990

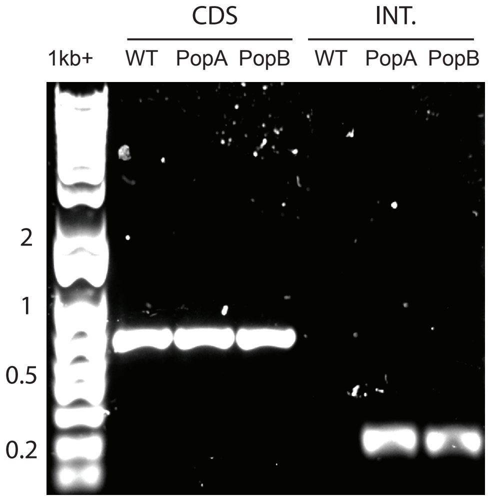


##### LmxM.31.1020

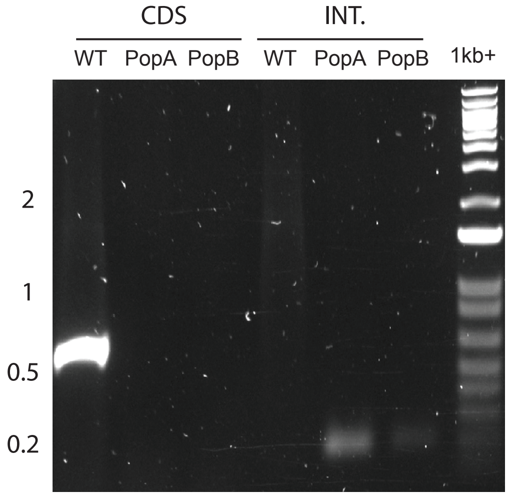


##### LmxM.04.0440

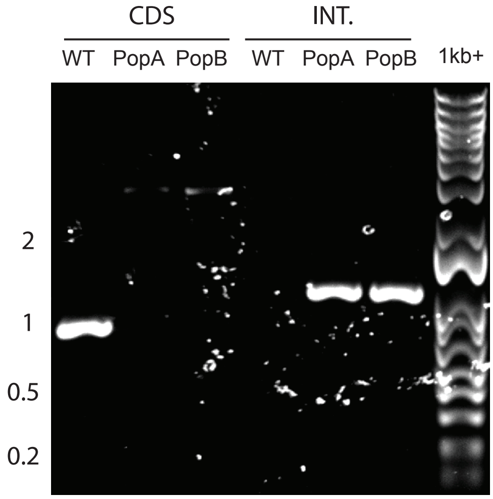


##### LmxM.05.0390

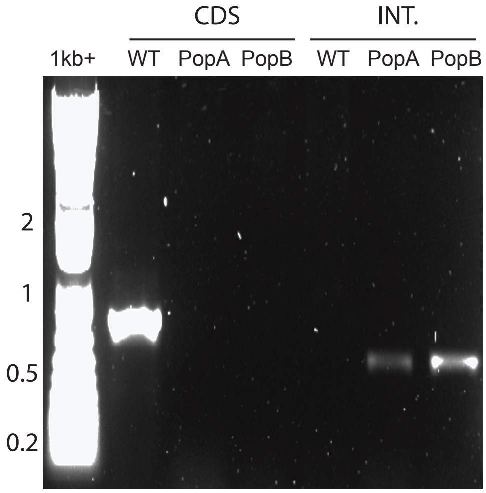


##### LmxM.06.0640

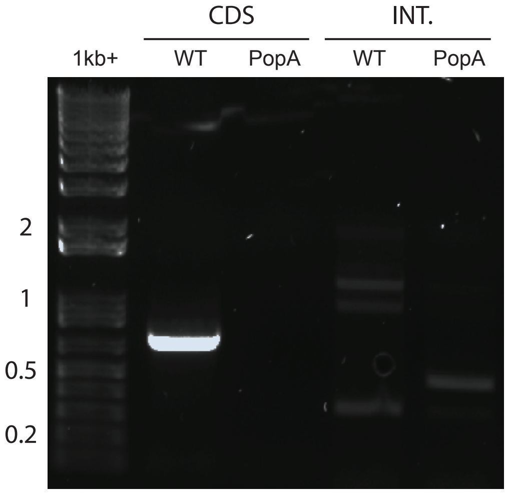


##### LmxM.07.0880

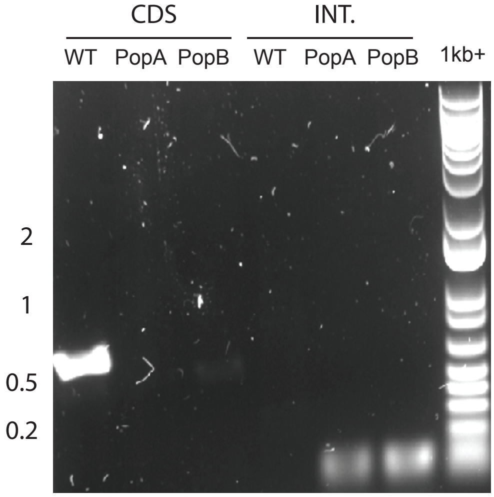


##### LmxM.19.0150

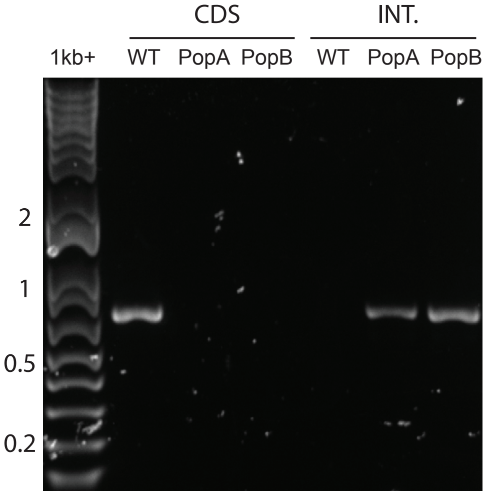


##### LmxM.21.0130

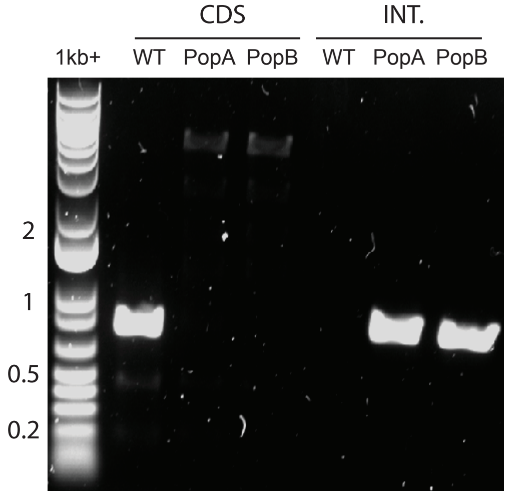


##### LmxM.24.1450

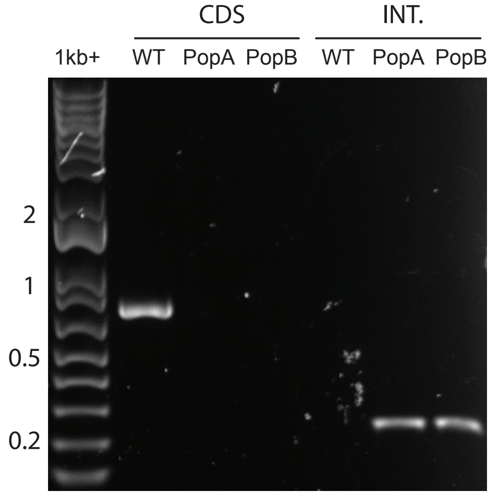

#### STE FAMILY

##### LmxM.26.1730

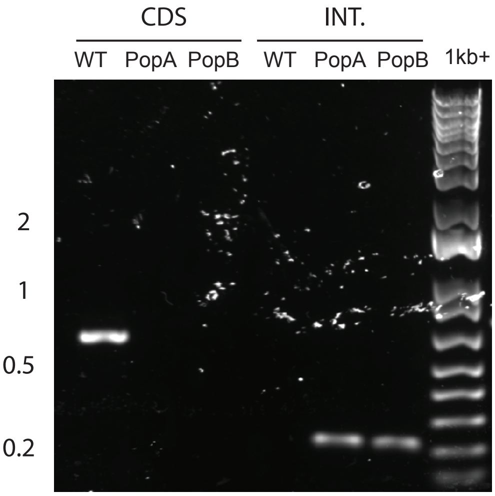


##### LmxM.29.0600

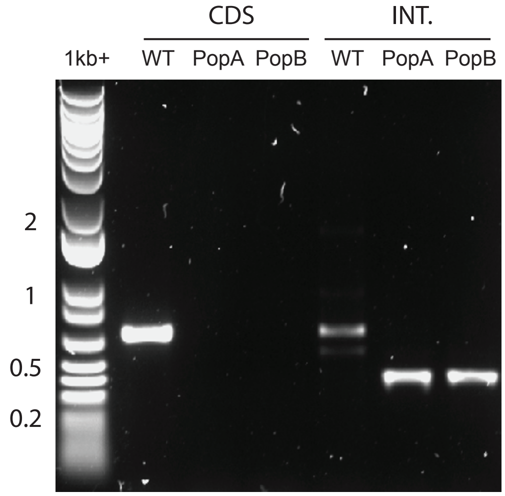


##### LmxM.29.3050

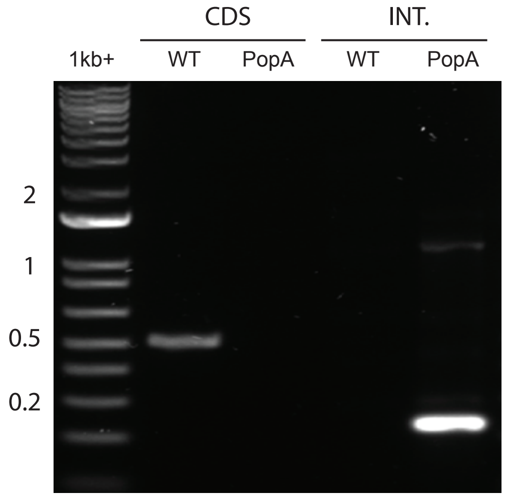


##### LmxM.31.0120

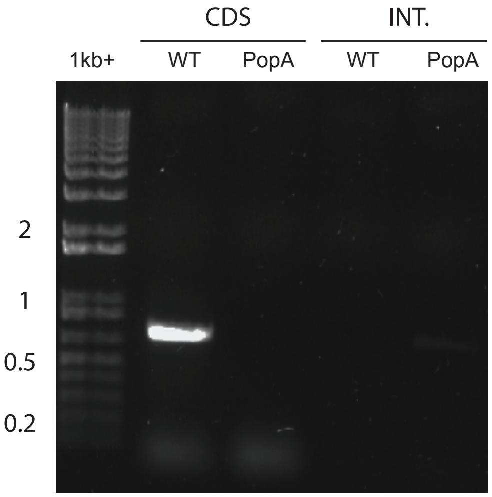


##### LmxM.31.0780

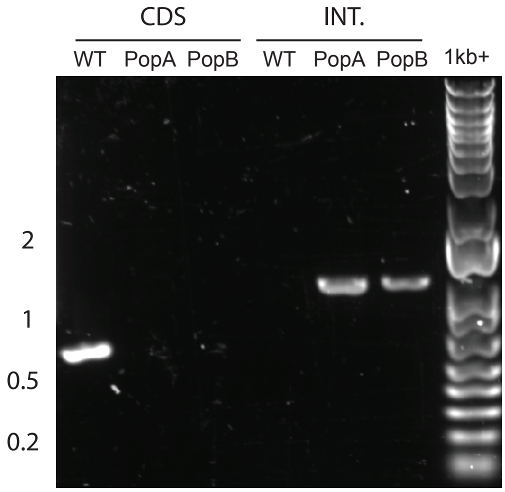


##### LmxM.31.0810

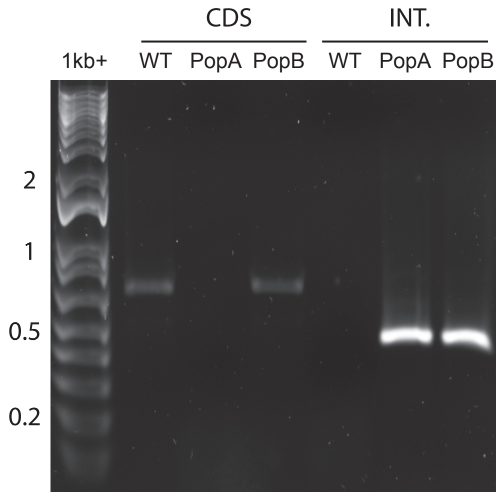


##### LmxM.32.1400

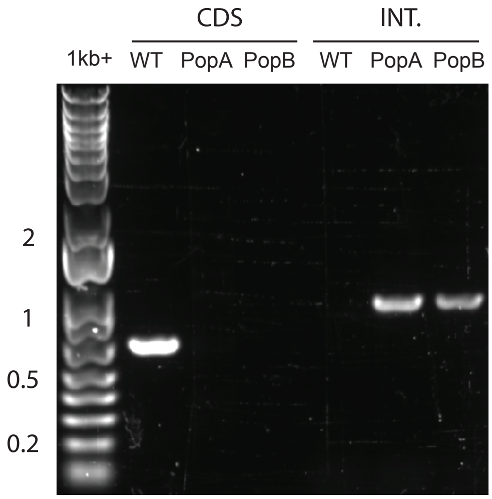


##### LmxM.32.2290

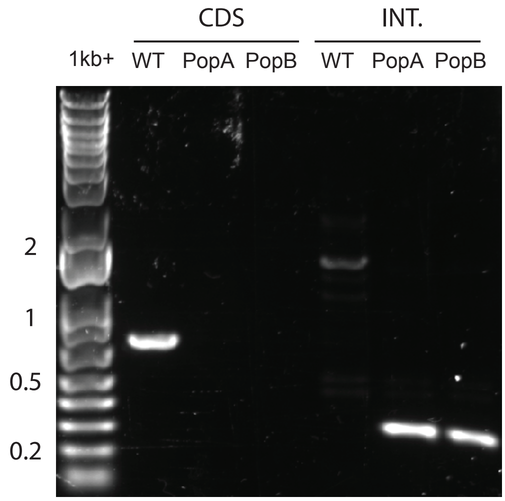


##### LmxM.36.0910

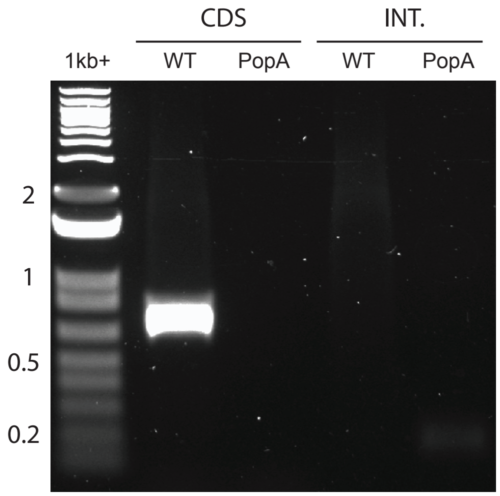

#### STE FAMILY

##### LmxM.16.0300

CDS | INT.

1kb+ WT PopA WT PopA

2
1
0.5
0.2

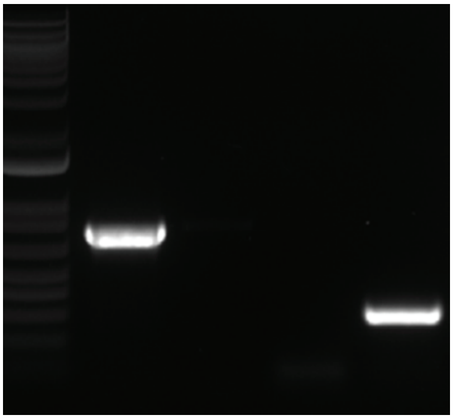

##### LmxM.07.0250

CDS | INT.

WT PopA PopB WT PopA PopB

2
1
0.5
0.2

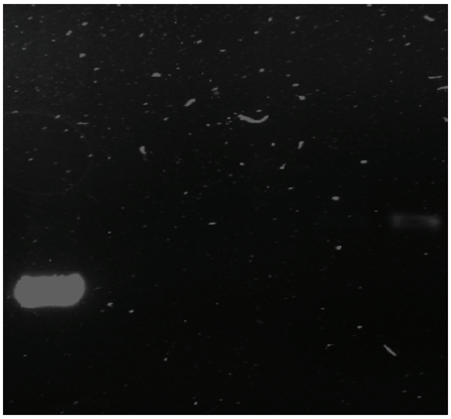

##### LmxM.19.1610

CDS | INT.

1kb+ WT PopA PopB WT PopA PopB

2
1
0.5
0.2

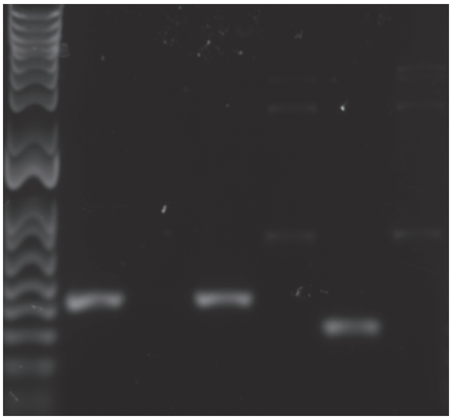

##### LmxM.33.2090

CDS | INT.

1kb+ WT PopA PopB WT PopA PopB

2
1
0.5
0.2

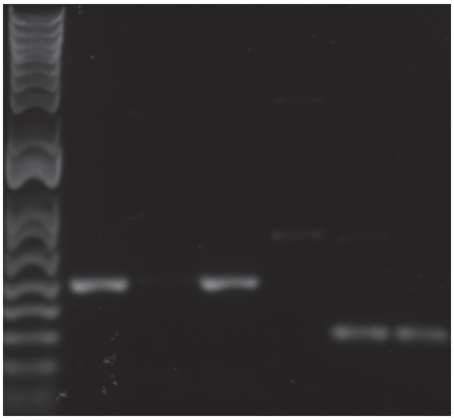

##### LmxM.34.4000

CDS | INT.

1kb+ WT PopA PopB WT PopA PopB

2
1
0.5
0.2

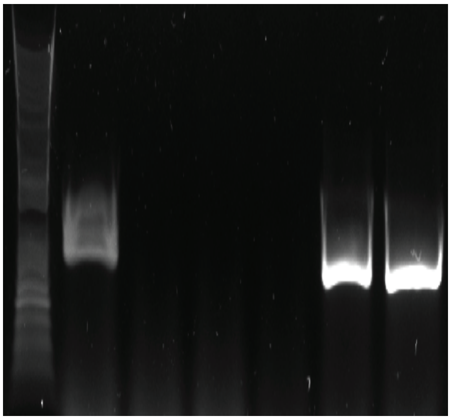

##### LmxM.36.3680

CDS | INT.

1kb+ WT PopA PopB WT PopA PopB

2
1
0.5
0.2

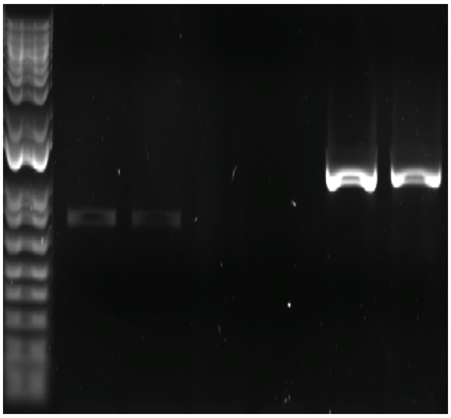

##### LmxM.30.1830

CDS | INT.

1kb+ WT PopA PopB WT PopA PopB

2
1
0.5
0.2

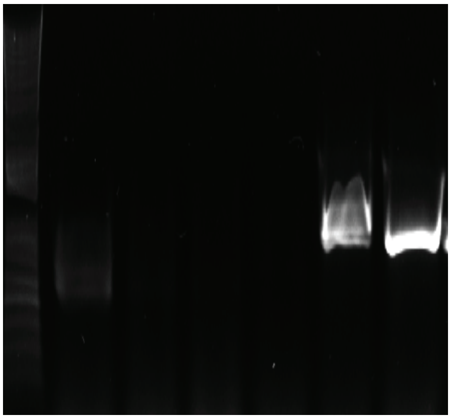

##### LmxM.30.1840

CDS | INT.

1kb+ WT PopA PopB WT PopA PopB

2
1
0.5

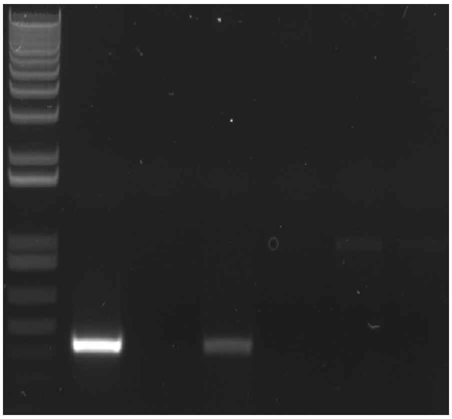

#### NEK FAMILY

##### LmxM.02.0290

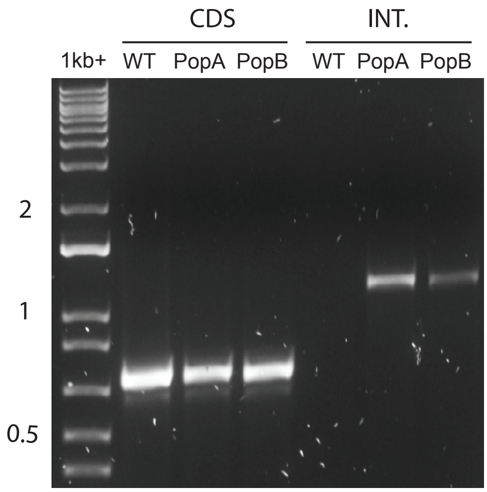


##### LmxM.07.0160

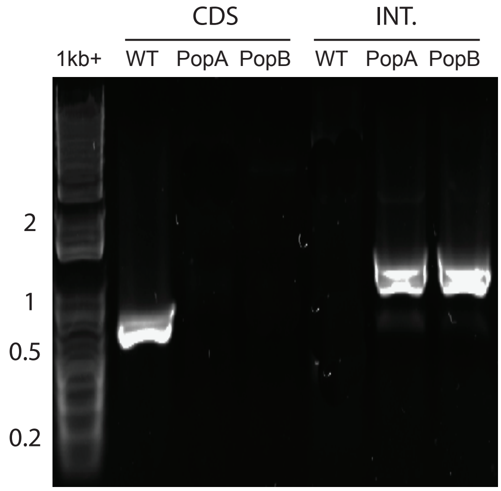


##### LmxM.07.0170

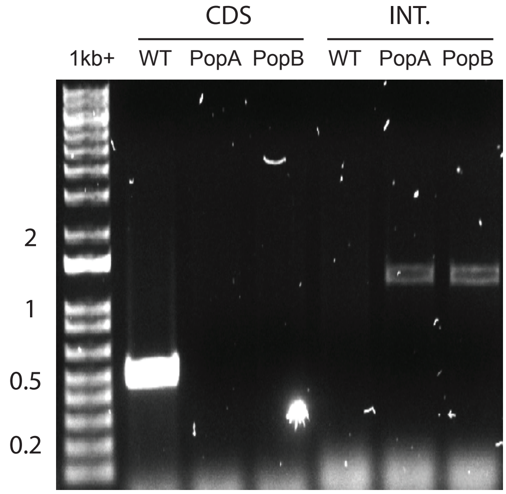


##### LmxM.08.0930

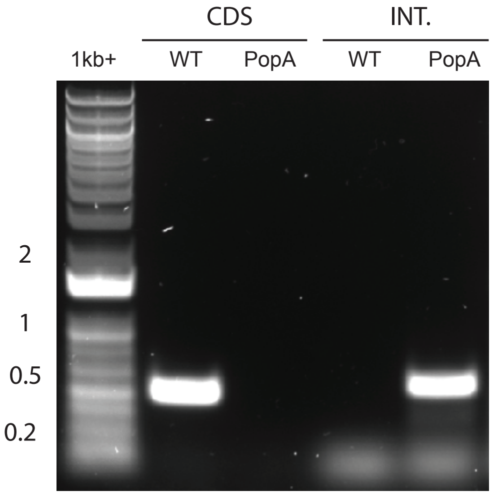


##### LmxM.14.1410

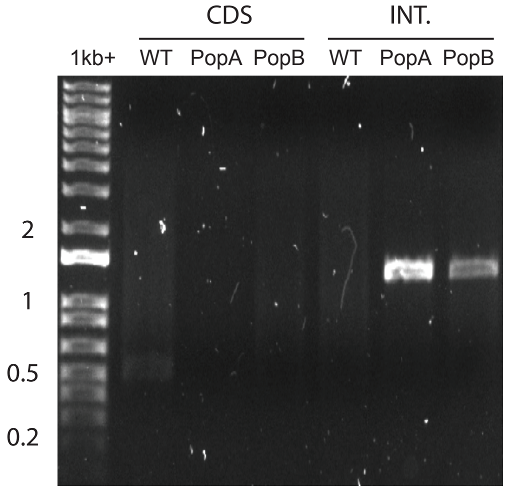


##### LmxM.21.0853

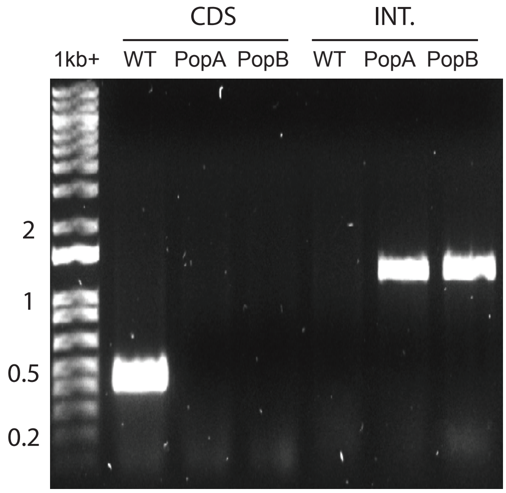


##### LmxM.21.1565

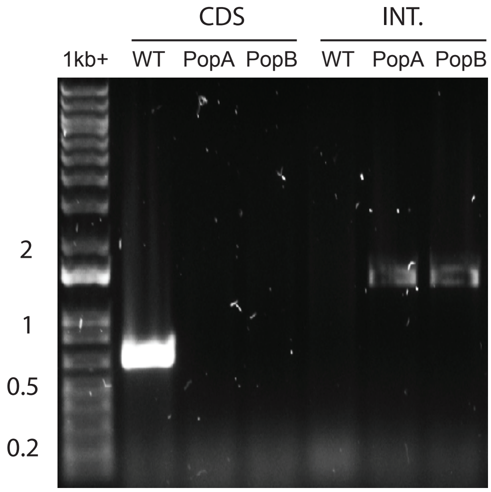


##### LmxM.22.0950

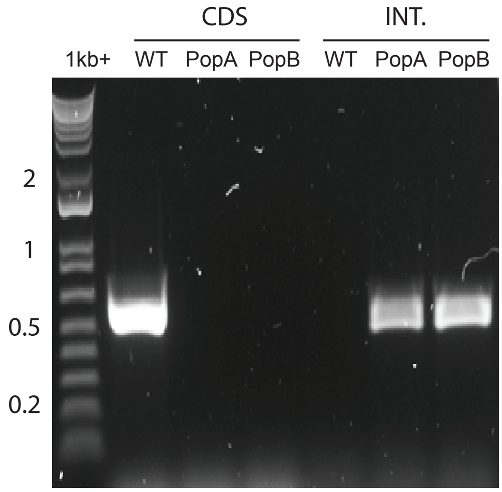


##### LmxM.26.2570

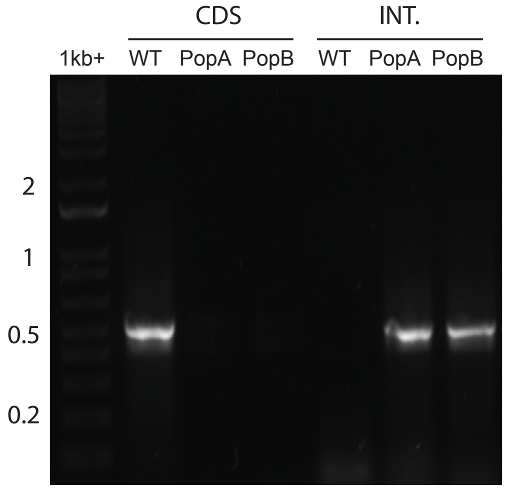

### NEK FAMILY

##### LmxM.28.3000

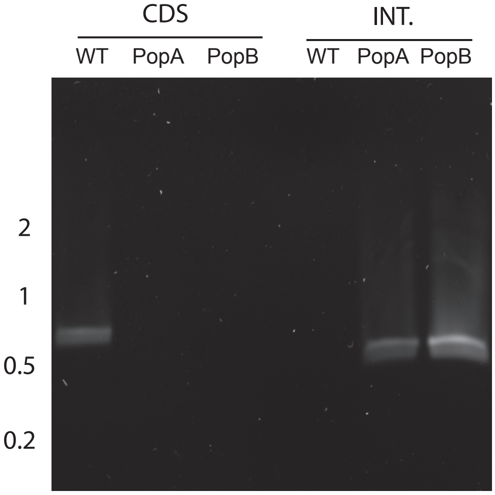


##### LmxM.08_29.2570

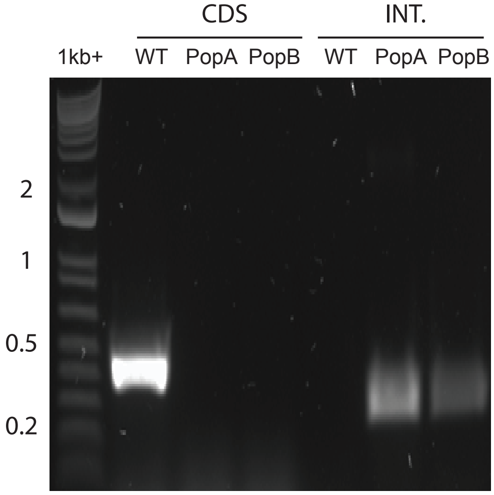


##### LmxM.08_29.2670

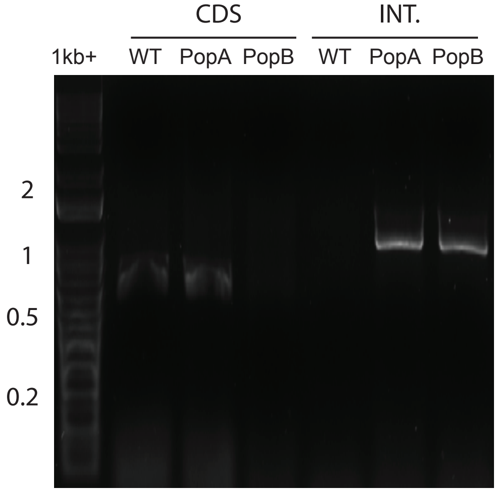


##### LmxM.29.2130

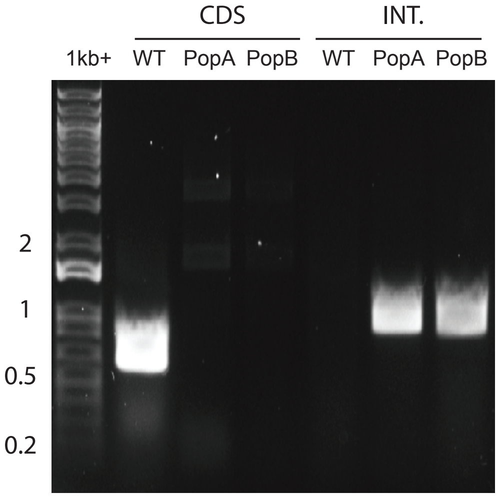


##### LmxM.30.2960

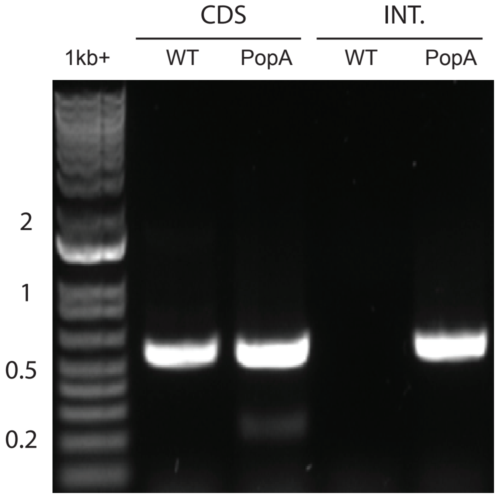


##### LmxM.30.3160

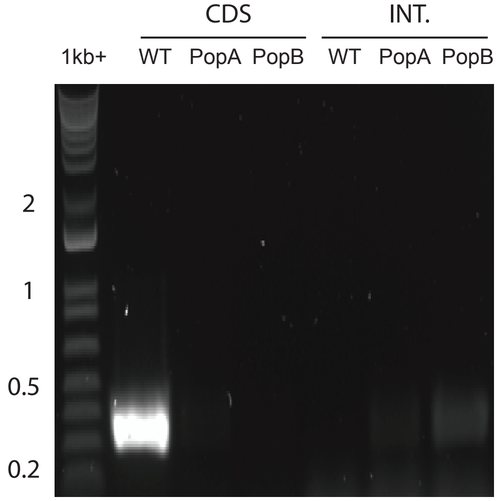


##### LmxM.31.0260

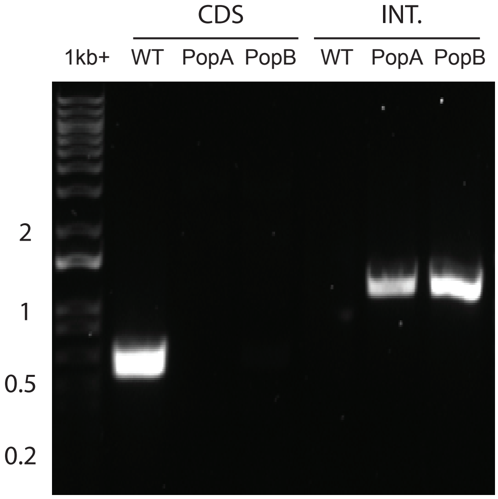


##### LmxM.31.1810

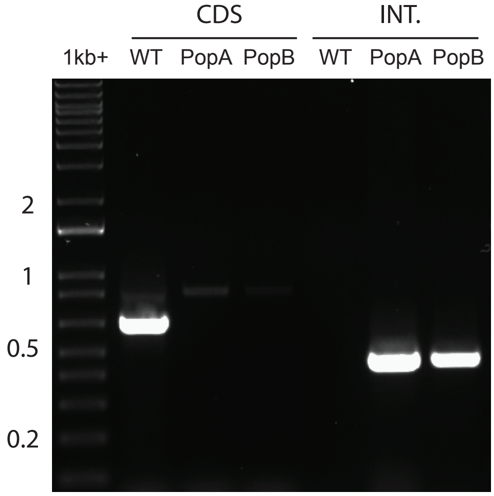


##### LmxM.32.1980

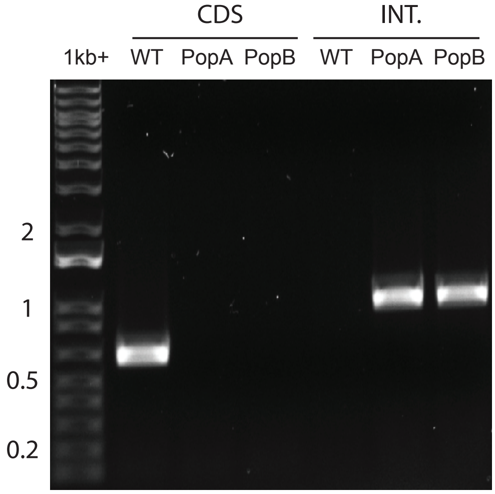

#### NEK FAMILY

**LmxM.34.5190**

CDS | INT.

1kb+ WT PopA PopB WT PopA PopB

2
1
0.5
0.2

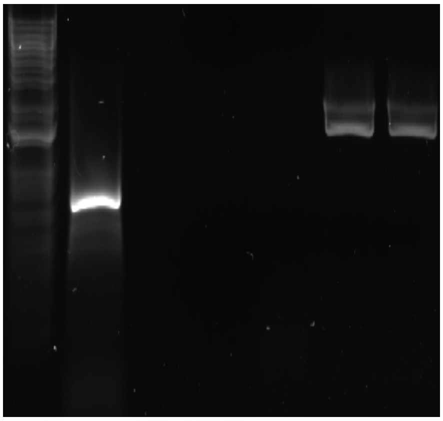

**LmxM.36.1520**

CDS | INT.

WT PopA PopB WT PopA PopB 1kb+

2
1
0.5
0.2

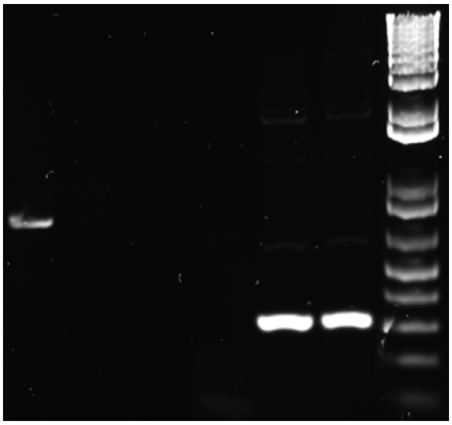

**LmxM.36.1530**

CDS | INT.

1kb+ WT PopA PopB WT PopA PopB

2
1
0.5
0.2

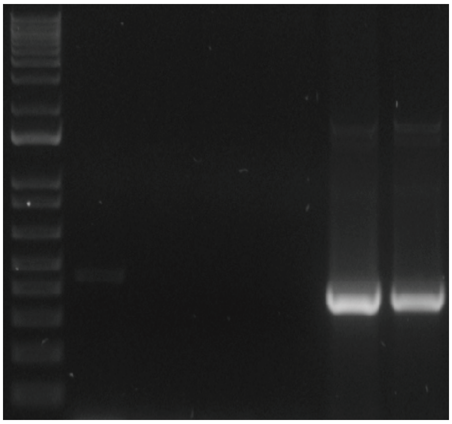

**LmxM.36.2290**

CDS | INT.

1kb+ WT PopA PopB WT PopA PopB

2
1
0.5
0.2

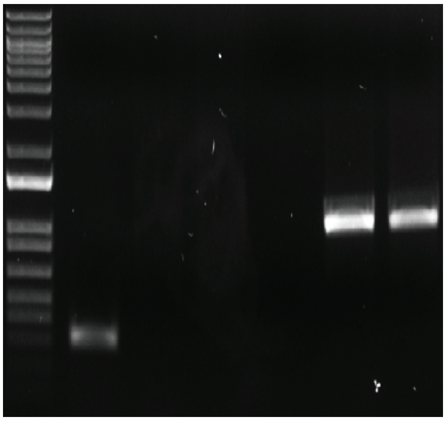

#### ORPHAN KINASES

**LmxM.03.0350**

CDS | INT.

1kb+ WT PopA WT PopA 1kb+

2
1
0.5
0.2

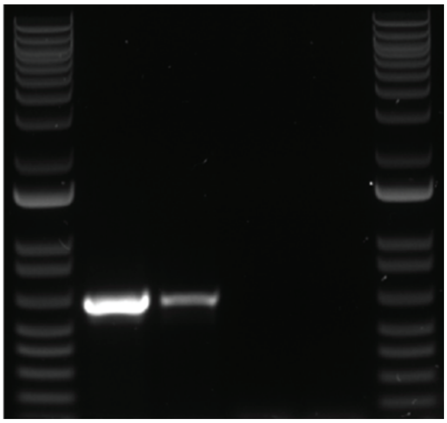

**LmxM.08_29.2490**

CDS | INT.

1kb+ WT PopA WT PopA 1kb+

2
1
0.5
0.2

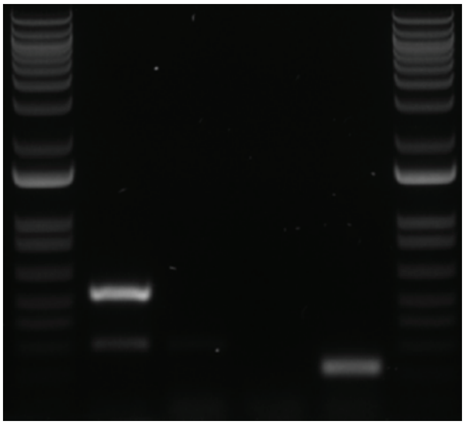

**LmxM.26.2110**

CDS | INT.

1kb+ WT PopA WT PopA 1kb+

2
1
0.5
0.2

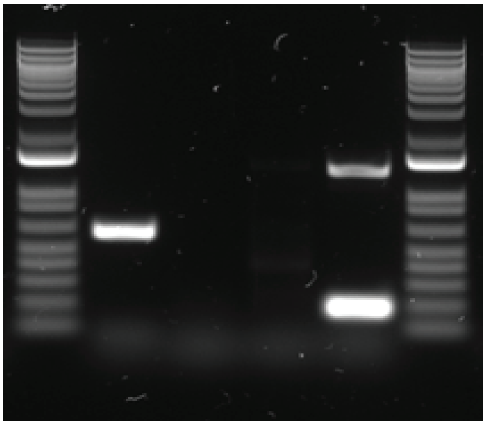

### ORPHAN KINASES

**LmxM.16.0870**

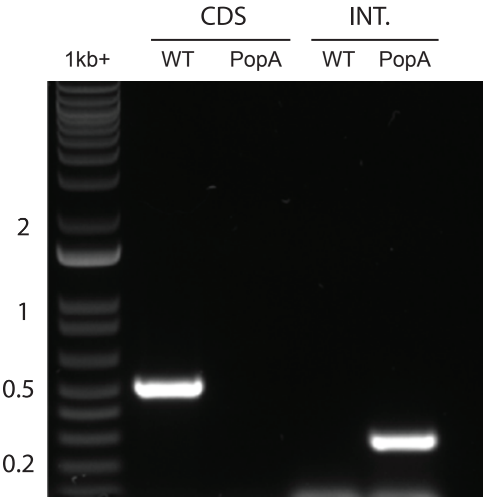


**LmxM.11.0510**

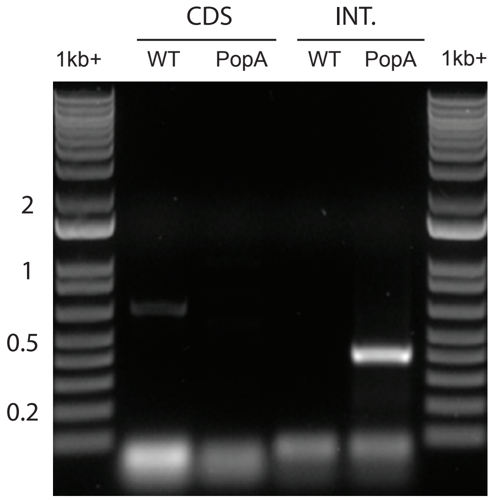


**LmxM.25.1520**

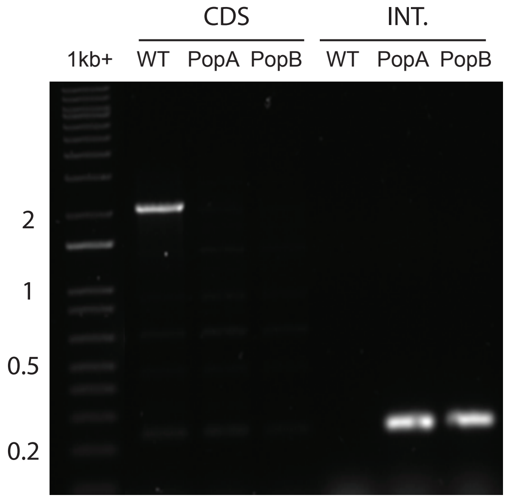


**LmxM.02.0570**

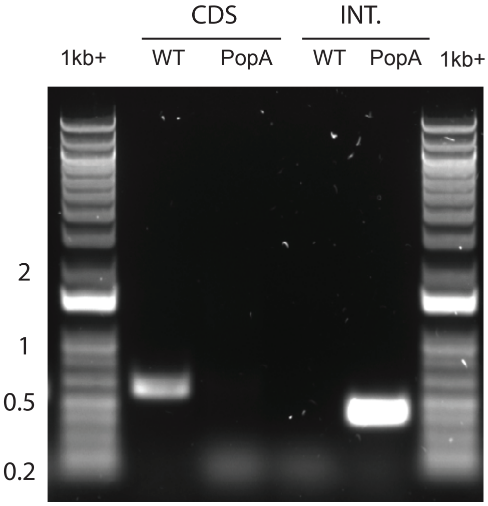


**LmxM.10.0830**

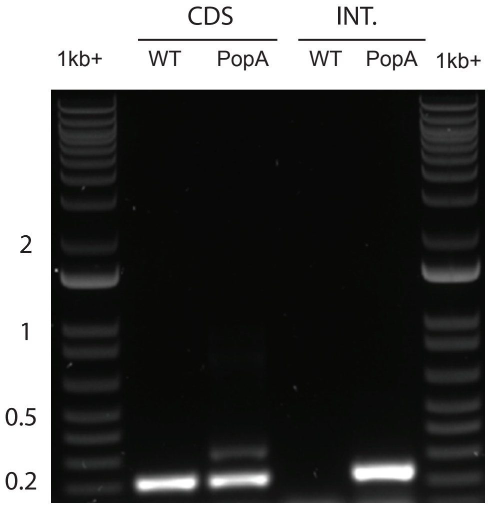


**LmxM.34.4620**

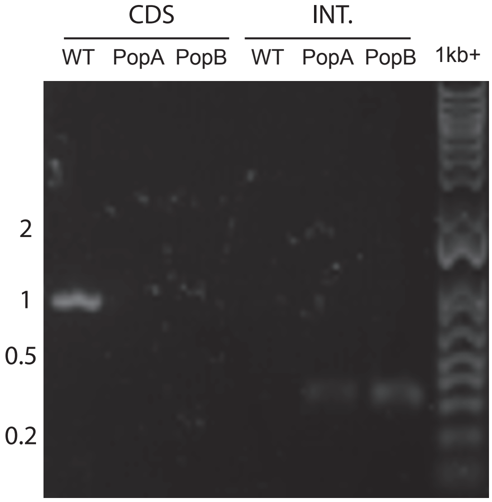


**LmxM.08.0660**

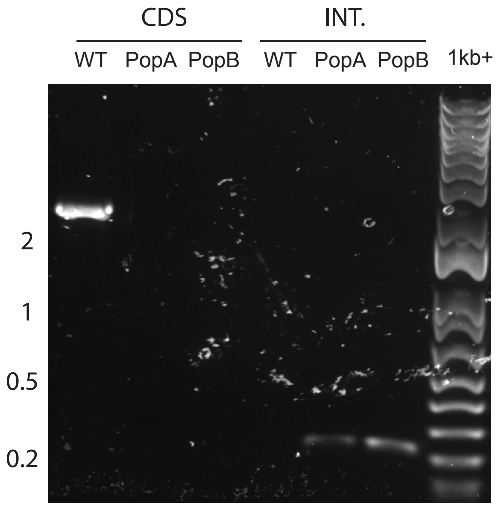


**LmxM.26.2060**

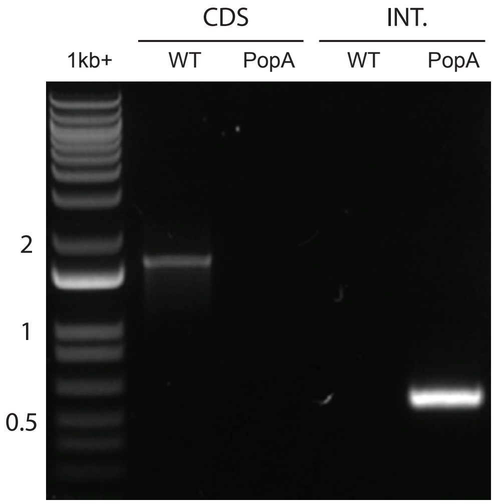


**LmxM.28.1650**

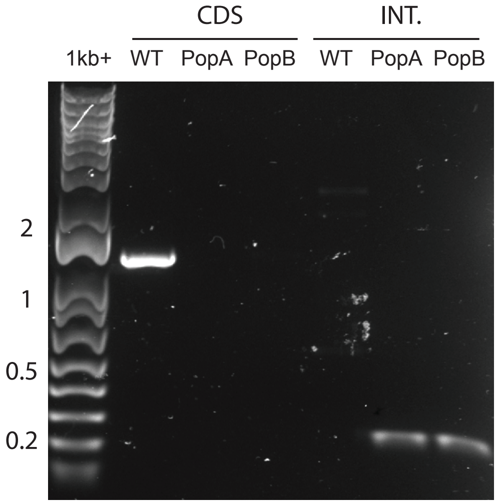

#### ORPHAN KINASES

**LmxM.31.1290**

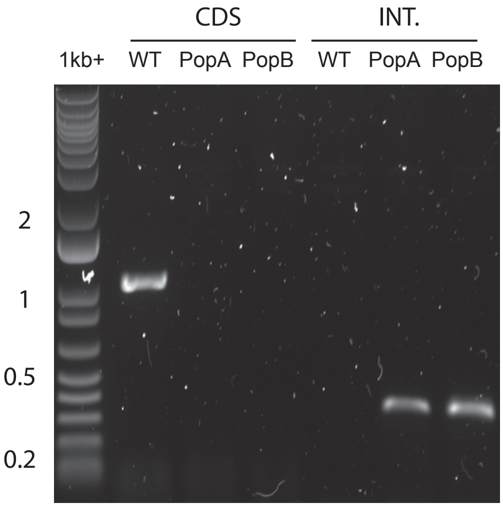


**LmxM.33.2190**

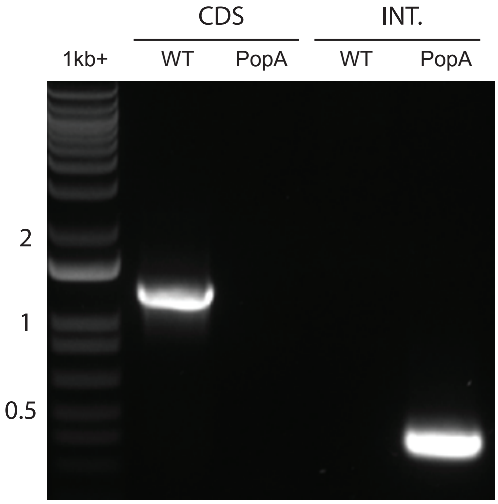


#### OTHER KINASES

**LmxM.20.1330**

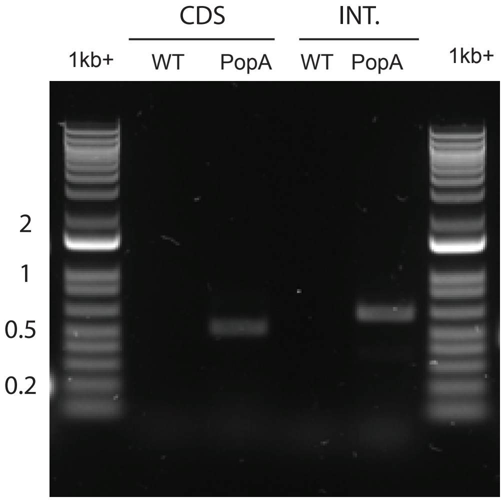


**LmxM.20.1340**

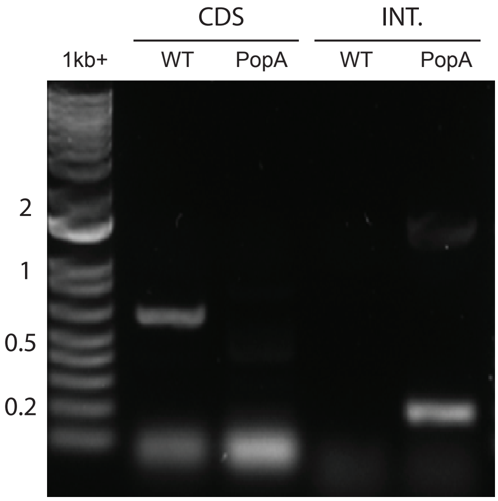


**LmxM.08_29.0370**

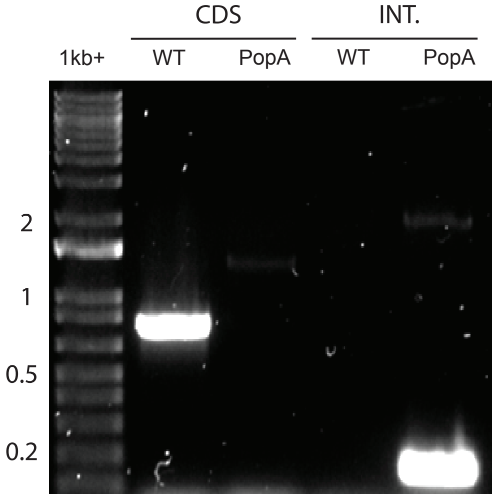


**LmxM.34.2870**

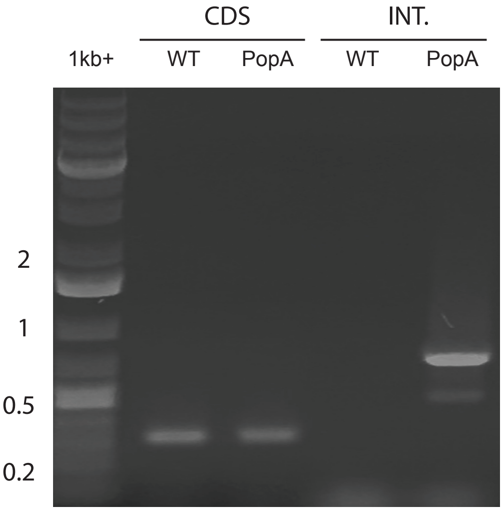


**LmxM.26.2440**

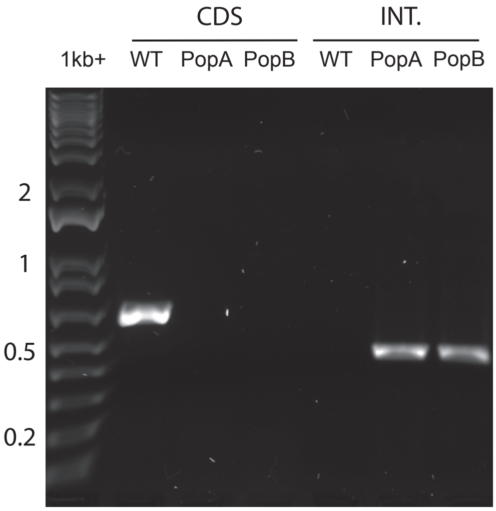


**LmxM.28.0520**

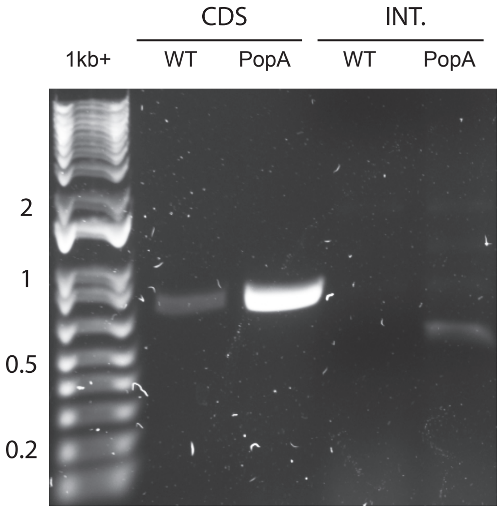

### OTHER KINASES

##### LmxM.08_29.1330

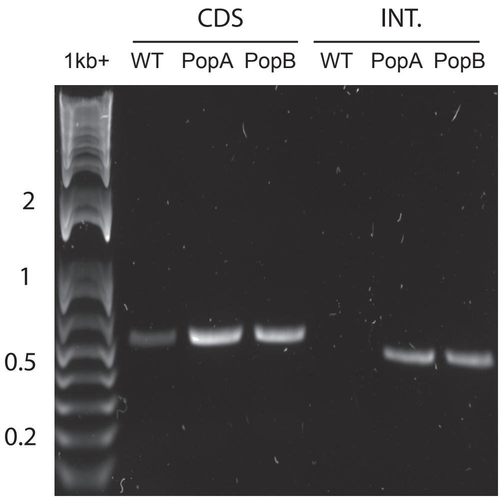


##### LmxM.19.0590

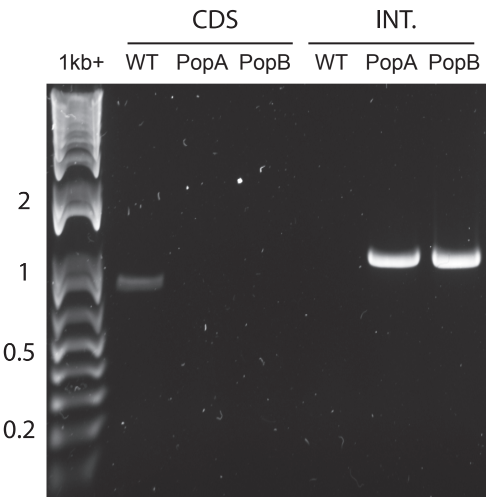


##### LmxM.11.0250

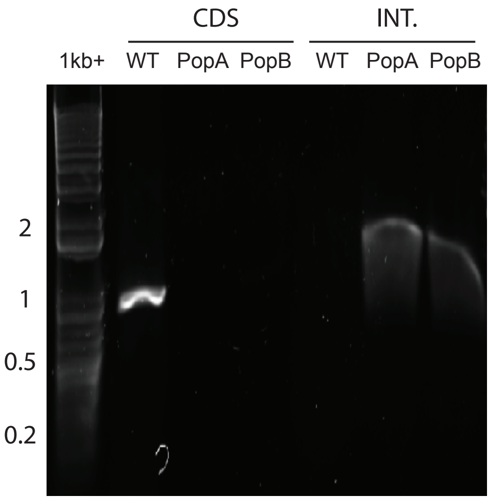


##### LmxM.24.1730

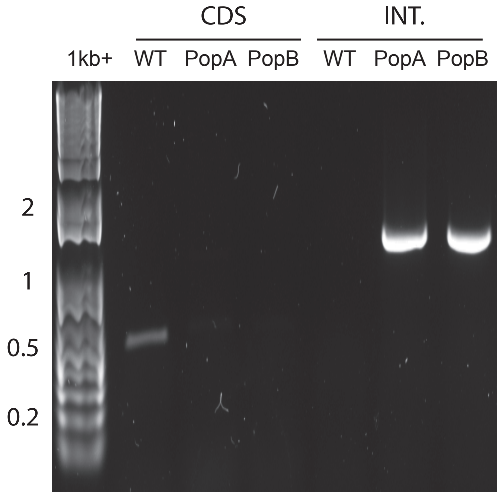


##### LmxM.34.2320

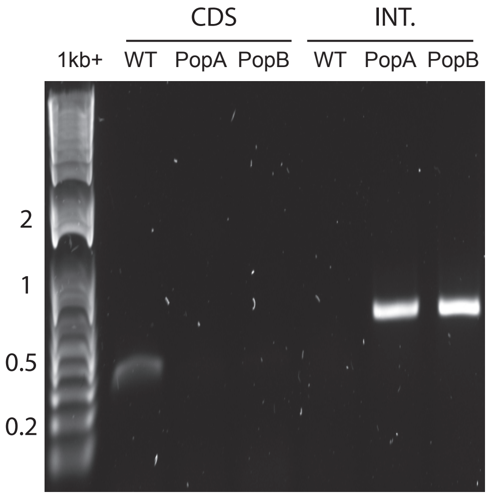


##### LmxM.34.1730

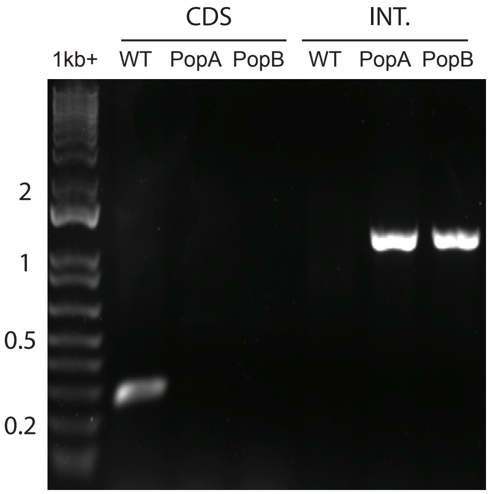


##### LmxM.02.0360

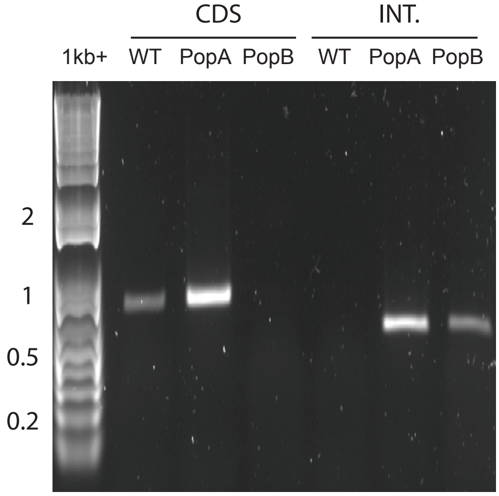


##### LmxM.15.0770

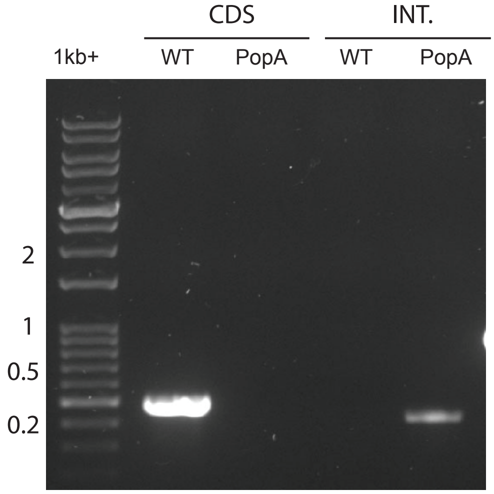


##### LmxM.33.0030

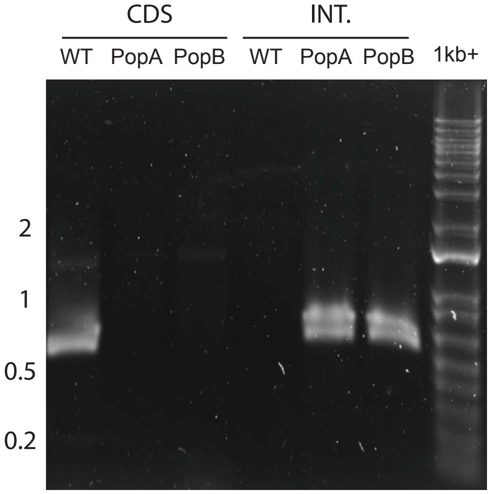

#### OTHER KINASES

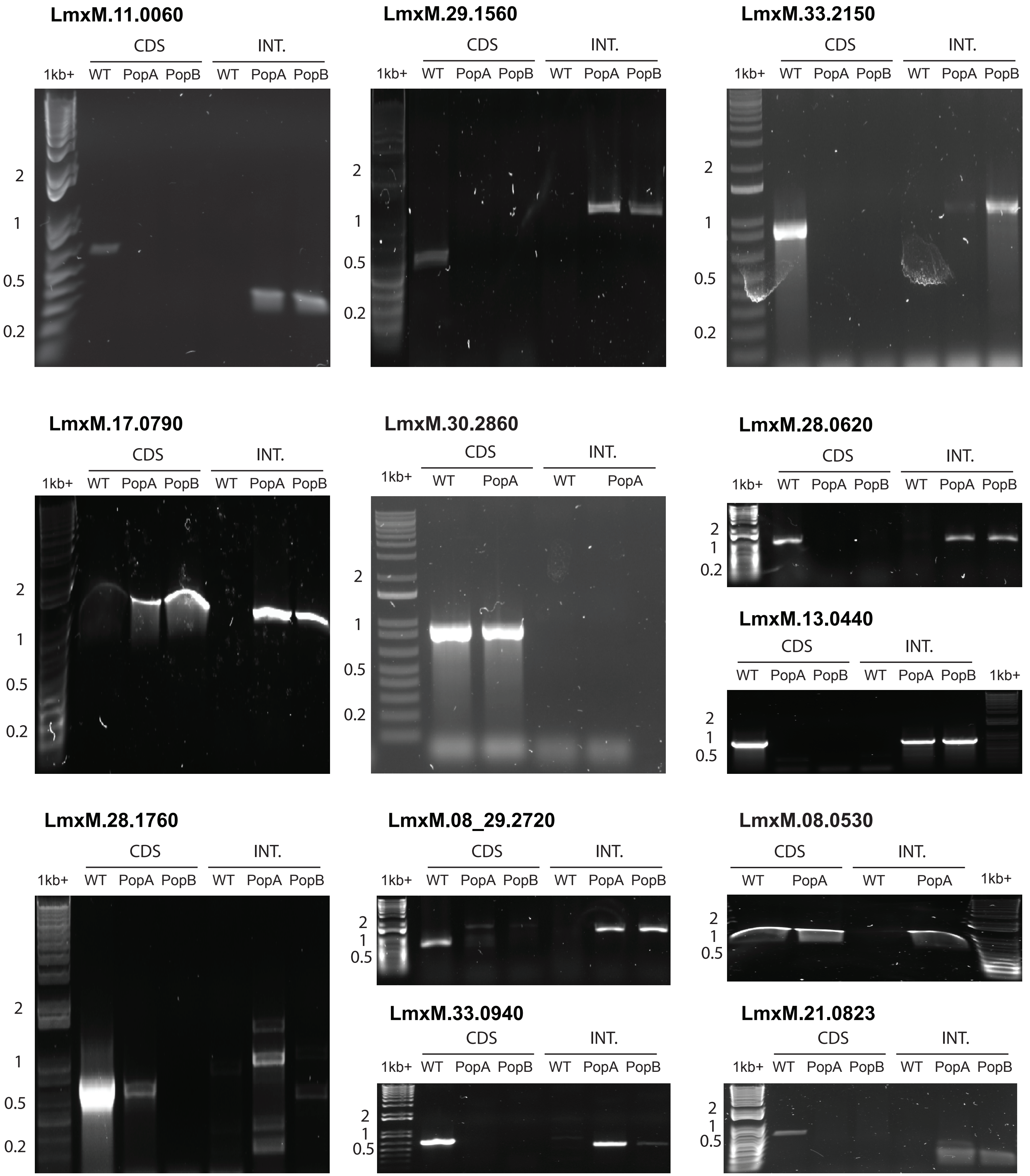

#### OTHER KINASES

##### LmxM.36.2630

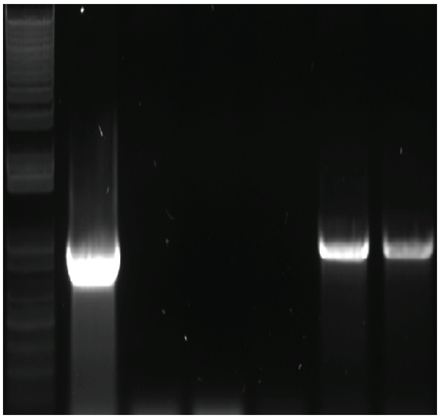


##### LmxM.22.1150

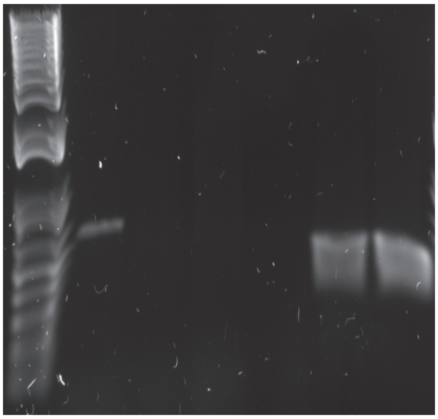


#### PIKK (aPK)

##### LmxM.08_29.1450

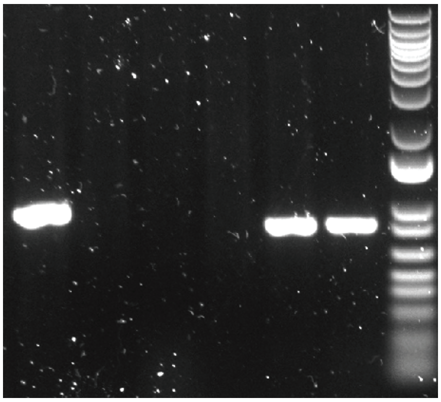


##### LmxM.24.2010

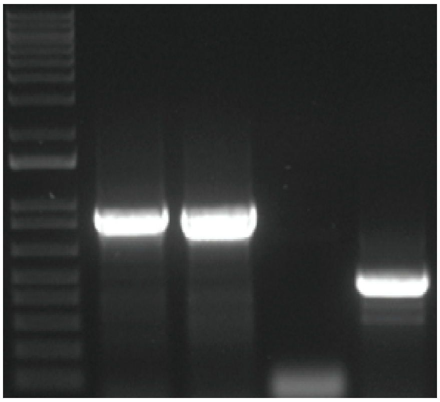


##### LmxM.33.3590

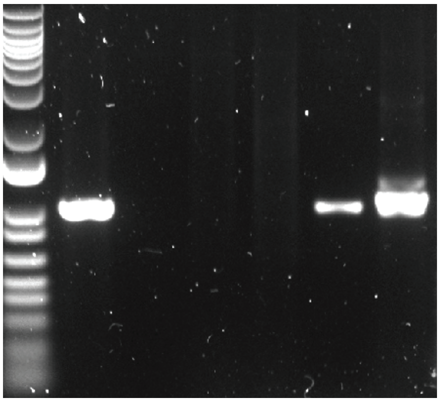


##### LmxM.27.0890

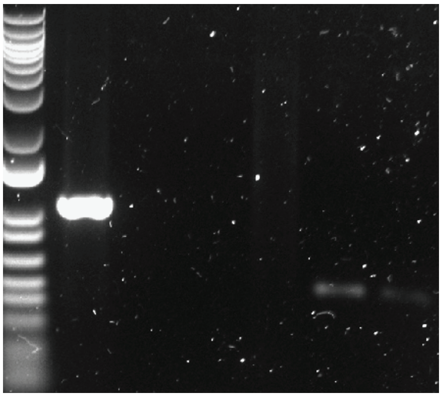


##### LmxM.33.3090

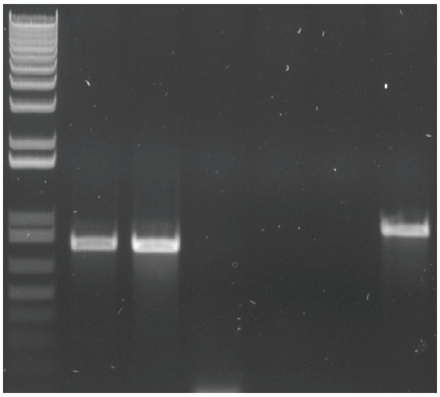


##### LmxM.34.0560

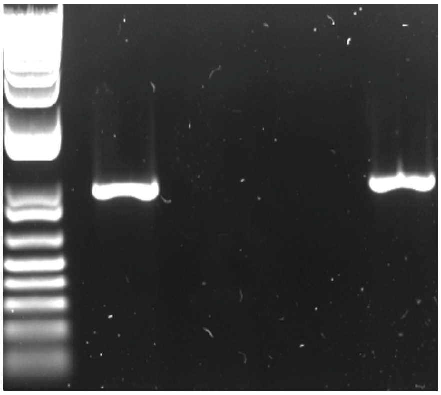

### OTHER KINASES

**LmxM.36.6320**

CDS | INT.

1kb+ WT PopA WT PopA

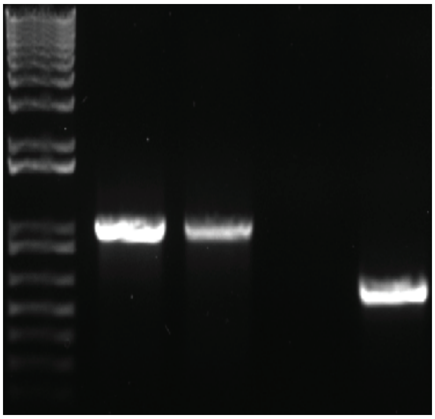

2 1 0.5 0.2

**LmxM.33.3940**

CDS | INT.

1kb+ WT PopA WT PopA

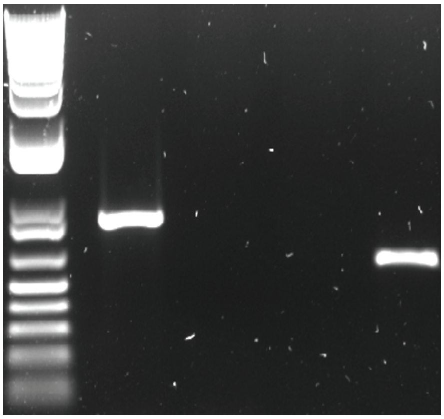

2 1 0.5 0.2

**LmxM.02.0120**

CDS | INT.

WT PopA PopB WT PopA PopB 1kb+

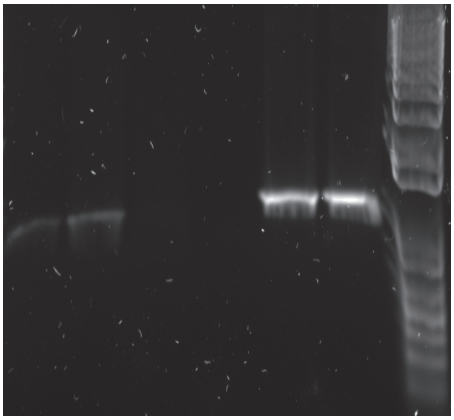

2 1 0.5 0.2

**LmxM.31.1460**

CDS | INT.

WT PopA PopB WT PopA PopB 1kb+

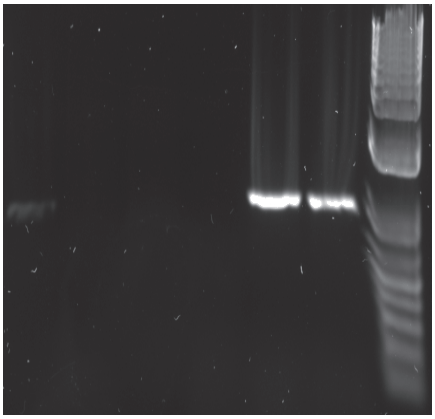

2 1 0.5 0.2
