## Supplementary Data 2 for "Systematic functional analysis of *Leishmania* protein kinases identifies regulators of differentiation or survival"

**Supplementary Data 2**. Localisation of mNeonGreen tagged protein kinases. An example image is shown for each protein kinase in the G1 phase of the cell cycle (1K 1N). Additional images are included for protein kinases that are stage specific or for which N- and C- terminal tagging differ. Images are ordered by the protein kinase family. Scale bar: 2 µm. Descriptors are based on those described in Halliday et al, Mol. Biochem. Parasitol. 230, 24–36 (2019).

### <u>AGC FAMILY</u>

#### LmxM.15.1550 (RAC1 – beta Serine/threonine kinase)

N-terminus

Primary localisation: mitochondrion

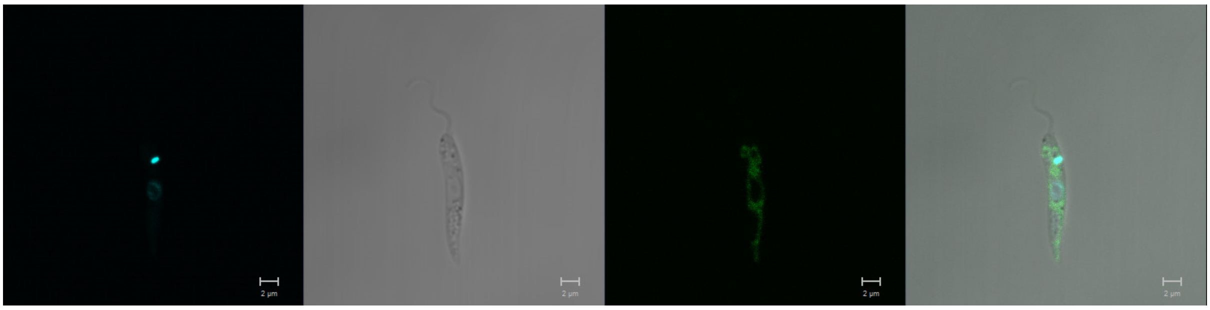

C-terminus

Primary localisation: mitochondrion

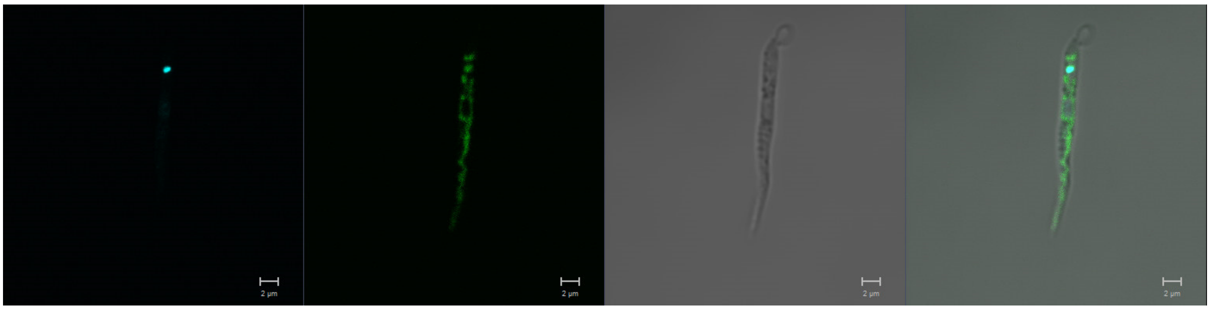

#### LmxM.25.2340 (AEK1, AGC essential kinase 1)

N-terminus

Primary localisation: cytoplasm

Secondary localisation: flagellum, basal body

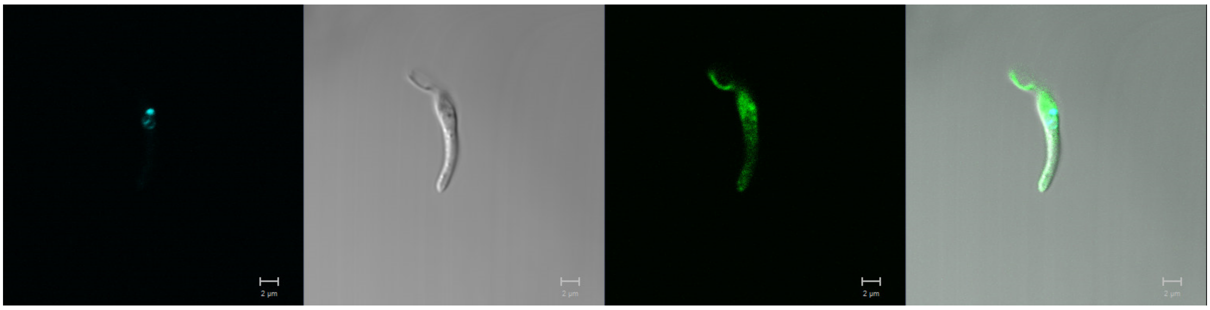

**LmxM.28.1670 (ZFK, differentiation inhibitory kinase)**
N-terminus

Primary localisation: lysosome

Secondary localisation: cytoplasm

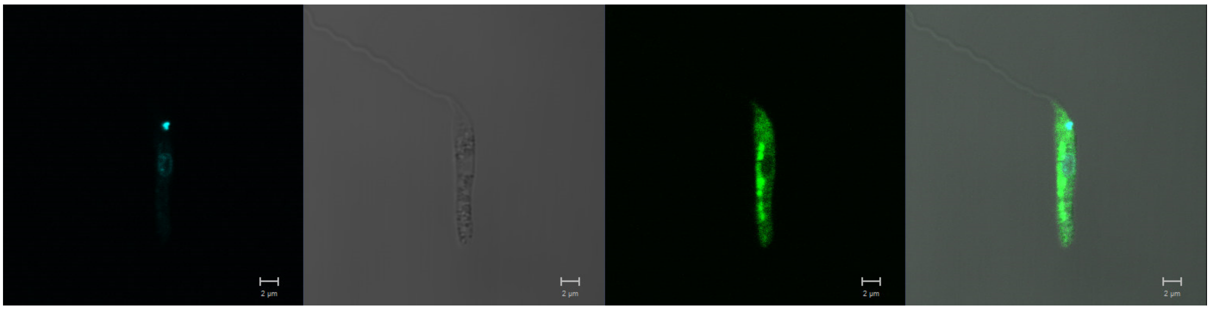

C-terminus

Primary localisation: endomembrane

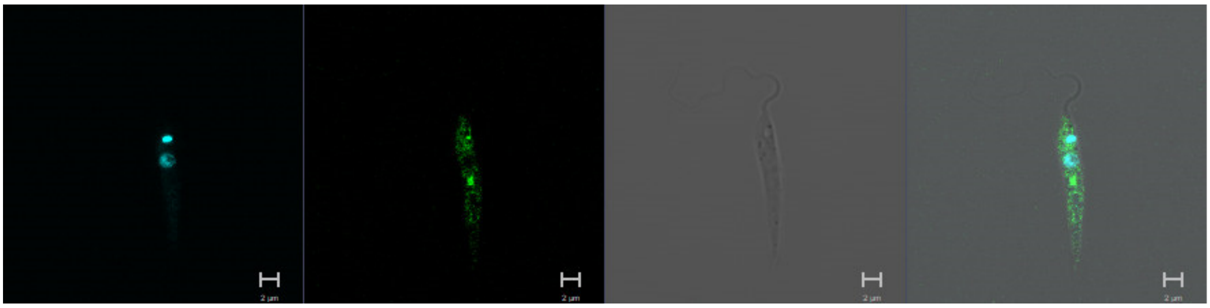

**LmxM.29.0800 (RAC2 serine-threonine kinase)**
N-terminus

Primary localisation: cytoplasm

Secondary localisation: flagellum, kinetoplast

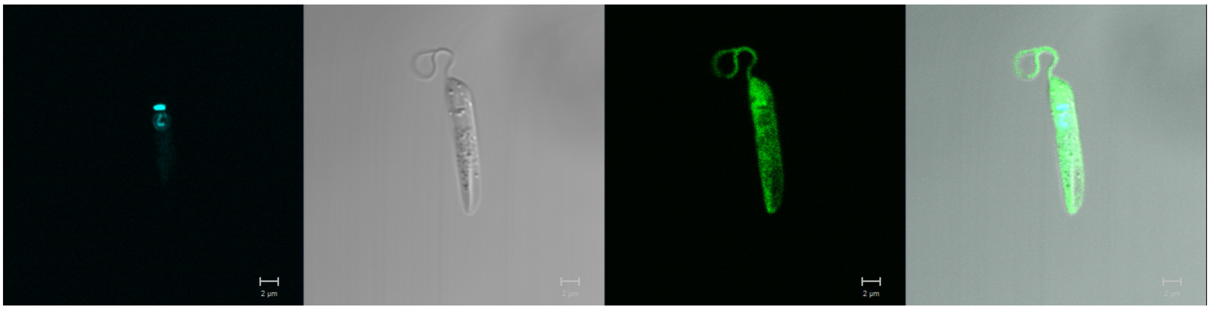

**LmxM.29.1000**
N-terminus

Primary localisation: cytoplasm

Secondary localisation: flagellum, nucleus, kinetoplast

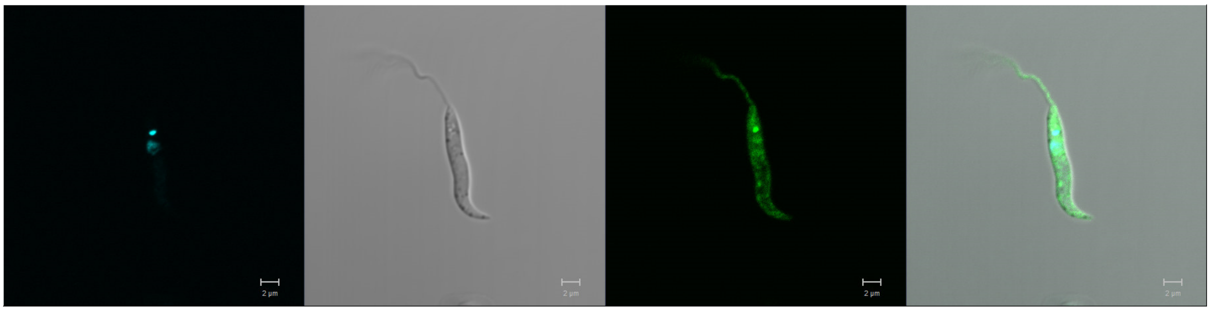

**LmxM.30.1530**
N-terminus

Primary localisation: cytoplasm

Secondary localisation: nucleus

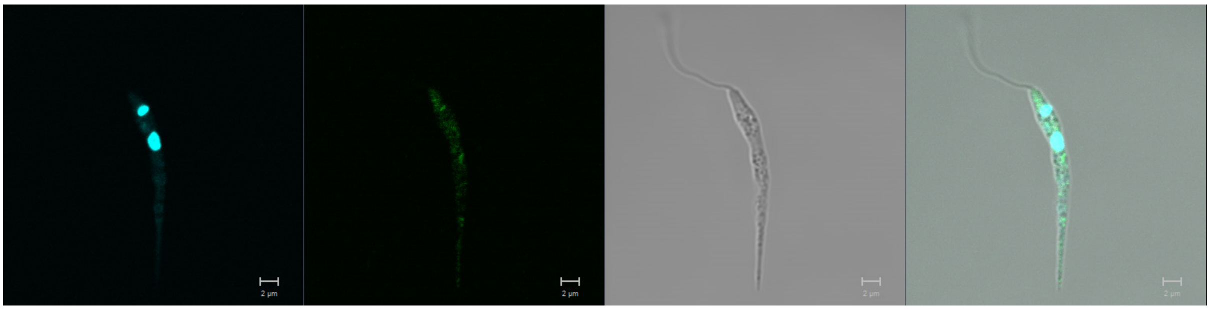

**LmxM.06.1180**
N-terminus

Primary localisation: cytoplasm

Secondary localisation: flagellum, nucleus

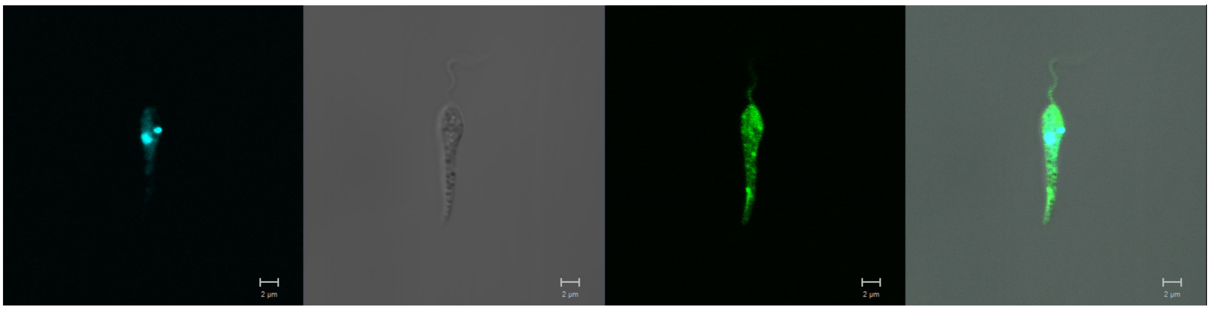

**LmxM.18.1080 (PKAC3, protein kinase A catalytic subunit 3)**
N-terminus

Primary localisation: pellicular membrane

Secondary localisation: cytoplasm, flagellum, nucleus

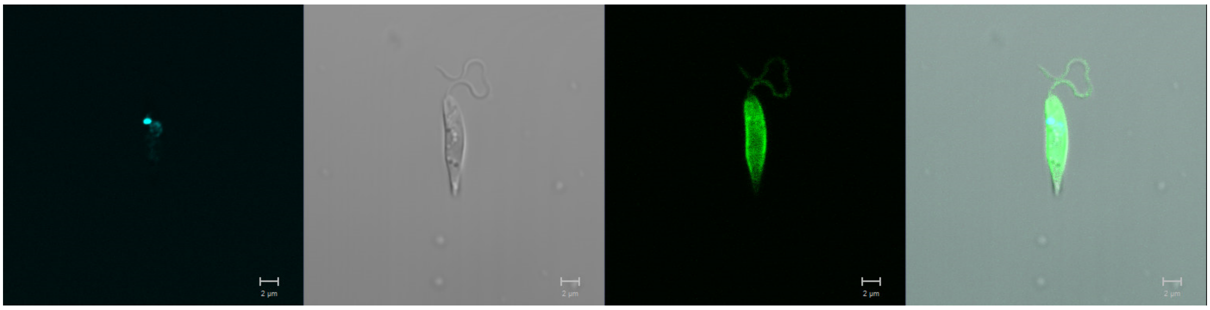

**LmxM.34.3960 (PKAC2, protein kinase A catalytic subunit isoform 2)**
N-terminus

Primary localisation: pellicular membrane

Secondary localisation: kinetoplast, cytoplasm, flagellum, nucleus

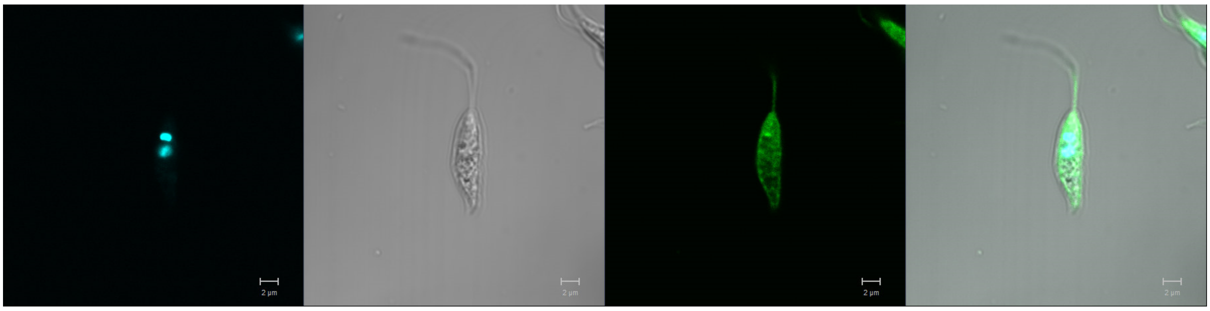

**LmxM.34.4010 (PKAC1, protein kinase A catalytic subunit isoform 1)**
N-terminus

Primary localisation: nucleus

Secondary localisation: cytoplasm (only during division)

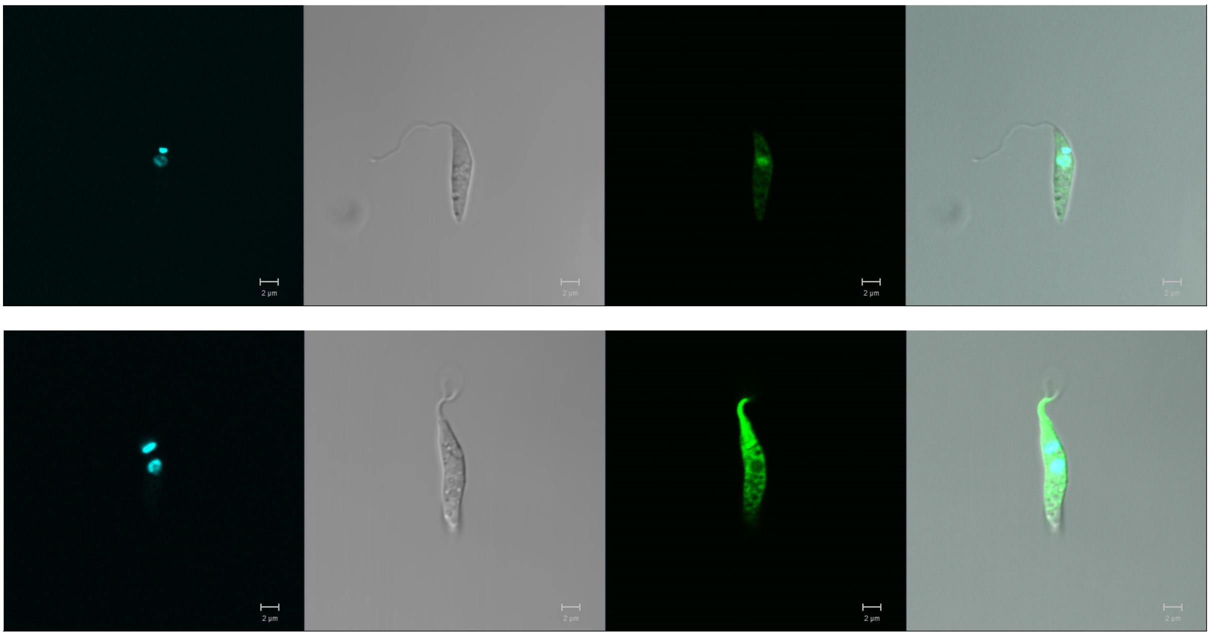

**LmxM.03.0780**
N-terminus

Primary localisation: cytoplasm

Secondary localisation: flagellum

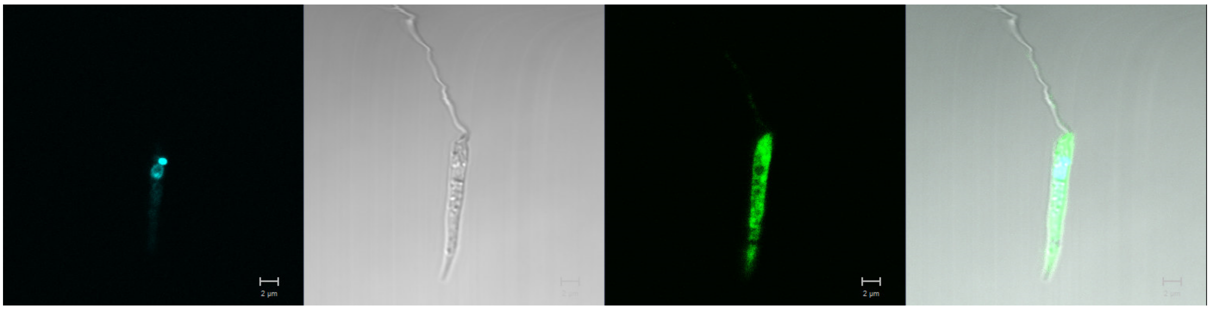

### CAMK FAMILY

**LmxM.27.2460**
N-terminus

Primary localisation: basal body

Secondary localisation: cytoplasm, flagellum, nucleus

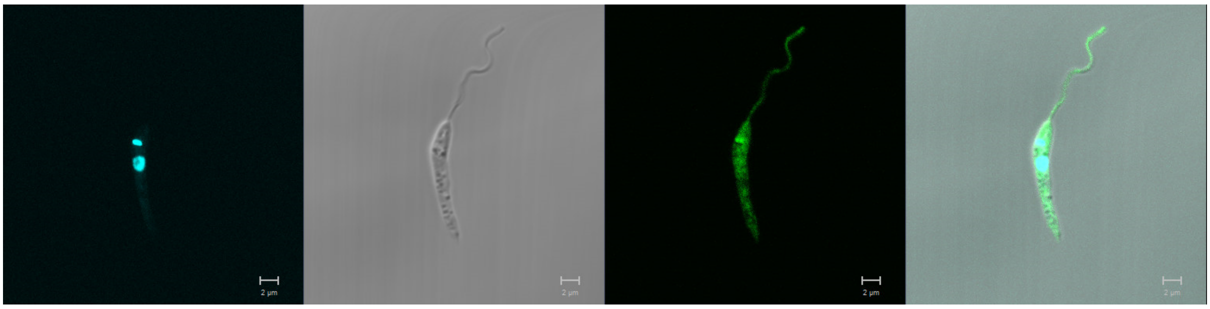

**LmxM.05.0130**
N-terminus

Primary localisation: nucleus

Secondary localisation: nucleus, cytoplasm, flagellum

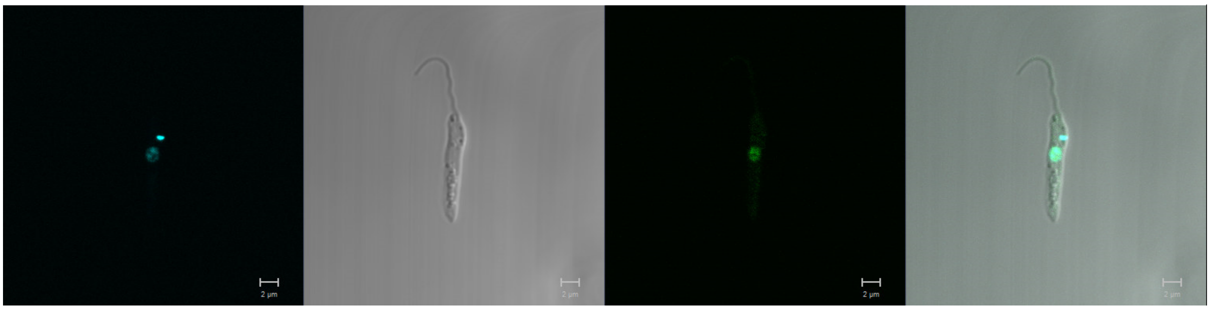

**LmxM.17.0060 (mitogen-activated protein kinase kinase 3 - MAPKK3)**
N-terminus

Primary localisation: cytoplasm

Secondary localisation: cytoplasm, flagellum, nucleus

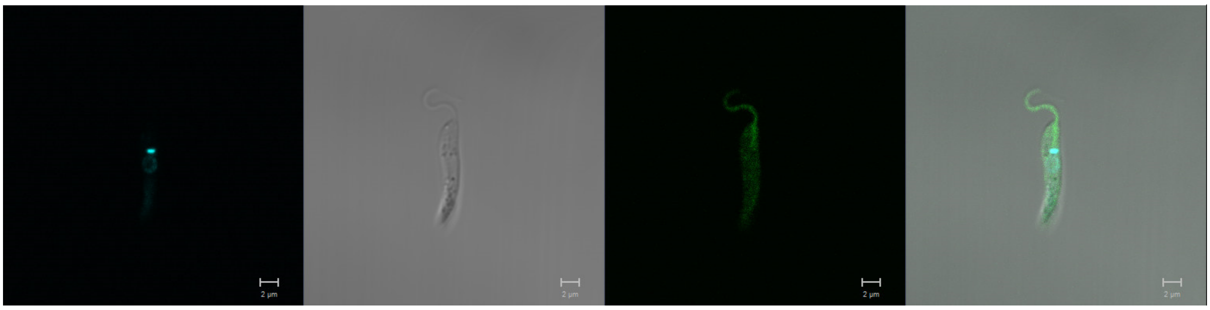

**LmxM.19.0140 (AKB1)**
N-terminus

Primary localisation: basal body

Secondary localisation: cytoplasm

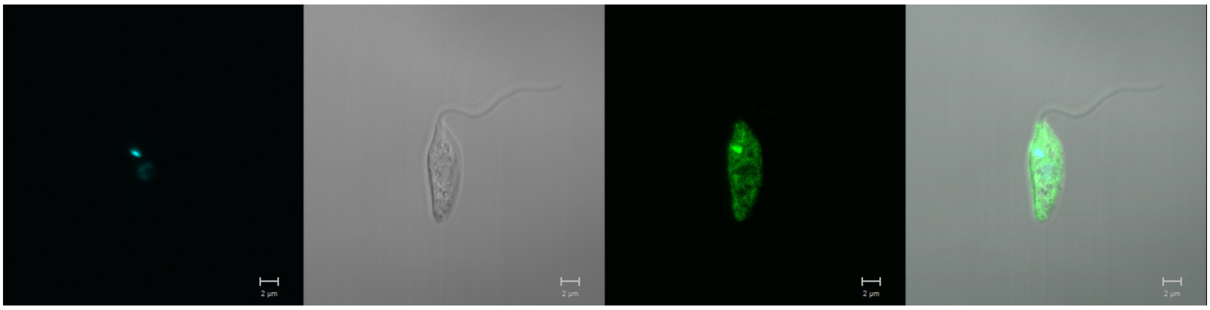

**LmxM.22.0810 (SOS2)**
N-terminus

Primary localisation: basal body

Secondary localisation: cytoplasm, flagellum, nucleus

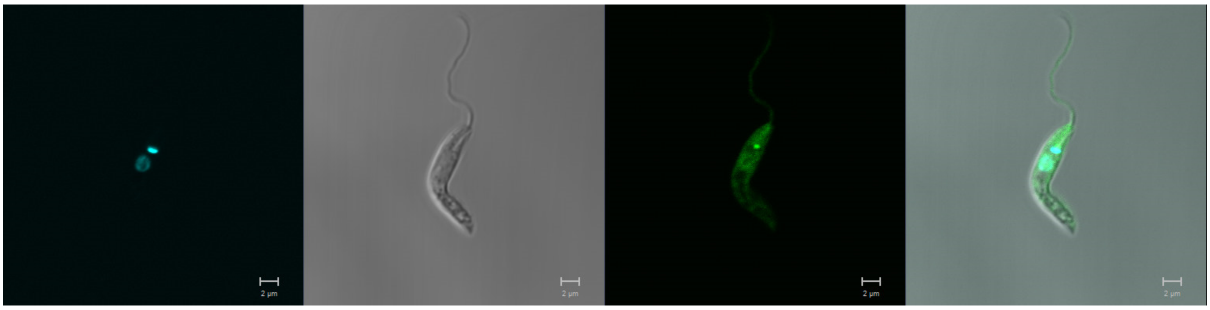

**LmxM.26.2510**
N-terminus

Primary localisation: cytoplasm

Secondary localisation: flagellum

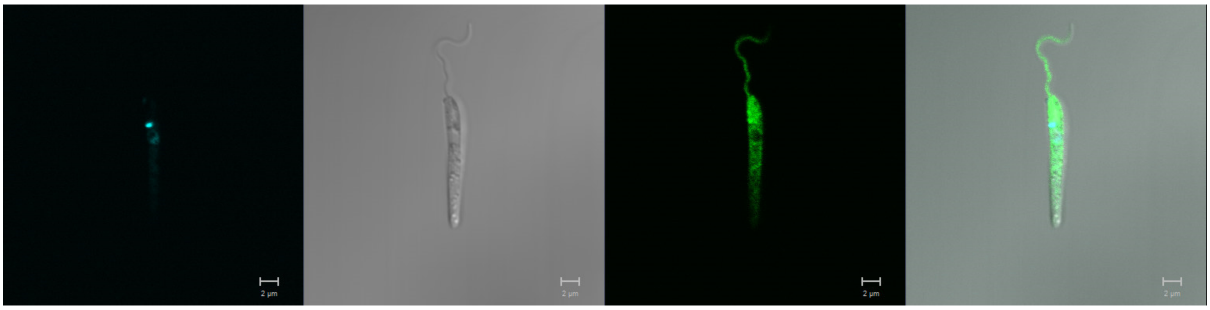

**LmxM.27.2470**
N-terminus

Primary localisation: basal body

Secondary localisation: cytoplasm, flagellum, nucleus

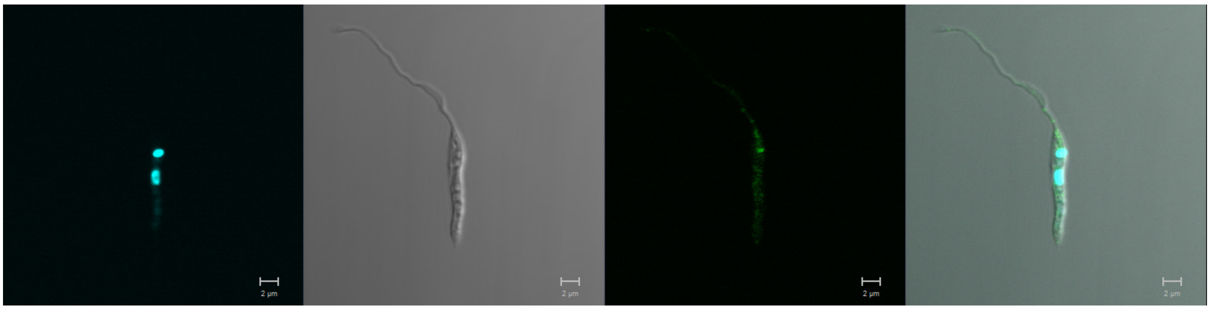

**LmxM.28.2000 (LDK)**
N-terminus

Primary localisation: cytoplasmic organelles (lipid droplet)

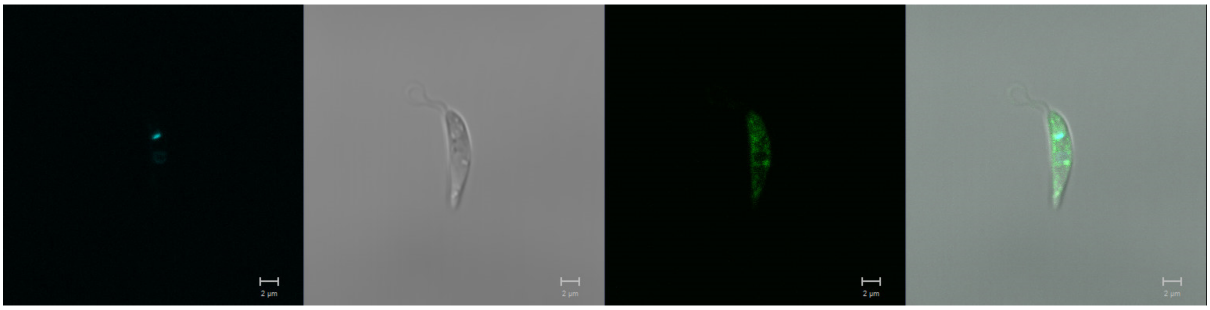

**LmxM.34.1050**
N-terminus

Primary localisation: cytoplasm

Secondary localisation: flagellum

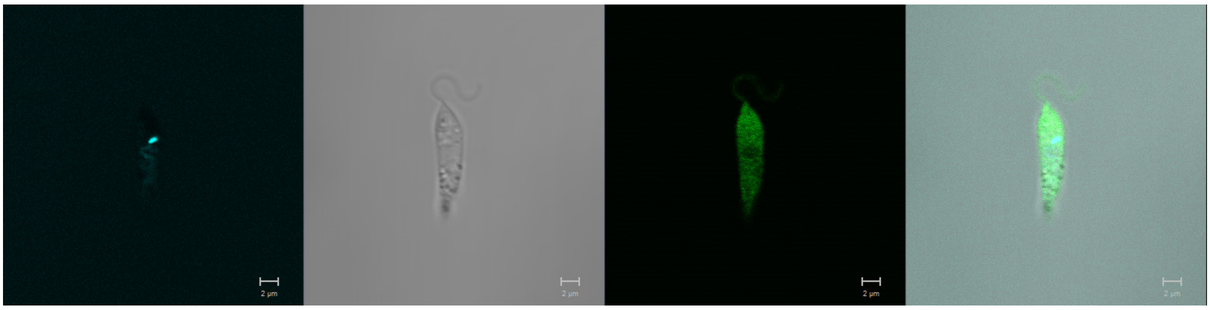

**LmxM.07.0900**
N-terminus

Primary localisation: basal body

Secondary localisation: cytoplasm, flagellum, basal body

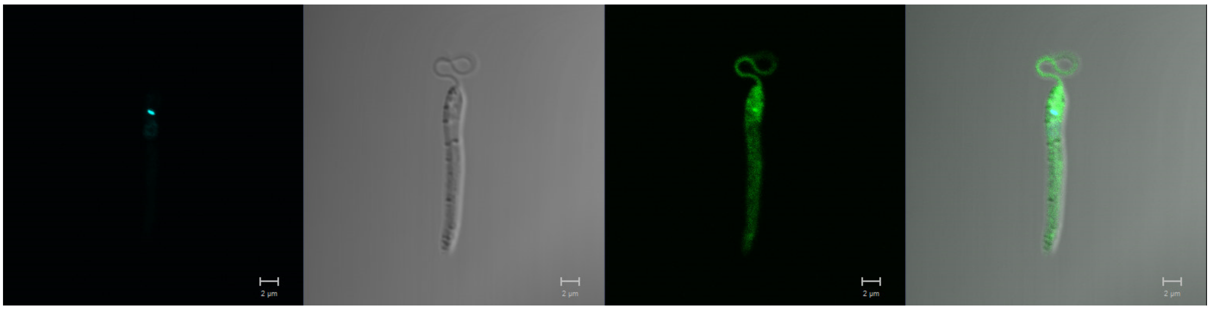

**LmxM.18.0640**
N-terminus

Primary localisation: basal body

Secondary localisation: cytoplasm, flagellum

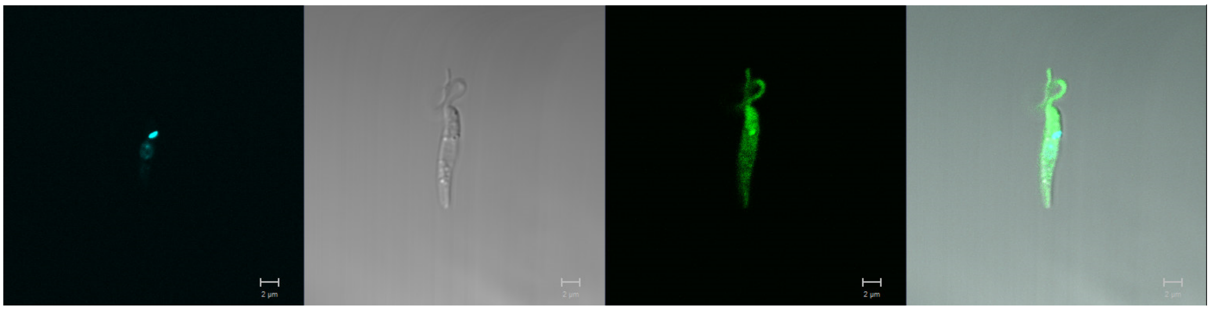

**LmxM.24.0230**
N-terminus

Primary localisation: nucleus

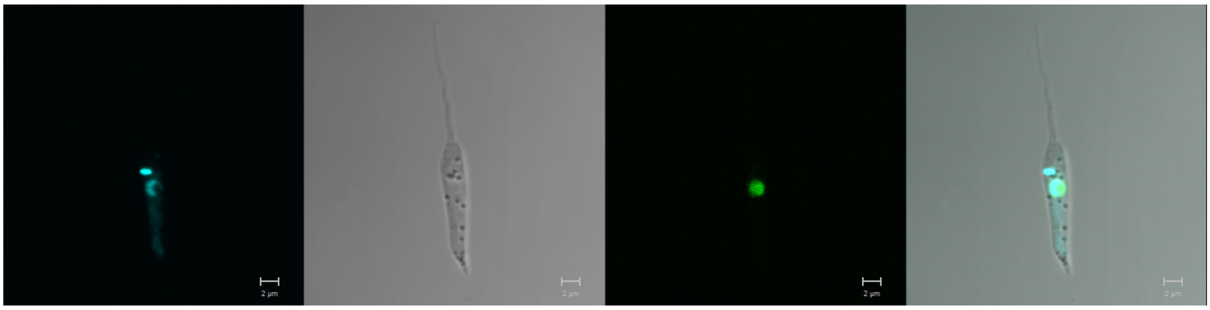

**LmxM.32.1710**
N-terminus

Primary localisation: cytoplasm

Secondary localisation: flagellum, nucleus, basal body

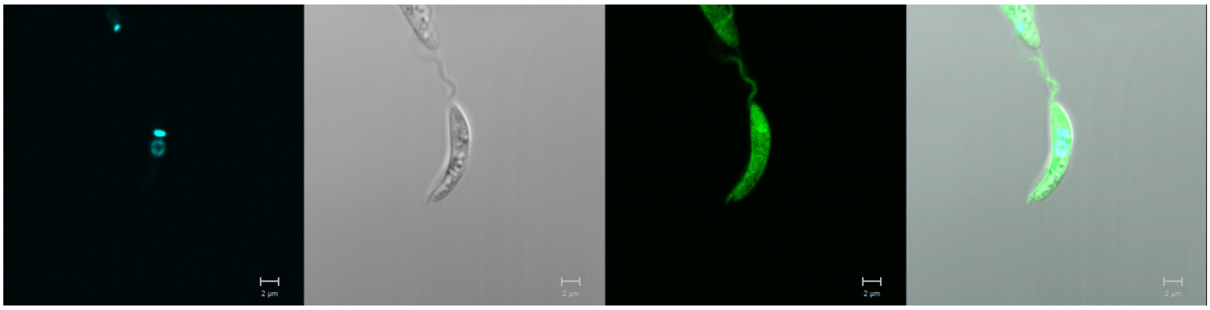

**LmxM.08_29.2020 (5'-AMP-activated protein kinase catalytic subunit alpha)**
N-terminus

Primary localisation: cytoplasm

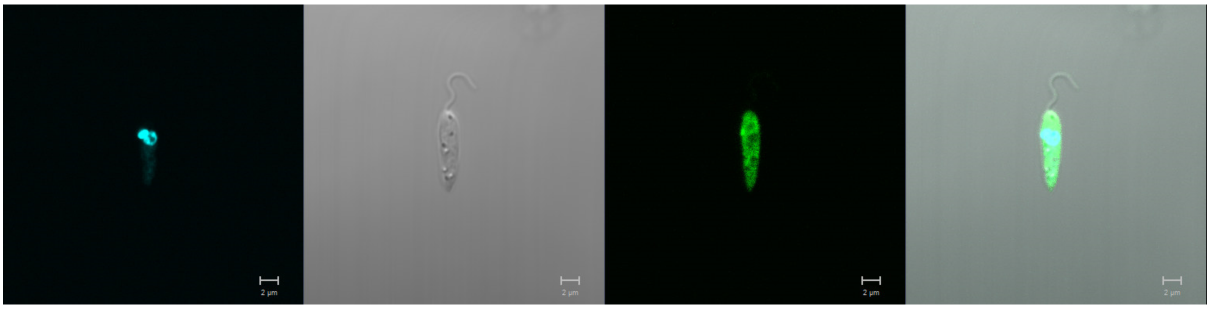

**LmxM.34.0490**
N-terminus

Primary localisation: cytoplasm

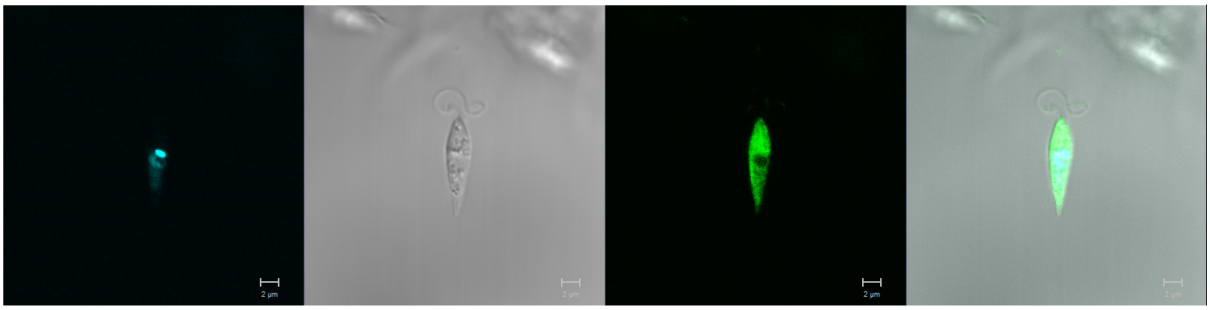

**LmxM.36.0900 (SNF1-related protein kinase)**
Missing – incomplete sequencing – cannot design primers.

**LmxM.21.0150**
N-terminus

Primary localisation: flagellar pocket

Secondary localisation: cytoplasm

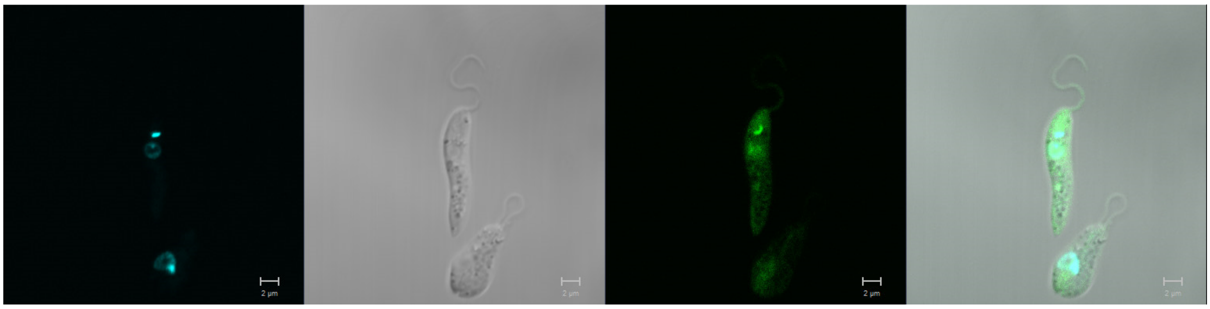

**LmxM.04.0650 (LUK1)**
N-terminus

Primary localisation: cytoplasm

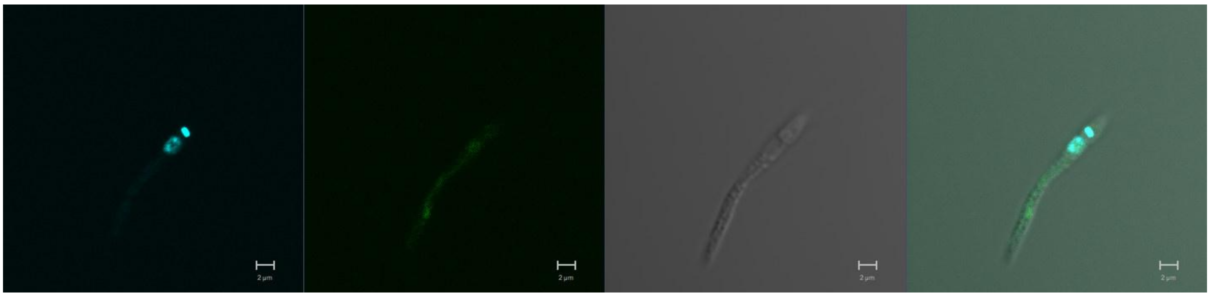

**LmxM.19.1470 (LUK3)**
N-terminus

Primary localisation: cytoplasm

Secondary localisation: basal body

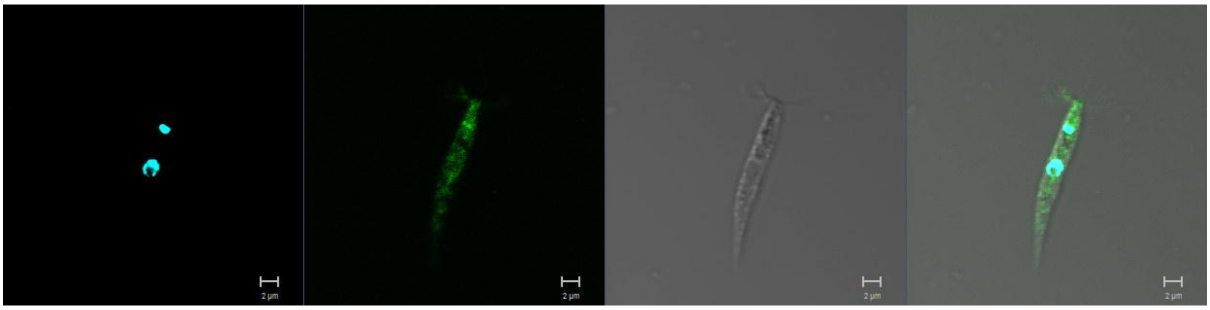

**LmxM.22.1170**
N-terminus

Primary localisation: basal body

Secondary localisation: flagellum, cytoplasm

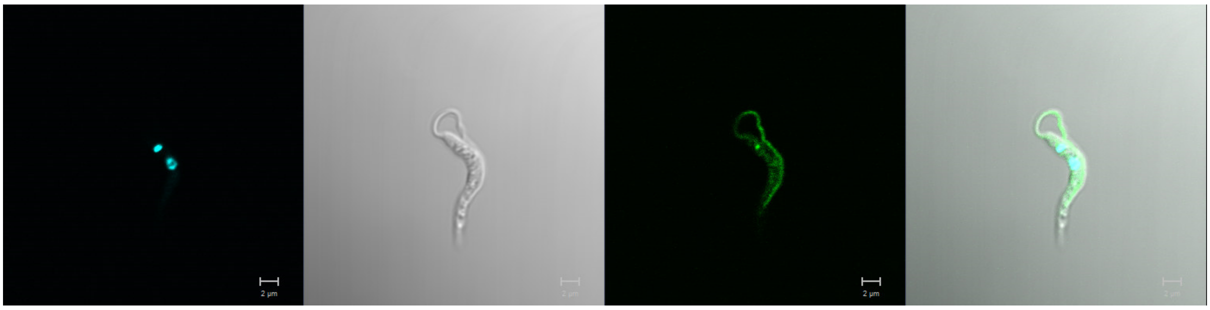

**LmxM.23.0490 (AMPKβ)**
N-terminus

Primary localisation: basal body

Secondary localisation: cytoplasm, basal body

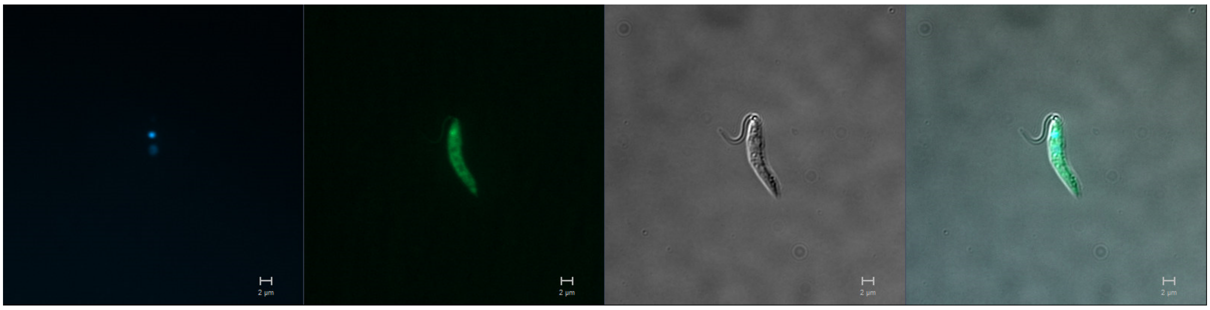

C-terminus

Primary localisation: cytoplasm

Secondary localisation: basal body

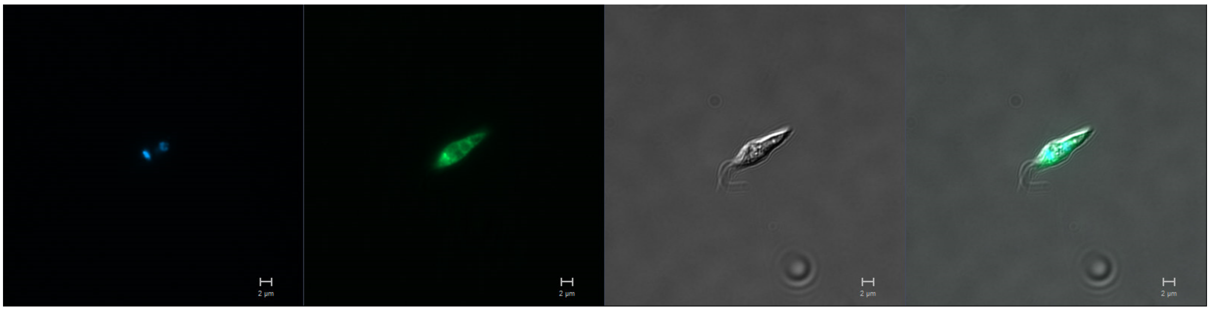

**LmxM.34.0760 (AMPKγ)**
N-terminus

Primary localisation: cytoplasm

Secondary localisation: basal body, flagellum

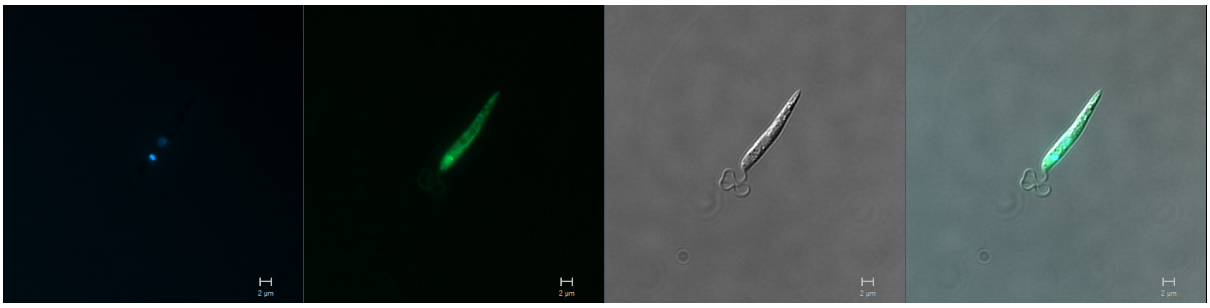

C-terminus

Primary localisation: cytoplasm

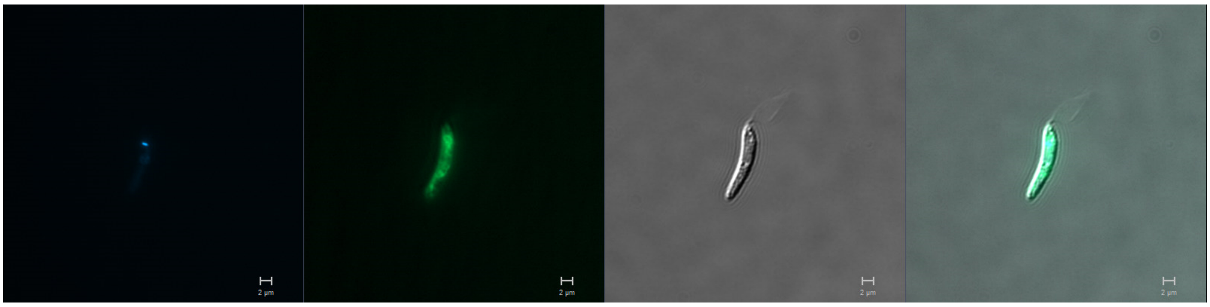

### CASEIN KINASE FAMILY

**LmxM.27.1780 (CK1.4)**
N-terminus

Primary localisation: flagellar pocket

Secondary localisation: cytoplasm

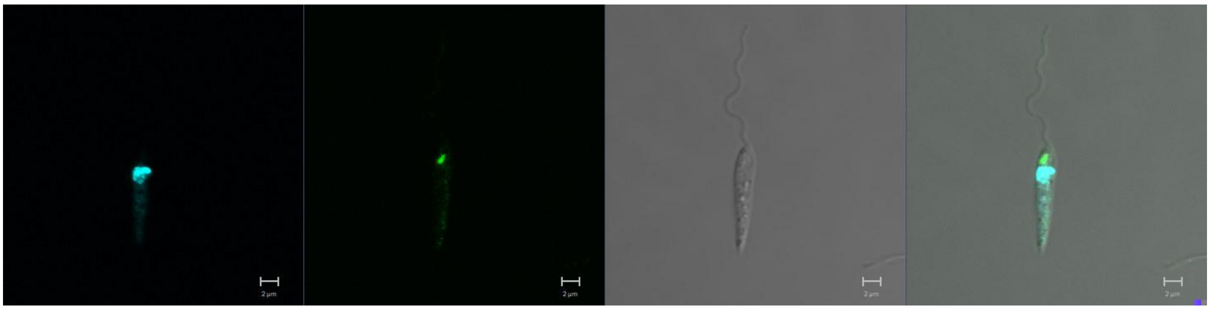

C-terminus

Primary localisation: cytoplasm

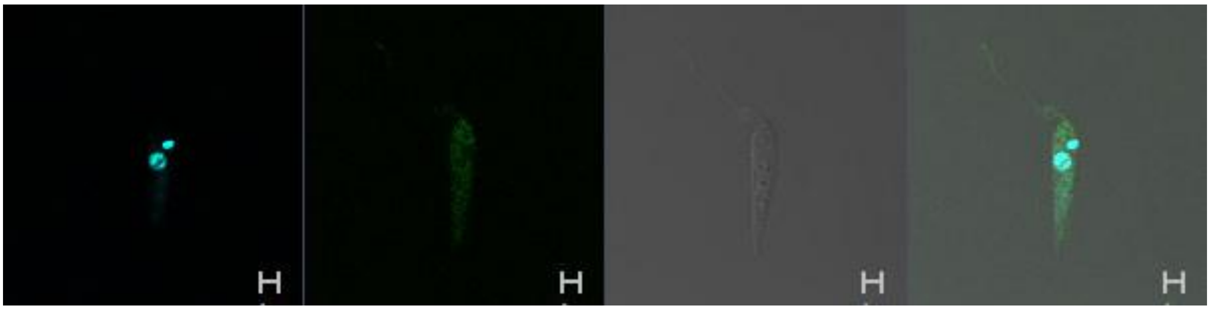

**LmxM.04.1210 (CK1.3)**
N-terminus

Primary localisation: no signal

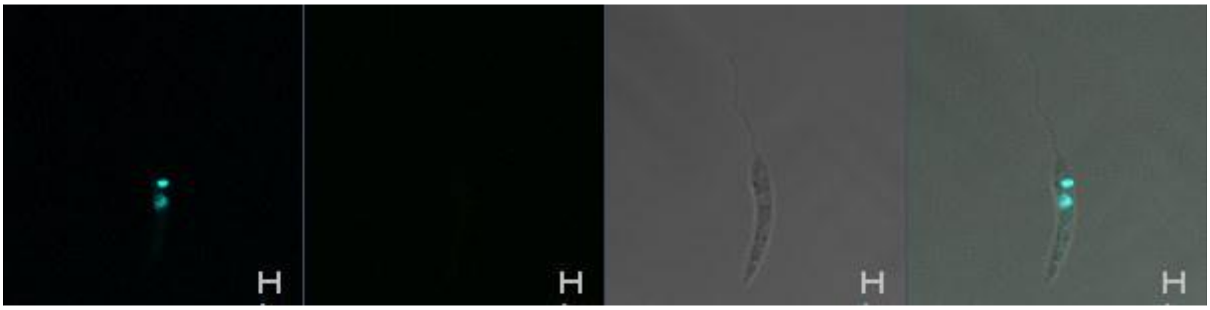

C-terminus

Primary localisation: cytoplasm

Secondary localisation: flagellum

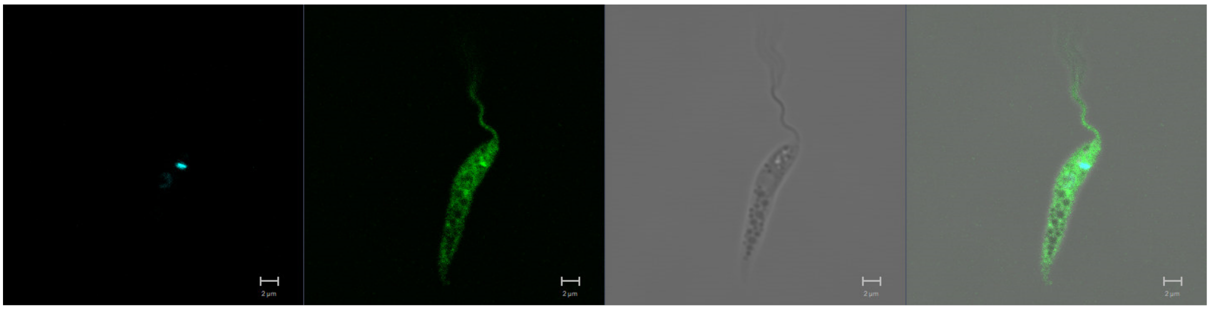

**LmxM.25.1580**
N-terminus

Primary localisation: flagellum

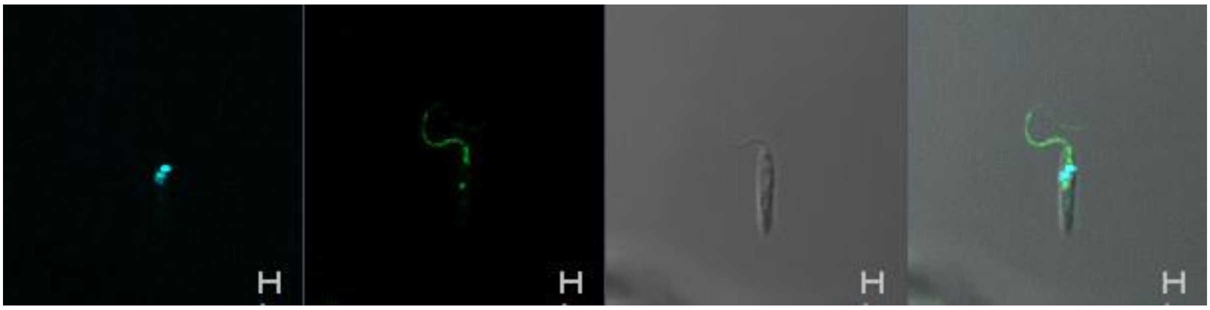

**LmxM.29.3470**
N-terminus

Primary localisation: cytoplasm

Secondary localisation: flagellum

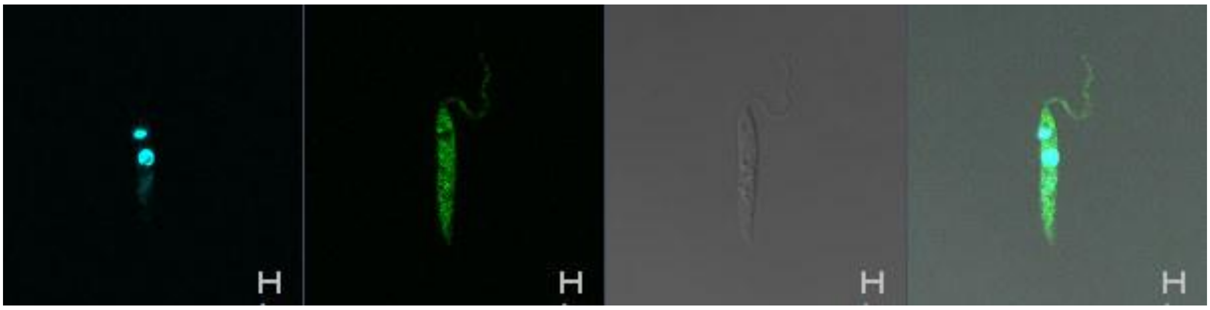

**LmxM.34.1000 (CK1.1)**
N-terminus

Primary localisation: cytoplasm

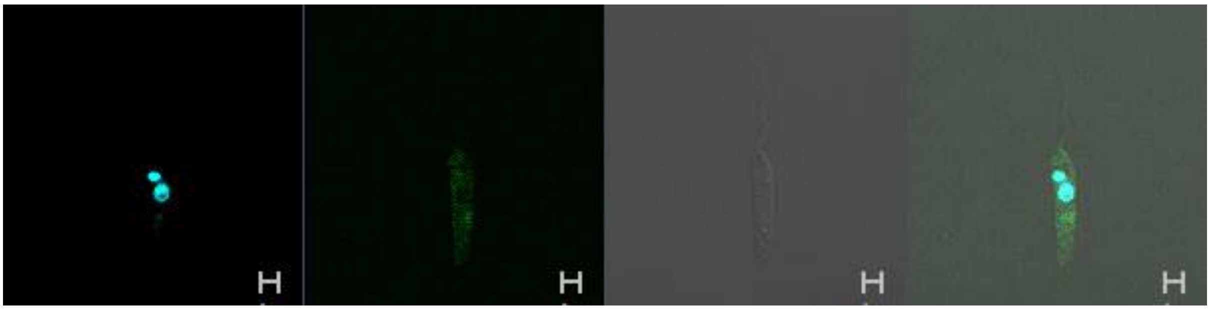

**LmxM.34.1010 (CK1.2)**
N-terminus

Primary localisation: cytoplasm

Secondary localisation: flagellum, basal body, nucleus

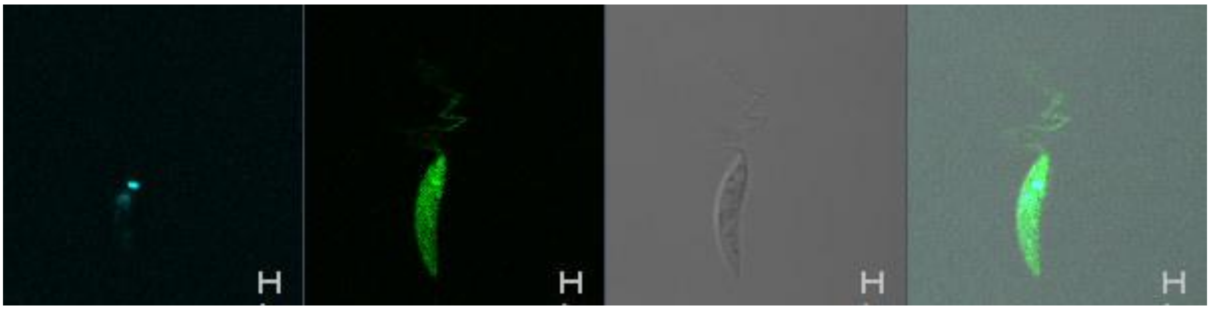

**LmxM.33.3020**
N-terminus

Primary localisation: flagellar pocket

Secondary localisation: cytoplasm

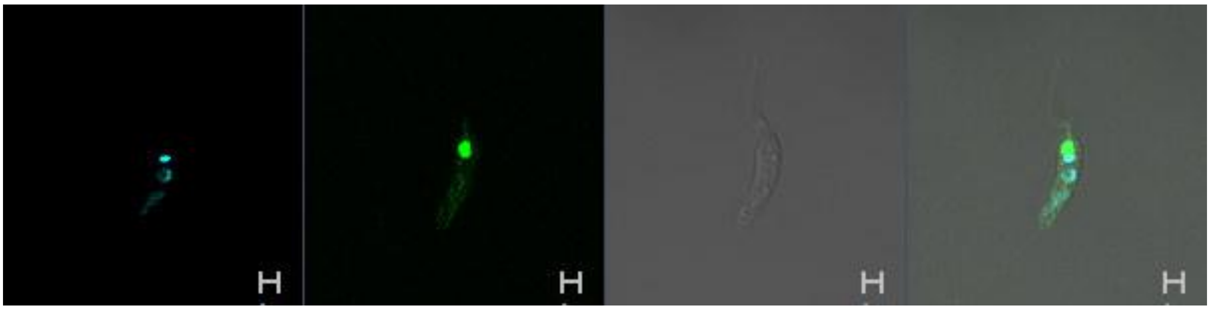

<u>**CMGC FAMILY**</u>

**LmxM.24.0670**
N-terminus

Primary localisation: nucleus

Secondary localisation: kinetoplast, cytoplasm

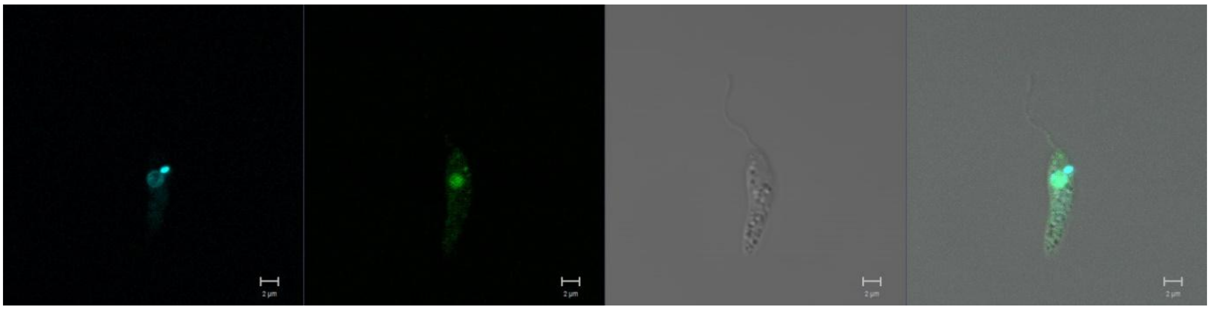

**LmxM.32.2070 (MPK15)**
N-terminus

Primary localisation: cytoplasm

Secondary localisation: cytoplasmic organelles, flagellum, nucleus

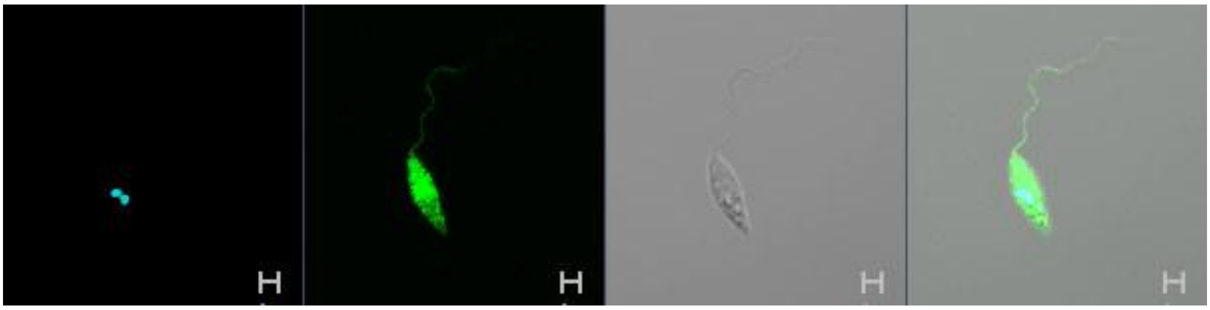

**LmxM.26.0040 (CRK7)**
N-terminus

Primary localisation: cytoplasm

Secondary localisation: flagellum, nucleus

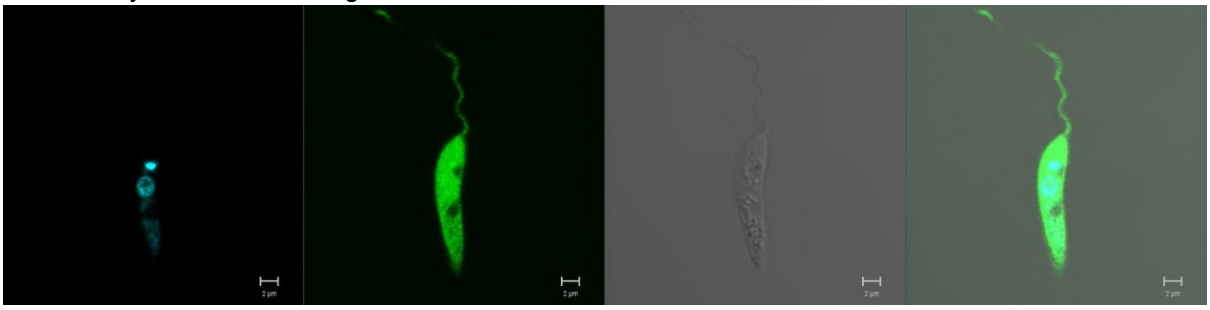

**LmxM.05.0550 (CRK2)**
N-terminus

Primary localisation: cytoplasm

Secondary localisation: nucleus, flagellum

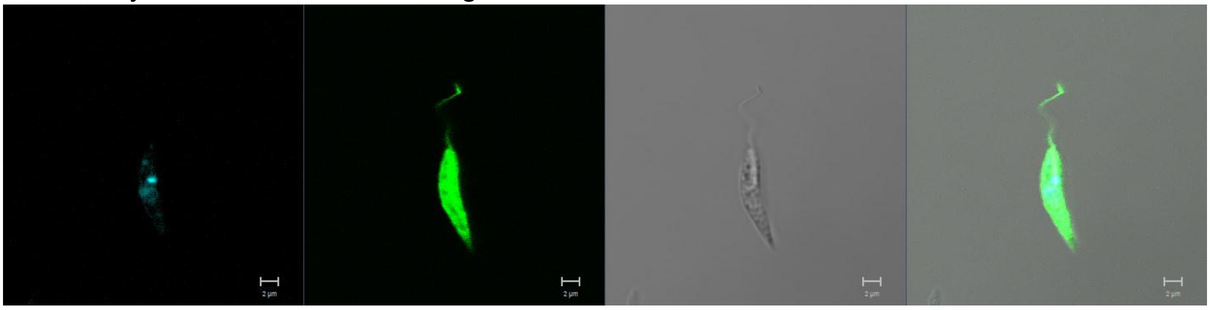

**LmxM.09.0310 (CRK12)**
N-terminus

Primary localisation: nucleus

Secondary localisation: kinetoplast, cytoplasm

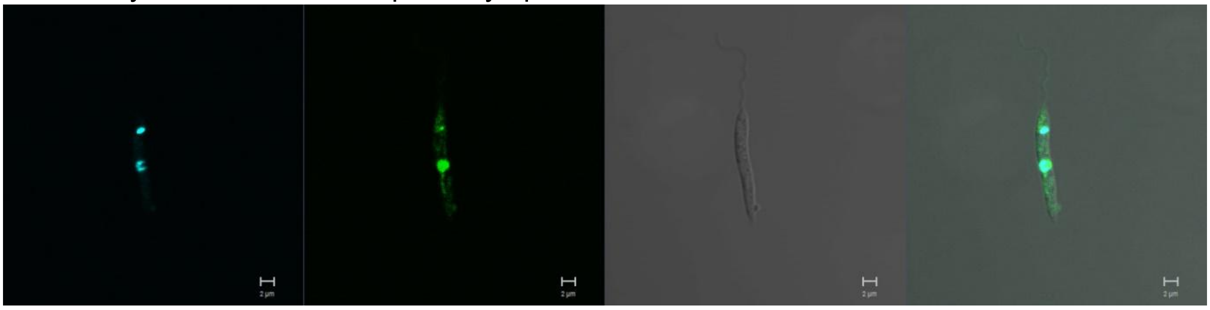

**LmxM.11.0110 (CRK8)**
N-terminus

Primary localisation: nucleus

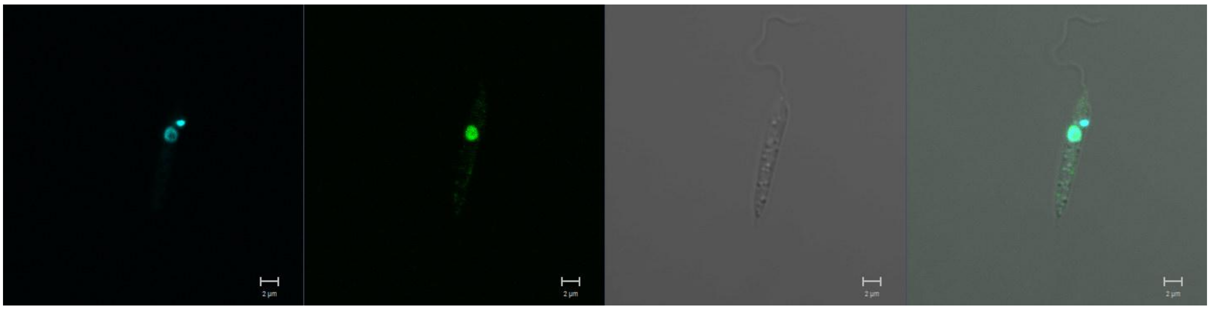

**LmxM.16.0990 (CRK4)**
N-terminus

Primary localisation: cytoplasm

Secondary localisation: flagellum

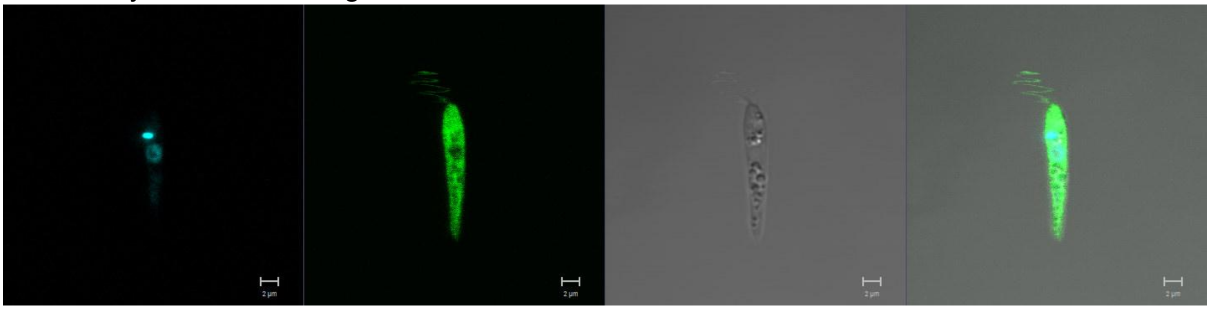

**LmxM.21.1080 (CRK1)**
N-terminus

Primary localisation: mitochondrion

Secondary localisation: cytoplasm

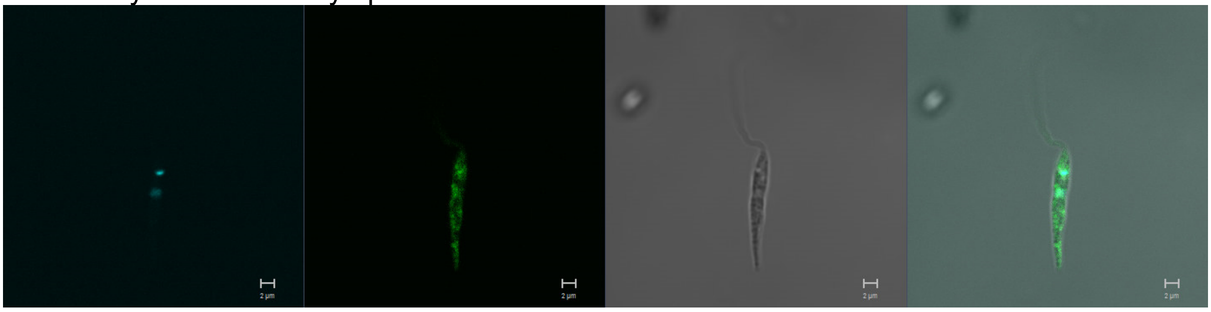

C-terminus

Primary localisation: mitochondrion

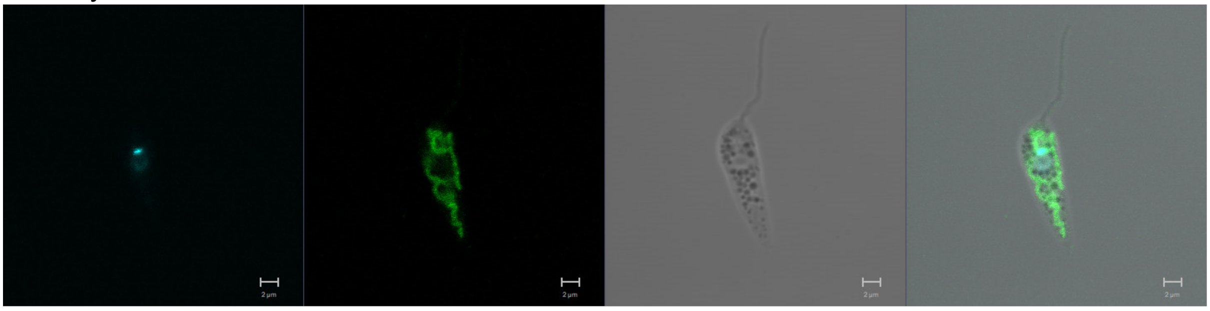

**LmxM.27.0560 (CRK6)**
N-terminus

Primary localisation: cytoplasm

Secondary localisation: flagellum, basal body

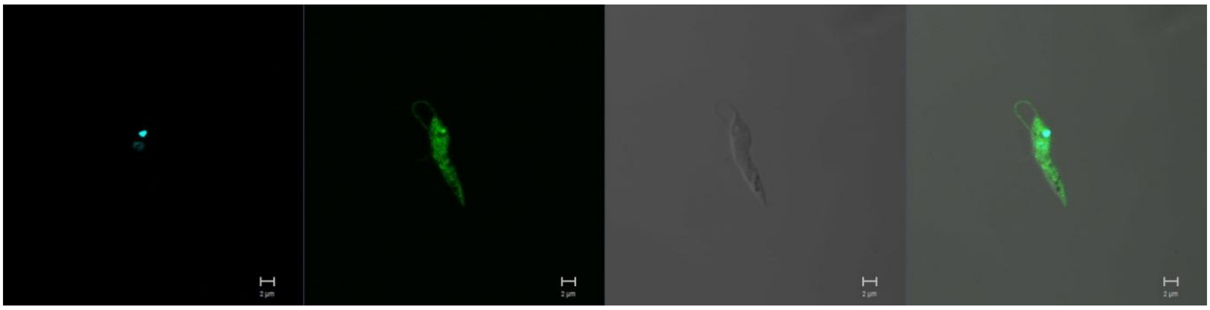

**LmxM.27.1940 (CRK9)**
C-terminus

Primary localisation: nucleus

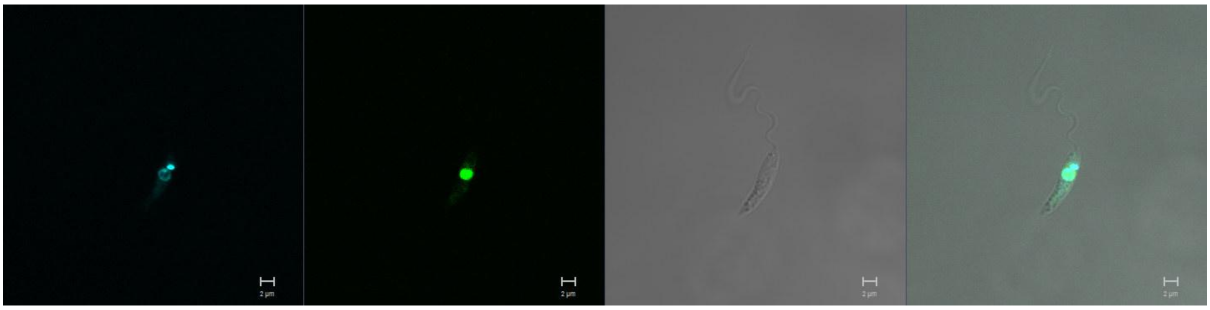

**LmxM.08_29.2150 (CRK10)**
N-terminus

Primary localisation: nucleus

Secondary localisation: kinetoplast, cytoplasm

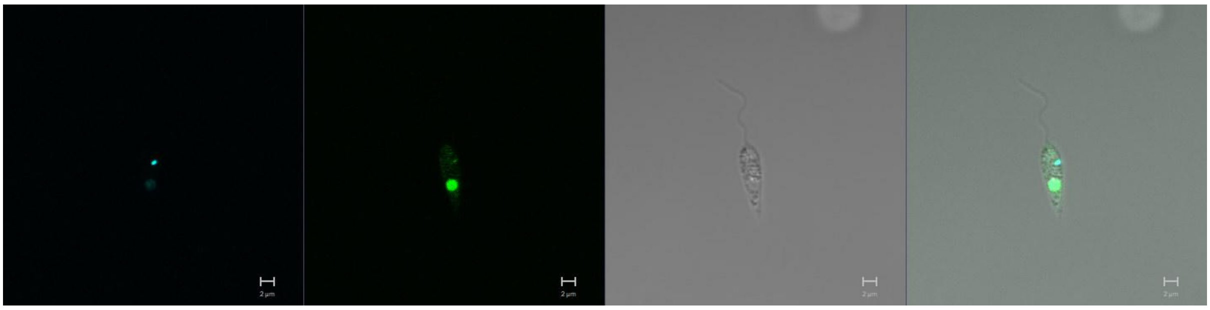

**LmxM.29.1780 (CRK11)**
C-terminus

Primary localisation: nucleus

Secondary localisation: cytoplasm

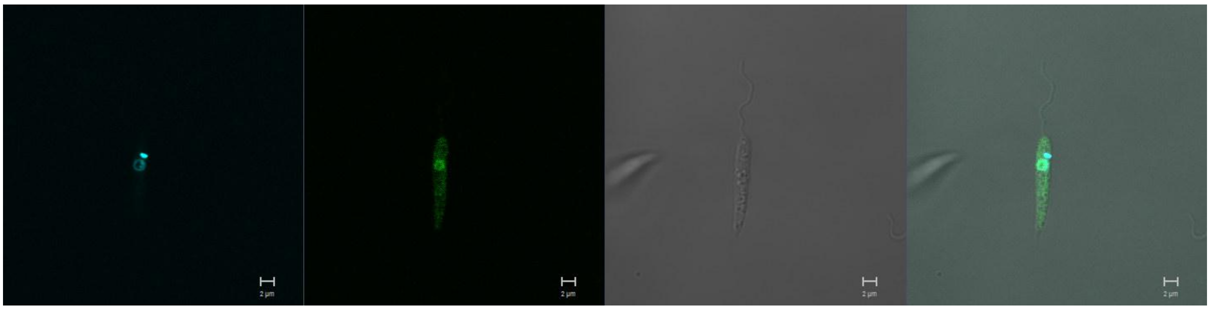

**LmxM.36.0550 (CRK3)**
N-terminus

Primary localisation: cytoplasm

Secondary localisation: flagellum, nucleus, kinetoplast

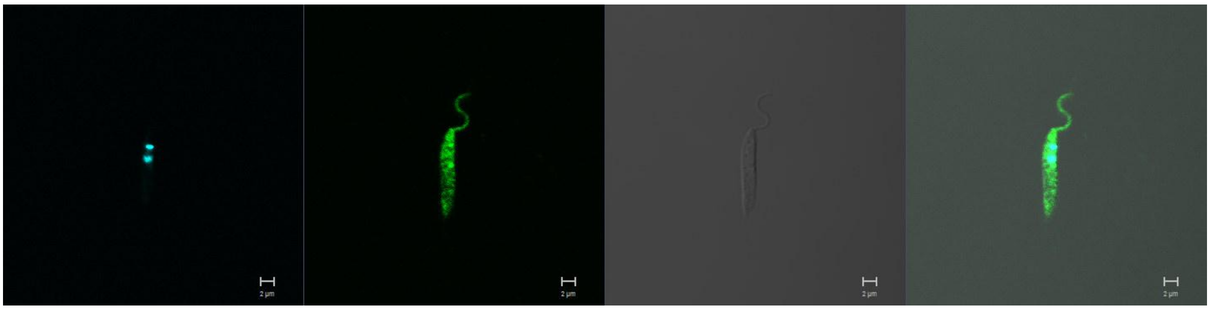

**LmxM.09.0400 (CLK1/KKT10)**
N-terminus

Primary localisation: nucleus

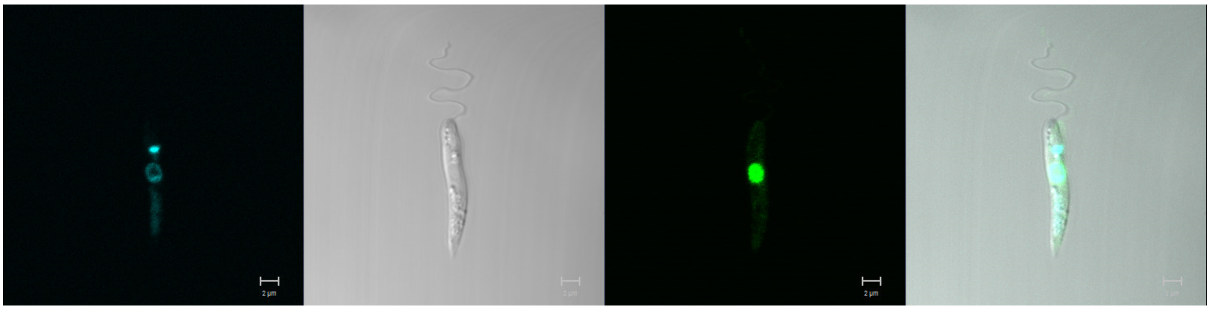

C-terminus

Primary localisation: nucleus

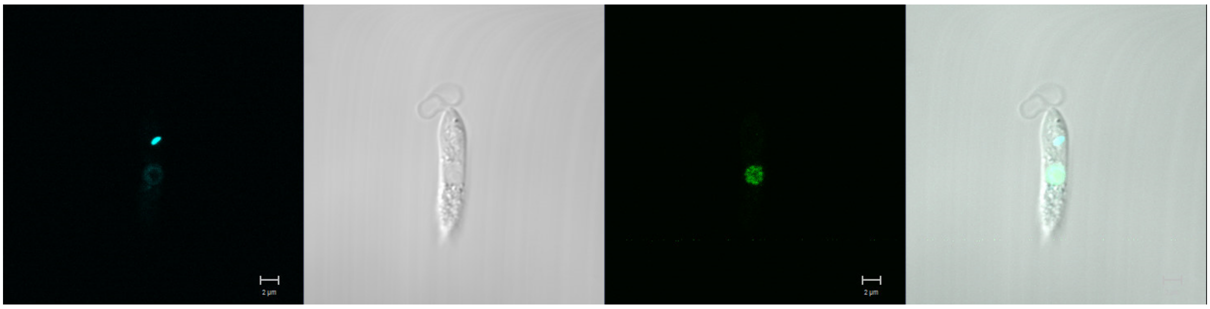

**LmxM.09.0410 (CLK2/KKT19)**
N-terminus

Primary localisation: nucleus

Secondary localisation: cytoplasm (only during division), kinetoplast

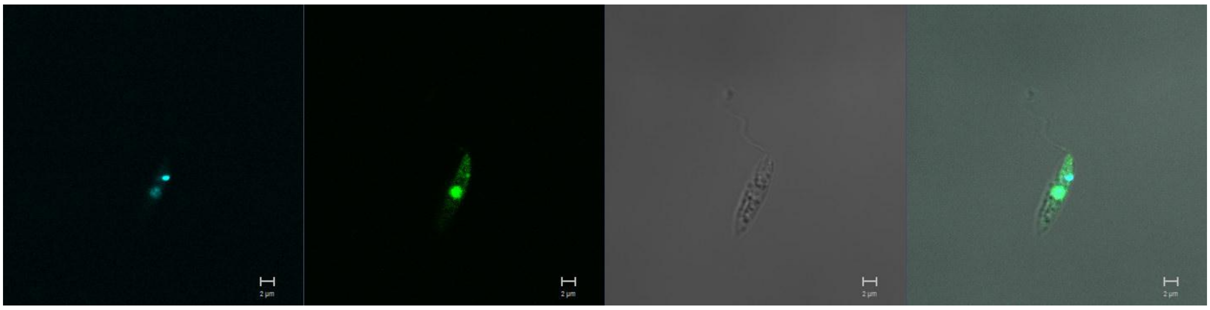

Field of view of dividing cells expressing CLK2/KKT19 in the cytoplasm (white arrows)

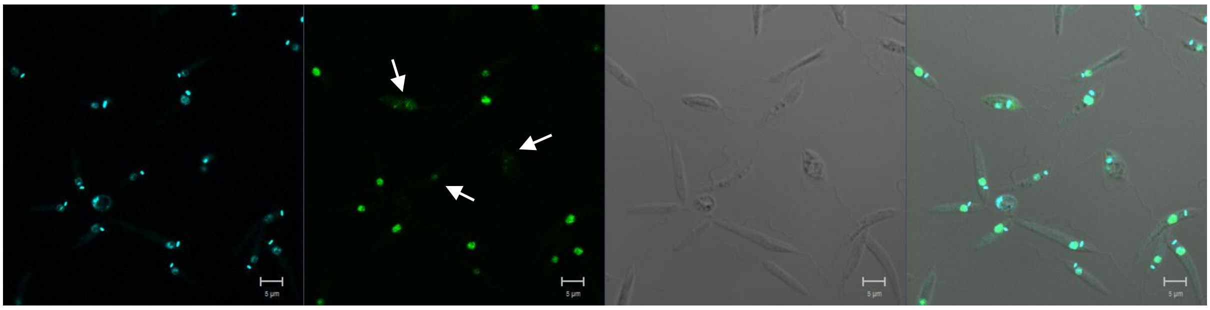

**LmxM.25.1560**
N-terminus

Primary localisation: cytoplasm

Secondary localisation: nucleus, kinetoplast

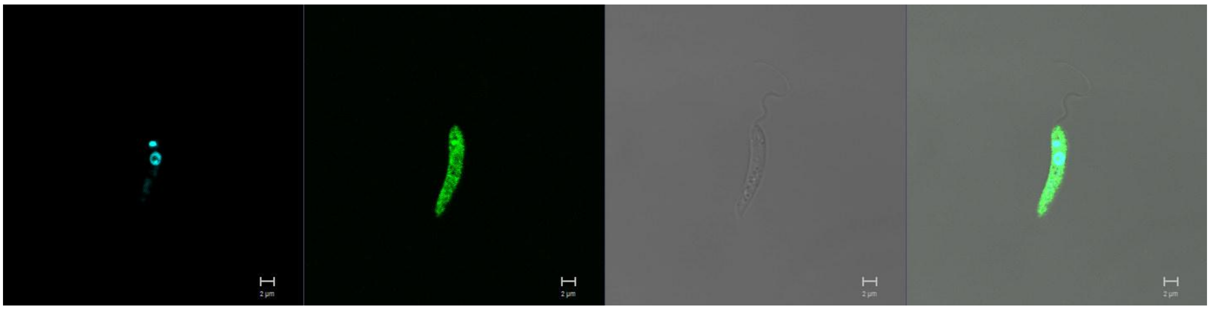

**LmxM.27.1800**
C-terminus

Primary localisation: endoplasmic reticulum

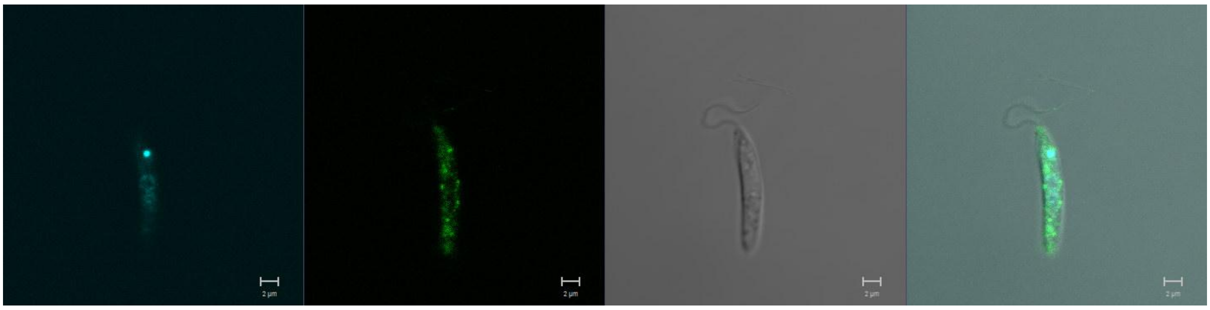

**LmxM.14.0830**
N-terminus

Primary localisation: basal body

Secondary localisation: cytoplasm

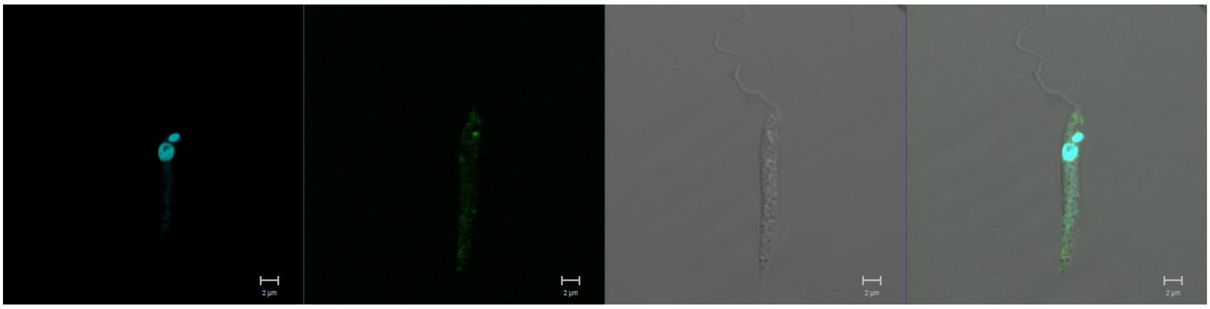

**LmxM.14.1070**
N-terminus

Primary localisation: basal body

Secondary localisation: flagellar pocket, lysosome, cytoplasm

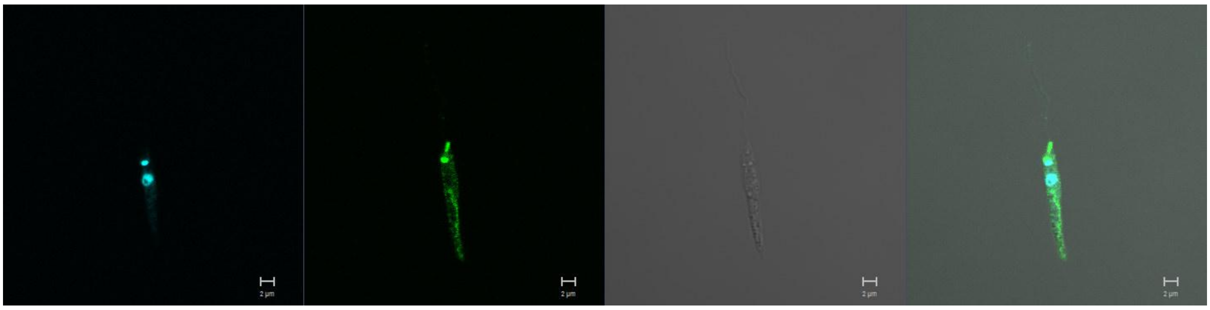

**LmxM.15.0180 (DYRK1)**
N-terminus

Primary localisation: lysosome

Secondary localisation: cytoplasm, endomembrane

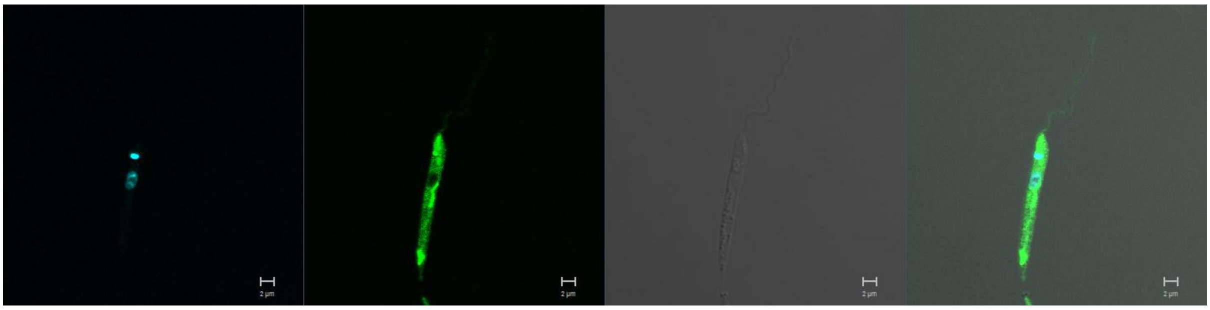

**LmxM.19.0360**
N-terminus

Primary localisation: cytoplasm

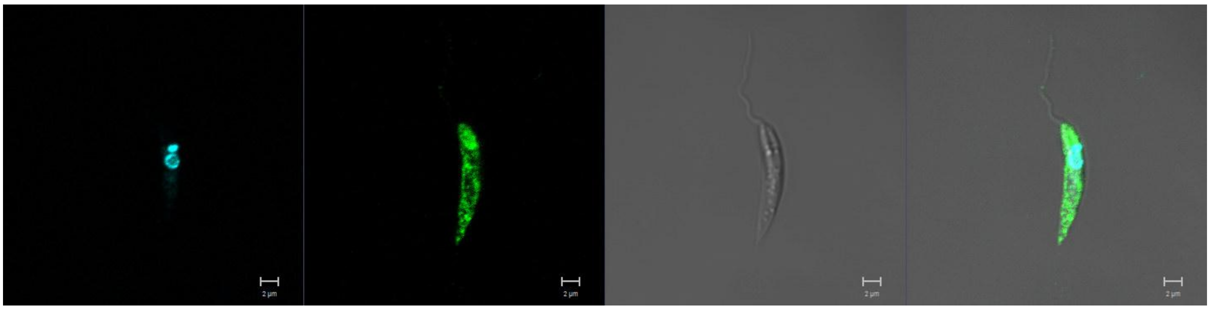

**LmxM.21.1650 (PK4)**
N-terminus

Primary localisation: cytoplasm (low expression)

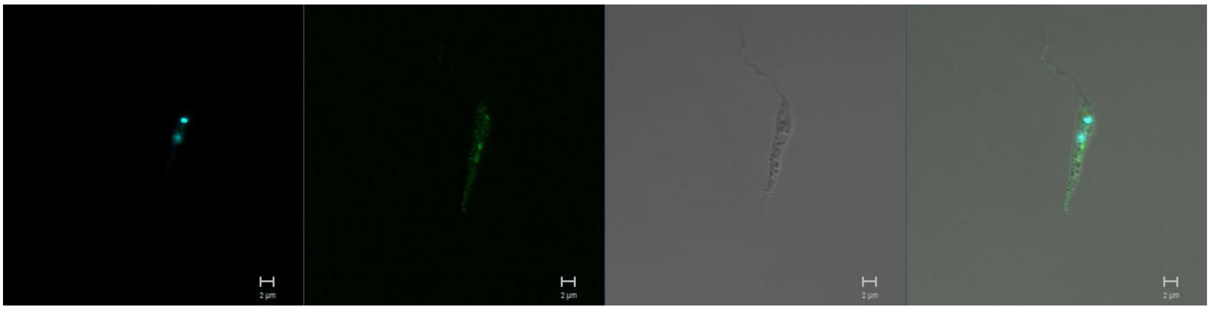

**LmxM.34.1860**
N-terminus

Primary localisation: flagellar pocket

Secondary localisation: basal body, cytoplasm, flagellum

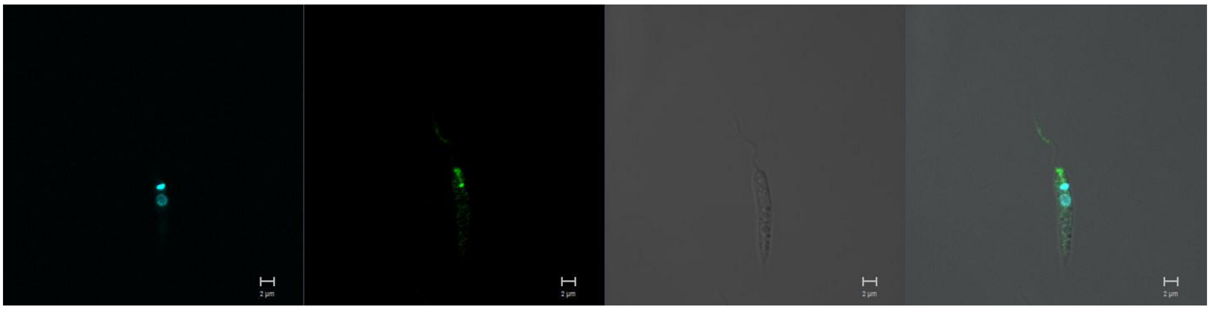

**LmxM.32.1830 (DYRK2)**
N-terminus

Primary localisation: cytoplasm (low expression)

Secondary localisation: flagellum (low expression)

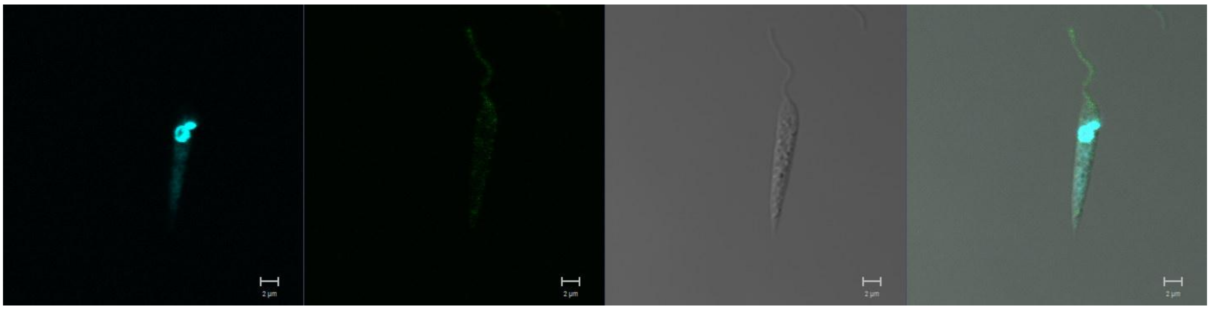

**LmxM.36.4250**
N-terminus

Primary localisation: no signal

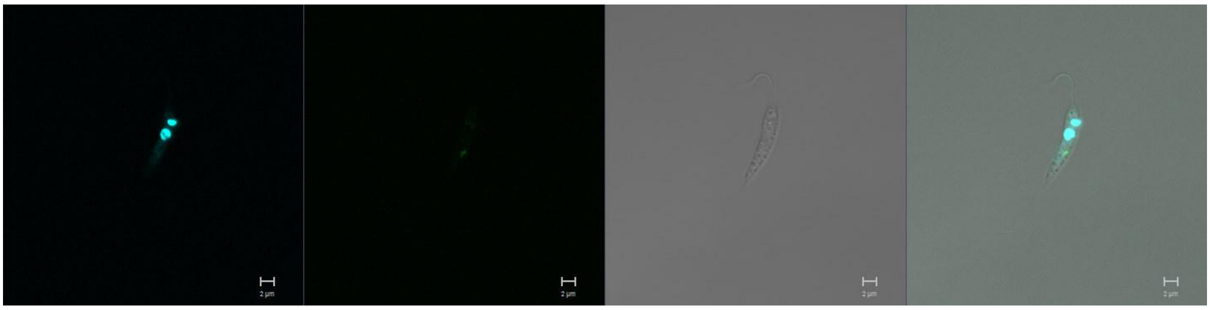

C-terminus

Primary localisation: no signal

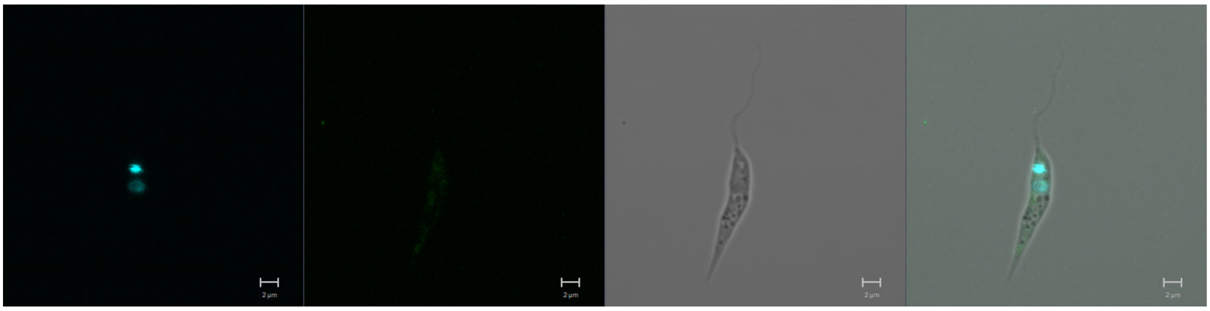

**LmxM.18.0270 (GSK3)**
N-terminus

Primary localisation: pellicular membrane

Secondary localisation: cytoplasm

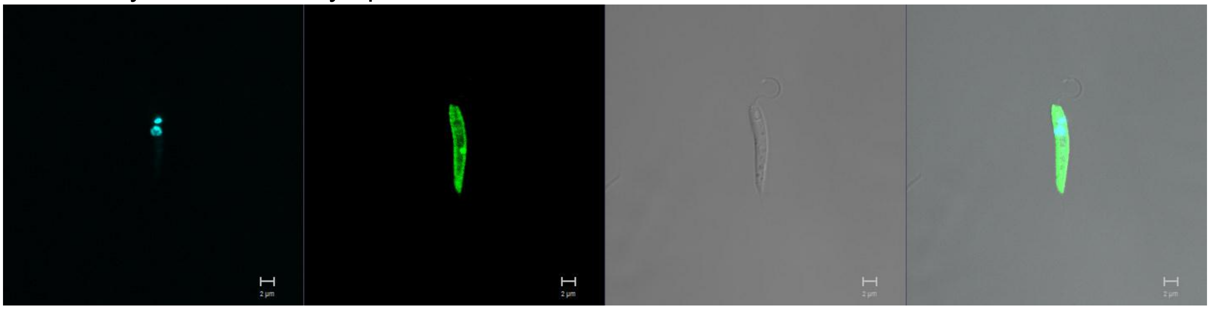

**LmxM.22.0490 (GSKA)**
N-terminus

Primary localisation: no signal

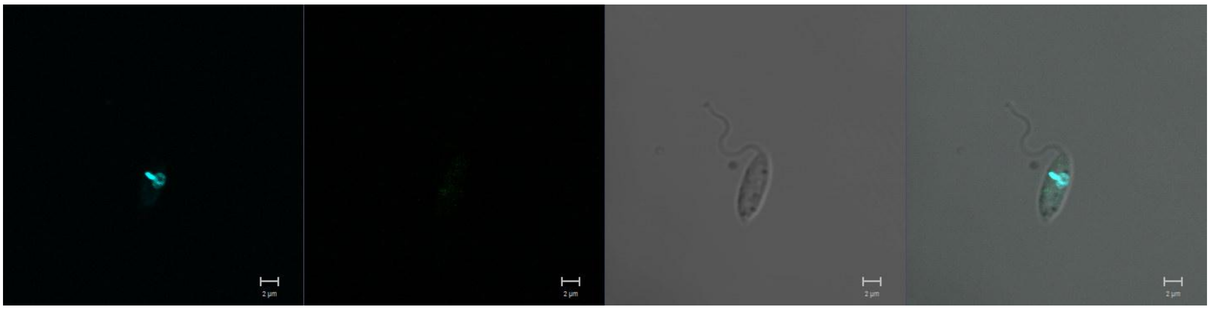

C-terminus

Primary localisation: no signal

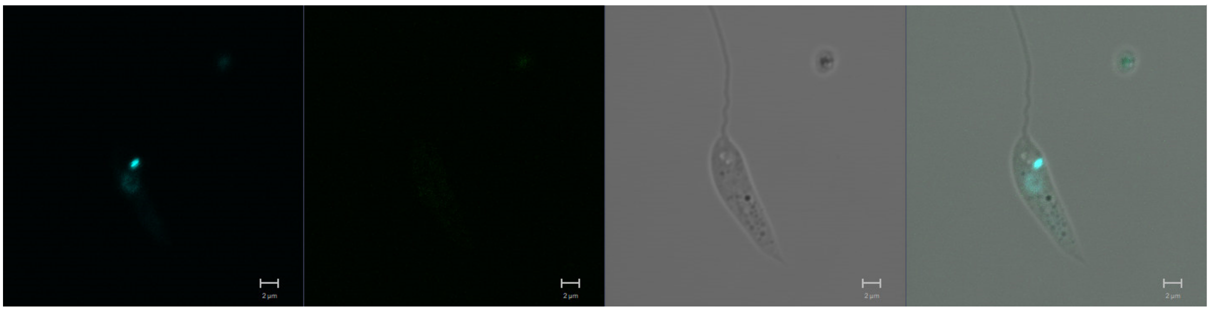

**LmxM.03.0210**
N-terminus

Primary localisation: cytoplasm

Secondary localisation: nucleus, flagellum

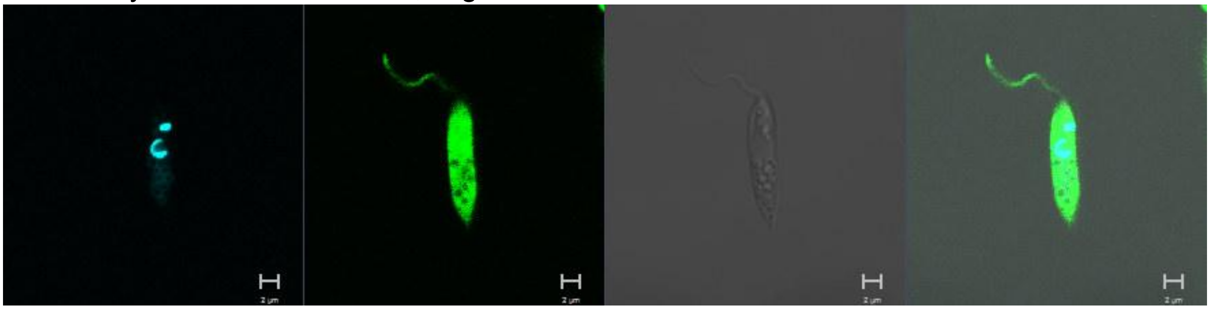

**LmxM.10.0200 (MPK10)**
N-terminus

Primary localisation: cytoplasm

Secondary localisation: flagellum, nucleus

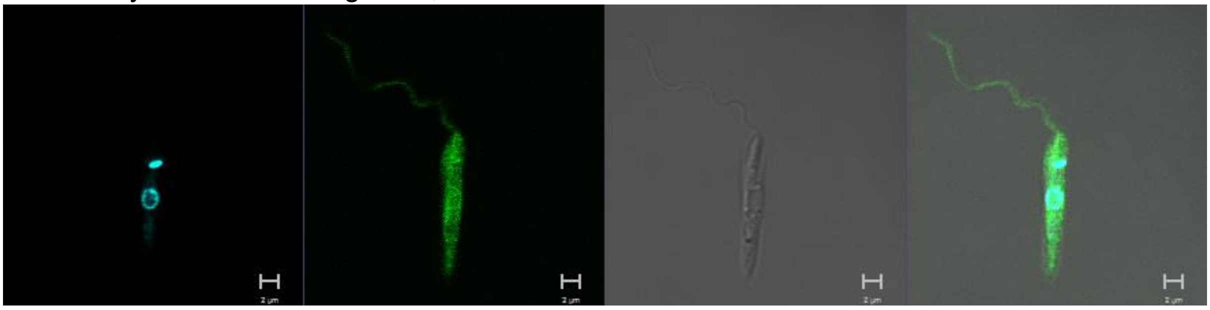

**LmxM.10.0490 (MPK3)**
N-terminus

Primary localisation: cytoplasm

Secondary localisation: flagellum

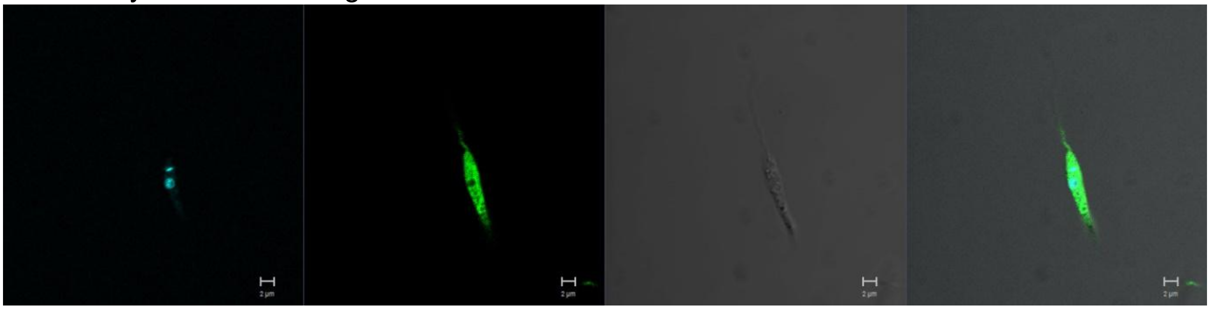

**LmxM.13.0780**
N-terminus

Primary localisation: cytoplasm

Secondary localisation: flagellum

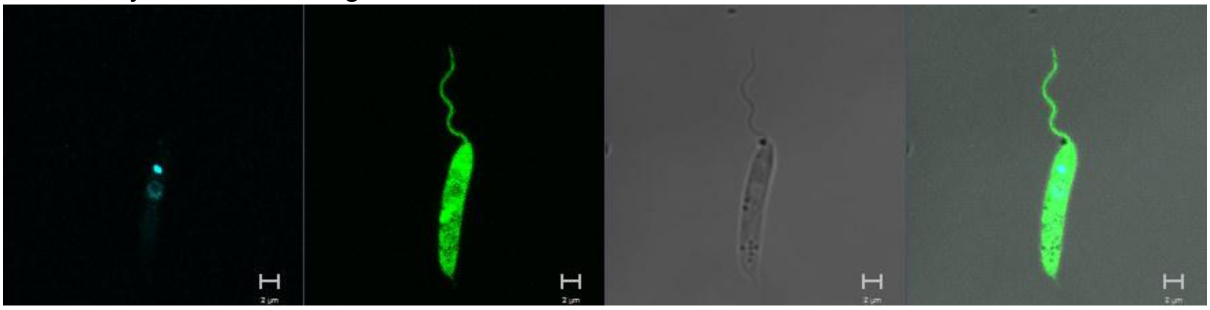

**LmxM.13.1640 (MPK7)**
N-terminus

Primary localisation: cytoplasm

Secondary localisation: nucleus, flagellum, basal body

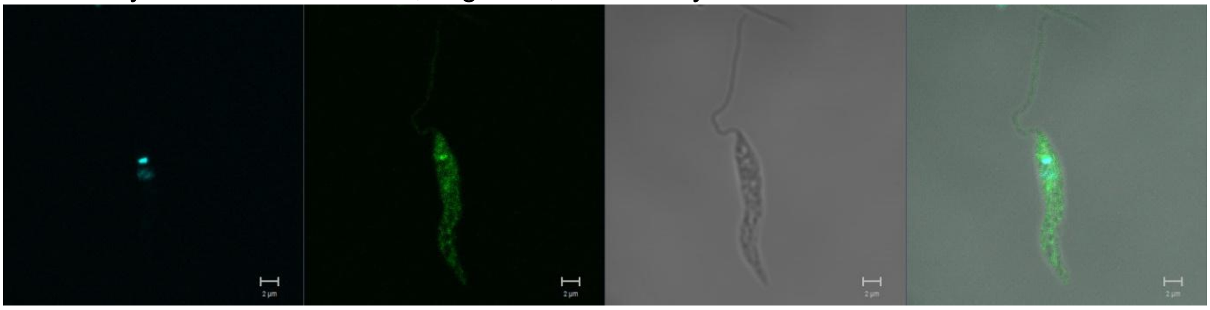

**LmxM.19.1440 (MPK4)**
N-terminus

Primary localisation: lysosome

Secondary localisation: cytoplasm, flagellum

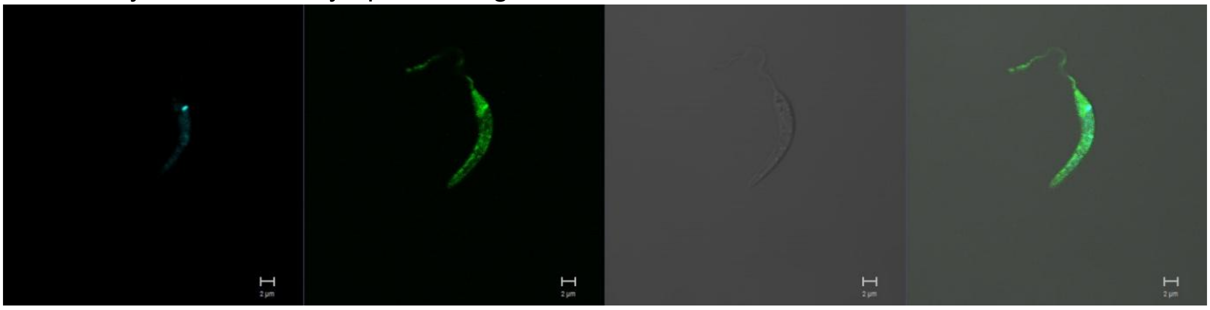

Field of view of MPK4 fluorescent mutants

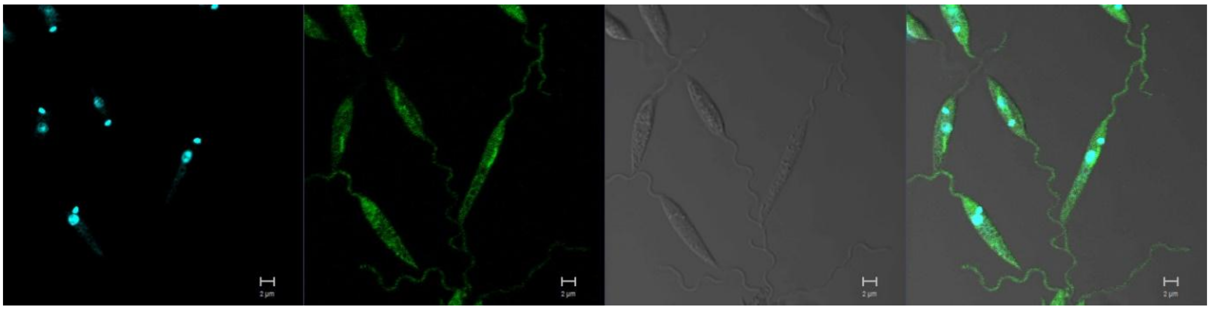

**LmxM.29.0370 (MPK12)**
N-terminus

Primary localisation: cytoplasm

Secondary localisation: flagellum, nucleus, kinetoplast

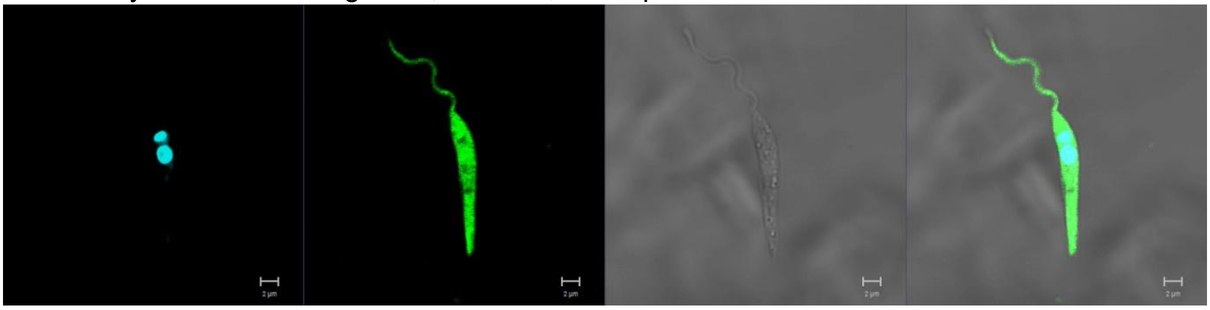

**LmxM.29.2910 (MPK5)**
N-terminus

Primary localisation: pellicular membrane

Secondary localisation: cytoplasm

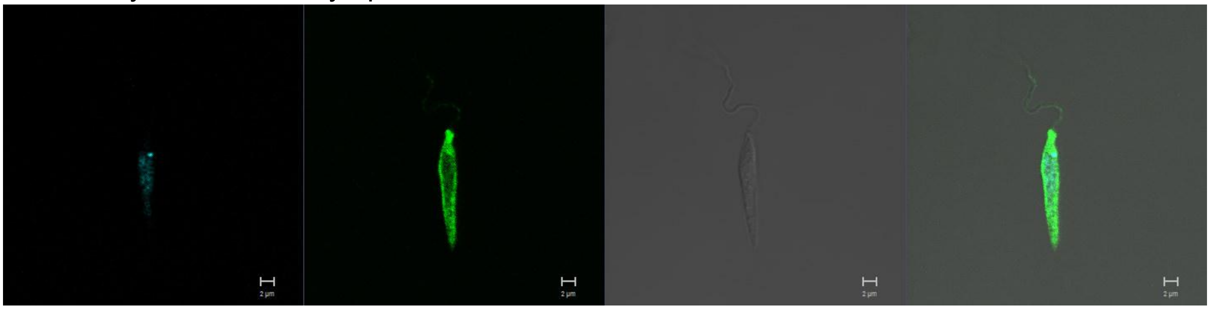

**LmxM.32.1380 (MPK11)**
N-terminus

Primary localisation: cytoplasm

Secondary localisation: nucleus, flagellum, basal body

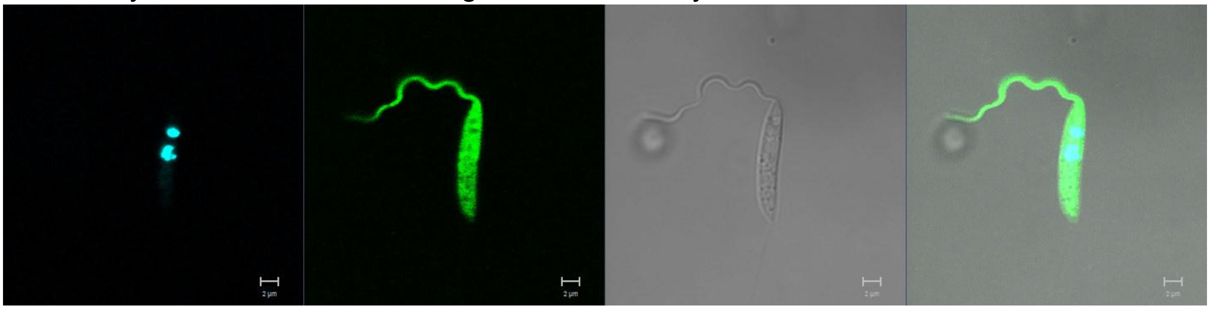

**LmxM.36.0720 (MPK2)**
N-terminus

Primary localisation: cytoplasm

Secondary localisation: flagellum, basal body

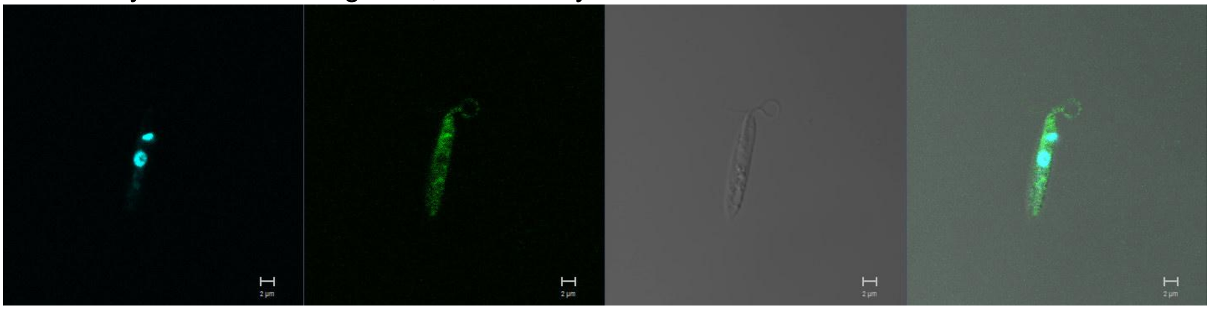

**LmxM.20_36.6470 (MPK1)**
N-terminus

Primary localisation: flagellum (anterior)

Secondary localisation: cytoplasm

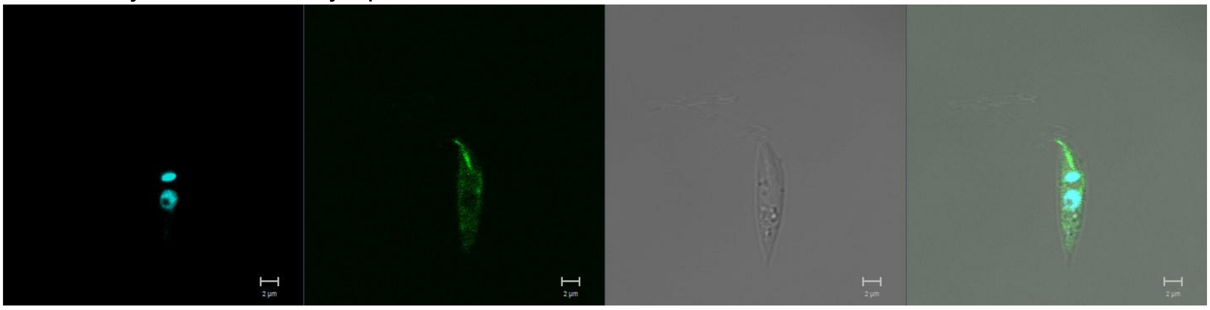

**LmxM.28.0580 (MPK8)**
N-terminus

Primary localisation: flagellum

Secondary localisation: cytoplasm

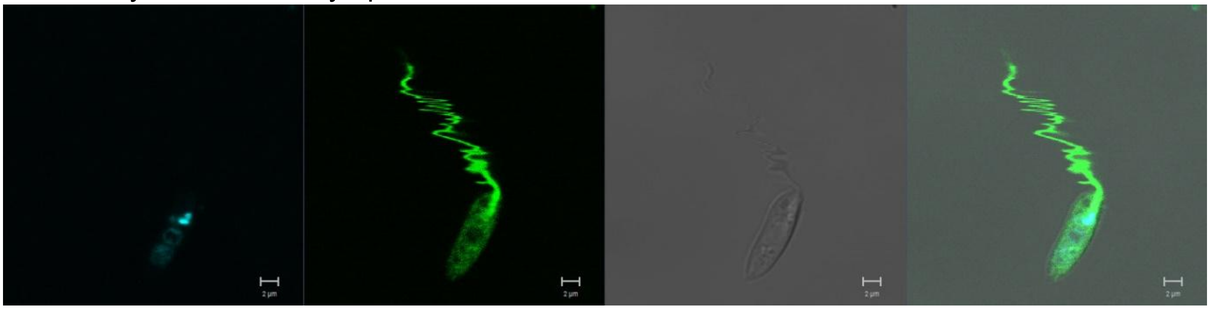

**LmxM.31.3250 (MPK6)**
N-terminus

Primary localisation in G1 phase: flagellum

Secondary localisation in G1 phase: cytoplasm

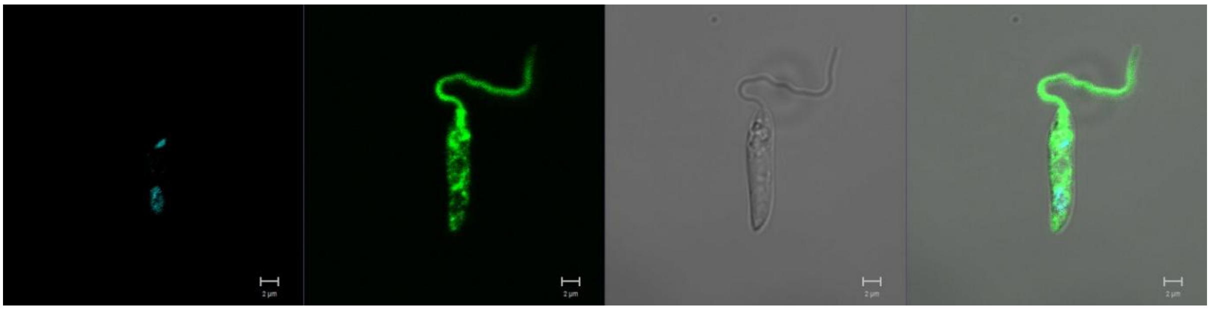

N-terminus

Primary localisation in S phase: pellicular membrane

Secondary localisation in S phase: flagellum, cytoplasm

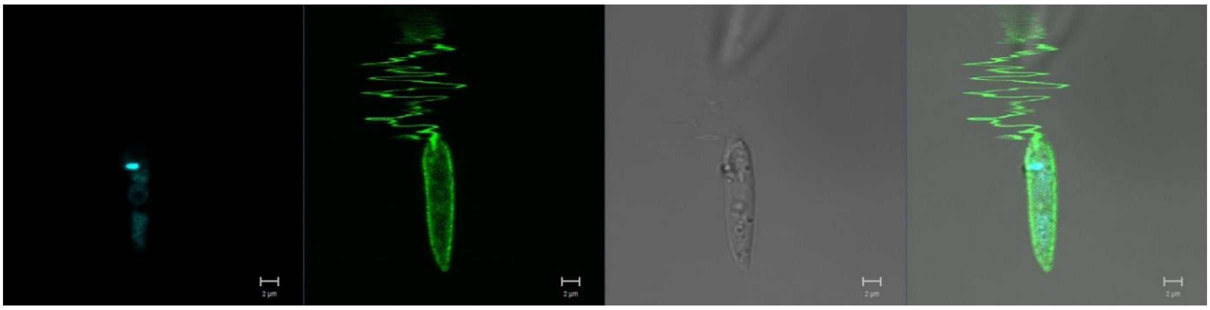

**LmxM.19.0180 (MPK9)**
N-terminus

Primary localisation: cytoplasm

Secondary localisation: lysosome

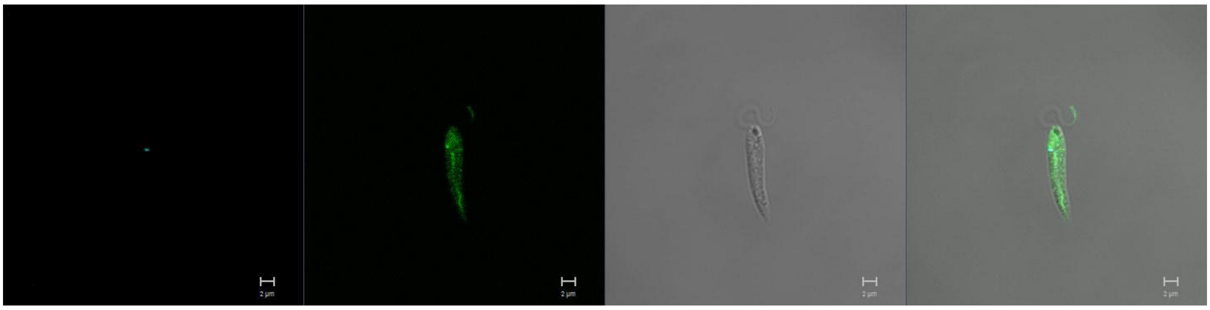

**LmxM.27.0100 (MPK14)**
N-terminus

Primary localisation: flagellum

Secondary localisation: basal body, cytoplasm

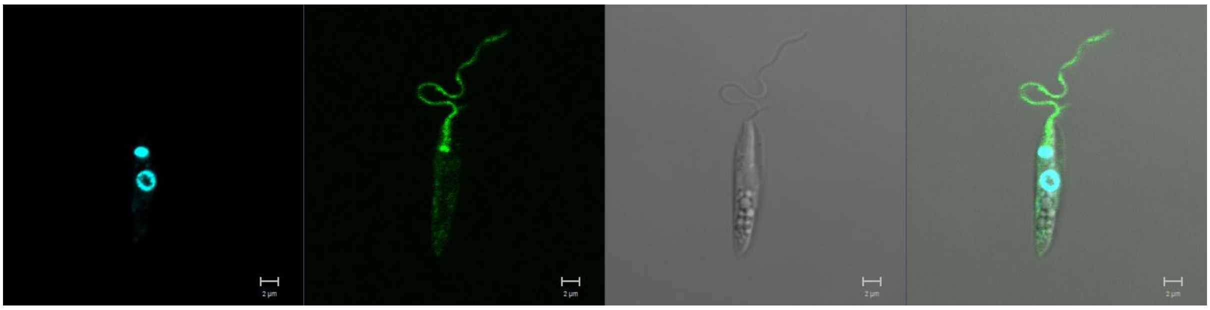

**LmxM.34.5010 (MPK13)**
N-terminus

Primary localisation: basal body

Secondary localisation: cytoplasm

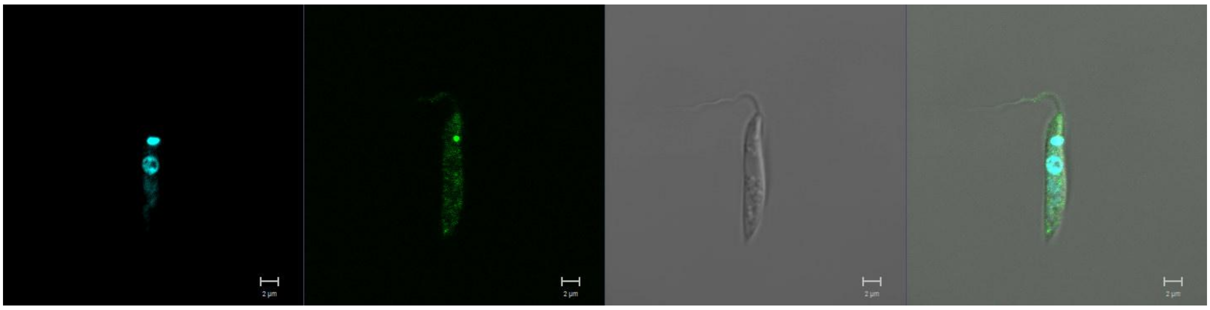

**LmxM.26.0980**
N-terminus

Primary localisation: cytoplasm

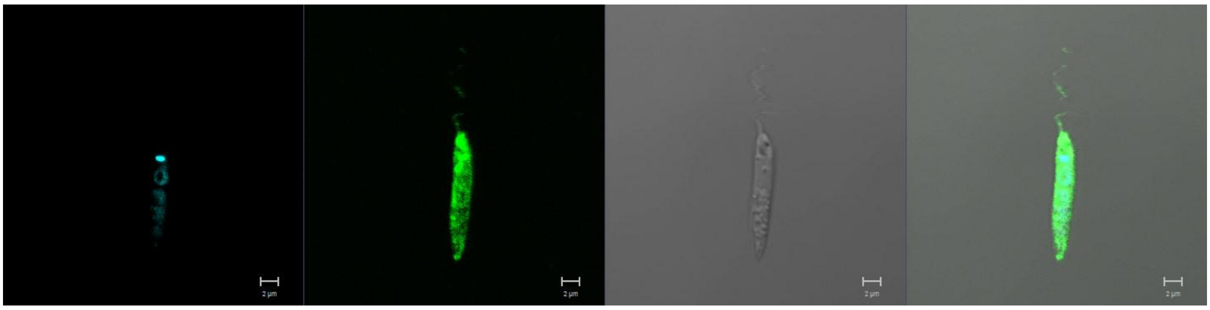

**LmxM.29.3580**
N-terminus

Primary localisation: cytoplasm

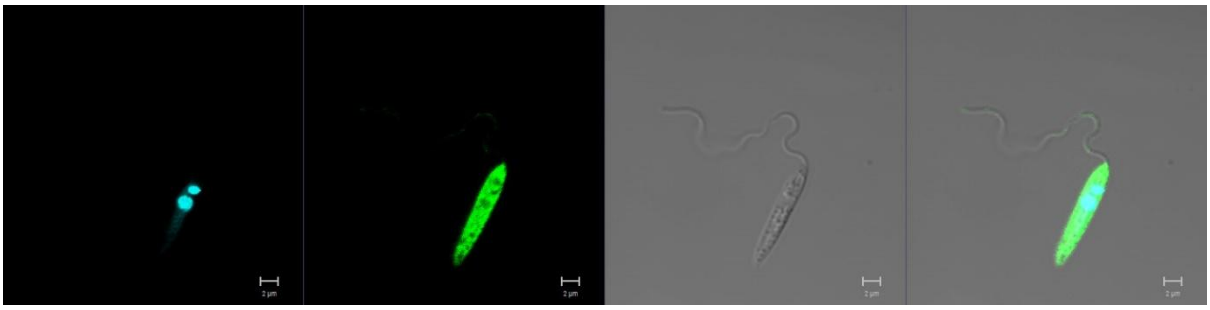

**LmxM.17.0670**
N-terminus

Primary localisation: basal body

Secondary localisation:  cytoplasm

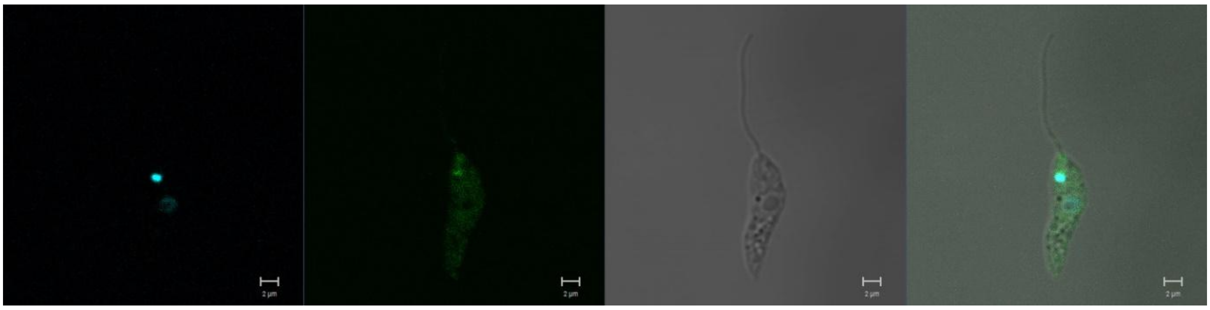

**LmxM.01.0750**
N-terminus

Primary localisation: cytoplasm

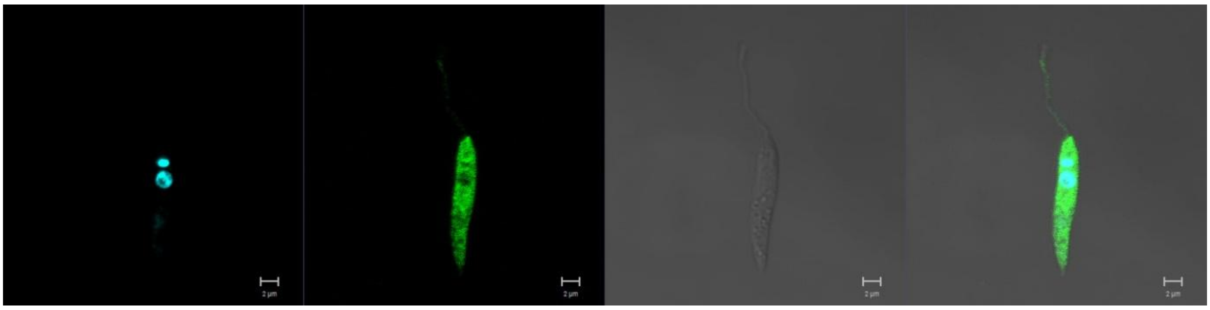

**LmxM.12.0130 (LUK2)**
N-terminus

Primary localisation: cytoplasm

Secondary localisation: flagellum, nucleus, kinetoplast

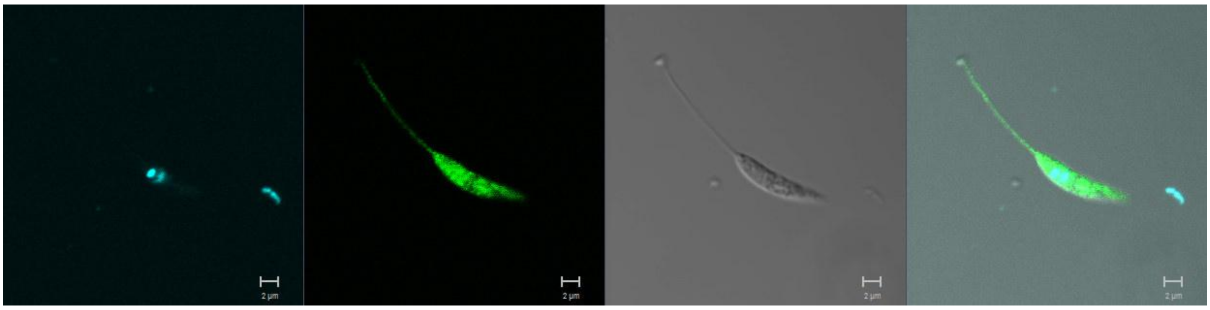

**STE FAMILY**

**LmxM.24.2320 (MKK4)**
N-terminus

Primary localisation: endomembrane

Secondary localisation: cytoplasm, flagellum

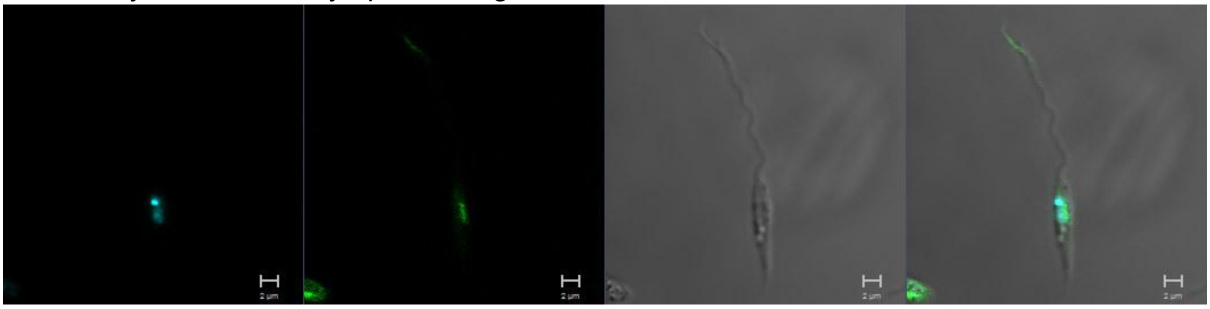

**LmxM.08_29.2320 (MKK1)**
C-terminus

Primary localisation: cytoplasm

Secondary localisation: flagellum

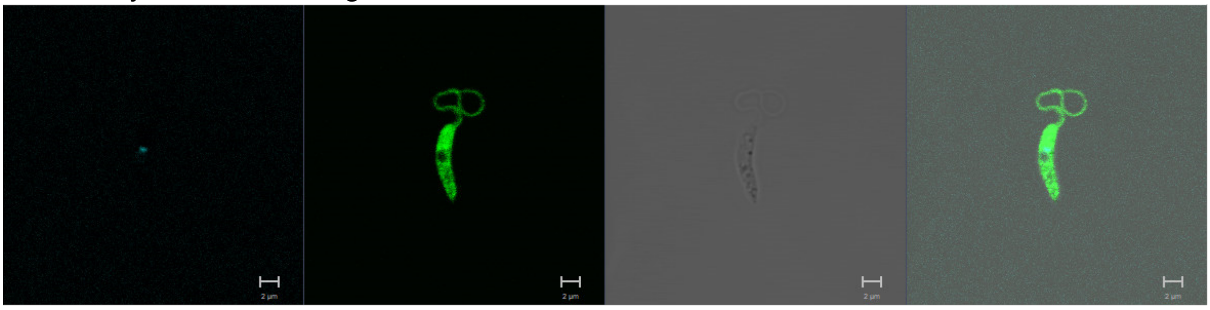

**LmxM.08.1228**

N-terminus

Primary localisation: cytoplasm

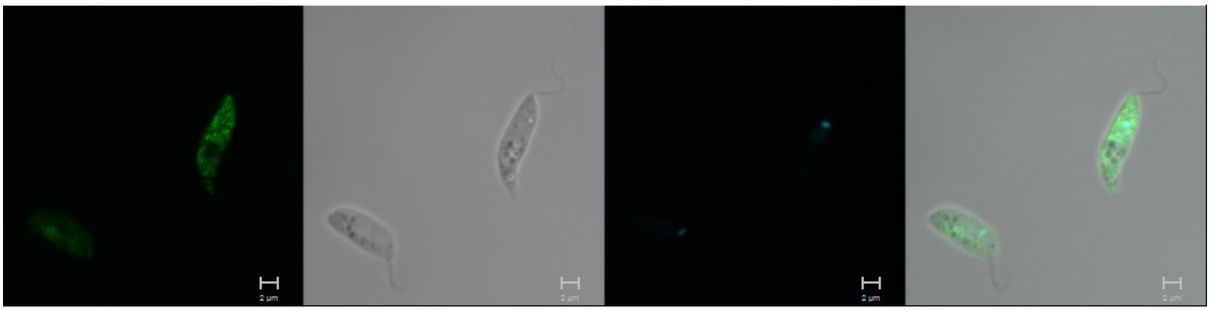

**LmxM.15.1200**
N-terminus

Primary localisation: cytoplasm

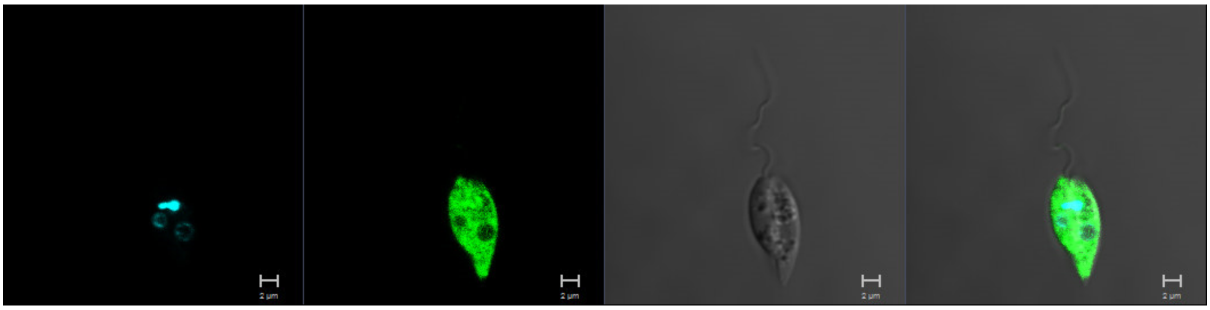

**LmxM.17.0390**
N-terminus

Primary localisation: basal body

Secondary localisation: cytoplasm

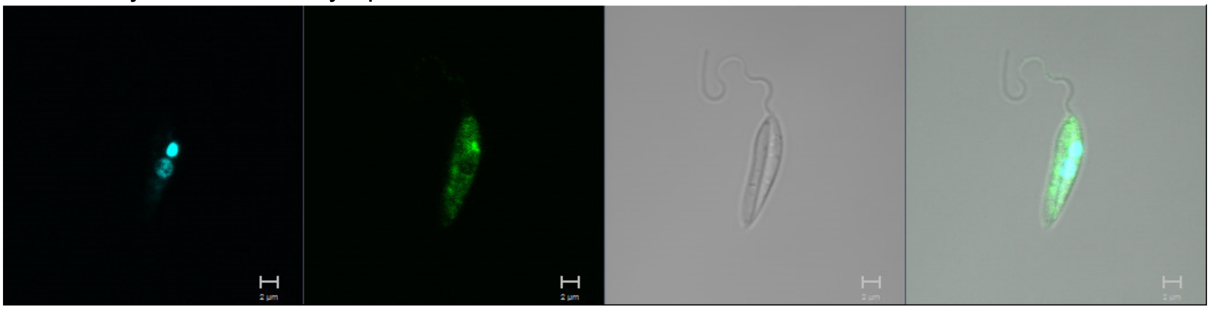

**LmxM.20.0770**
N-terminus

Primary localisation: pellicular membrane

Secondary localisation: cytoplasm

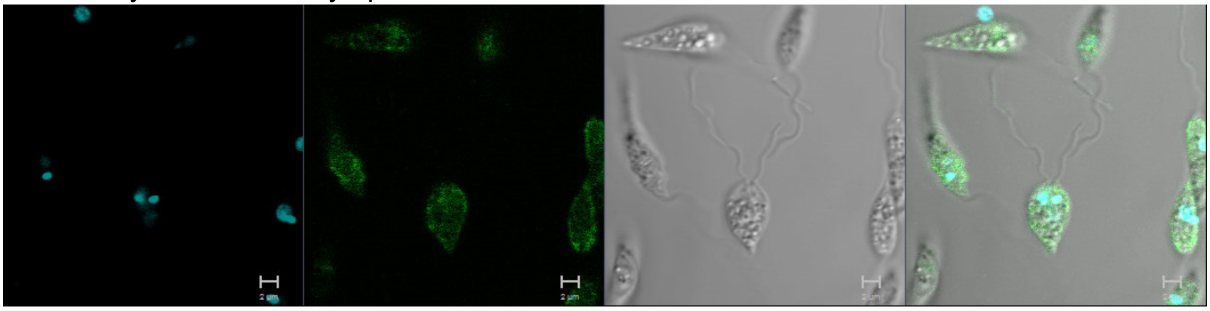

**LmxM.21.0270**
N-terminus

Primary localisation: cytoplasm

Secondary localisation: basal body, flagellum

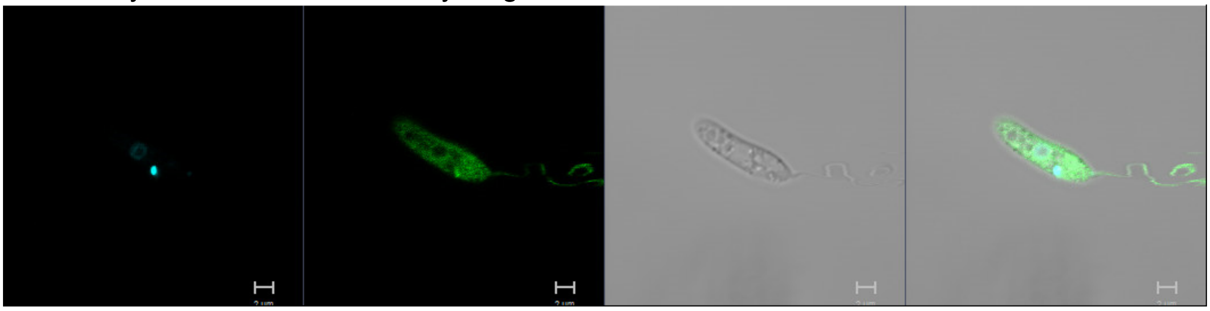

**LmxM.25.1990**
N-terminus

Primary localisation: endomembrane

Secondary localisation: lysosome, cytoplasm

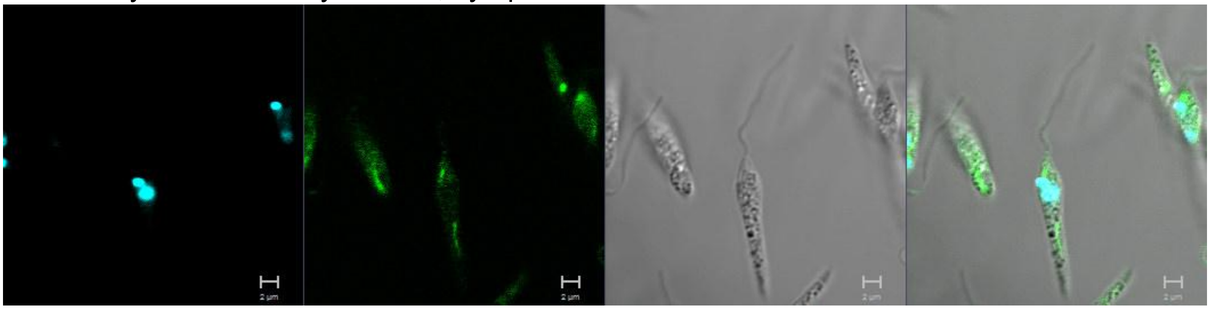

**LmxM.31.1020**
C-terminus

Primary localisation: cytoplasm

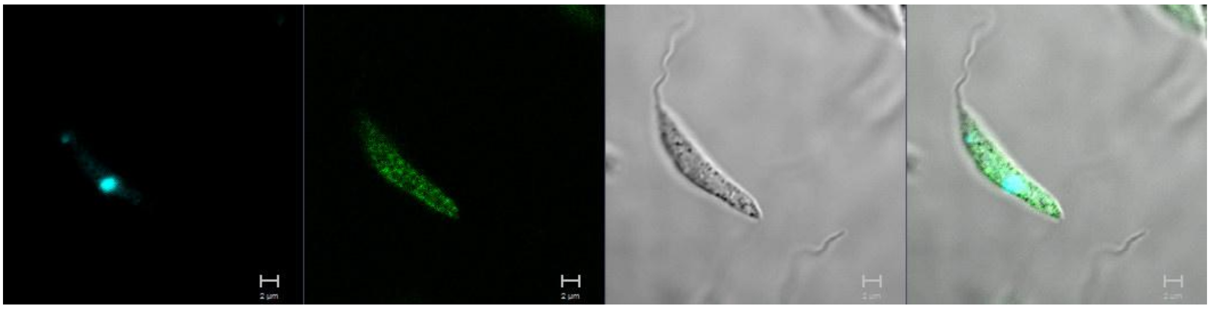

**LmxM.34.3170**
N-terminus

Primary localisation: cytoplasm

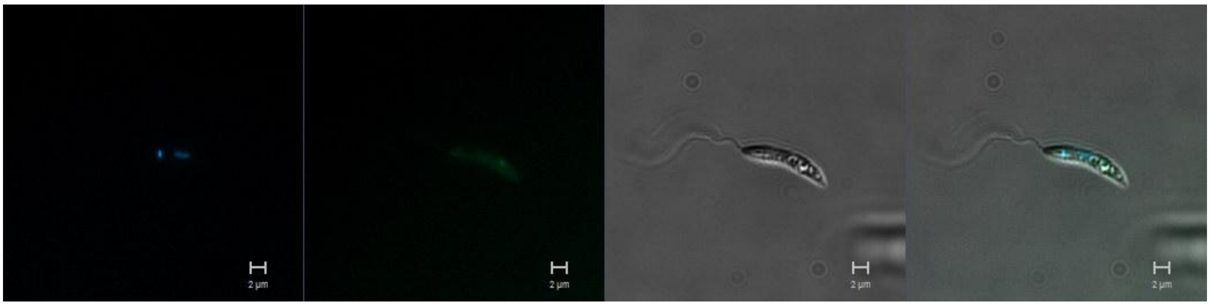

**LmxM.04.0440**
N-terminus

Primary localisation: flagellar pocket (collar region)

Secondary localisation: lysosomal, pellicular membrane, cytoplasm

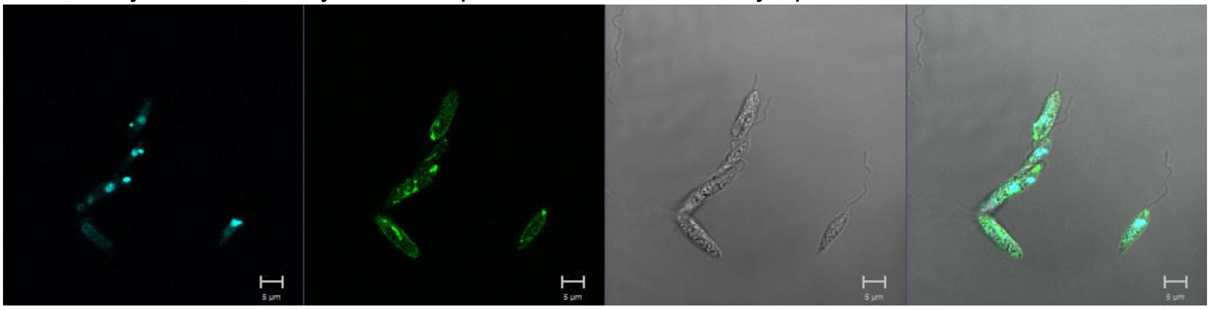

**LmxM.05.0390**
N-terminus

Primary localisation: basal body

Secondary localisation: flagellar pocket, cytoplasm, flagellum

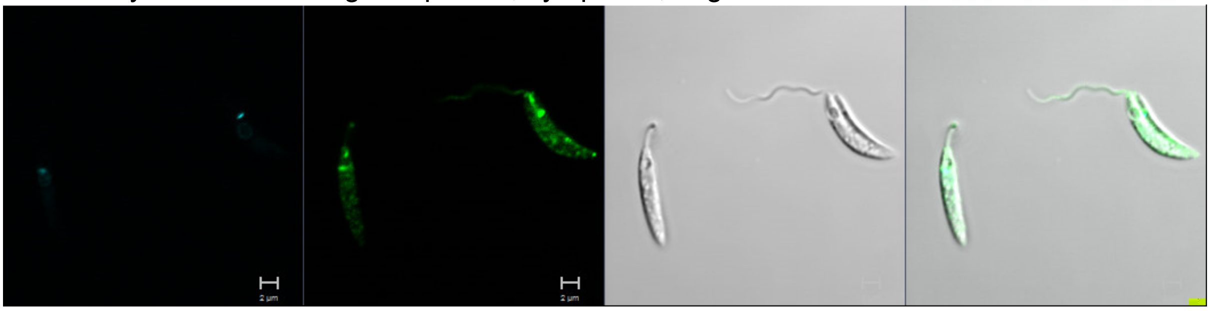

C-terminus

Primary localisation: basal body

Secondary localisation: flagellar pocket, cytoplasm

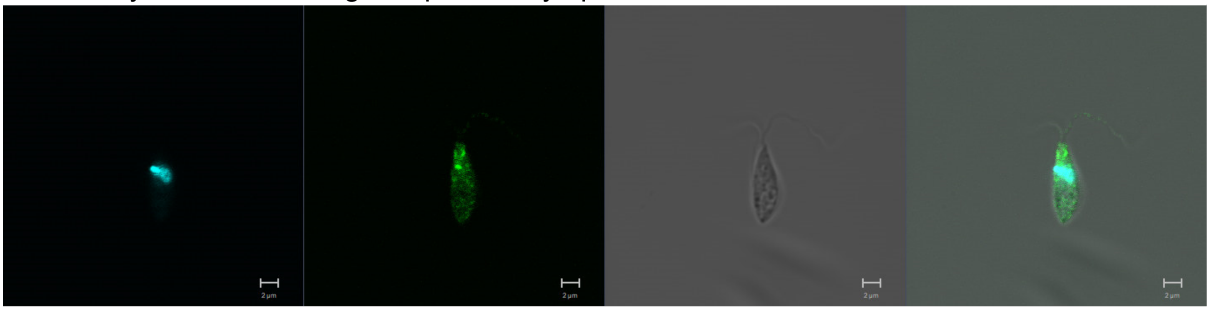

**LmxM.06.0640**
N-terminus

Primary localisation: lysosome

Secondary localisation: endomembrane

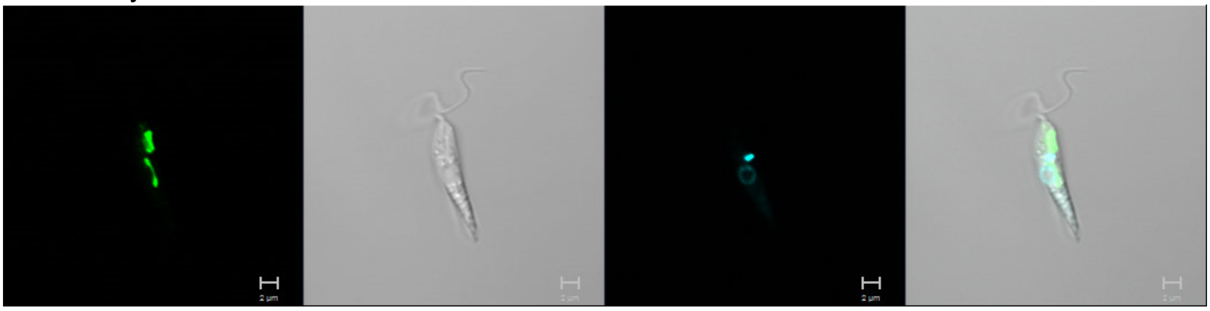

**LmxM.07.0690**
N-terminus

Primary localisation: cytoplasm

Secondary localisation: nucleus

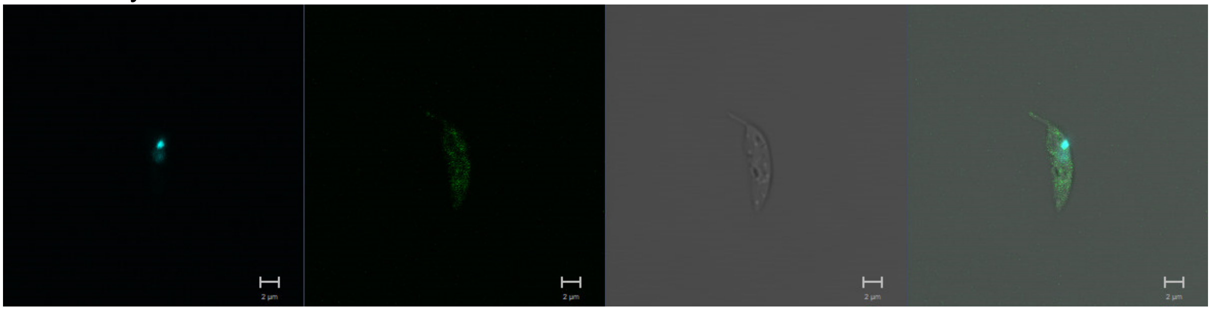

**LmxM.07.0880**
N-terminus

Primary localisation: cytoplasm

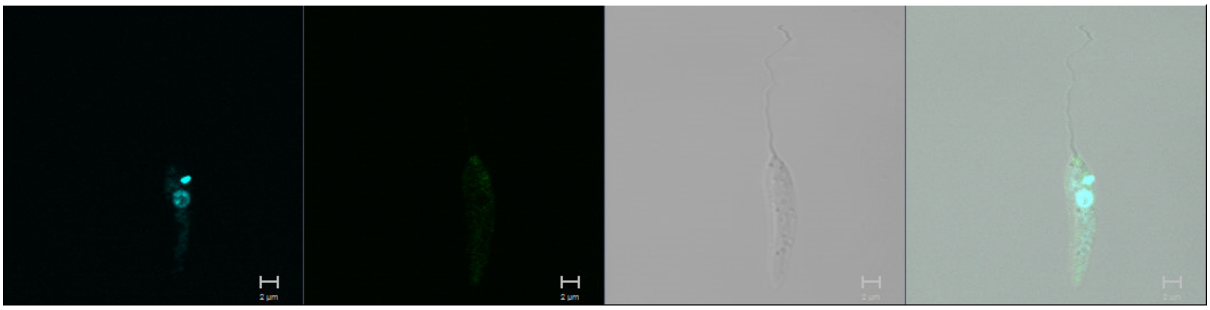

**LmxM.14.1300**
N-terminus

Primary localisation: lysosome

Secondary localisation: endomembrane

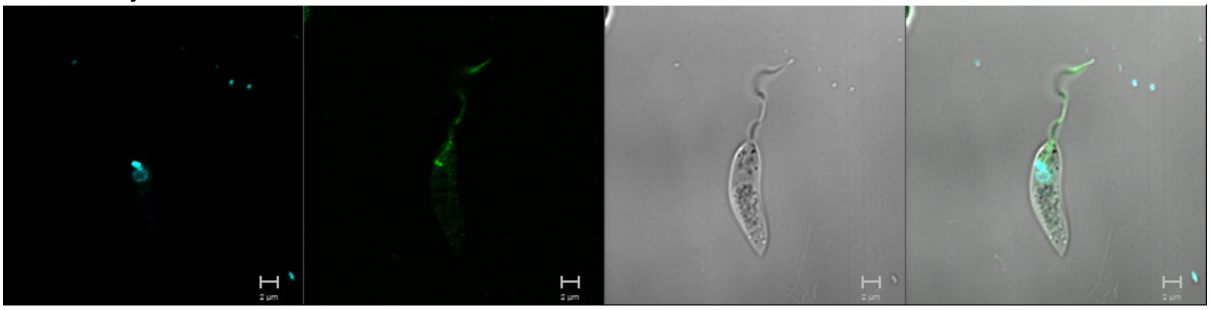

**LmxM.17.0490**
N-terminus

Primary localisation: flagellum

Secondary localisation: cytoplasm

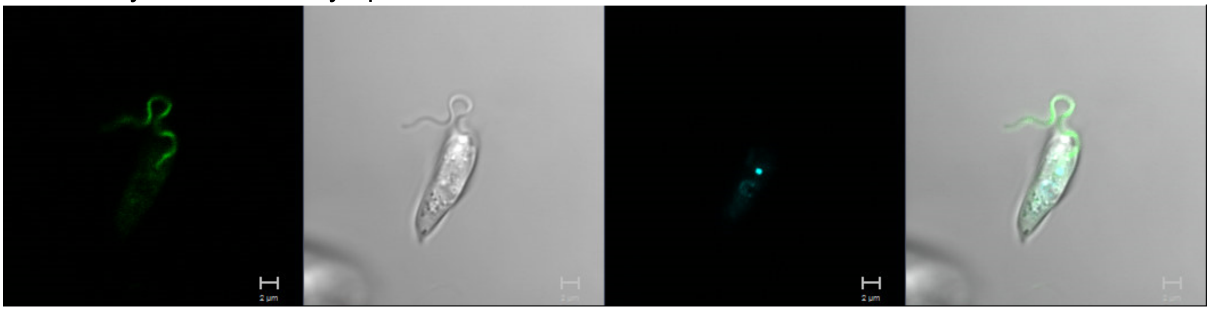

**LmxM.19.0150 (CBPK1)**
N-terminus

Primary localisation: cytoplasm

Secondary localisation: flagellum

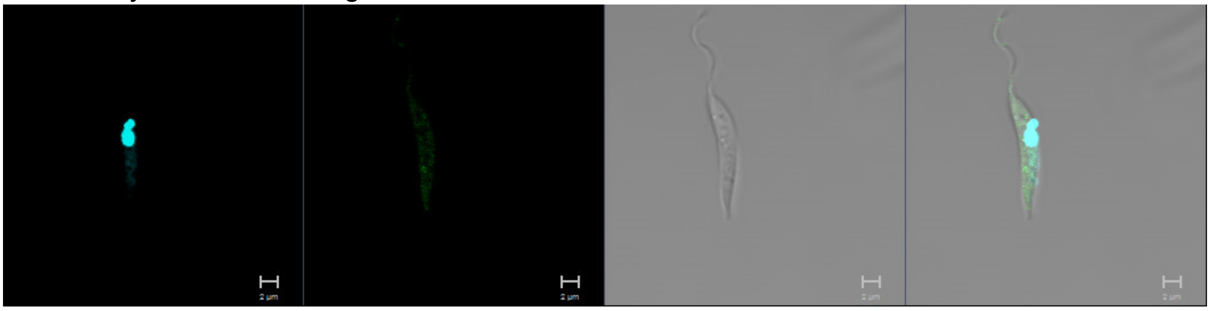

**LmxM.21.0130**
N-terminus

Primary localisation: endomembrane

Secondary localisation: golgi apparatus, cytoplasm

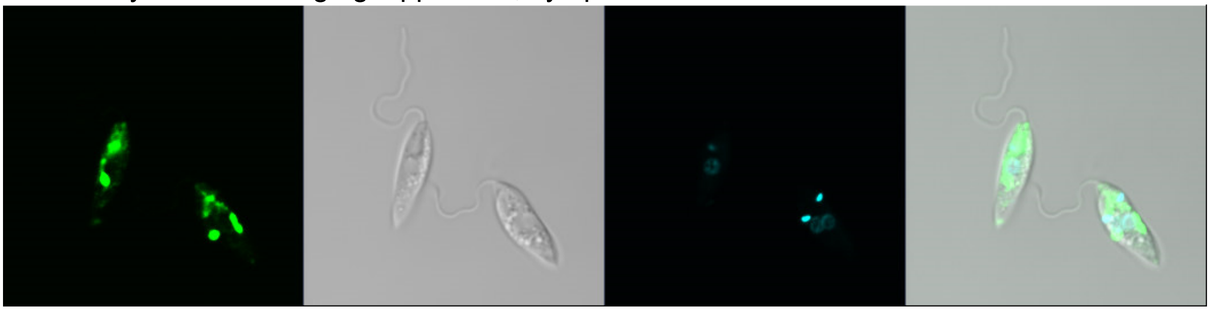

**LmxM.24.1450**
N-terminus

Primary localisation: cytoplasm

Secondary localisation: flagellum, nucleus, kinetoplast

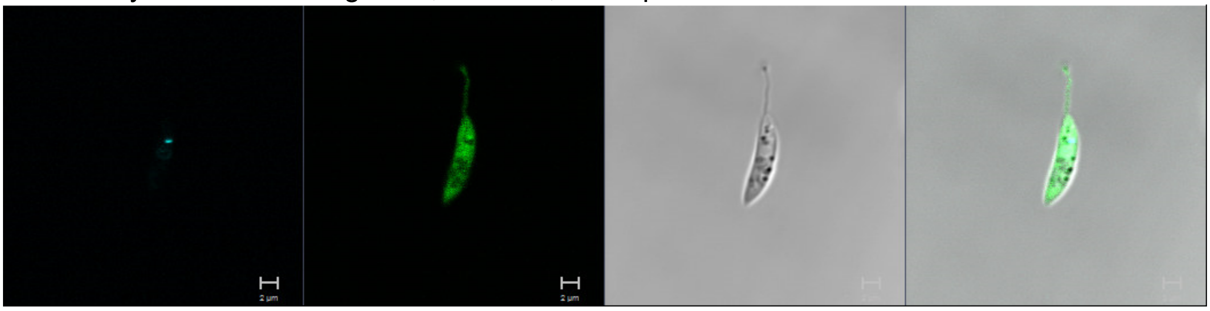

**LmxM.26.1730**
N-terminus

Primary localisation: lysosome

Secondary localisation: flagellum

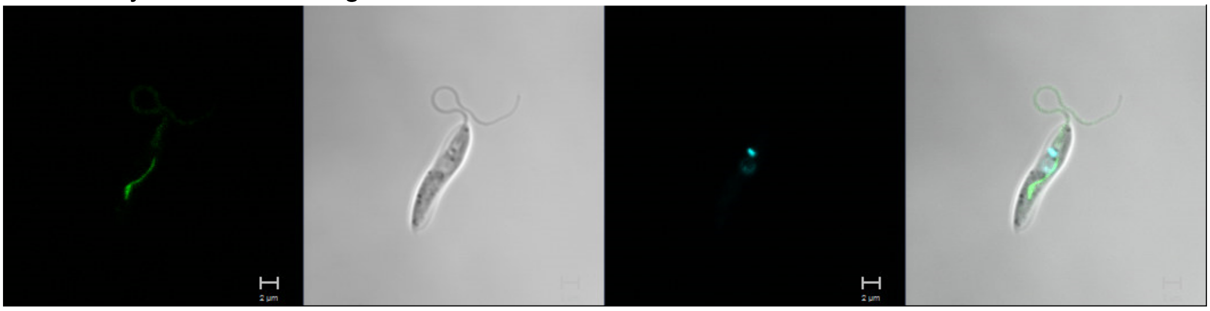

**LmxM.27.1370**
N-terminus

Primary localisation: endomembrane

Secondary localisation: cytoplasm

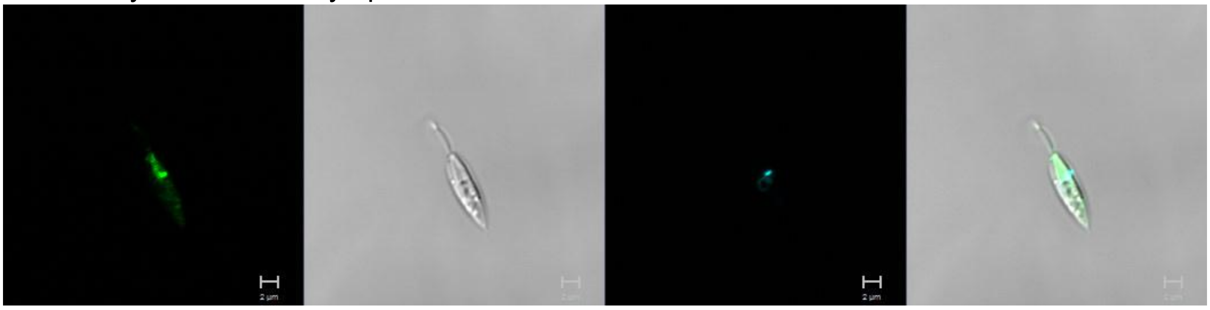

**LmxM.29.0600**
N-terminus

Primary localisation: flagellum

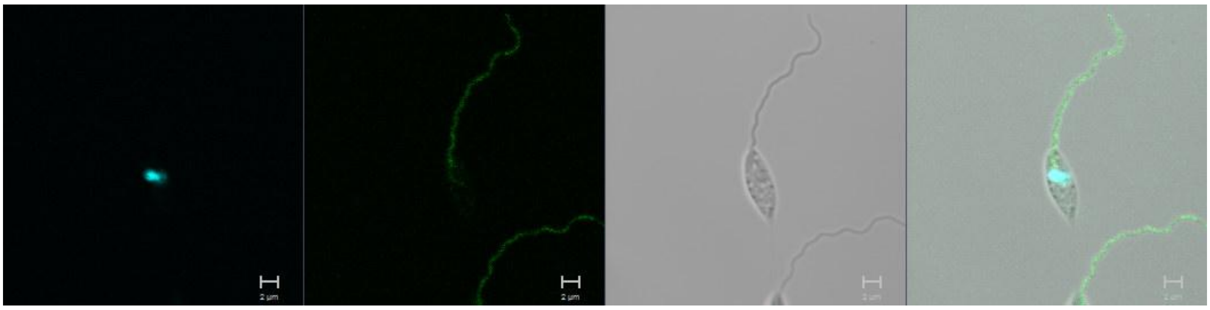

**LmxM.29.3050**
N-terminus

Primary localisation: cytoplasm (low expression)

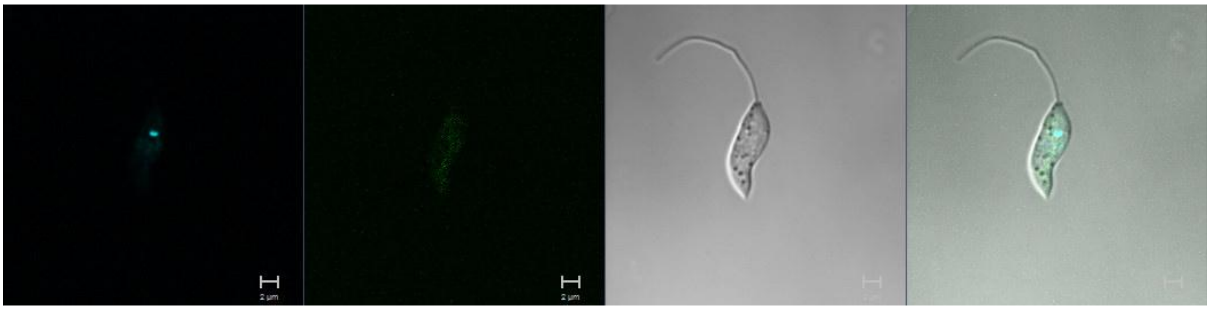

**LmxM.31.0120 (MRK1)**
N-terminus

Primary localisation: cytoplasm

Secondary localisation: kinetoplast

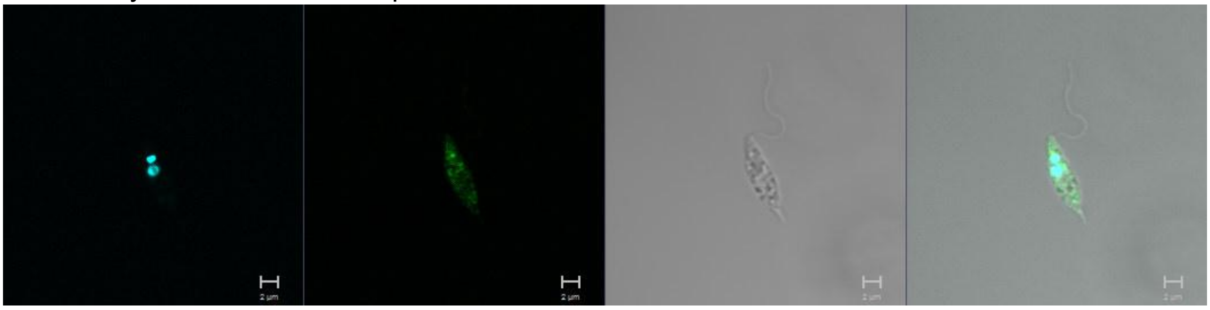

**LmxM.31.0780**
N-terminus

Primary localisation: basal body

Secondary localisation: cytoplasm

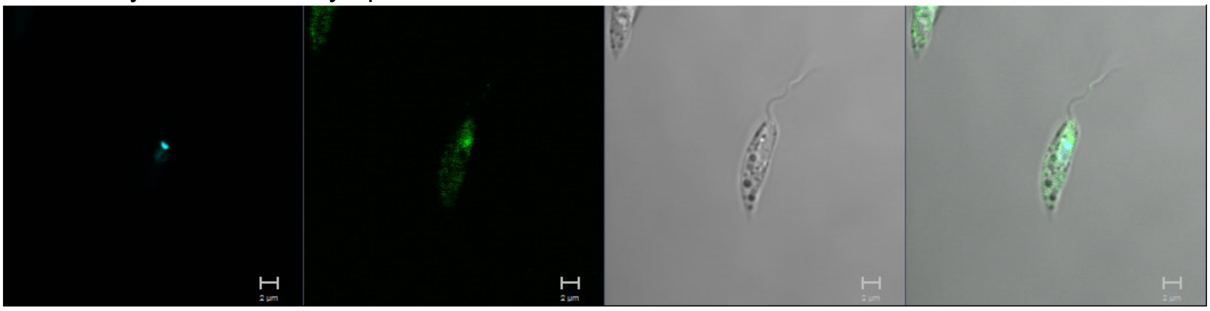

**LmxM.31.0810 (RDK1)**
N-terminus

Primary localisation: cytoplasm

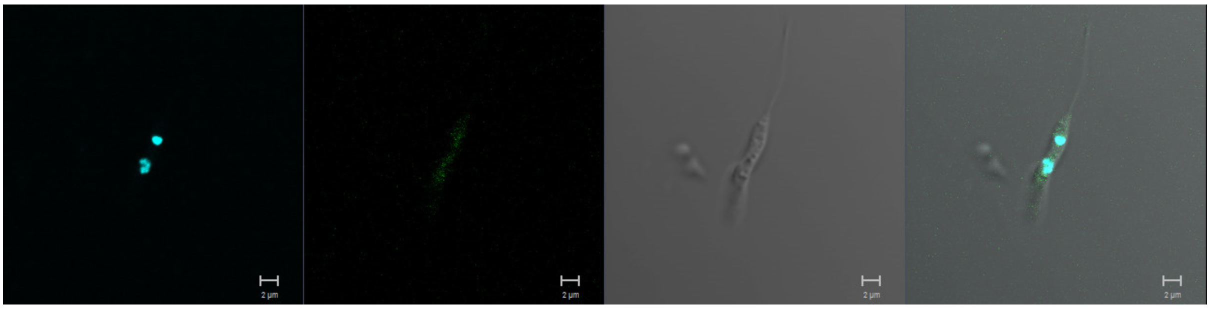

**LmxM.32.1400**
C-terminus

Primary localisation: nucleus

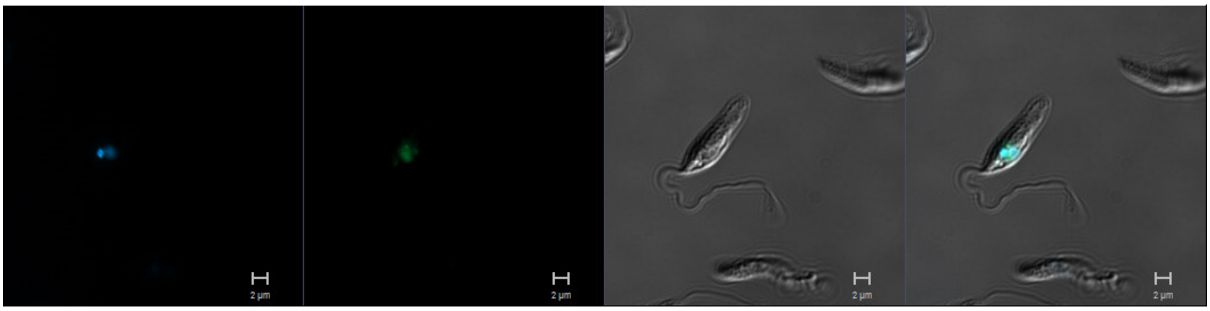

**LmxM.32.2290**
N-terminus

Primary localisation: lysosome

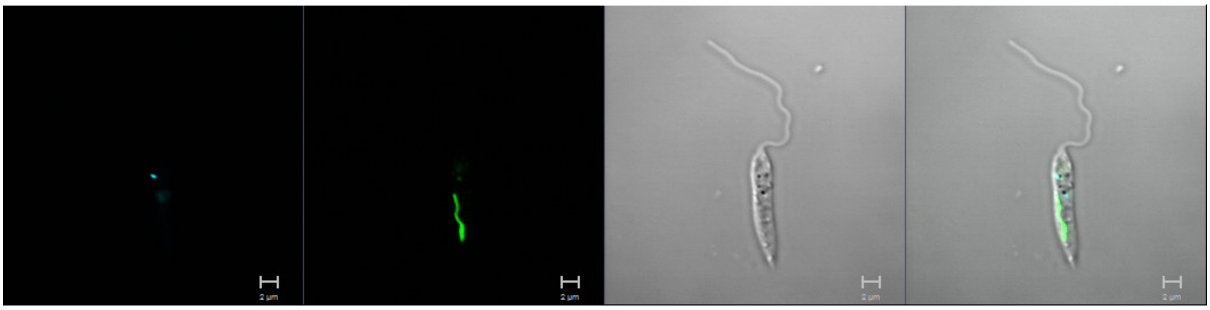

**LmxM.36.0910**
N-terminus

Primary localisation in log-phase culture: endomembrane

Secondary localisation: flagellum

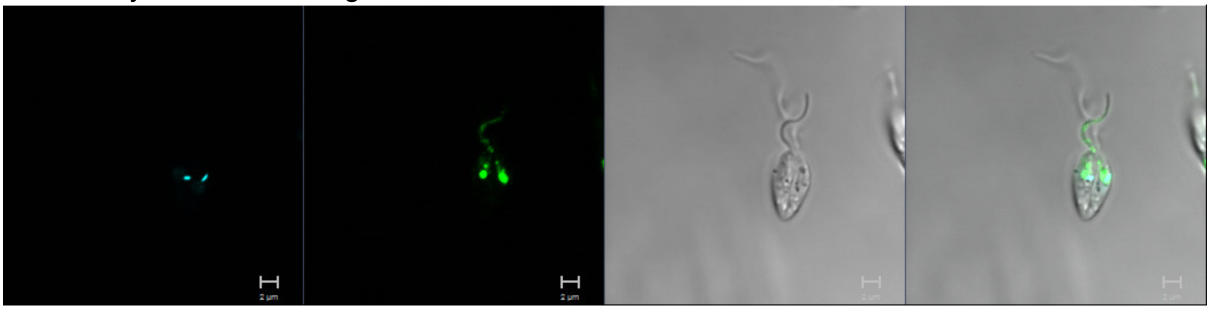

Primary localisation in stationary phase culture: lysosome

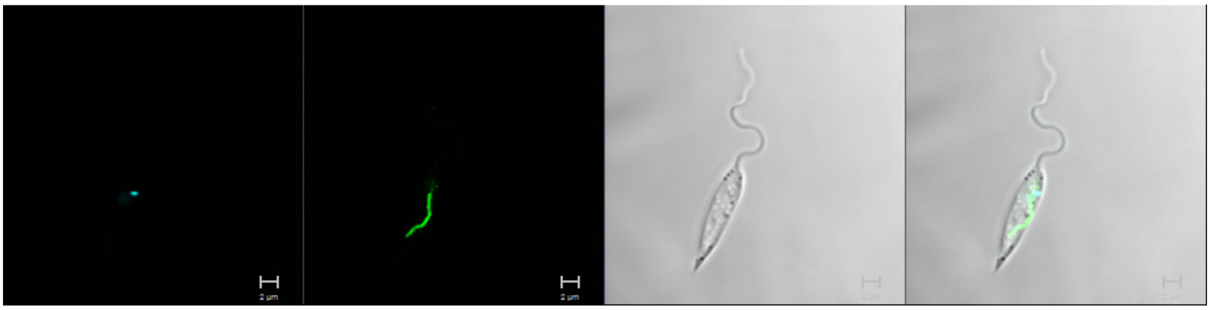

**LmxM.16.0300**
N-terminus

Primary localisation: basal body

Secondary localisation: cytoplasm

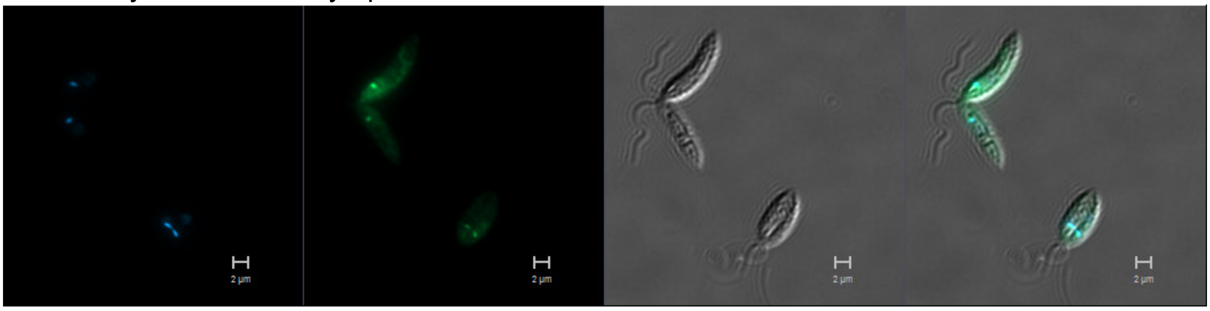


C-terminus

Primary localisation: basal body

Secondary localisation: cytoplasm

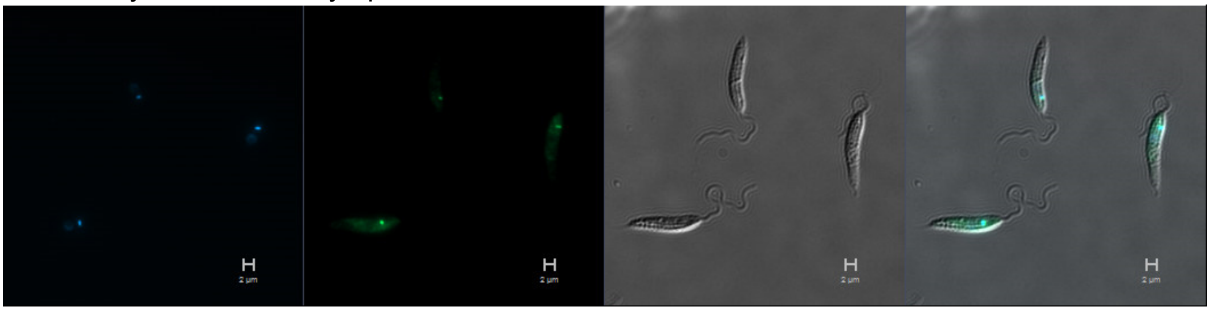


**LmxM.07.0250**
C-terminus

Primary localisation: cytoplasm

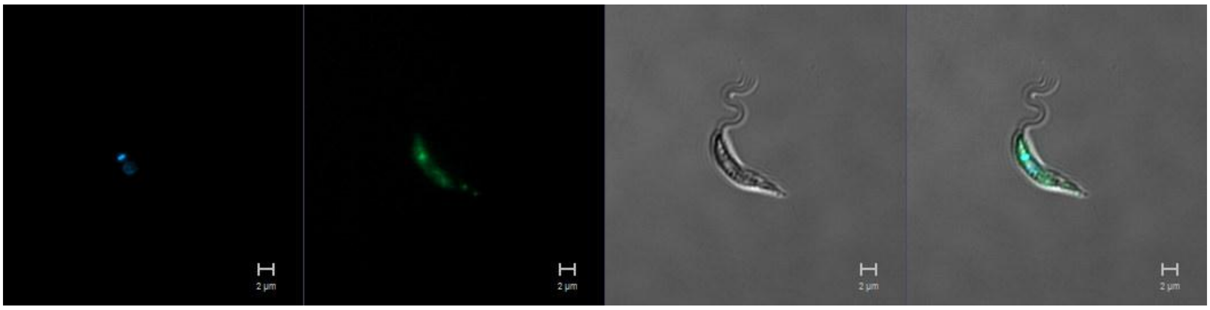

**LmxM.36.0860 (MKK5)**
C-terminus

Primary localisation: cytoplasm

Secondary localisation: flagellum

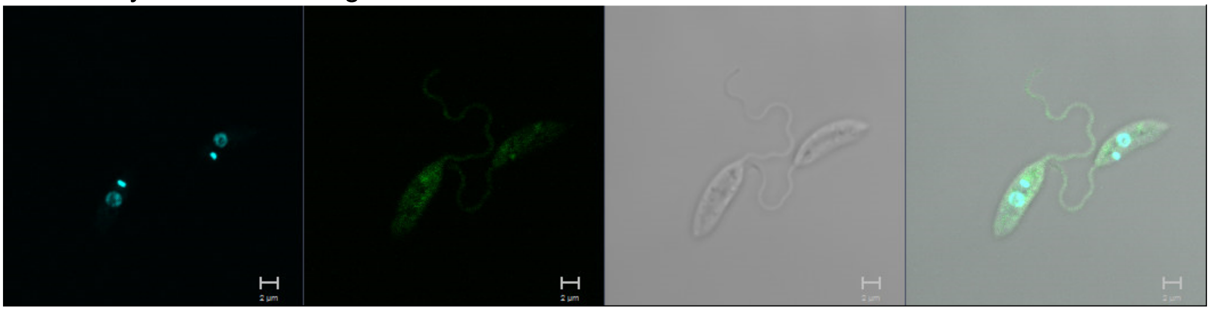

**LmxM.19.1610**
C-terminus

Primary localisation: cytoplasm

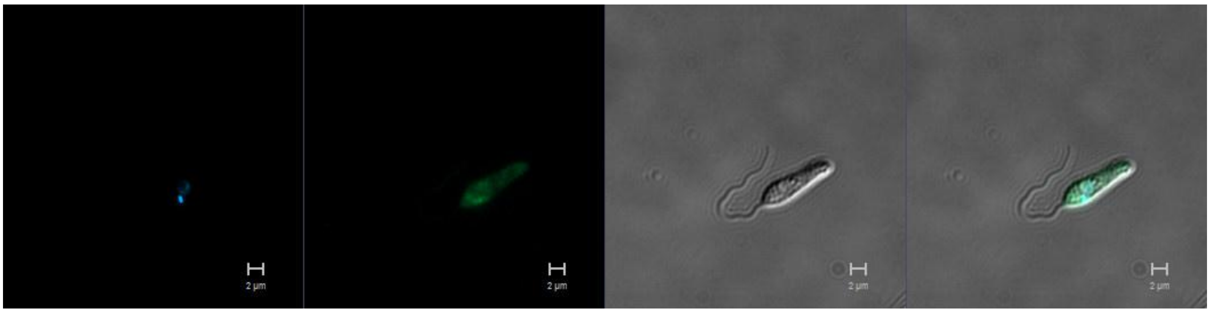

**LmxM.33.2090**
N-terminus

Primary localisation: cytoplasm

Secondary localisation: flagellum

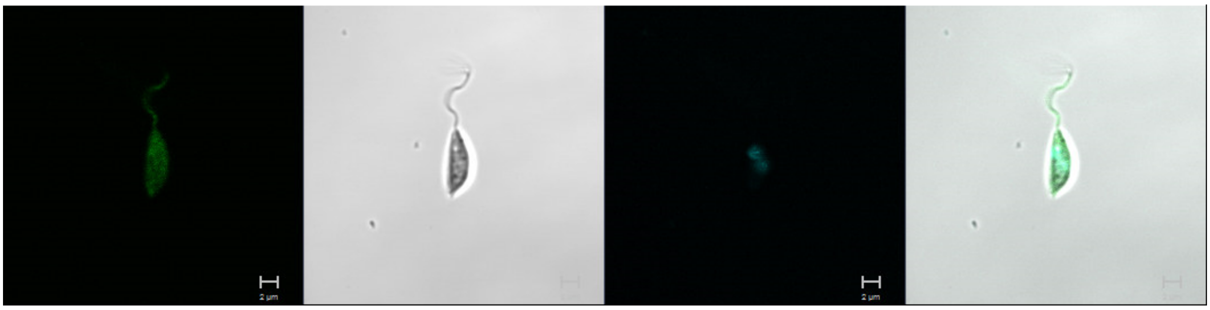

**LmxM.34.4000**
N-terminus

Primary localisation: cytoplasm

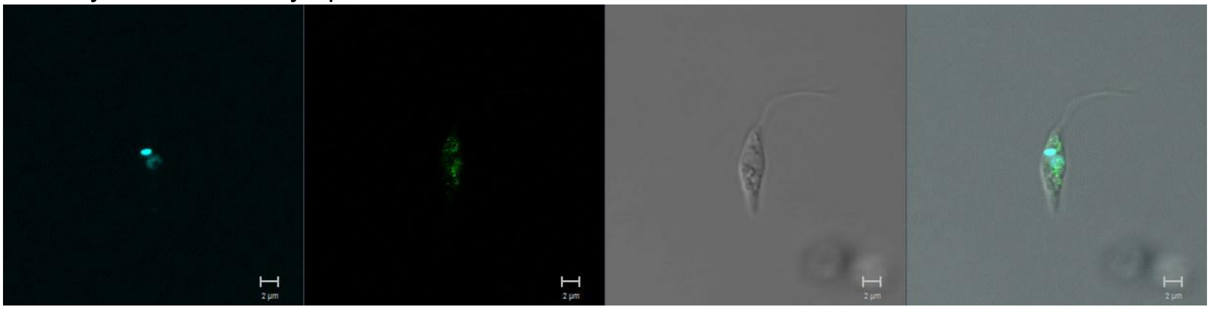

**LmxM.36.3680**
N-terminus

Primary localisation: endomembrane

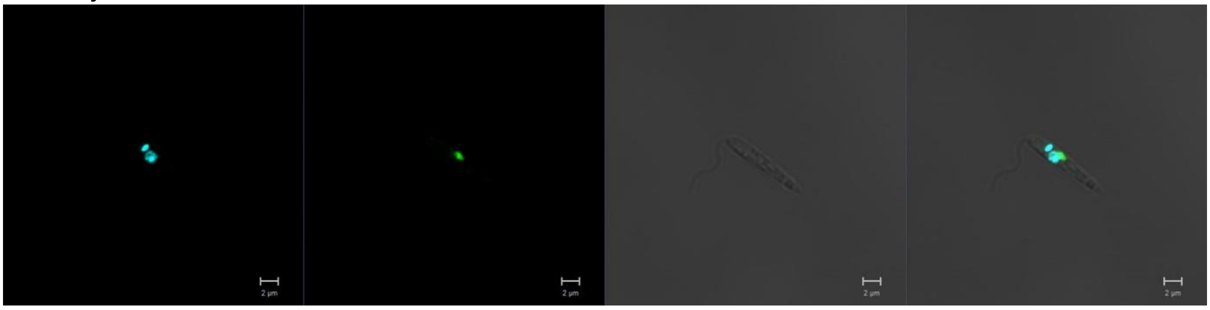

**LmxM.30.1830**
N-terminus

Primary localisation: endomembrane

Secondary localisation: cytoplasm

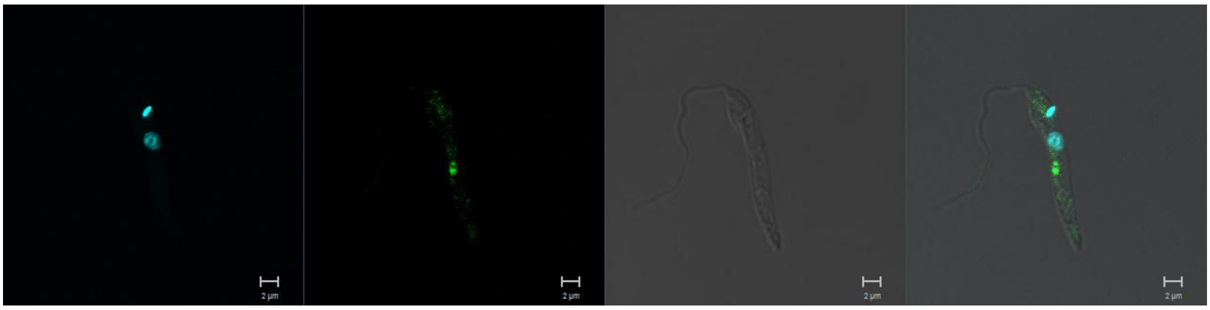

**LmxM.30.1840**
N-terminus

Primary localisation: cytoplasm

Secondary localisation: nucleus, flagellum, basal body

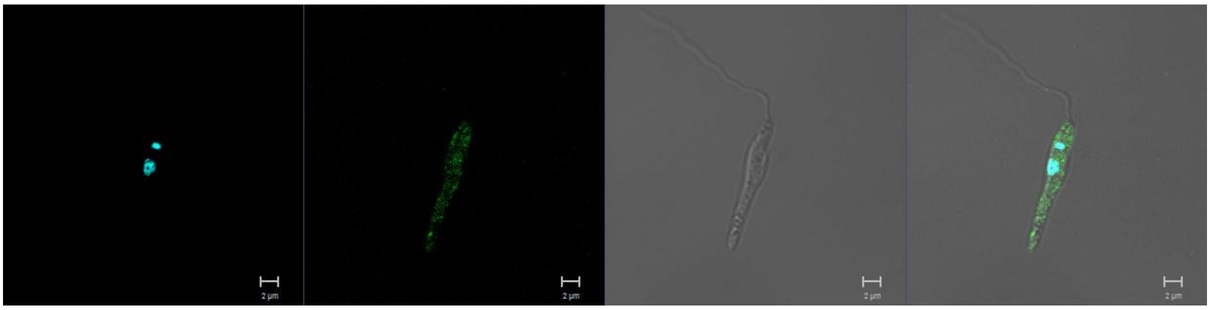

**NEK FAMILY**

**LmxM.02.0290**
N-terminus

Primary localisation: cytoplasm

Secondary localisation: nucleus, flagellum

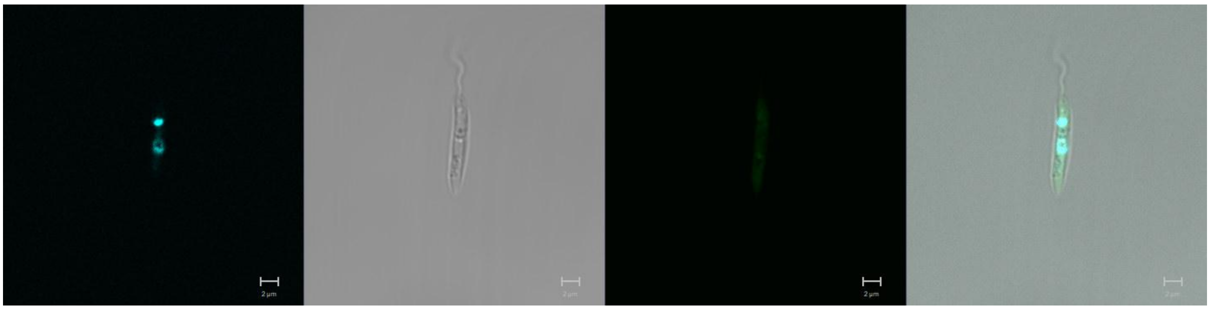

Primary localisation in dividing cells: basal body

Secondary localisation in dividing cells: cytoplasm, flagellum

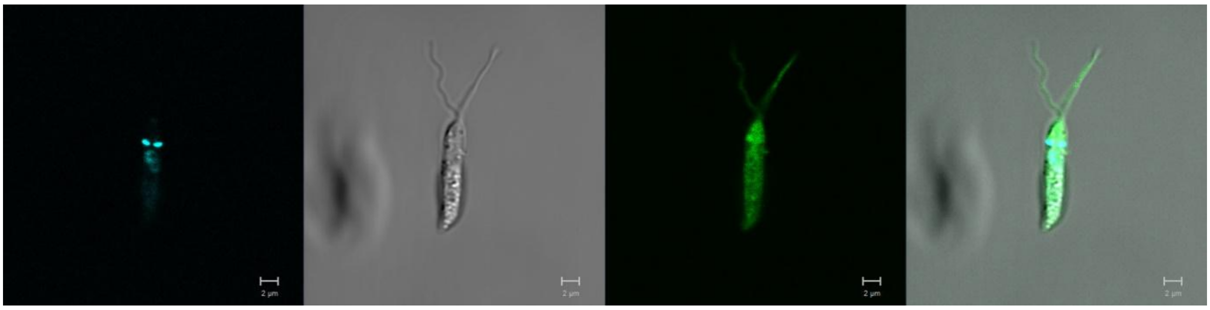

**LmxM.07.0160**
N-terminus

Primary localisation: endomembrane

Secondary localisation: lysosome (region)

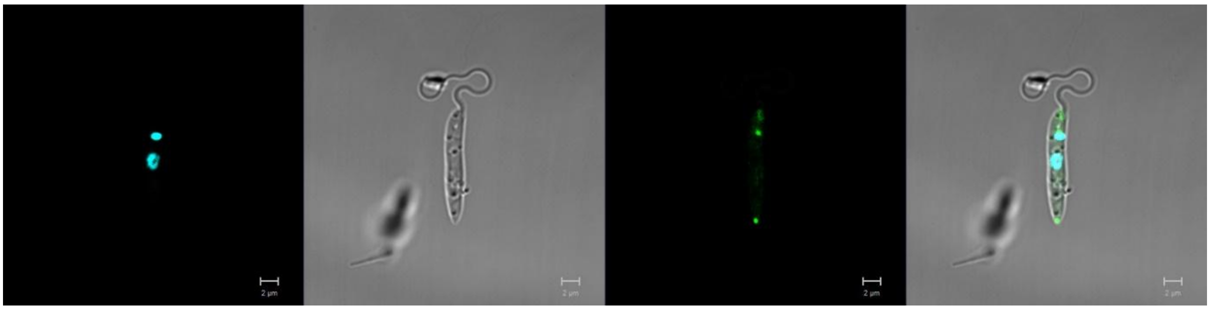

**LmxM.07.0170**
N-terminus

Primary localisation: basal body

Secondary localisation: cytoplasm, flagellum

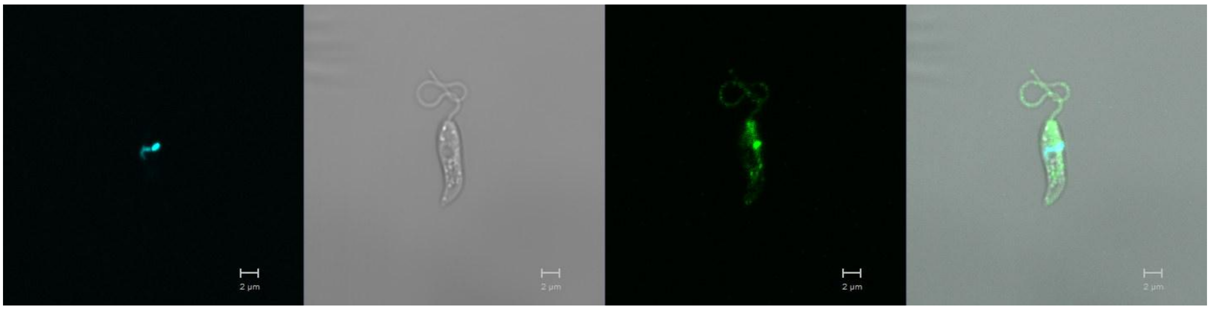

**LmxM.08.0930**
N-terminus

Primary localisation: flagellum

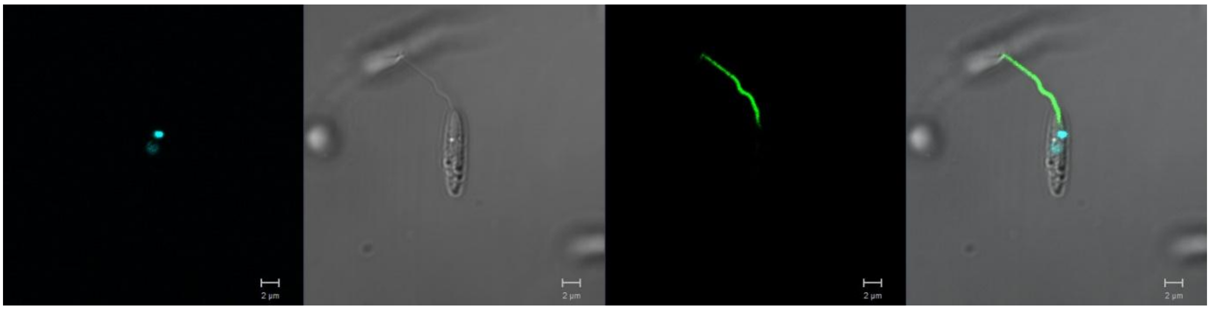

**LmxM.14.1410**
N-terminus

Primary localisation: flagellum

Secondary localisation: cytoplasm

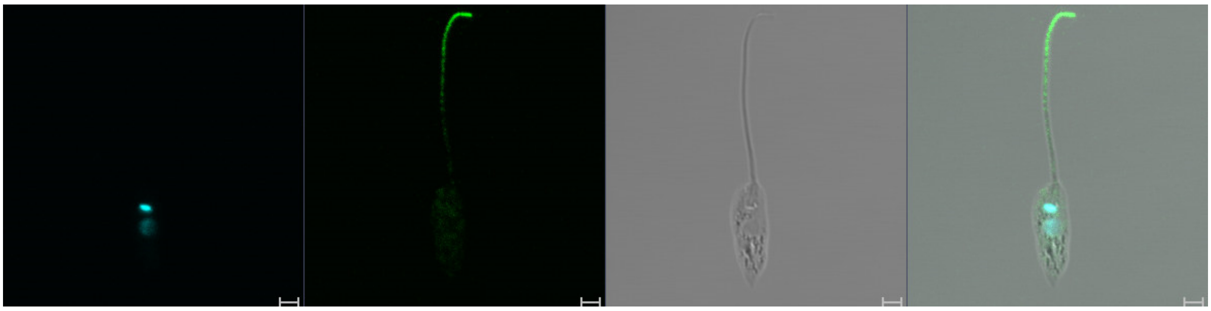

**LmxM.21.0853**
N-terminus

Primary localisation: cytoplasm

Secondary localisation: flagellum

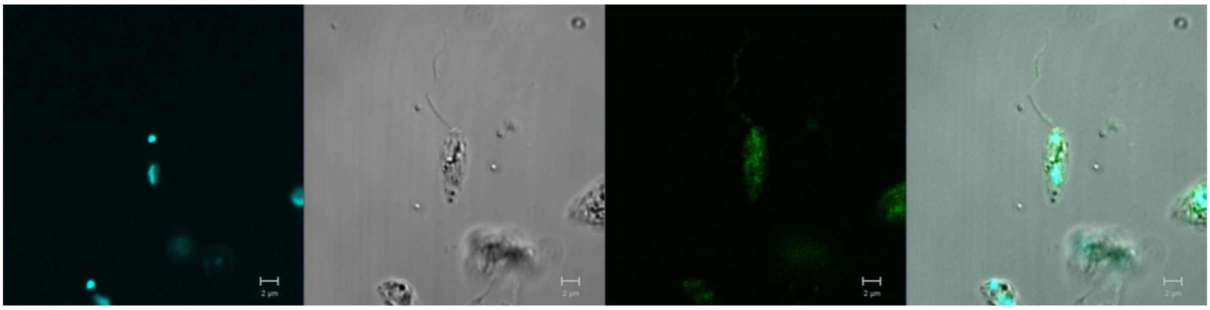

**LmxM.21.1565**
N-terminus

Primary localisation: basal body

Secondary localisation: cytoplasm, flagellum

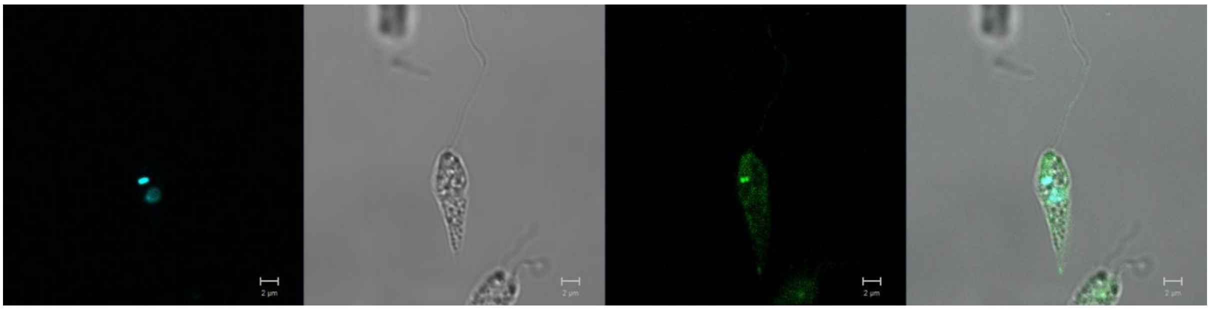

**LmxM.22.0950**
N-terminus

Primary localisation: cytoplasm

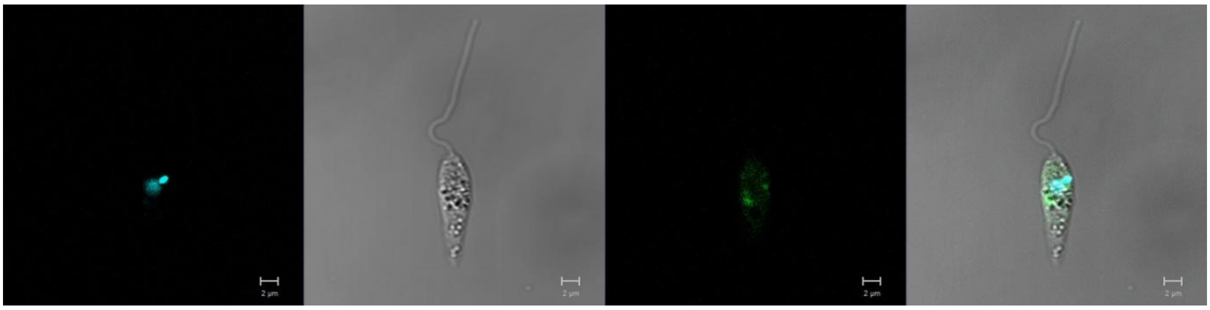

**LmxM.26.2570**
N-terminus

Primary localisation: basal body

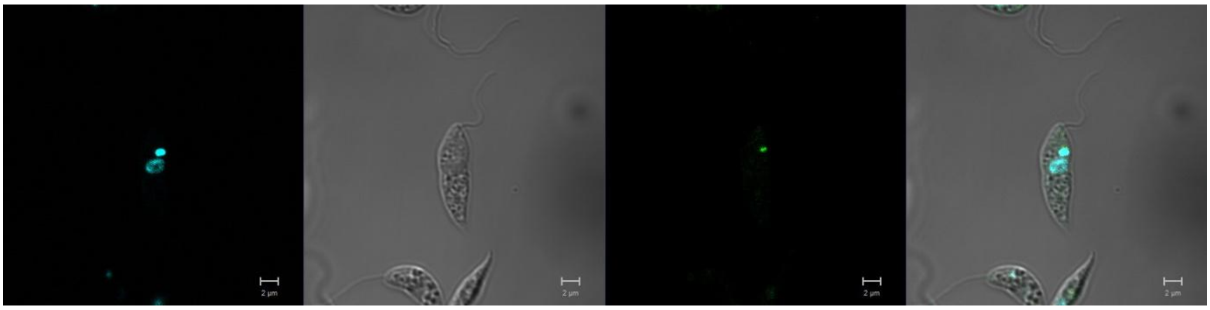

**LmxM.28.3000**
N-terminus

Primary localisation: cytoplasm

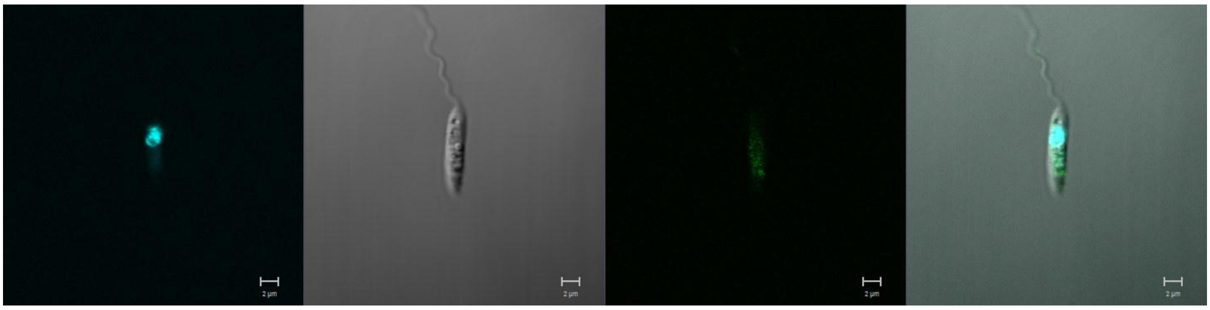

**LmxM.08_29.2570**
N-terminus

Primary localisation: cytoplasm

Secondary localisation: nucleus, flagellum

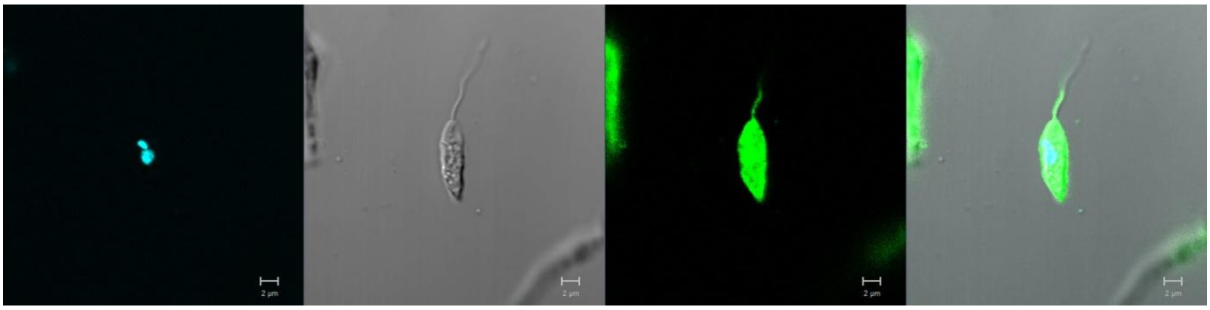

**LmxM.08_29.2670**
N-terminus

Primary localisation: cytoplasm

Secondary localisation: flagellum

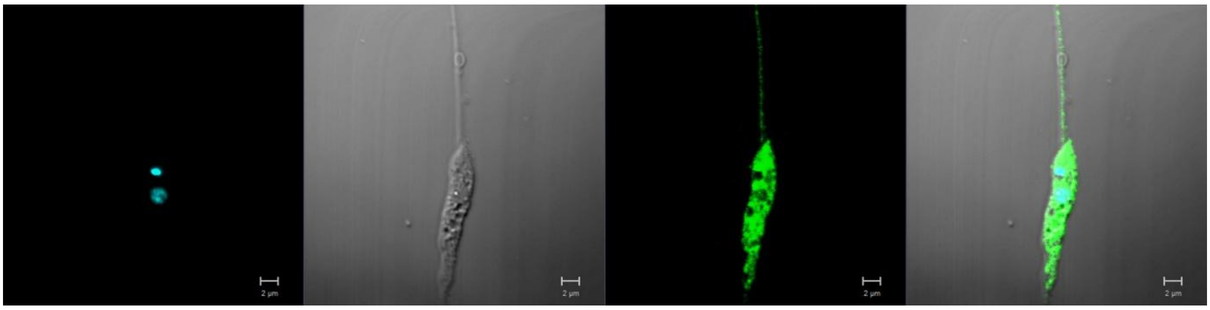

**LmxM.29.2130**
N-terminus

Primary localisation: basal body

Secondary localisation: cytoplasm, flagellum

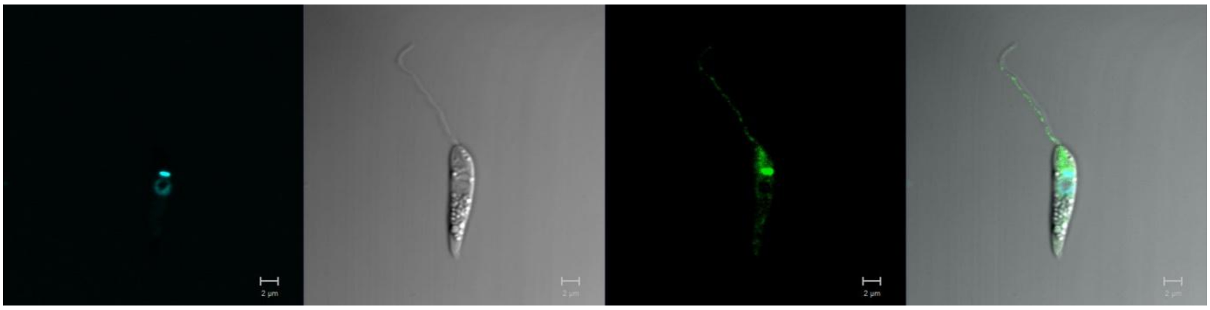

**LmxM.30.2960 (RDK2)**
N-terminus

Primary localisation: cytoplasm

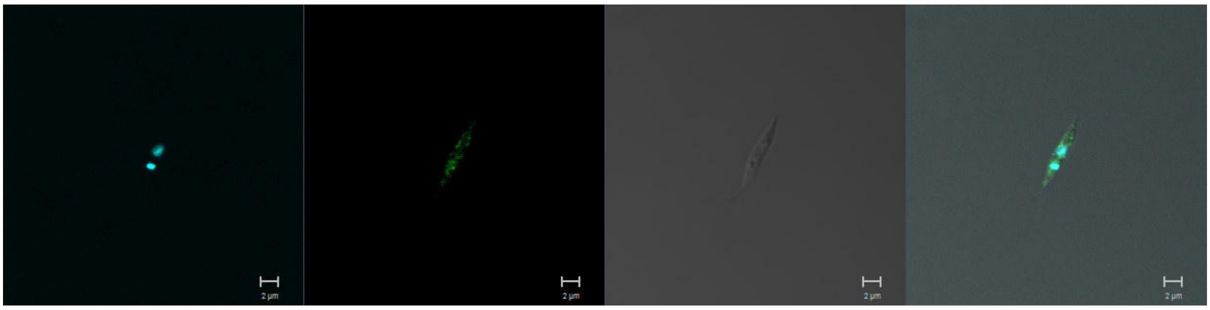

**LmxM.30.3160**
N-terminus

Primary localisation: cytoplasm

Secondary localisation: nucleus, flagellum

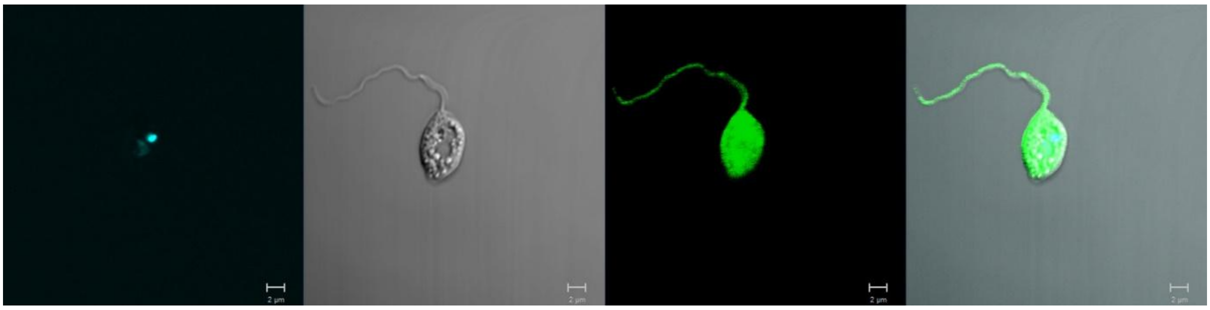

C-terminus

Primary localisation: cytoplasm

Secondary localisation: flagellum, basal body

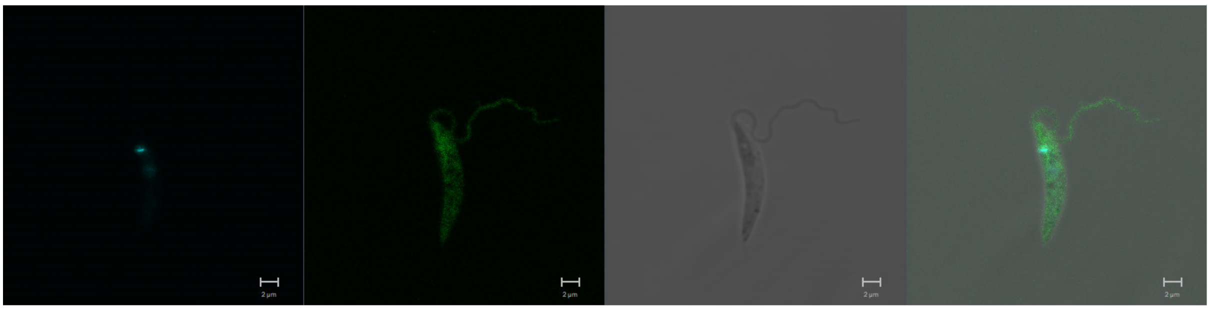

**LmxM.31.0260**
N-terminus

Primary localisation: flagellum

Secondary localisation: basal body, cytoplasm

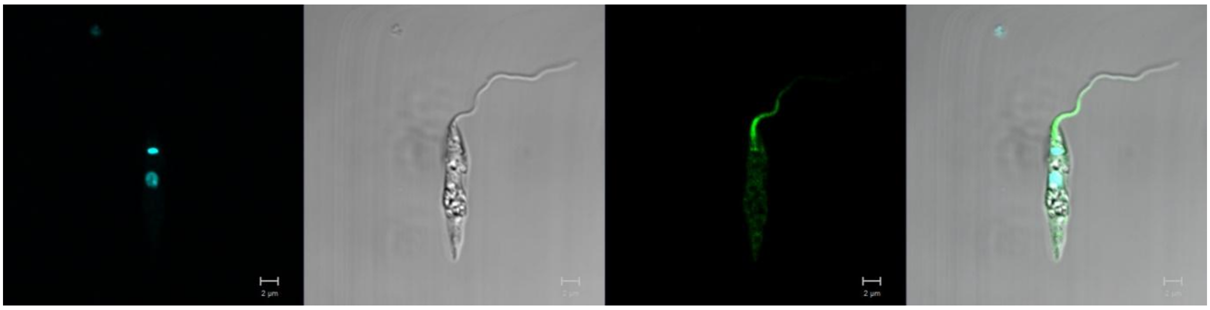

**LmxM.31.1810**
N-terminus

Primary localisation: cytoplasm

Secondary localisation: nucleus, flagellum

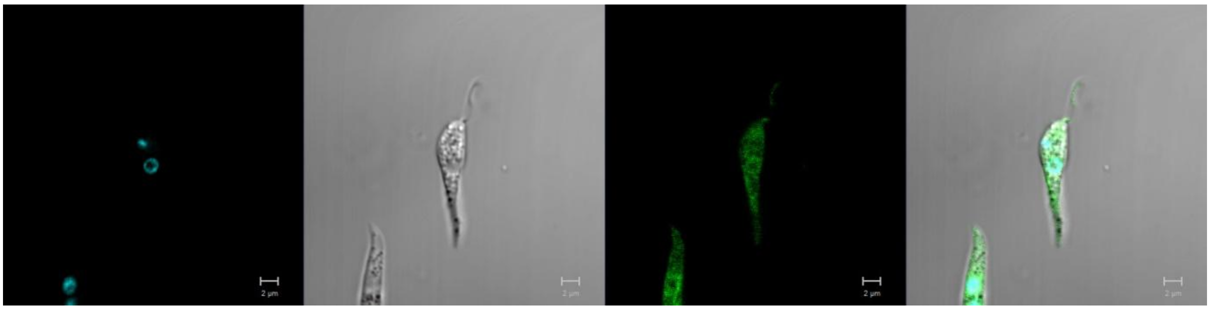

**LmxM.32.1980**
N-terminus

Primary localisation: endomembrane

Secondary localisation: cytoplasm

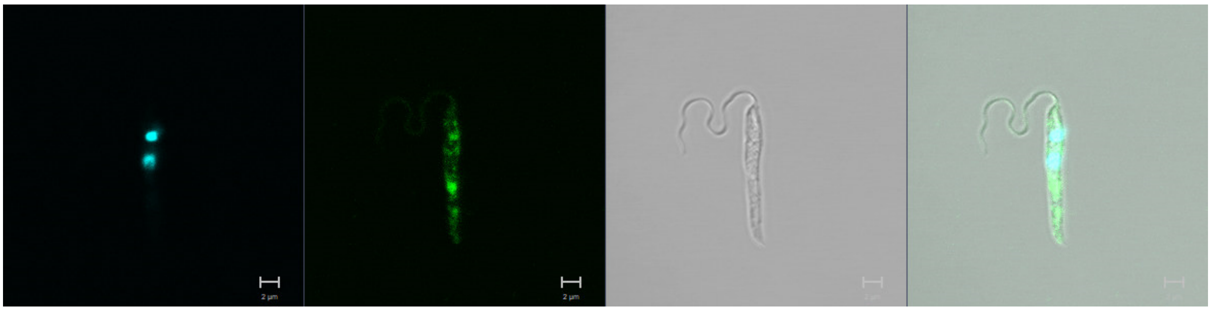

**LmxM.34.5190**
N-terminus

Primary localisation: cytoplasm

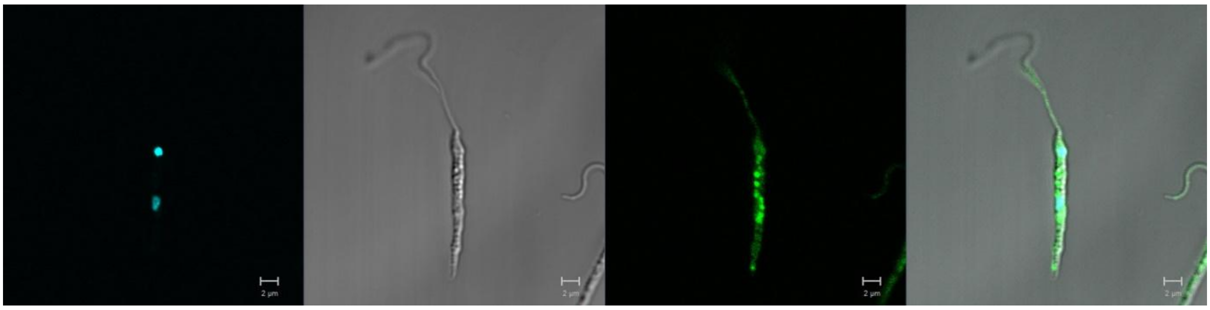

**LmxM.36.1520**
N-terminus

Primary localisation: lysosome

Secondary localisation: cytoplasm, flagellum

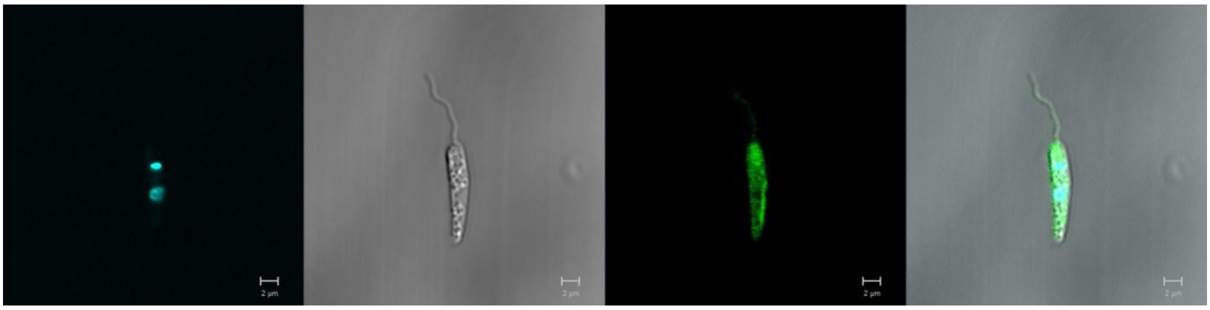

**LmxM.36.1530**
N-terminus

Primary localisation: cytoplasm

Secondary localisation: flagellum, endomembrane

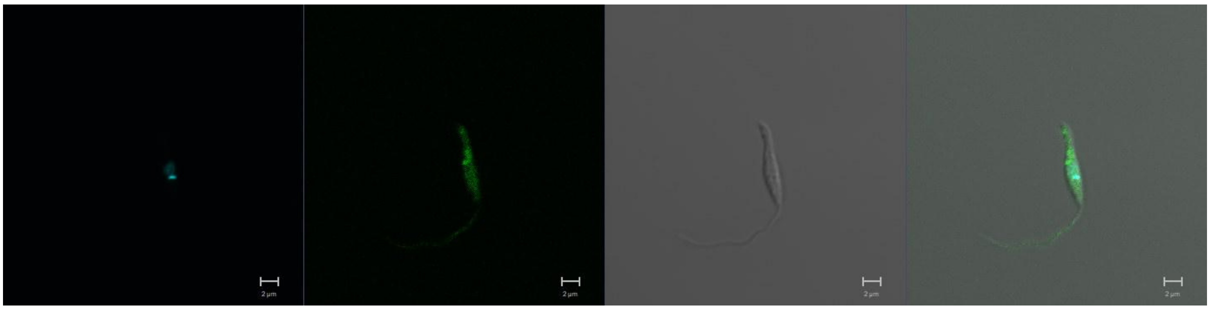

**LmxM.36.2290**
N-terminus

Primary localisation: lysosome

Secondary localisation: cytoplasm

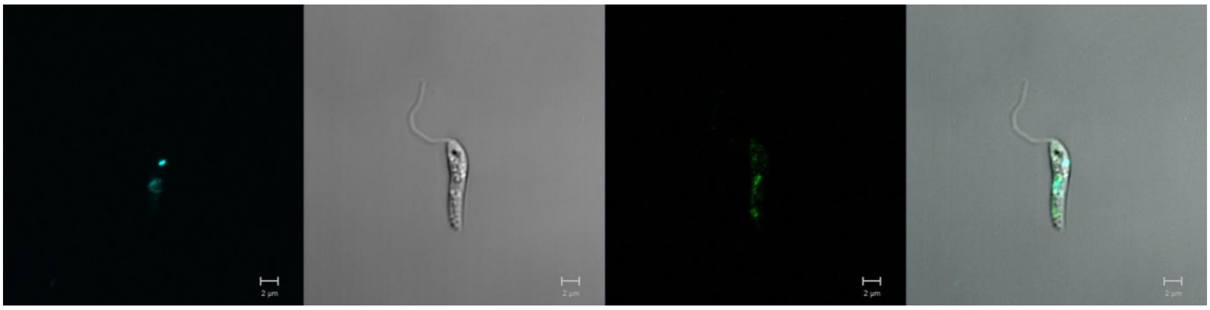

<u>**ORPHAN FAMILY**</u>

**LmxM.02.0570**
N-terminus

Primary localisation: flagellum

Secondary localisation: lysosome, cytoplasm

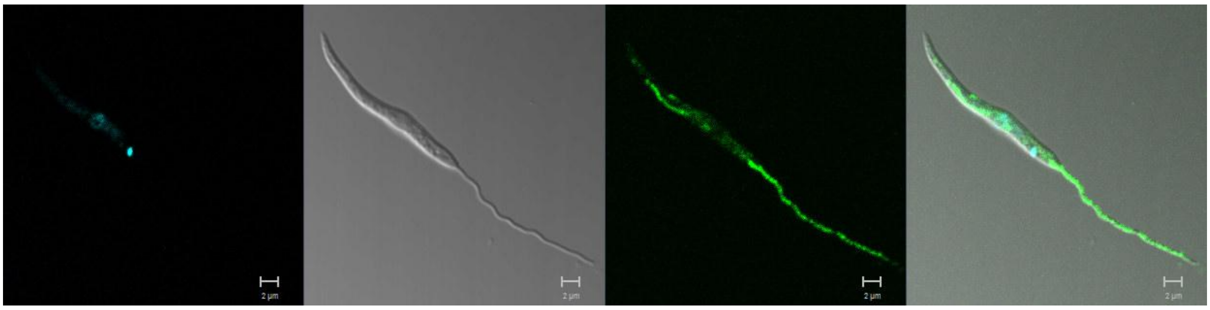

**LmxM.03.0350**
N-terminus

Primary localisation: cytoplasm

Secondary localisation: endomembrane

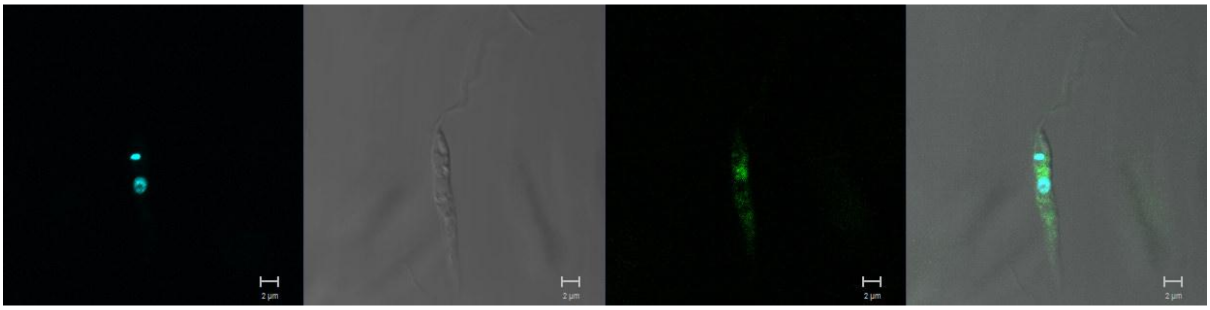

**LmxM.10.0830**
C-terminus

Primary localisation: nucleus

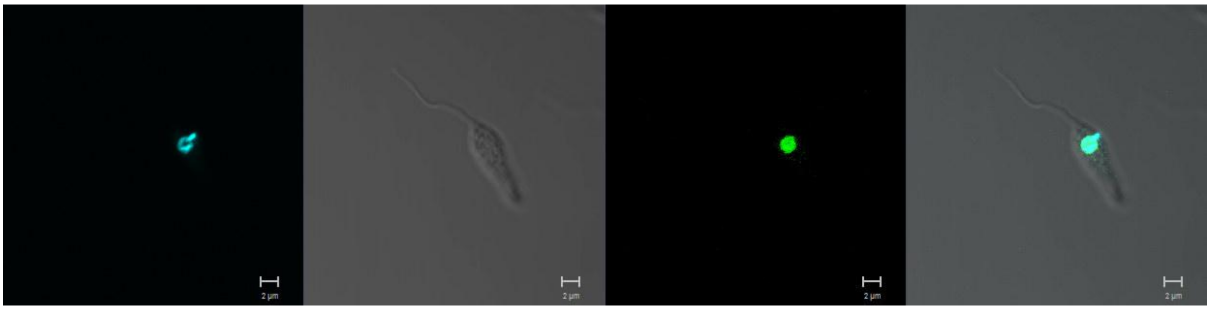

**LmxM.11.0510**
N-terminus

Primary localisation: lysosome

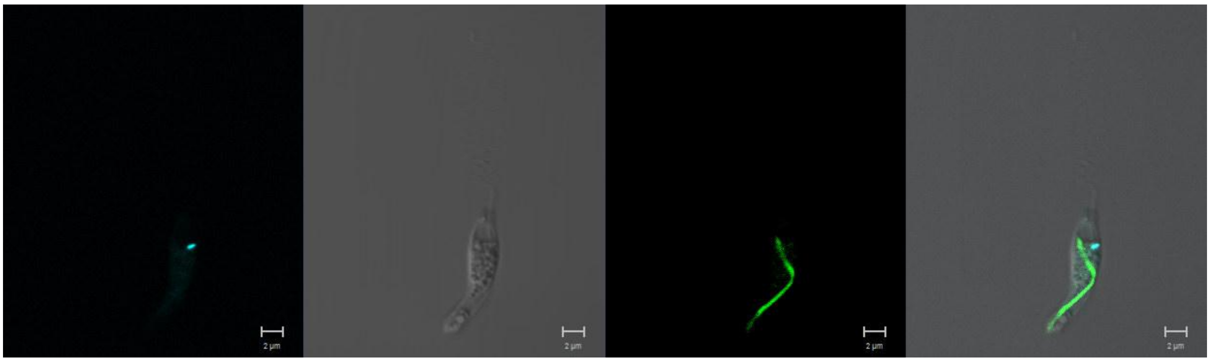

**LmxM.16.0870**
N-terminus

Primary localisation: cytoplasm

Secondary localisation: flagellum, nucleus, kinetoplast

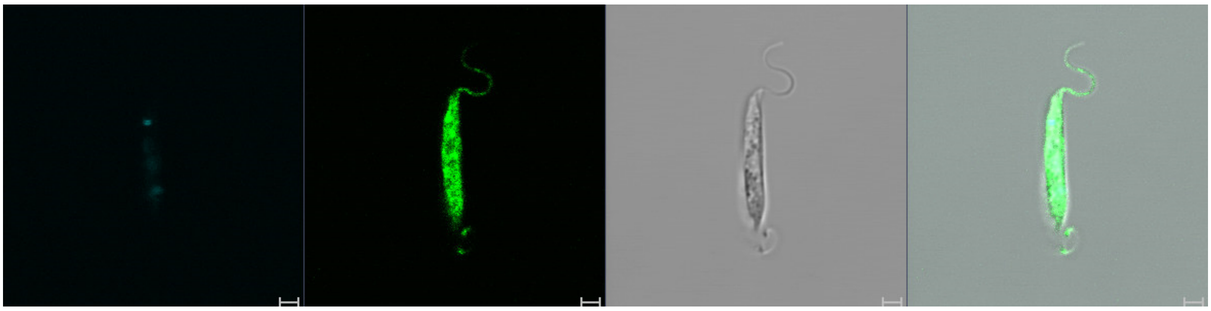

**LmxM.25.1520**
N-terminus

Primary localisation: endomembrane

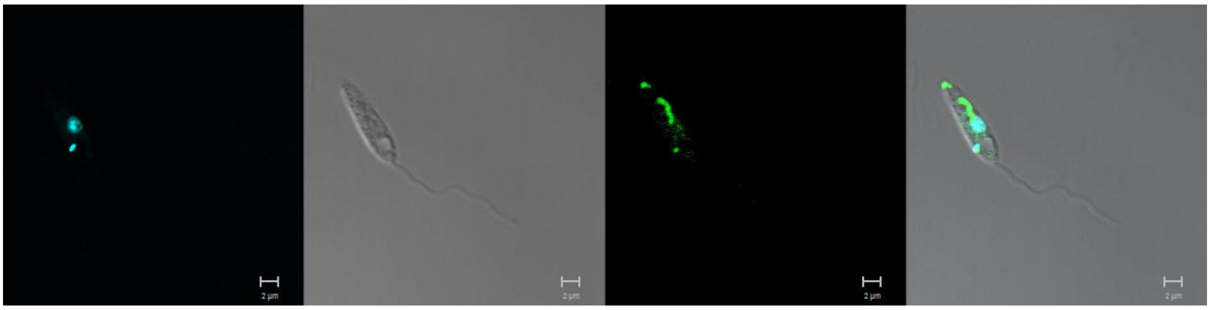

**LmxM.26.2110**
N-terminus

Primary localisation: lysosome

Secondary localisation: cytoplasm

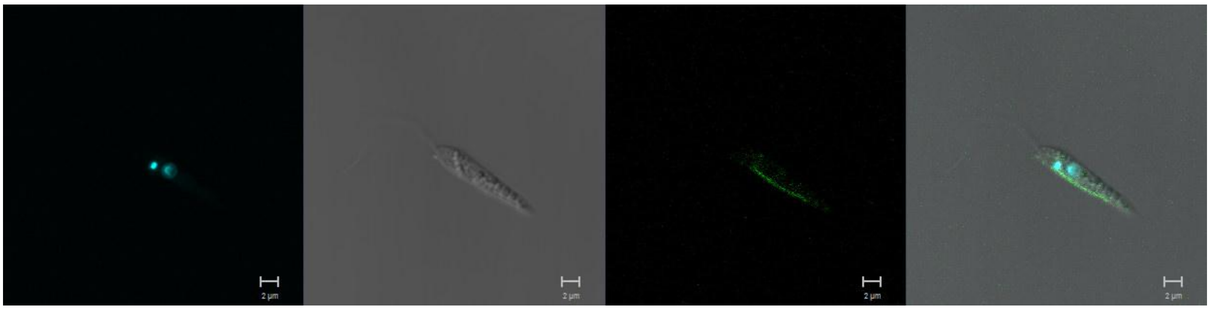

**LmxM.08_29.2490**
N-terminus

Primary localisation: cytoplasmic organelles

Secondary localisation: cytoplasm

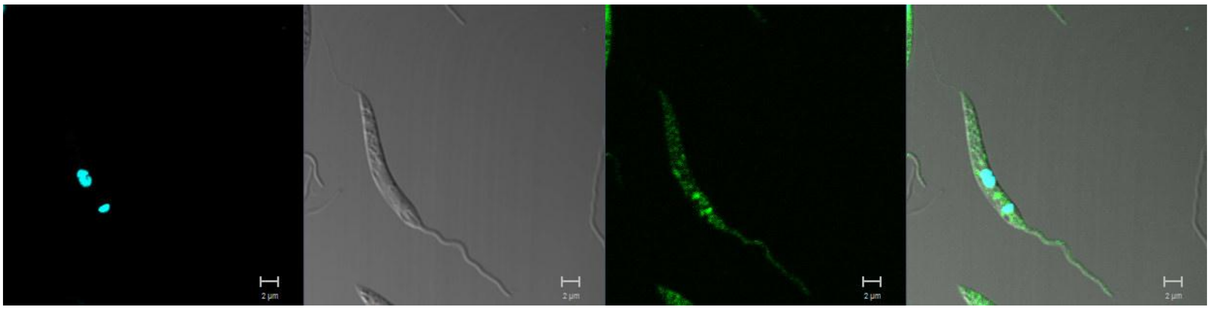

**LmxM.34.4050 (KKT3)**
C-terminus

Primary localisation: nucleus

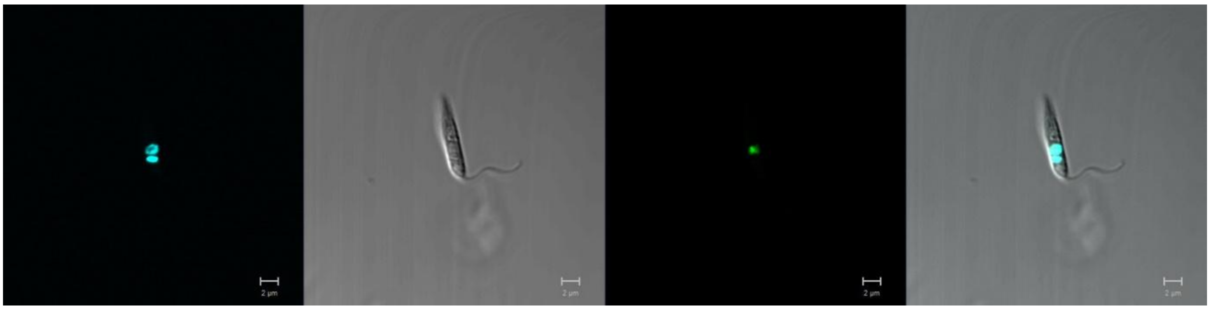

**LmxM.36.5350 (KKT2)**
C-terminus

Primary localisation: nucleus

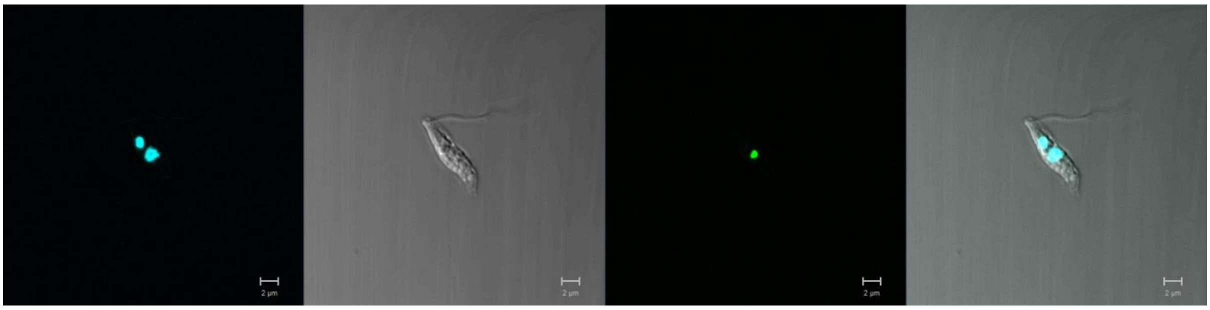

**LmxM.34.4620**
Not tagged
**LmxM.08.0660**
Not tagged
**LmxM.26.2060**
Not tagged
**LmxM.28.1650**
Not tagged
**LmxM.31.1290**
Not tagged
**LmxM.33.2190**
Not tagged

**<u>OTHERS FAMILY</u>**

**LmxM.20.1330**
N-terminus

Primary localisation: endomembrane

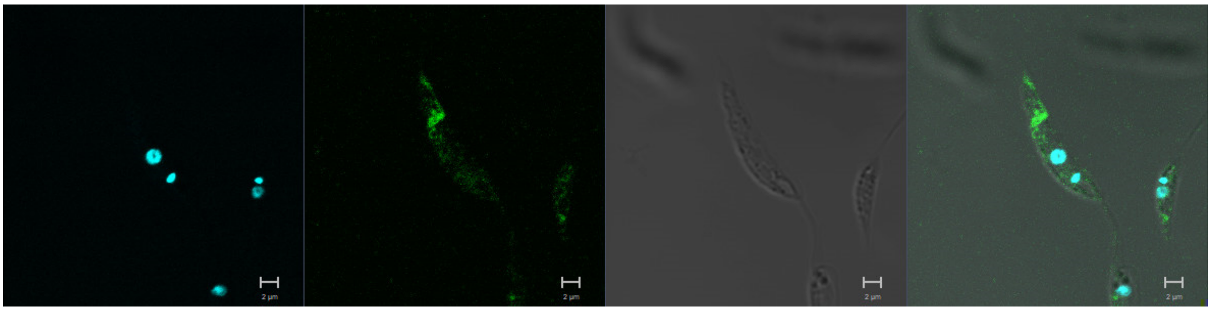

**LmxM.20.1340**
N-terminus

Primary localisation: cytoplasm

Secondary localisation: endomembrane

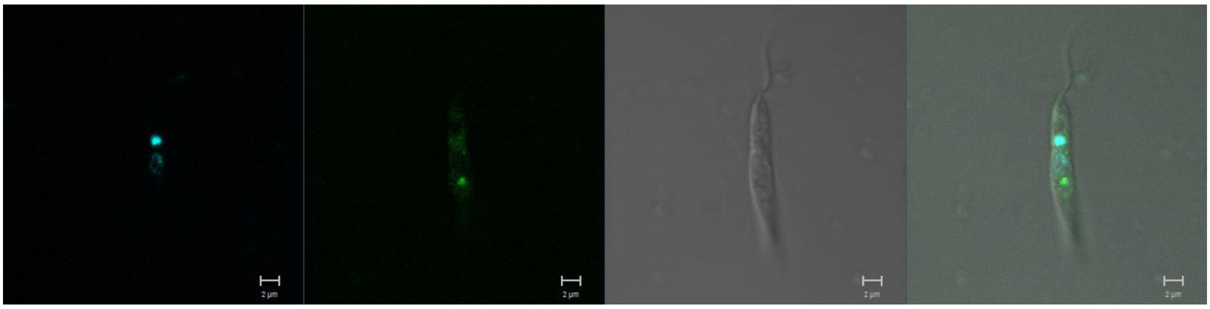

**LmxM.08_29.0370**
N-terminus

Primary localisation: cytoplasm

Secondary localisation: flagellum, endomembrane

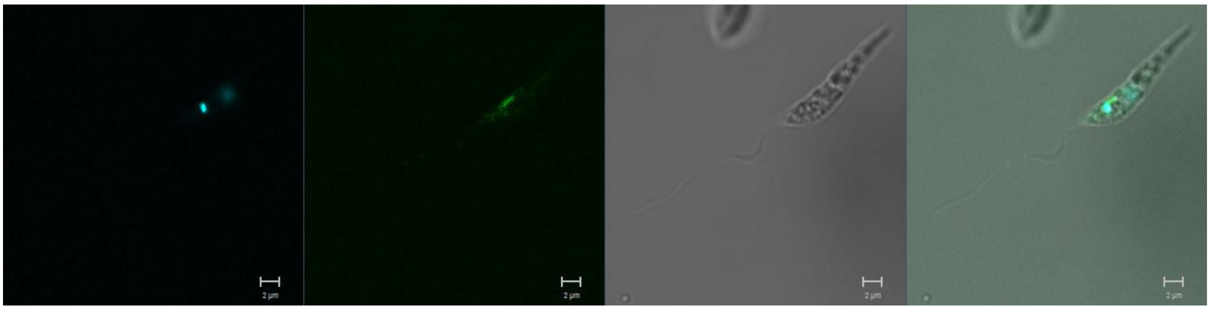

**LmxM.34.2870**
N-terminus

Primary localisation: cytoplasm

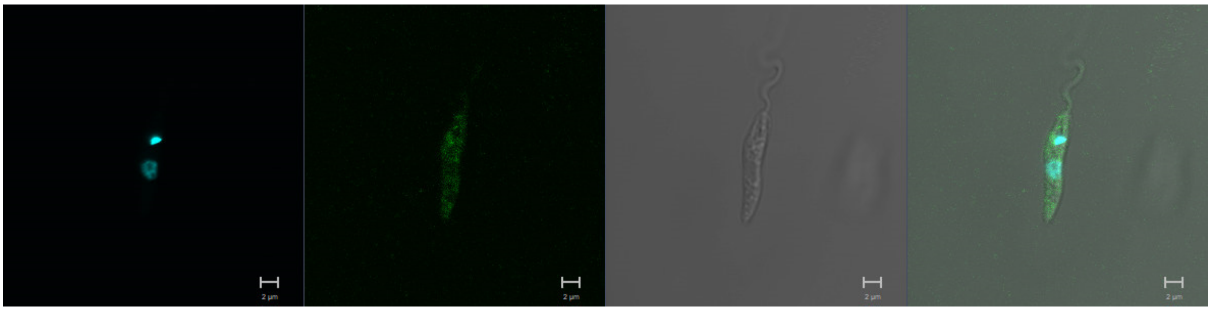

**LmxM.26.2440 (AUK3)**
N-terminus

Primary localisation: nucleus

Secondary localisation: endomembrane

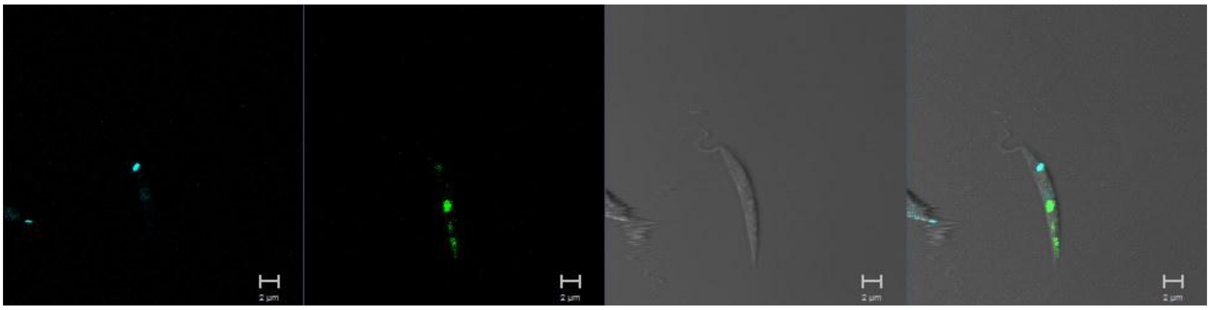

**LmxM.28.0520 (AIRK)**
N-terminus

Primary localisation: nucleus

Secondary localisation: endomembrane

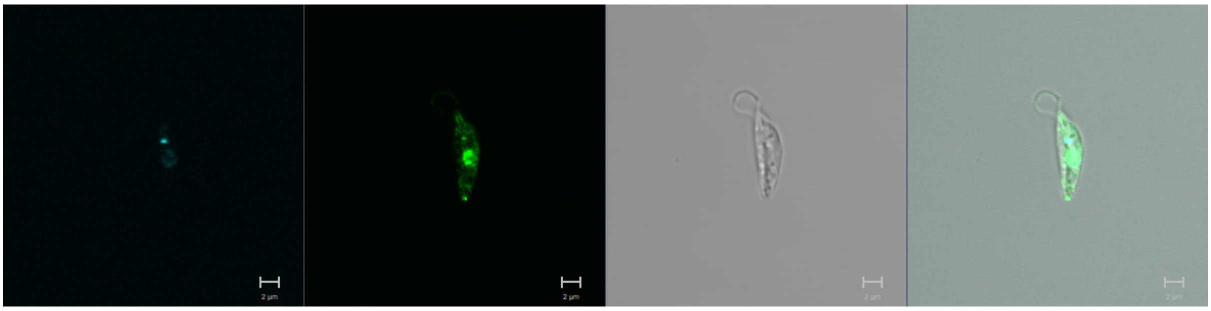

**LmxM.08_29.1330 (AUK2)**
N-terminus

Primary localisation: cytoplasm

Secondary localisation: endomembrane

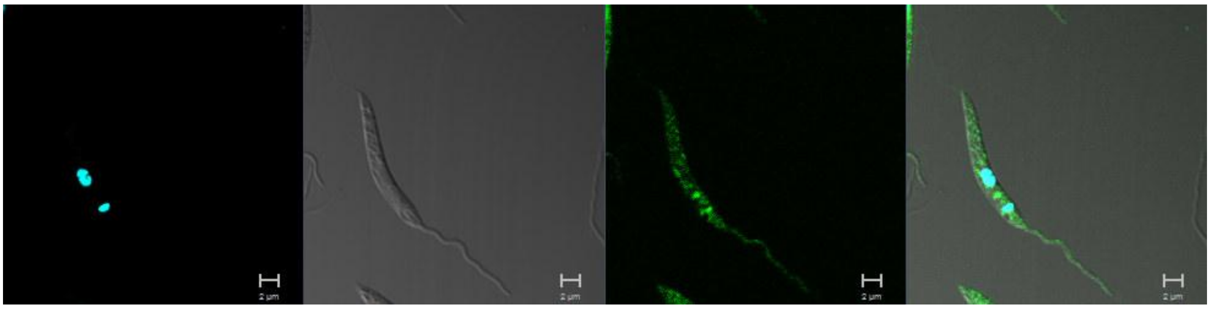

**LmxM.19.0590**
N-terminus

Primary localisation: cytoplasm

Secondary localisation: endomembrane

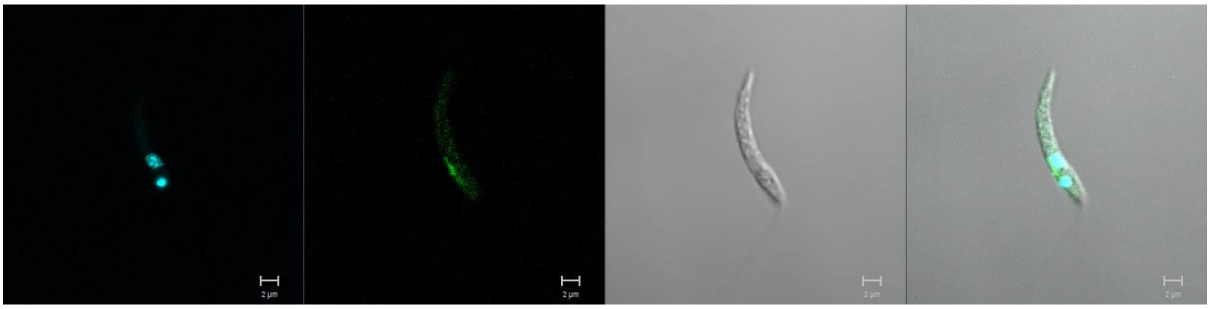

**LmxM.11.0250**
N-terminus

Primary localisation: cytoplasm

Secondary localisation: lysosome

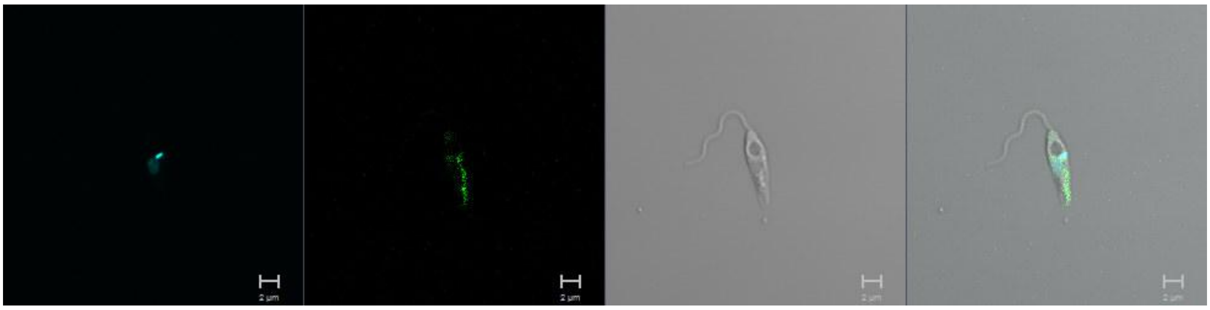

**LmxM.24.1730**
N-terminus

Primary localisation: cytoplasm

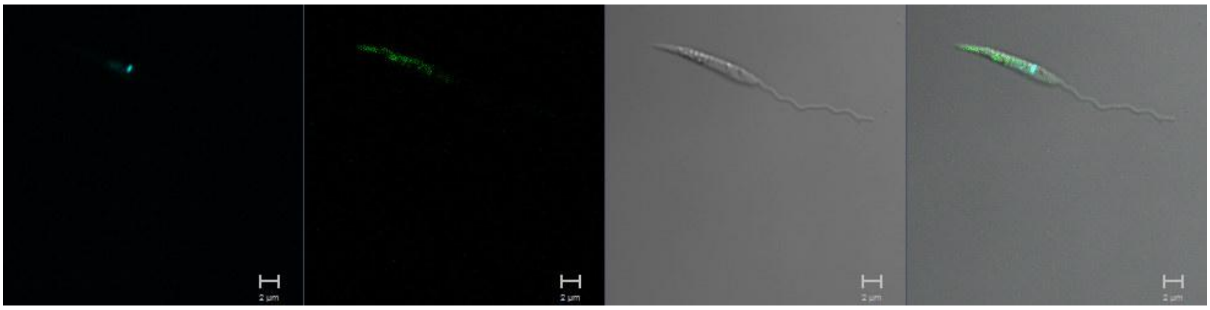

**LmxM.34.2320**
N-terminus

Primary localisation: flagellum

Secondary localisation: cytoplasm

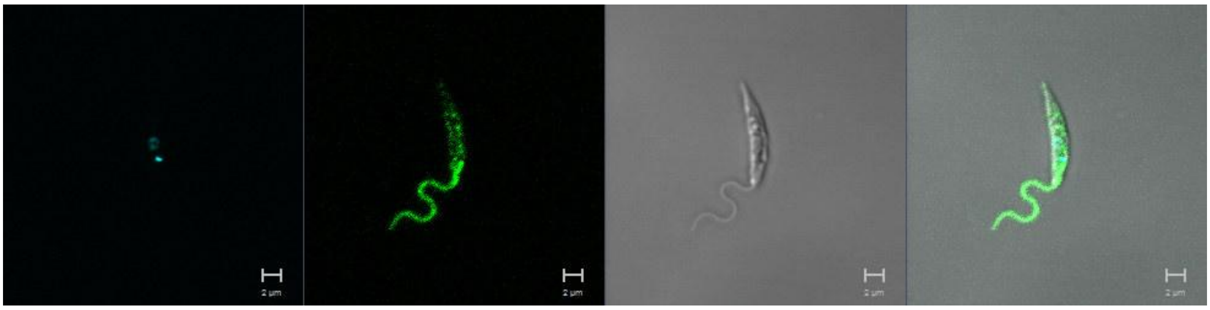

**LmxM.34.1730 (CK2A1)**
N-terminus

Primary localisation: cytoplasm

Secondary localisation: nucleus, kinetoplast

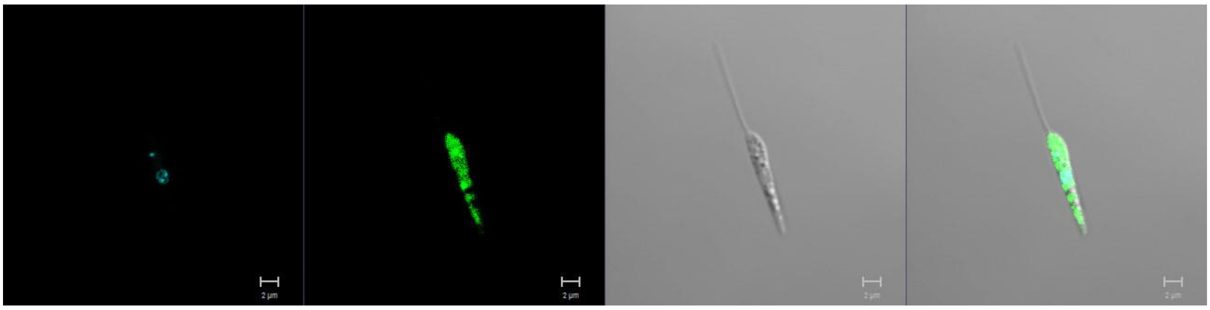

**LmxM.02.0360 (CK2A2)**
N-terminus

Primary localisation: nucleus

Secondary localisation: lysosome, cytoplasm

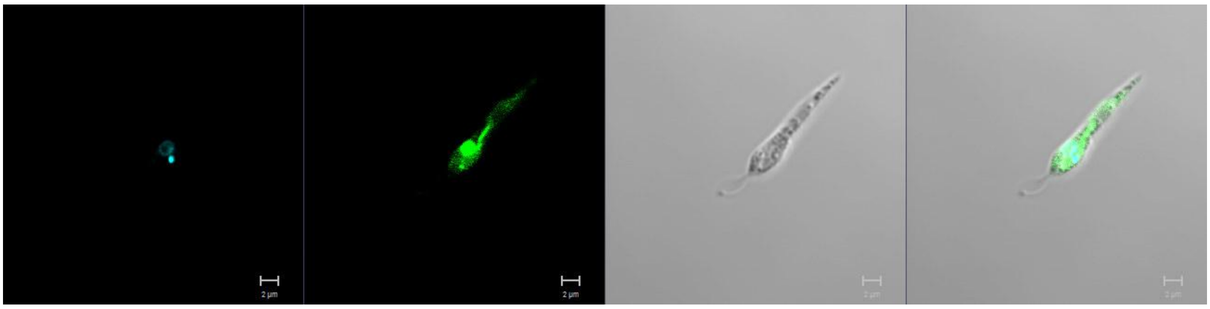

**LmxM.15.0770**
N-terminus

Primary localisation: cytoplasm

Secondary localisation: flagellum, endomembrane

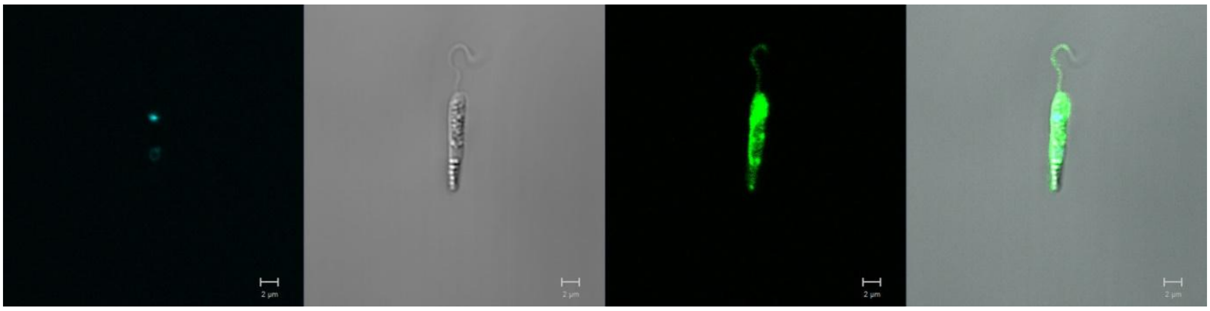

**LmxM.33.0030**
N-terminus

Primary localisation: cytoplasm

Secondary localisation: flagellum

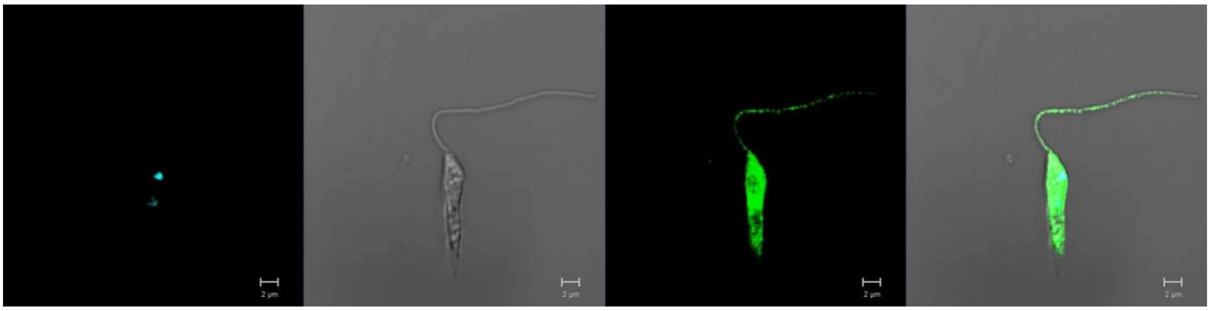

**LmxM.11.0060 (EF2A1)**
N-terminus

Primary localisation: cytoplasm

Secondary localisation: endoplasmic reticulum

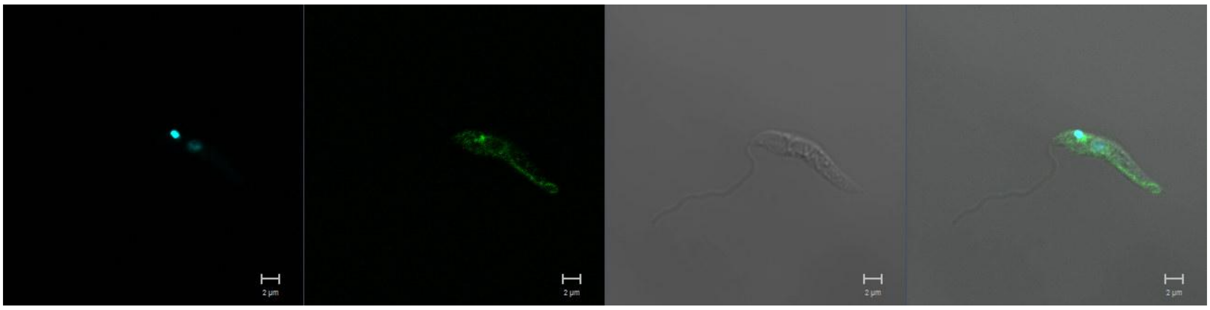

**LmxM.29.1560 (EF2A3)**
N term - N-terminus

Primary localisation: cytoplasm (low expression)

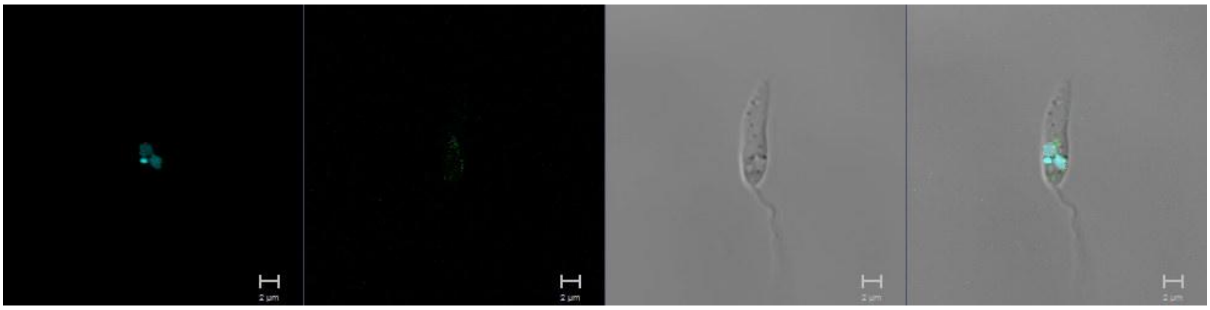

**LmxM.33.2150 (EF2A2)**
N-terminus

Primary localisation: cytoplasmic organelles

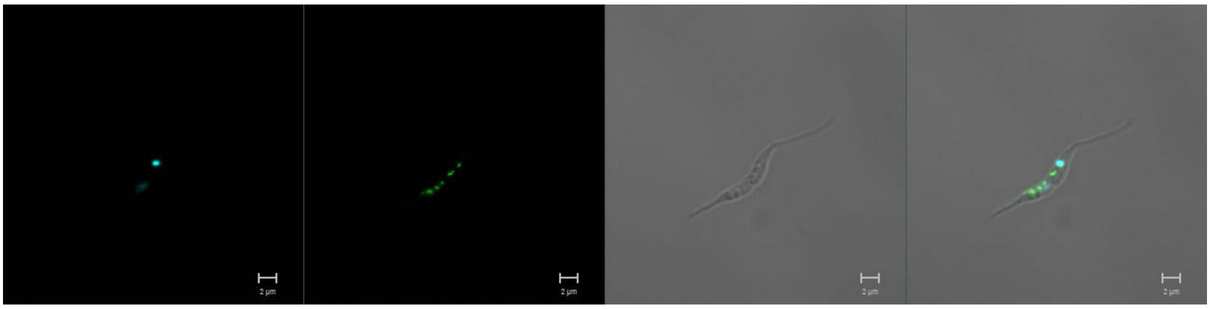

**LmxM.17.0790 (PKL)**
N-terminus

Primary localisation: cytoplasm (low expression)

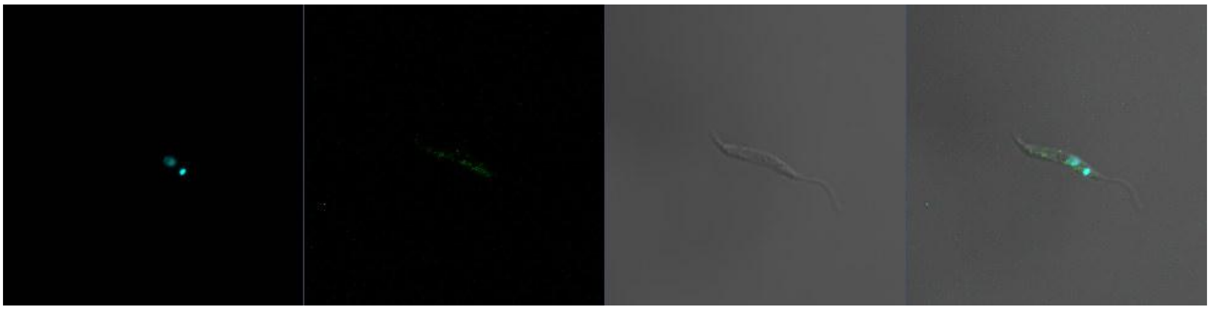

C-terminus

Primary localisation: basal body, endomembrane, cytoplasm (specific to early G1-phase)

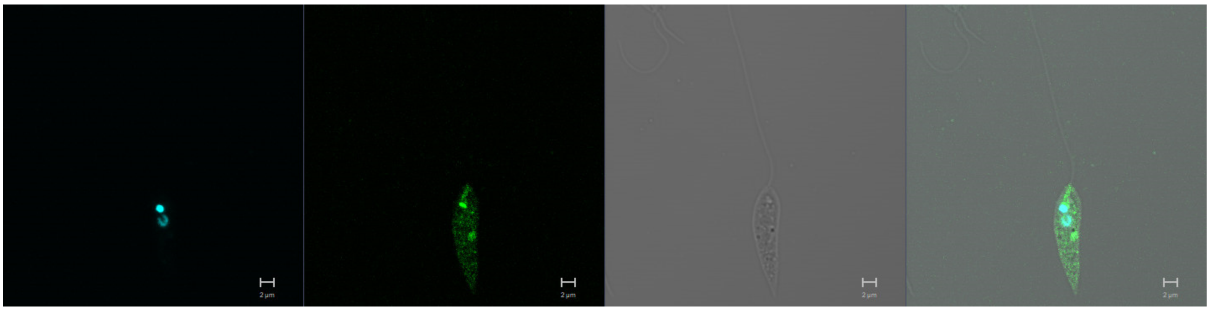

**LmxM.30.2860 (TLK)**
N-terminus

Primary localisation: cytoplasm (low expression)

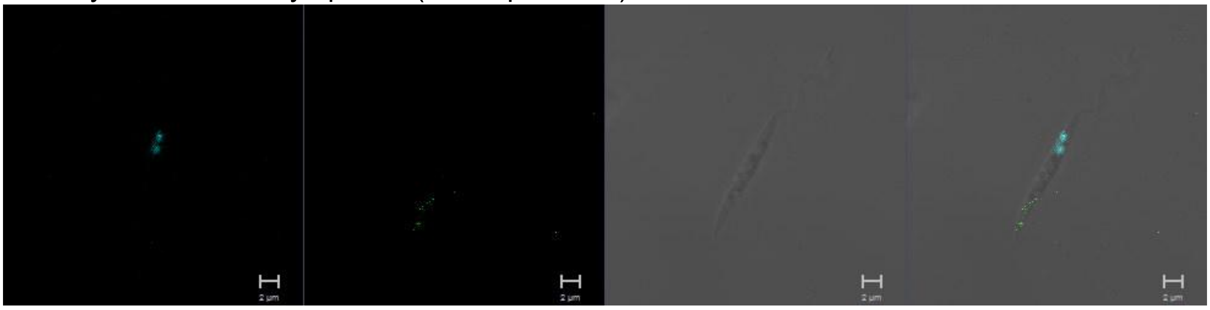

C-terminus

Primary localisation: nucleus

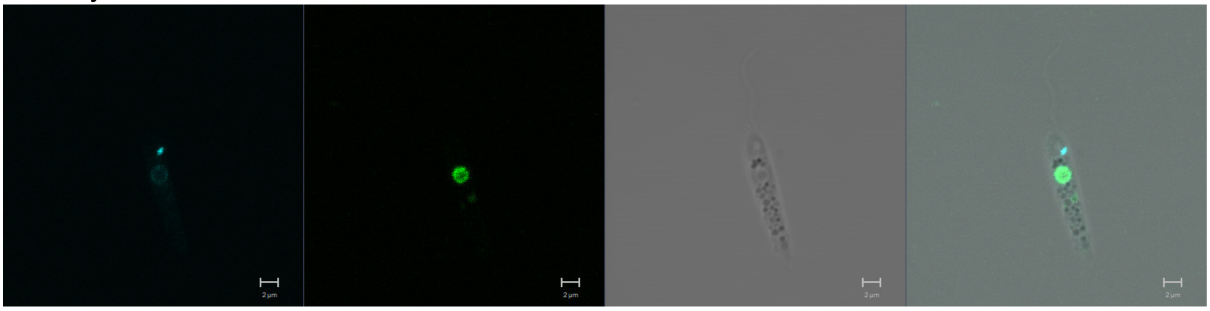

**LmxM.28.0620 (ULK4)**
N-terminus

Primary localisation: no signal

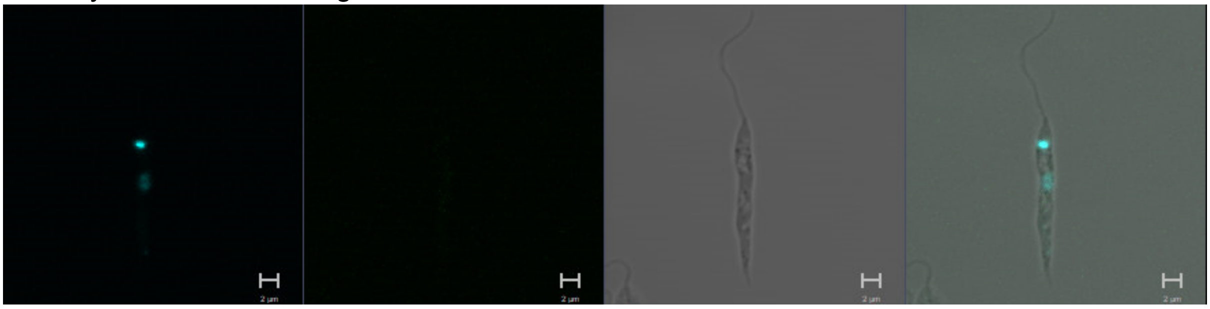

C-terminus

Primary localisation: basal body

Secondary localisation: flagellum tip, cytoplasm, flagellum, nucleus

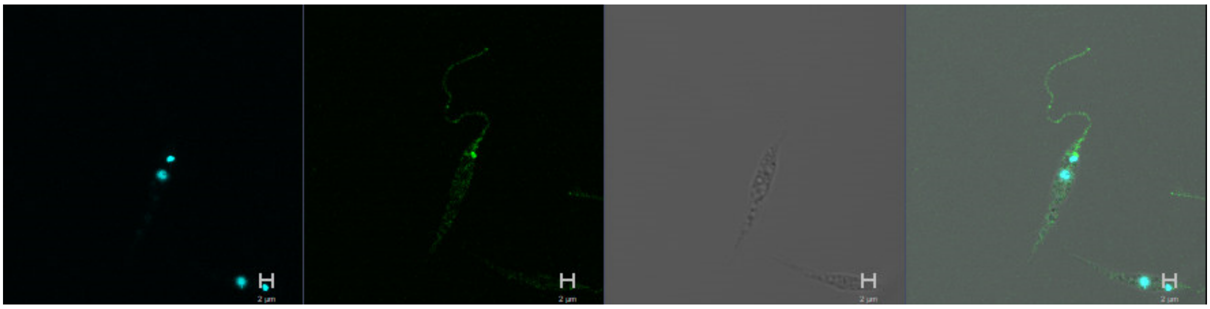

**LmxM.13.0440 (STK36)**
N-terminus

Primary localisation: no signal

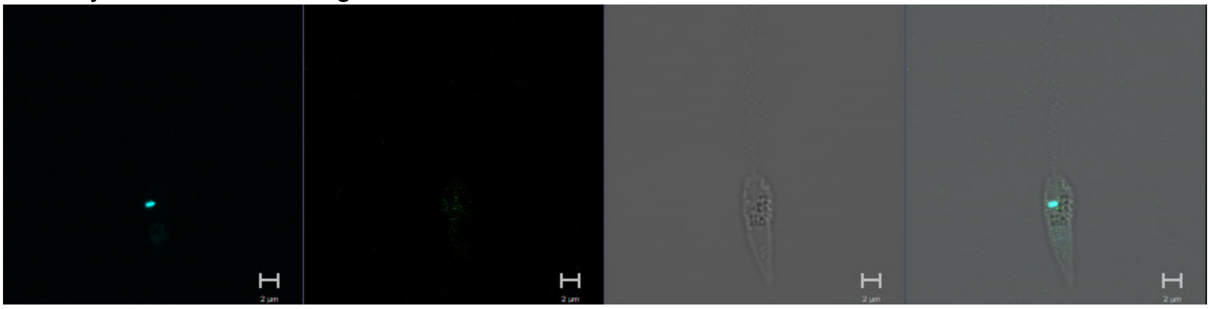

C-terminus

Primary localisation: basal body

Secondary localisation: flagellum tip, cytoplasm, flagellum, nucleus

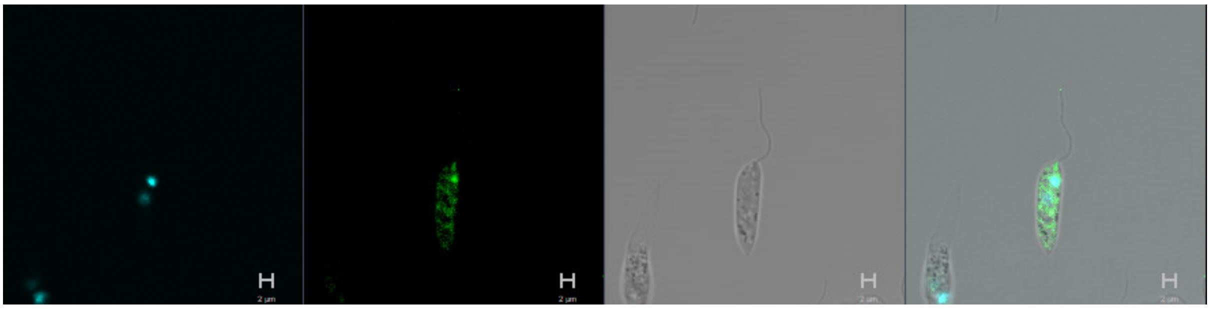

**LmxM.28.1760**
N-terminus

Primary localisation: failed attempts

**LmxM.08_29.2720**
N-terminus

Primary localisation: cytoplasm

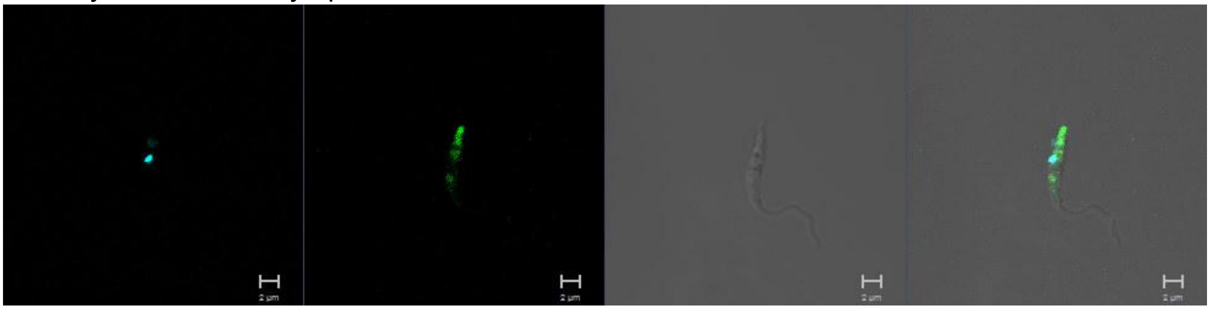

**LmxM.33.0940 (WEE1)**
N-terminus

Primary localisation: cytoplasm

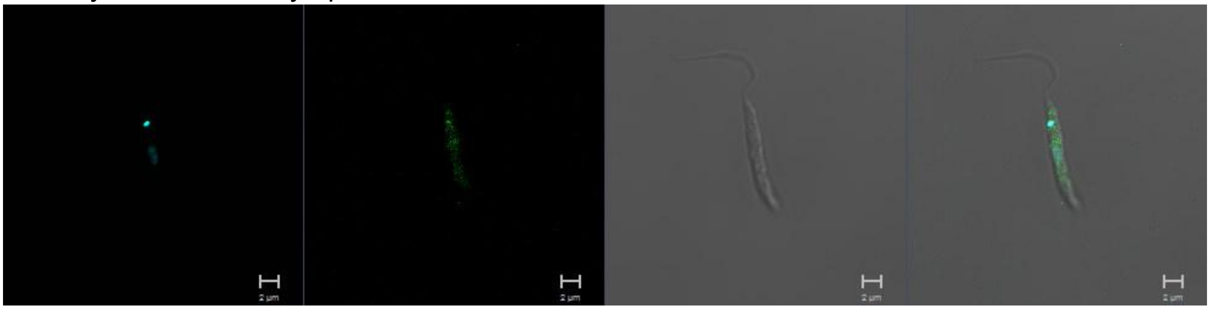

**LmxM.08.0530**

N-terminus

Primary localisation: endomembrane

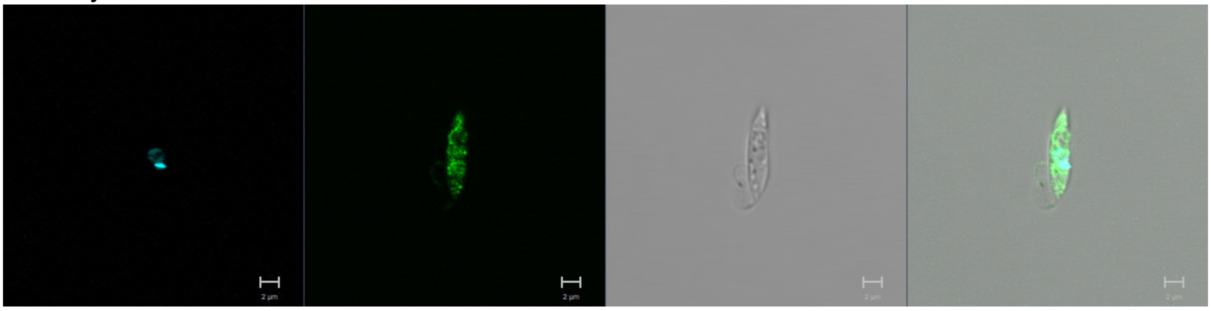

**LmxM.20.0960**

N-terminus

Primary localisation: cytoplasm (low expression)

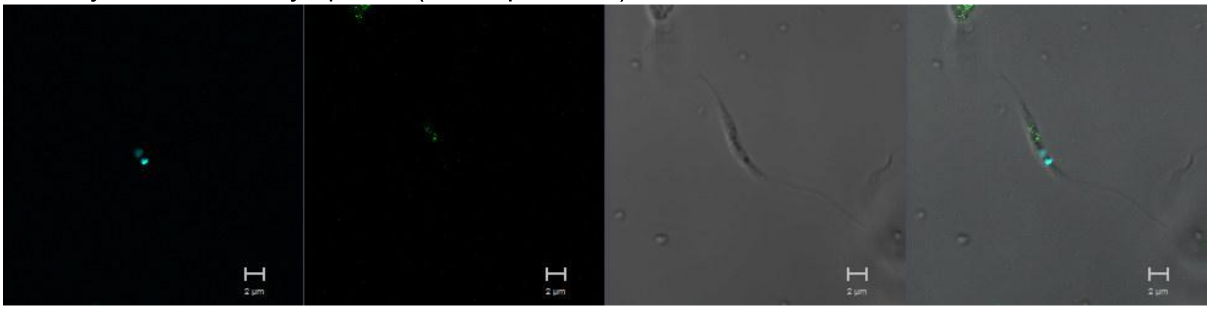

C-terminus

Primary localisation: endomembrane

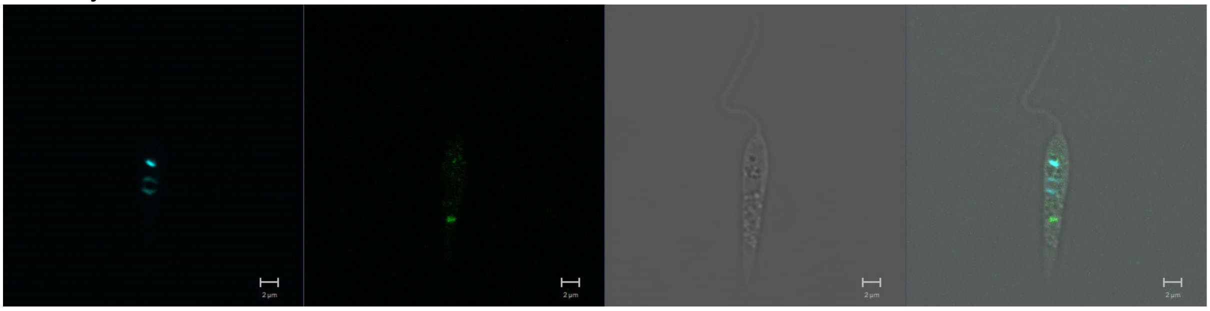

**LmxM.21.0823**
N-terminus

Primary localisation: lysosome

Secondary localisation: endomembrane

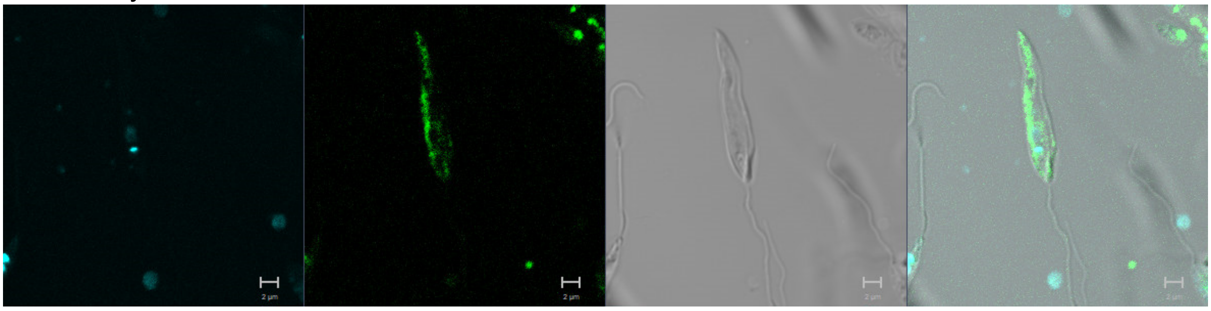

**LmxM.36.2630**
N-terminus

Primary localisation: endomembrane

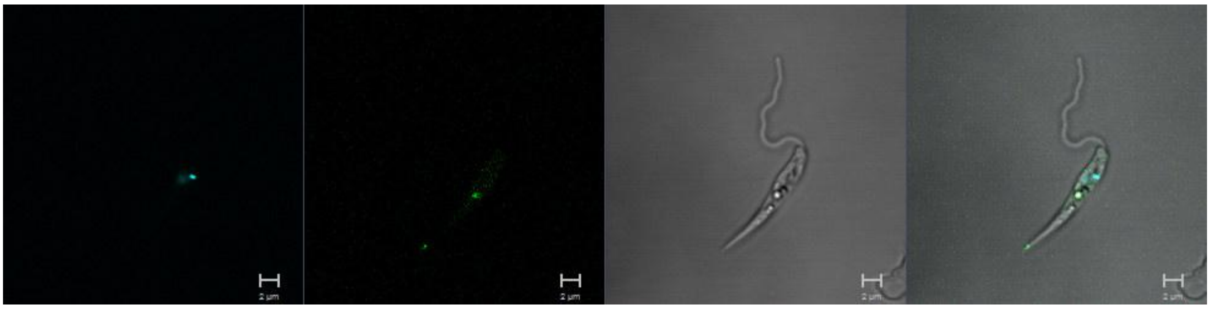

**LmxM.22.1150**
N-terminus

Primary localisation: cytoplasm

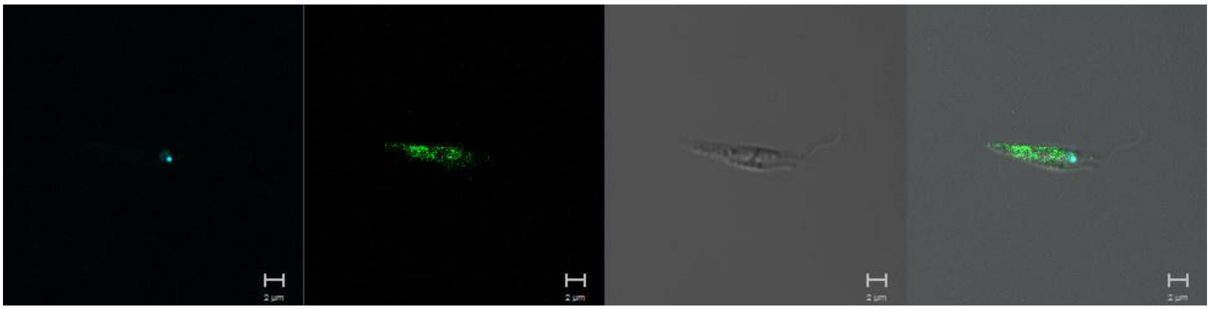

<u>**ATYPICAL PROTEIN KINASES**</u>

**LmxM.08_29.1450 (PIK)**
N-terminus

Primary localisation: basal body

Secondary localisation: cytoplasm

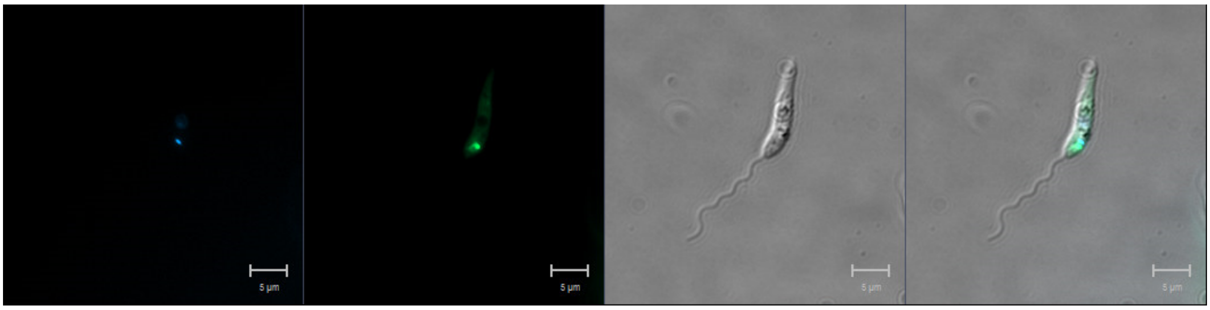

N-terminus

Primary localisation: lysosome

Secondary localisation: endomembrane

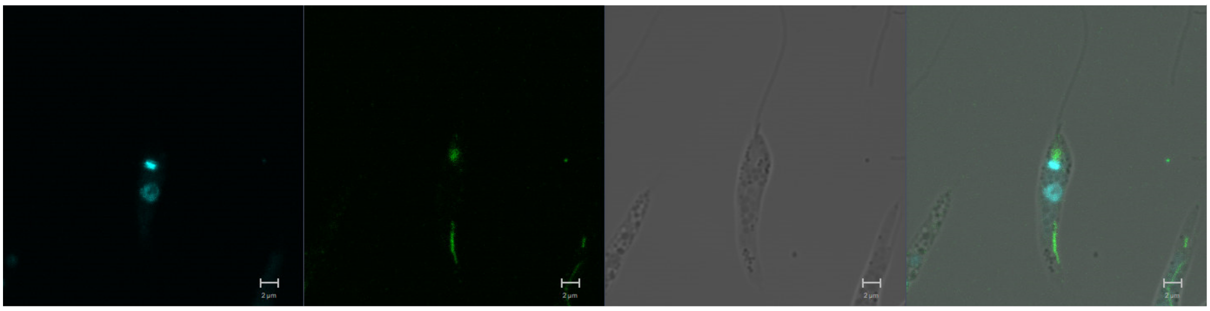

**LmxM.24.2010 (PI3K)**
N-terminus

Primary localisation: cytoplasm

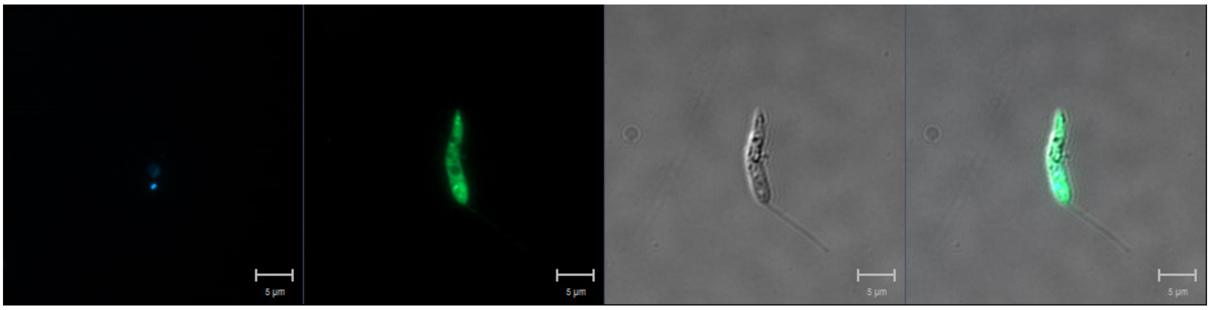

**LmxM.33.3590 (PI4K)**
N-terminus

Primary localisation: cytoplasm

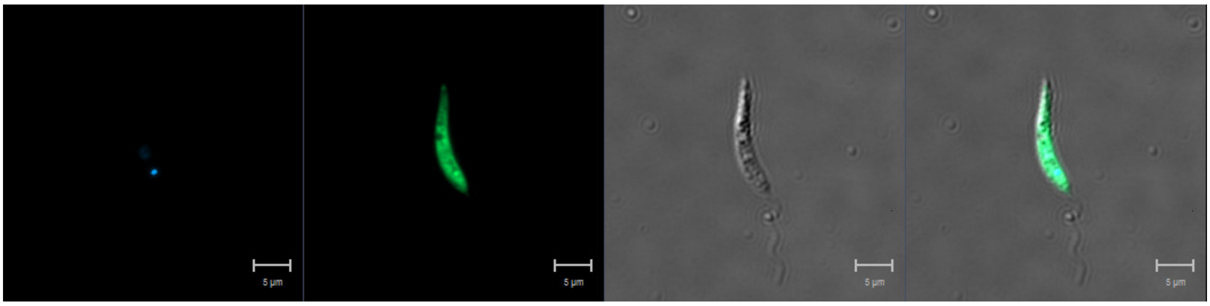


**LmxM.27.0890 (PI3,5K)**
N-terminus

Primary localisation: cytoplasm

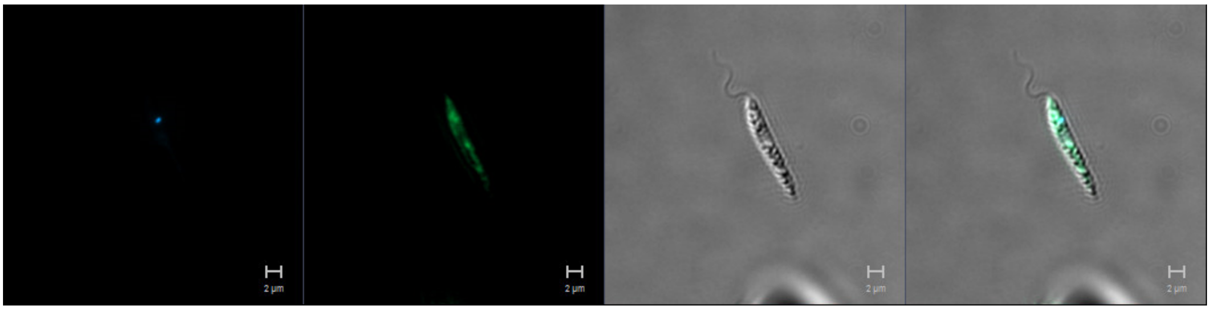


**LmxM.33.3090 (PIPKalpha)**
C-terminus

Primary localisation: cytoplasm

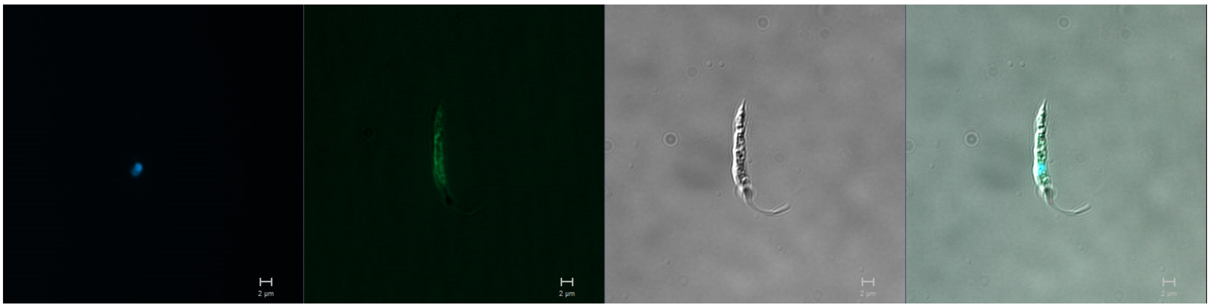

**LmxM.34.0560 (PIP5K)**
N-terminus

Primary localisation: flagellar pocket

Secondary localisation: cytoplasm

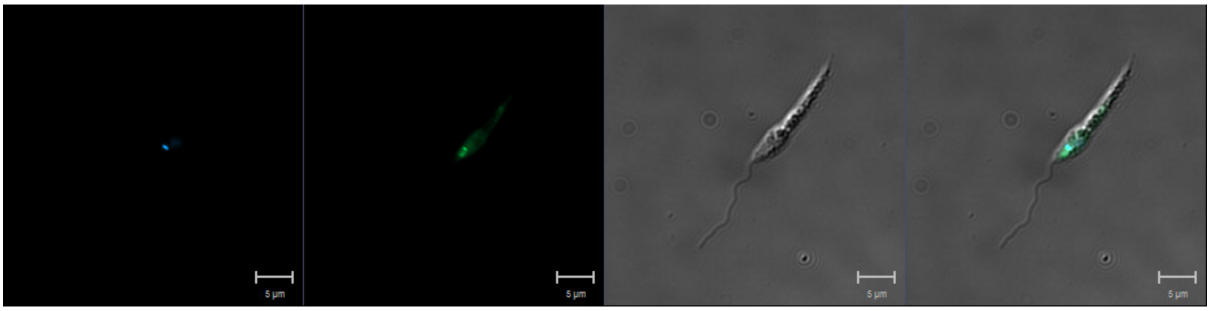

**LmxM.36.6320 (TOR1)**
N-terminus

Primary localisation: cytoplasm

Secondary localisation: endomembrane

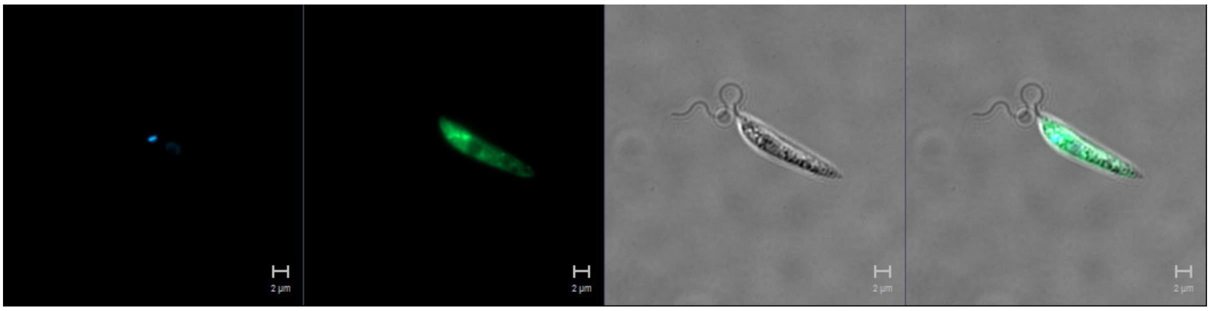

**LmxM.33.4530 (TOR2)**
N-terminus

Primary localisation: cytoplasmic organelles

Secondary localisation: cytoplasm

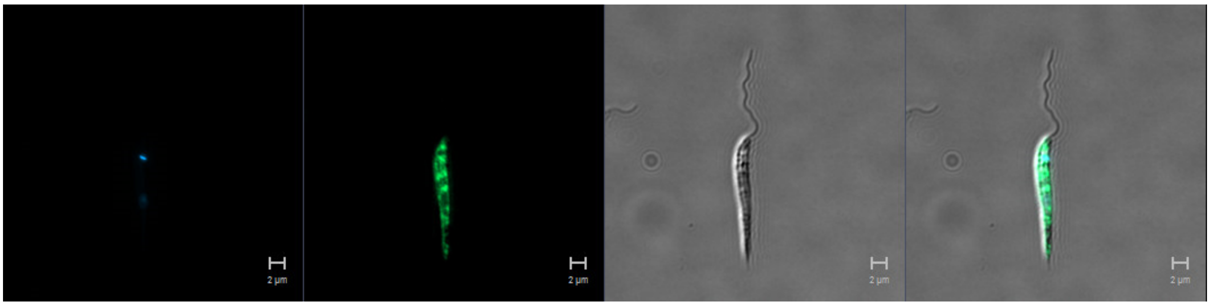

**LmxM.33.3940 (TOR3)**
N-terminus

Primary localisation: cytoplasm

Secondary localisation: kinetoplast

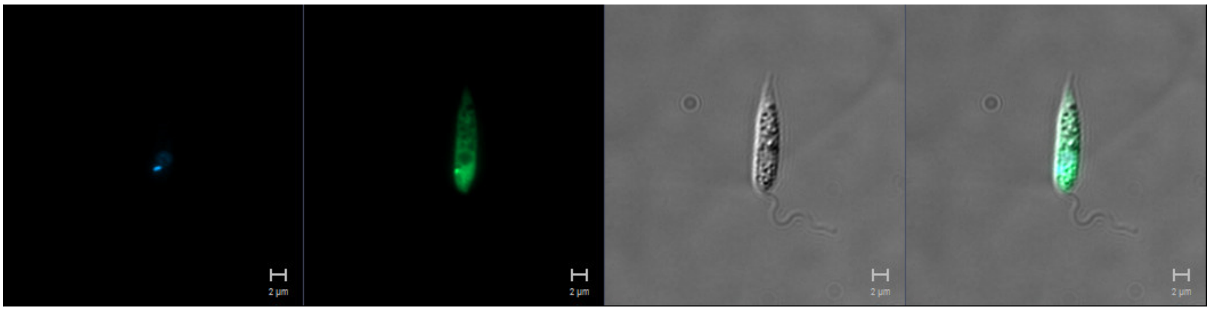

**LmxM.02.0120 (ATM)**
N-terminus

Primary localisation: nucleus

Secondary localisation: cytoplasm

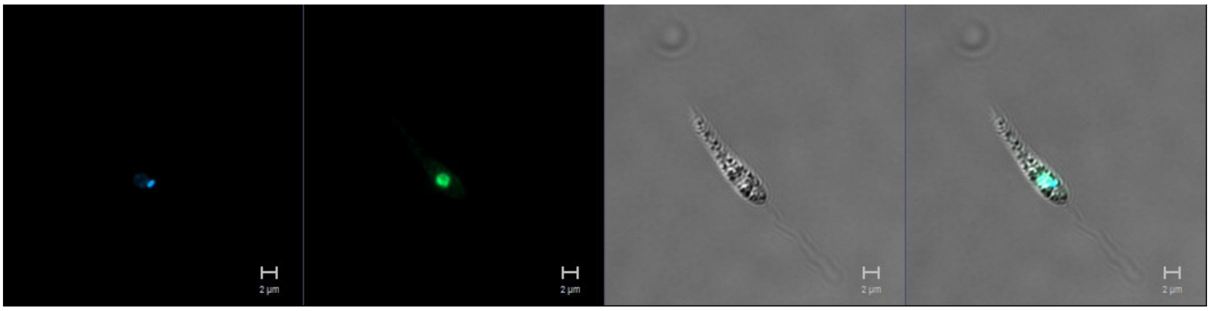

**LmxM.31.1460 (ATR)**
C-terminus

Primary localisation: cytoplasm

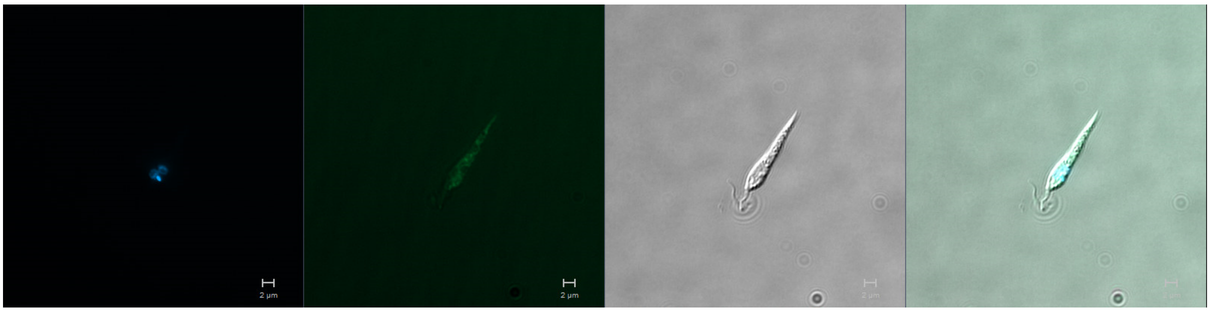
