## Supplementary Data 3 for "Systematic functional analysis of *Leishmania* protein kinases identifies regulators of differentiation or survival"

### Supplementary Data 3a

#### Heat maps for Axenic amastigote data

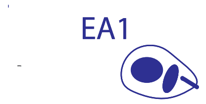


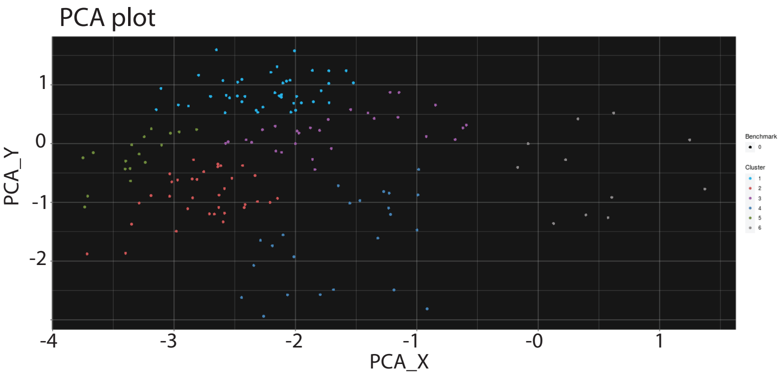


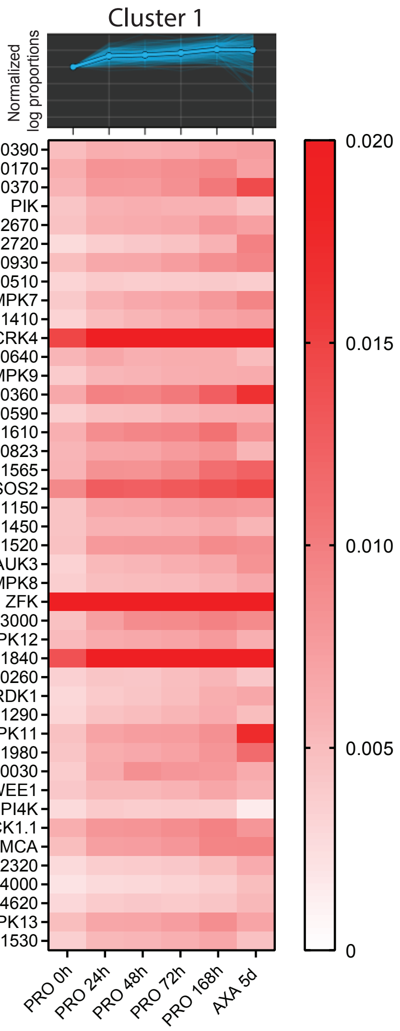


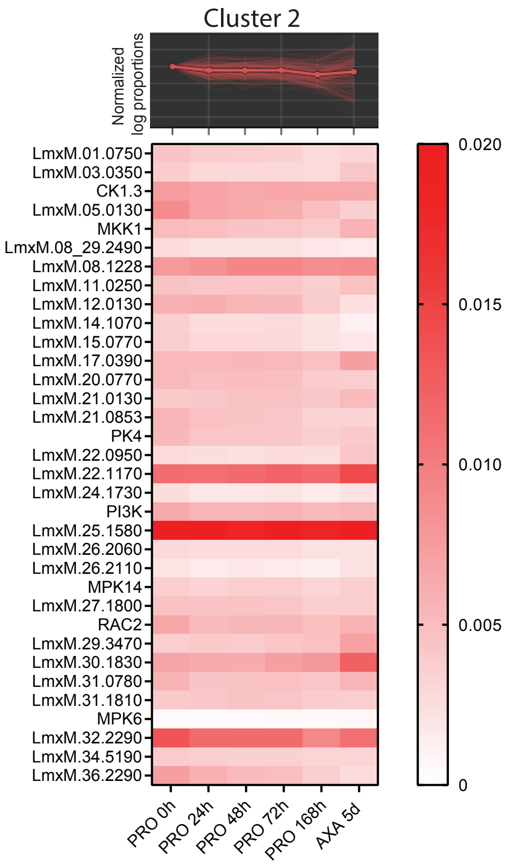


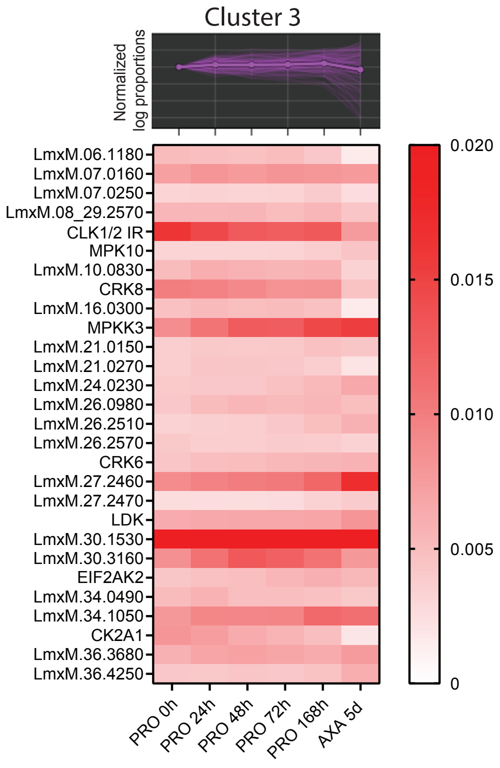


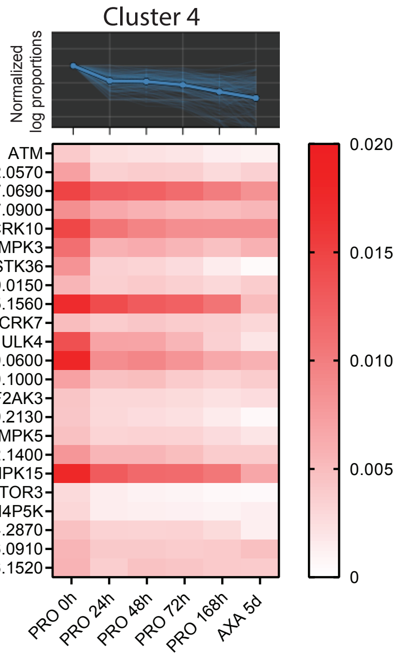


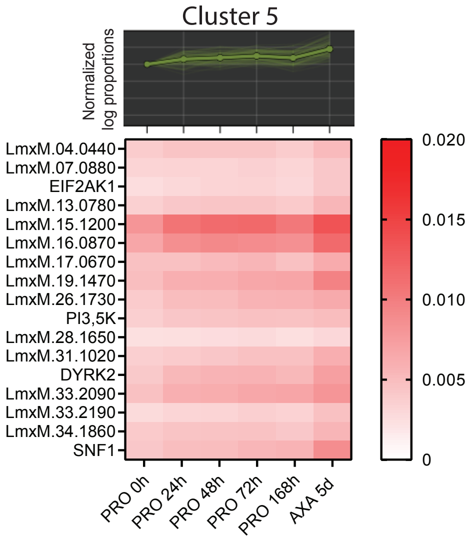


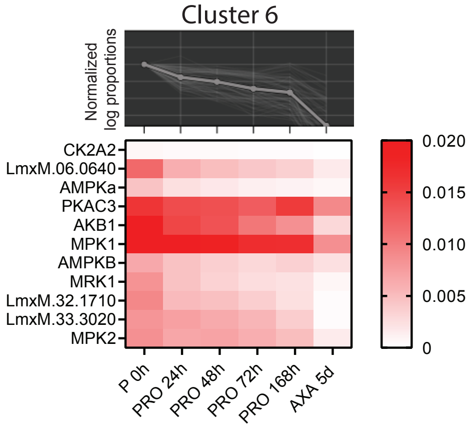

### Supplementary Data 3b
Individual plots of Axenic amastigote data

EA1

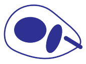

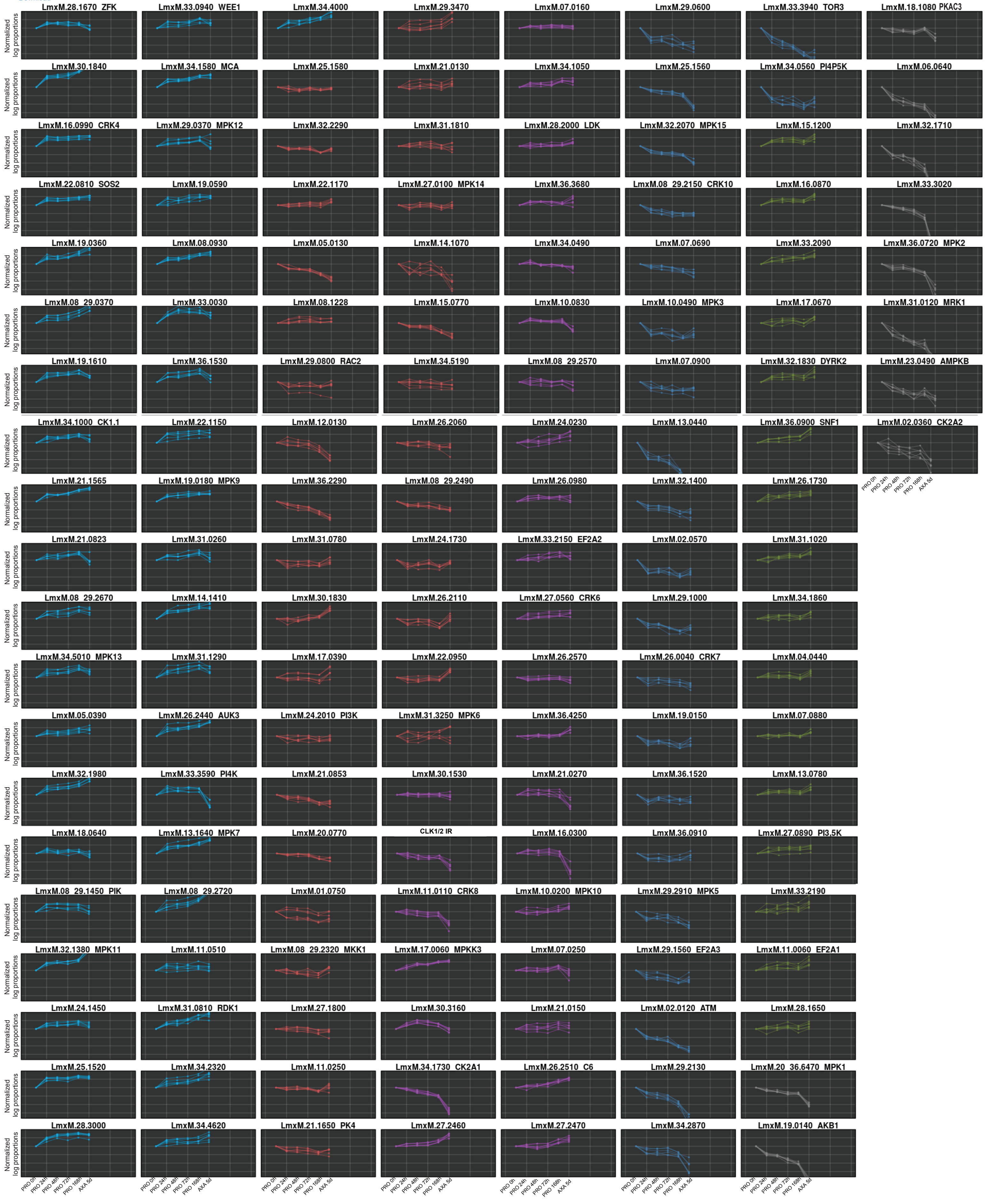

### Supplementary Data 3c

Heat maps for macrophage infection data

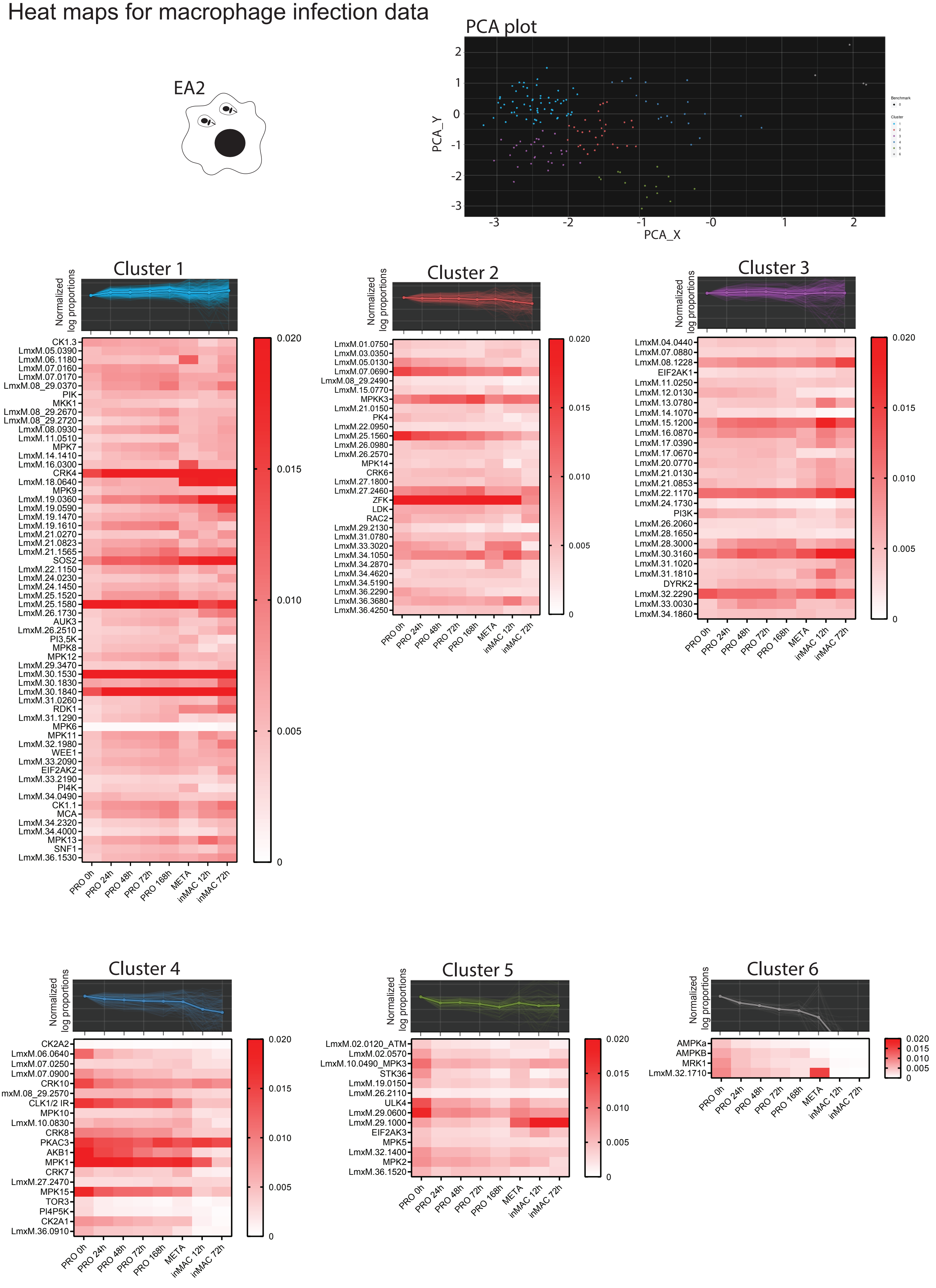

### Supplementary Data 3d
Individual plots of macrophage infection data

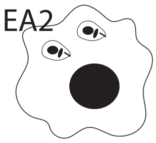


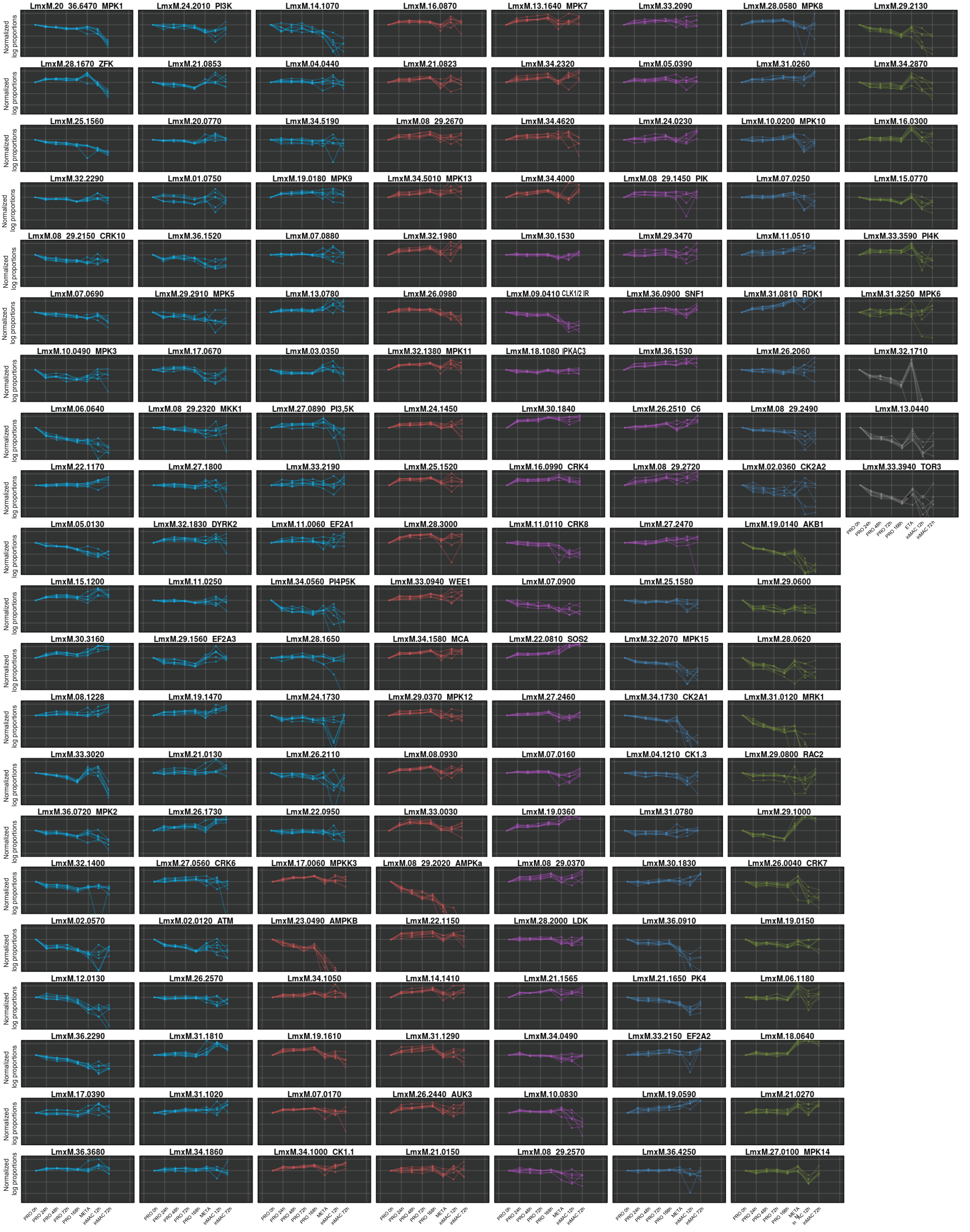

**Supplementary Data 3e**

Heat maps for mouse footpad infection data

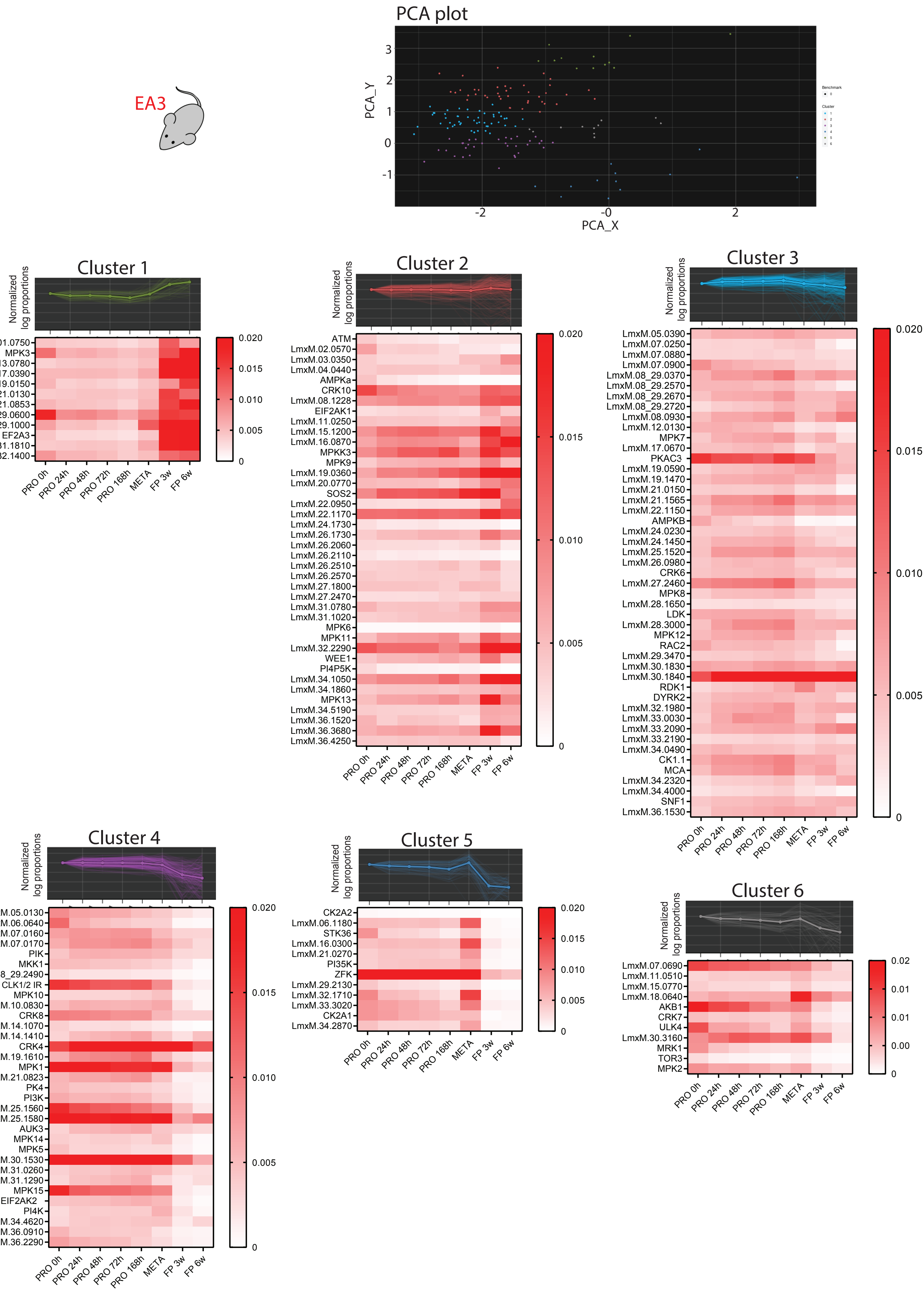

### Supplementary Data 3f

Individual plots of  mouse footpad infection data

EA3

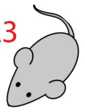

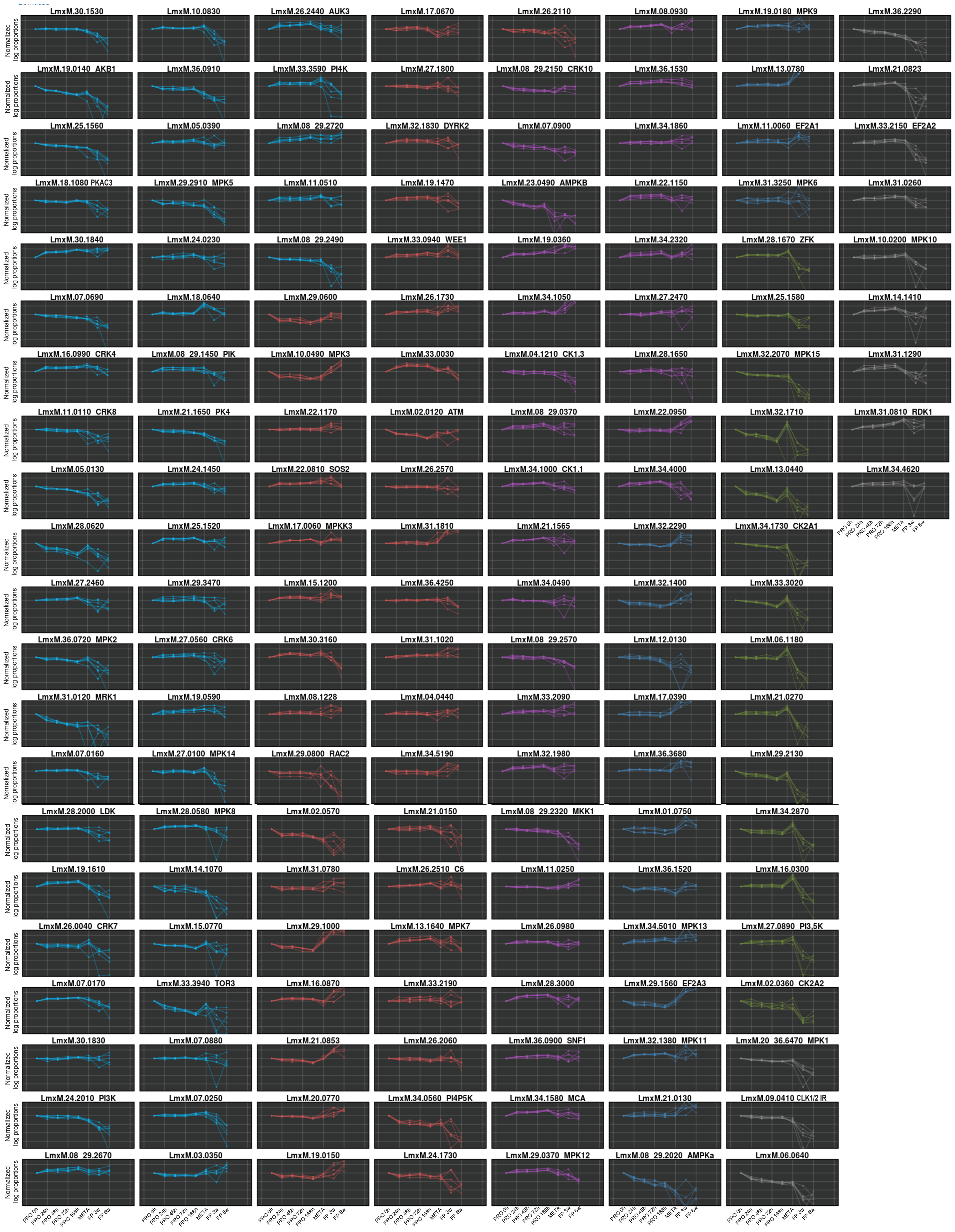
