## Supplementary equation 1 for "Systematic functional analysis of *Leishmania* protein kinases identifies regulators of differentiation or survival"

### Pseudocode for Projection Pursuit Clustering

Baker et al.

August 19, 2020

**Abstract**

The following pseudocode represents an informal sketch of the algorithm used to allocate the mutants to clusters.

1. **Input data.**

(1.1) The algorithm is provided with PCR counts $c_{i,j,t}$ for each mutant (indexed by $i = 1, \ldots, N$), each experiment replicate ($j = 1, \ldots, R$) and each experiment observation point ($t = 1, \ldots, T$). Use $c_{i,t} = \sum_j c_{i,t}$ to denote the sum of counts over replicates. The number of clusters $K$ for aggregating the mutants is also specified.

2. **Initialization.**

(2.1) Compute expectations and variances for log-proportions for each mutant, replicate and time point,

$$\begin{aligned} \mathbb{E}(\log p_{i,j,t}) &\leftarrow \gamma(c_{i,j,t}) - \gamma(c_{i,t}), \\ \mathrm{var}(\log p_{i,j,t}) &\leftarrow \psi(c_{i,j,t}) - \psi(c_{i,t}), \quad i = 1, \ldots, N,\ j = 1, \ldots, R,\ t = 1, \ldots, T \end{aligned}$$

where $\gamma$ and $\psi$ are the gamma and digamma functions, respectively. These are commonly available in scientific and statistical software packages.

(2.2) Aggregate expectations by taking a precision-weighted mean over replicates for each mutant and time point

$$\mathbb{E}(\log p_{i,t}) \leftarrow \sum_j \hat{w}_{i,j,t} \mathbb{E}(\log p_{i,j,t}) \quad i = 1, \ldots, N,\ t = 1, \ldots, T$$

where the weights

$$\begin{aligned} \hat{w}_{i,j,t} &\leftarrow w_{i,j,t} / \sum_j w_{i,j,t}, \\ w_{i,j,t} &\leftarrow 1/\mathrm{var}(\log p_{i,j,t}), \quad i = 1, \ldots, N,\ j = 1, \ldots, R,\ t = 1, \ldots, T \end{aligned}$$

are normalized reciprocals of the variances.

(2.3) Compute first differences for the series so that the clustering algorithm distinguishes between trajectories on the basis of just their shape rather than their shape and displacement,

$$d_{i,t} \leftarrow \mathbb{E}(\log p_{i,t}) - \mathbb{E}(\log p_{i,t-1}) \quad i = 1, \ldots, N,\ t = 2, \ldots, T.$$

(2.4) Encode the differences for each mutant as vectors $d_i$ in a $(T-1)$-dimensional Euclidean space.

(2.5) Pick arbitrary a pair of orthonormal vectors in $(T-1)$ dimensions. Use these to form the columns of a matrix $U$.

3. **Iterative clustering procedure.**

(3.1) Project the difference vectors onto the two-dimensional subspace spanned by the columns of a matrix $U$,

$$x_i \leftarrow U^T d_i \quad i = 1, \ldots, N.$$

(3.2) Propose a clustering solution for the projected coordinates. We have used a simple hierarchical clustering algorithm that sequentially aggregates groups of mutants that are closest together as measured according to Ward's criterion . Aggregation is halted when the $K$ clusters remain.

(3.3) If the clustering solution has changed since the previous iteration (or if this is the first iteration) proceed to the next step. Else, break the cycle of iterations and proceed to step 4.

(3.4) Compute cluster counts, means and variances for the mutants in the high-dimensional difference space.

$$\begin{aligned} n_k \leftarrow & \sum_{\text{mutant } i \in \text{cluster } k} 1, \\ m_k \leftarrow & n_k^{-1} \sum_{\text{mutant } i \in \text{cluster } k} d_i, \\ S_k \leftarrow & n_k^{-1} \sum_{\text{mutant } i \in \text{cluster } k} (d_i - m_k)(d_i - m_k)^T, \quad k = 1, \dots, K. \end{aligned}$$

(3.5) Compute mean of cluster means

$$m \leftarrow N^{-1} \sum_k n_k m_k.$$

(3.6) Compute within-cluster and between-cluster variance matrices

$$W \leftarrow N^{-1} \sum_k n_k S_k, \qquad B \leftarrow N^{-1} \sum_k n_k (m_k - m)(m_k - m)^T.$$

(3.7) Redefine the projection space as that spanned by the first two eigenvectors of the variance ratio matrix,

$$U \leftarrow Eigen(W^{-1}B, 2).$$

This step identifies the projection that best separates the clusters in the high-dimensional space.

(3.8) Return to step (3.1).

4. **Terminate and return output.**

(4.1) Output the labels from the last clustering step and the projected coordinates.
